# IRF5 mediates adaptive immunity via altered glutamine metabolism, mTORC1 signaling and post-transcriptional regulation following T cell receptor activation

**DOI:** 10.1101/2024.08.26.609422

**Authors:** Zarina Brune, Ailing Lu, Matthew Moss, Leianna Brune, Amanda Huang, Bharati Matta, Betsy J Barnes

## Abstract

Although dynamic alterations in transcriptional, translational, and metabolic programs have been described in T cells, the factors and pathways guiding these molecular shifts are poorly understood, with recent studies revealing a disassociation between transcriptional responses and protein expression following T cell receptor (TCR) stimulation. Previous studies identified interferon regulatory factor 5 (IRF5) in the transcriptional regulation of cytokines, chemotactic molecules and T effector transcription factors following TCR signaling. In this study, we identified T cell intrinsic IRF5 regulation of mTORC1 activity as a key modulator of CD40L protein expression. We further demonstrated a global shift in T cell metabolism, with alterations in glutamine metabolism accompanied by shifts in T cell populations at the single cell level due to loss of *Irf5*. T cell conditional *Irf5* knockout mice in a murine model of experimental autoimmune encephalomyelitis (EAE) demonstrated protection from clinical disease with conserved defects in mTORC1 activity and glutamine regulation. Together, these findings expand our mechanistic understanding of IRF5 as an intrinsic regulator of T effector function(s) and support the therapeutic targeting of IRF5 in multiple sclerosis.

**Sentence Summary:** Findings provide new insight into the mechanisms by which T cell intrinsic IRF5 regulates the adaptive immune response via modulation of mTORC1 signaling, glutamine metabolism, and protein translation.

## INTRODUCTION

The transcription factor interferon (IFN) regulatory factor 5 (IRF5) has been characterized as a regulator of type I IFNs and a key mediator of proinflammatory cytokine expression in response to Toll-like receptor (TLR) signaling. Dysregulation of IRF5 has been linked to infection, autoimmune disease, cancer, metabolic diseases, and neuropathic pain (*1–3*). Initial mechanistic studies on IRF5 were performed in myeloid and B cells; however, more recent studies support a role for IRF5 in CD4 T cells (*3, 4*). Loss of IRF5 expression in CD4 T cells drives Th2 skewing in the context of Th1 and Th17 polarizing conditions, alters inflammatory cytokine production, regulates apoptosis, and inhibits T cell proliferation and chemotaxis (*4*). Many of these IRF5-mediated functions have been reported downstream of IRF5 transcriptional regulation, yet the role of IRF5 in T cell metabolism and translational reprogramming, global shifts of which are required to assume effector function(s), has yet to be examined. Indeed, a lack of correlation between transcriptional responses and protein expression following T cell activation has been documented, with regulation of the T cell proteome occurring far more rapidly than can be explained by *de novo* transcription (*5*). Conversely, rapid metabolic changes in response to T cell stimulation correlate with the kinetics of T activation. One of the key mediators of both normal and pathologic metabolic and translational responses is the mammalian target of rapamycin (mTOR) signaling complex (*6*).

Signaling from mTOR kinase complexes, mTORC1 and mTORC2, regulates T cell effector programs, migration, proliferation, and survival. Similar to previous observations in *Irf5^−/−^* mice showing skewing towards IL4 secreting Th2 cells, reductions in Th1 and Th17 cells, and increased Treg generation (*4*), reduced mTORC1 signaling is correlated with increased regulatory T cells (Tregs) and a reduction in Th1 and Th17 generation (*7*). In addition, mTORC1 is a key post-transcriptional and translational regulator. Prior studies revealed that naïve CD4 T cells have reserves of ribosomal machinery which, upon TCR stimulation, are activated by mTORC1 to support increased protein synthesis (*8, 9*). Not only has repression of protein synthesis and mTORC1 signaling been linked to T cell quiescence, but effector immune signaling pathways are also regulated by these translational molecules (*10, 11*). Further studies elucidating how manipulation of ribosomal machinery regulates T cell function remain to be completed.

The emerging importance of metabolic pathways, mTOR signaling and translational regulation in T cells has inspired a new generation of drug development and target discovery. *In vitro* studies using 2-deoxyglucose and metformin to reprogram reactive T cells from patients with systemic lupus erythematosus (SLE) were met with success (*12*). In the murine model of multiple sclerosis (MS), experimental autoimmune encephalomyelitis (EAE), mTORC1 inhibition with rapamycin protected mice from classical disease development and reversed symptom onset (*7, 13, 14*). In addition, manipulation of the Th17-Treg axis through alterations in glutamine metabolism is protective in EAE (*15*). Through a combination of targeted inhibition studies, scRNAseq, flow cytometry, and unbiased metabolomics, we identify IRF5 in the regulation of CD4 T cell metabolism, mTOR signaling and protein translation, and demonstrate a disease modifying role for T cell intrinsic IRF5 in EAE.

## RESULTS

### IRF5 regulates T cell support of B cell adaptive responses

Previous studies by our lab and others revealed defects in plasmablast (PB) generation and IgG2a/c production in mouse and human *Irf5*-deficient B cells by *in vitro* culture and *in vivo* immunization (*16, 17*). Although these defects were attributed to B cell intrinsic IRF5 function, more recent studies suggest a role for T cell intrinsic IRF5 in the regulation of B cell adaptive immune responses (*4*) We thus utilized the T cell-dependent antigen NP-conjugated chicken gamma globulin (CGG) emulsified in Complete Freund’s Adjuvant (CFA) to assess PB differentiation in 8-10 weeks-old *Irf5^+/+^* (WT) and *Irf5^−/−^* (KO) littermate mice. 7 days following immunization, WT and KO spleens were harvested for immunohistochemistry (IHC) and flow cytometric analysis. We detected a significant reduction in B220^+^ staining in KO spleens with no difference in spleen size (**Fig. 1A, B, Supp. 1A, B**). Flow cytometric analysis of KO splenocytes showed reductions in T follicular helper (Tfh) cells (BCL6^+^CXCR5^+^) and significantly increased regulatory T cells (Tregs) (CD25^+^FoxP3^+^) (**Fig. 1C-E**). Further examination in non-immunized mice revealed significant reductions in splenic Tfh cells in KO compared to WT mice (**Supp. 1C**). Prior studies showed KO T cells to have decreased activation and proliferation following anti-CD3/CD28 stimulation. We confirmed these findings in purified CD4 T cells (**Supp. 1D-G**). Additionally, although KO mice had no significant reductions in NP-specific PBs (CD19^+^IgD^low^CD138^+^), there were significant reductions in IgG2a production (**Fig. 1F-H**). Next, we examined by *in vitro* co-culture assay if KO CD4 T cells were sufficient to replicate the observed B cell defects in the NP-CGG CFA immunization model. B cells (CD45^+^B220^+^) and CD4 T cells (CD45^+^CD4^+^) were sorted from WT and KO splenocytes and co-cultured for 4 days in the presence of anti-IgM, CpG-B and anti-CD3/CD28 Dynabeads. Of interest, significant reductions in CD19^+^IgD^low^CD138^+^ PB generation were observed in KO B cell:KO T cell, WT B cell:KO T cell and KO B cell:WT T cell cocultures as compared to WT B cell:WT T cell cultures with only slight differences in IgG2a production (**Fig. 1I, J, Supp. 1H, I**). Together, these data support a role for CD4 T cell intrinsic *Irf5* in B cell adaptive immunity.

**Fig. 1.**
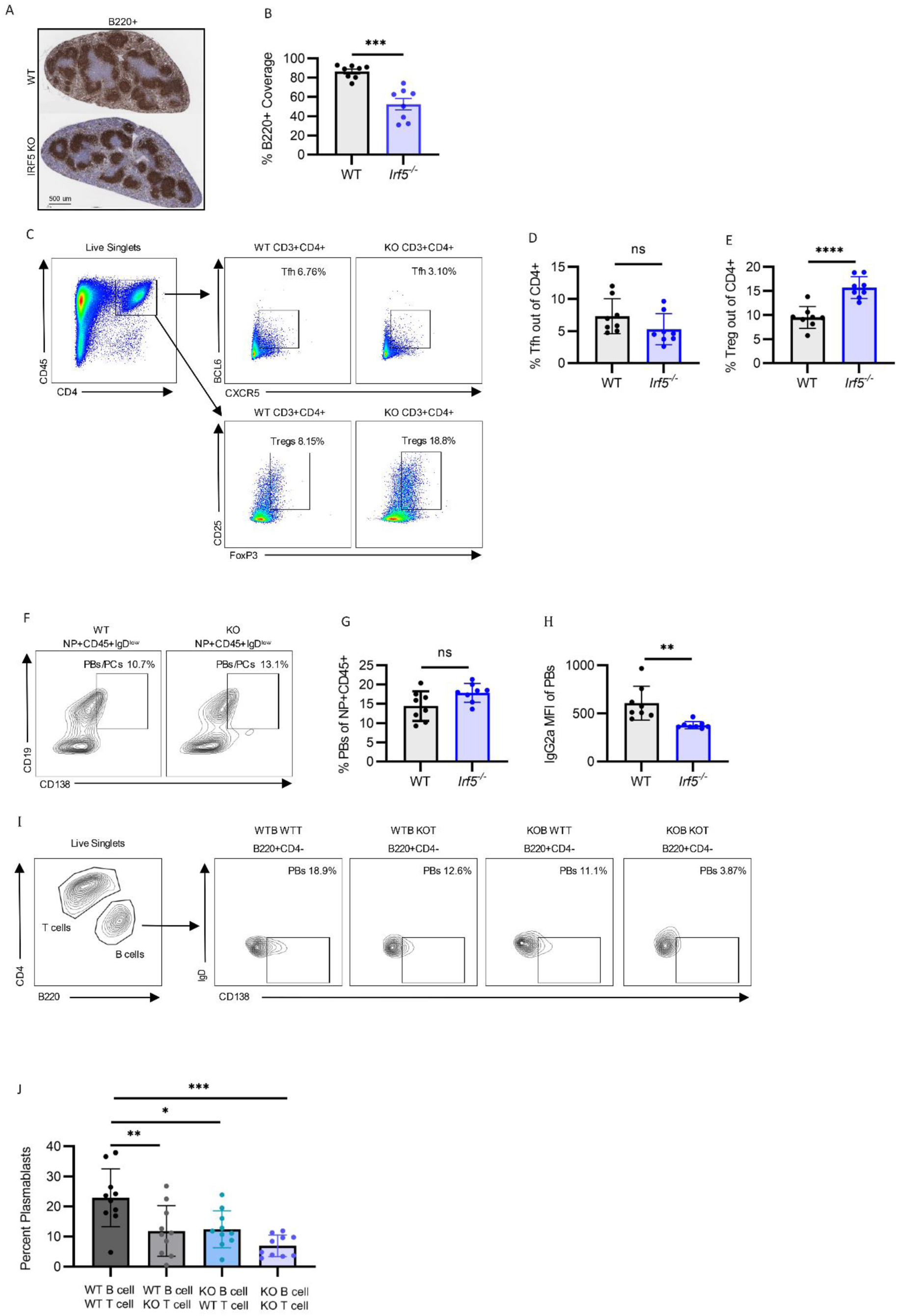
T cell intrinsic IRF5 regulates T cell dependent B cell maturation. Mice were immunized intraperitoneally with 50 ug NP-CGG/CFA. Spleens harvested 7 days following immunization. (**A**) Representative histology and (**B**) quantification of B220^+^ staining from WT and *Irf5^−/−^* spleens; scale bar represents 500µm. (**C**) Representative flow gating strategies for Tfh (CD3^+^CD4^+^BCL6^+^CXCR5^+^) and Tregs (CD3^+^CD4^+^CD25^+^FoxP3^+^) and their respective quantification (**D, E**). (**F**) Representative flow cytometric gating strategy for PBs (CD19^+^IgD^low^CD138^+^). (**G**) Quantification of NP specific PBs (NP^+^CD45^+^CD19^+^CD138^+^IgD^low^) and (**H**) IgG2a production from (**G**). *P* values calculated by two-way unpaired *t* test. (**I**) Representative flow gating strategy and (**J**) summary graphs of B220^+^CD138^+^IgD^low^ plasmablast generation following 4-day *in vitro* coculture. *P* value calculated by one-way analysis of variance (ANOVA) with Tukey’s post-hoc test for multiple comparisons. Bar graphs show means +/- SEMs. **p < 0.01, ***p < 0.001, ****p <0.0001, ns = not significant. Data pooled from two to three independent experiments with each point representing an independent biological replicate.

### scRNAseq reveals alterations in metabolism, ribosome biogenesis and identifies novel IRF5 transcriptional targets

Current dogma in the field suggests that loss of *Irf5* inhibits Th1 and Th17 effector subsets, enhances Treg differentiation and Th2 responses, and alters Tfh function through transcriptional regulation (*3, 4, 18, 19*). To gain further insight into the cellular pathways by which IRF5 regulates T cell differentiation/function, we performed scRNAseq on 5,000-7,000 TCR-stimulated WT and KO CD4 T cells. Using Seurat, we identified seven distinct T cell clusters (**Fig. 2A**). The subsets were classified as follows: activated naïve (*Nr4a1, Nfkbid, Cd69, Egr3*), naïve (*Sell, Lef1, Klf2, Ifit3*), memory-like (T_MEM_) (*Spry1, Fabp5*, *Ccr7*) (*20, 21*), IFN Enriched (*Gbp2, Gbp5*, *Trat1, Nme1, Nme2)* (*22–25*), Tfh (*Tigit, Il21, Cxcr5, Pdcd1*), Th Complex (*Rora, Ccr5, Ccr2, Serpin6b6)* (*26*), and Treg (*FoxP3*, *Ikzf2, Il2ra, Il2rb*) (**Fig. 2A, B**). Further subclustering revealed two distinct populations comprising both Treg (termed Treg 0 and Treg 1) and Tfh clusters (identified as Tfh 0 and Tfh 1) (**Supp. Table 1**).

**Fig. 2.**
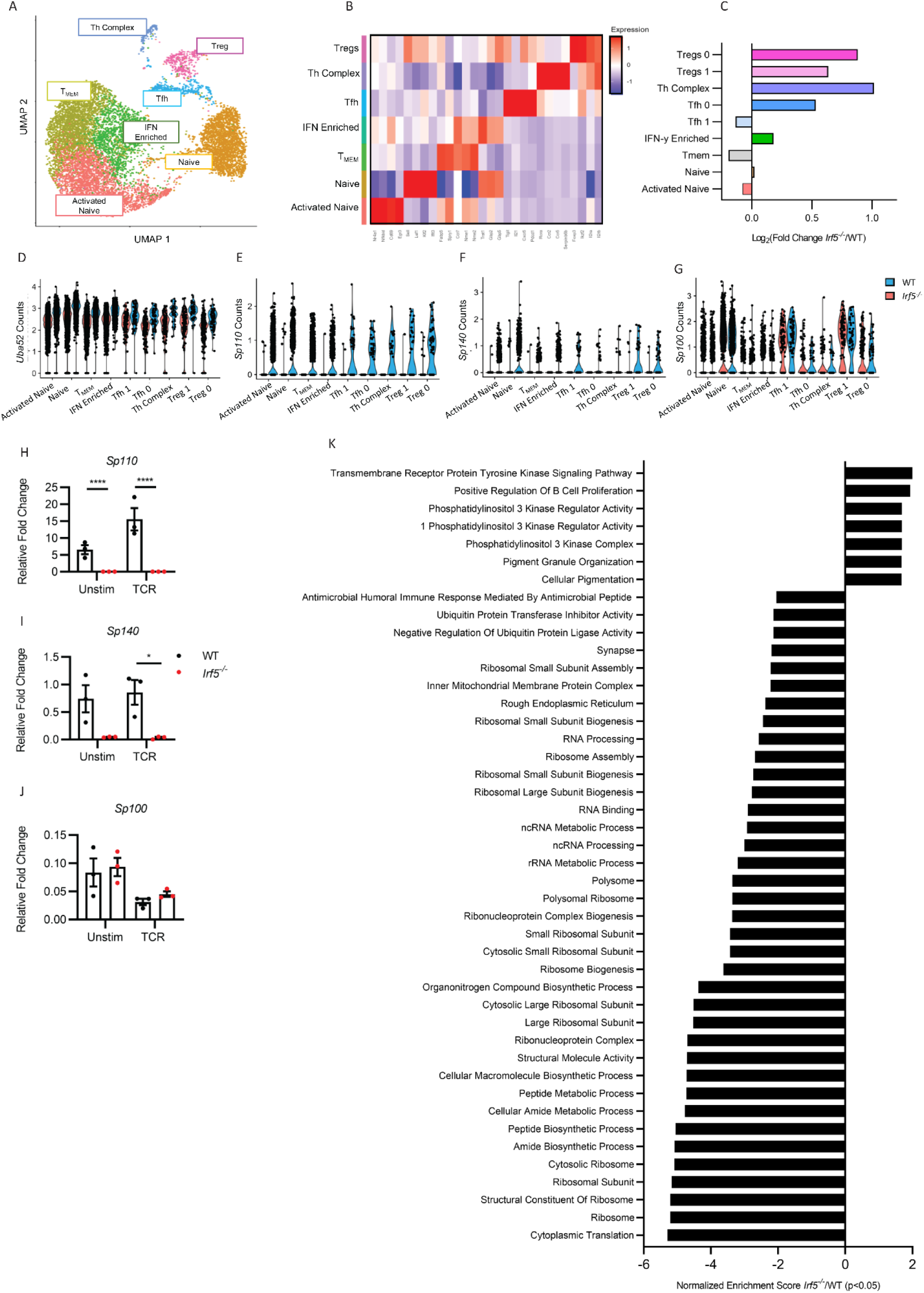
Loss of IRF5 alters CD4 single cell transcriptional landscape. (A) Uniform Manifold Approximation and Projection (UMAP) of 5000 - 7000 CD4^+^ single cells. Each dot corresponds to a single cell, colored according to cell type. Representative graph of two WT and two *Irf5^−/−^* biological replicates, with one male and one female representing each genotype. (B) Heatmap of genes used to identify CD4 T cell clusters. Data are colored according to expression levels. Legend is labeled in log scale. (**C**) Log2 fold change enrichment values of *Irf5^−/−^* relative to WT T cell clusters. (**D**-**G**) Violin plots showing the transcript expression of (**D**) *Uba52*, (**E**) *Sp110*, (**F**) *Sp140* and (**G**) *Sp100* in WT and *Irf5^−/−^* CD4 T cells in indicated T cell cluster. (**H**-**J**) Sorted CD3^+^CD4^+^ T cells from WT and *Irf5^−/−^* splenocytes were activated for 6-hours *in vitro* with anti-CD3/CD28 (TCR). (**H**) *Sp110*, (**I**) *Sp140* and (**J**) *Sp100* transcript expression normalized to *β-Actin.* Three biological replicates, representative of two independent experiments. *P* value calculated by two-way unpaired *t* tests. (**K**) Preranked gene set enrichment analyses using C5 gene ontology (GO) gene sets. Top gene ontology (GO) terms sorted by Normalized Enrichment Score (NES). Padj < 0.05. \**P* < 0.05, \*\*\*\**P* < 0.0001. Each point represents an independent biological replicate.

Pseudotime trajectory analysis was performed using Slingshot (*27*) to examine the relationships between each subset. Four trajectory lineages were identified in WT and KO CD4 T cells (**Supp. 2A**). The Tfh and Treg, Th complex subset and IFN enriched, T_MEM_ and activated naïve T cell clusters were predicted to exist within independent lineages. Paired with cluster enrichment analysis (**Fig. 2C**), the pseudotime trajectory findings support that loss of *Irf5* skews T cells toward the specific Tfh/Treg lineage and away from activated naïve and T_MEM_ trajectories. Indeed, further examination of subset enrichment in KO compared to WT CD4 T cells showed KO mice to have reduced activated naïve cells, enrichment in both Treg 0 and Treg 1 and a decrease in T_MEM_ populations, supporting previously described flow cytometric CD4 T cell subset profiling by our lab and others (*4*). There was also a distinct increase in the Tfh 0 and a slight decrease in Tfh 1 KO T cell populations. The Th complex subset, named thus due to increased expression of genes encoding migratory receptors, inflammatory cytokines, inhibitory molecules, and transcription factors, was also highly enriched in KO CD4 T cells (**Fig. 2C, Supp. 2B**). Of interest, a significant proportion of genes dysregulated in the KO Th complex subset comprised of ribosomal transcripts and alternative splicing machinery. In addition, KO Th complex cells had downregulation of proinflammatory and metabolic genes, including *Il18r1*, *Tomm5*, *Nkg7*, *Mif*, *Klrk1* and *Irf8* (**Supp. 2B**) (*28–32*). Unlike the Th complex subset, despite the clear shifts in the Tfh and Treg subcluster distribution, few transcripts within each of these clusters demonstrated significant differences with loss of *Irf5* (**Fig. 2C, Supp. 2C, D**). DEG analysis identified only three genes with clear dysregulation across all KO T cell clusters: *Uba52*, *Speckled protein (Sp) 110* (*Sp110*) and *Speckled protein 140* (*Sp140*) (**Fig. 2D-F**). Analysis of the third SP family member, *Sp100*, revealed no change in expression (**Fig. 2G**) (*33*). The specific and dramatic reductions in *Sp110* and *Sp140*, but not *Sp100*, were independently confirmed by *qPCR* in purified WT and KO CD4 T cells (**Fig. 2H-J)**. Of note, previous studies in myeloid cells demonstrated that IRF4 can bind to and regulate similar target genes as IRF5 (*34*). To assess specificity of IRF5 in *Sp* regulation, we examined *Sp100*, *Sp110* and *Sp140* expression in *Irf4^−/−^* T cells. No significant differences in either *Sp100*, *Sp110* or *Sp140* transcript expression were detected (**Supp. 2E-G**).

*SP110* and *SP140* are genes of interest in both inflammatory and autoimmune diseases. Mutations in these factors are associated with Crohn’s disease, chronic lymphocytic leukemia, and MS, while hyperactivation of SP110 and SP140 is associated with SLE and MS (*33*). Given the striking reduction in *Sp110* and S*p140* expression in *Irf5^−/−^* T cells and the implications of elevated IRF5 expression and hyperactivation as a driver of SLE (*35, 36*), we examined *Sp110* and *Sp140* expression in a published RNAseq dataset (GSE149050) from healthy donors and SLE patients with differing IFN levels (*37*). Interestingly, we found increased *SP110* expression in PBMCs from IFN high expressing SLE patients compared to both IFN low SLE patients and healthy controls (HC). Similar trends were observed for *SP140* (**Supp. 2H, I)**. Previous studies from our lab and others demonstrated that SLE is mediated in part by the aberrant production of autoantibodies regulated by B cell intrinsic IRF5 (*16, 38*). Notably, one of the pathologies associated with speckled protein inactivating mutations is inhibition of B cell antibody production (*39*). Analysis of *Sp110* expression in KO B cells revealed a similar reduction in transcript expression as KO T cells (**Supp. 2J**).

To further assess functional differences between WT and KO T cells, we performed Gene Set Enrichment Analysis (GSEA) using the C5 Gene Ontology (GO) sub-collection. We found significant downregulation in gene sets for Cytoplasmic Translation, Ribosomal Synthesis, Ribosome Metabolism, Ribosome Assembly and Function, RNA Processing, Mitochondrial Depolarization and Mitochondrial structure in KO CD4 T cells (**Fig. 2K**). GSEA using C2 KEGG also revealed significant downregulation of Ribosomes, Oxidative Phosphorylation and Systemic Lupus Erythematosus, supporting conserved regulatory roles for IRF5 in oxidative phosphorylation, mitochondrial function, and ribosome regulation (**Supp. 2K, Supp. Table 2)**. Given these striking reductions in cytoplasmic translation and ribosomal synthesis, we re-examined a previously unpublished study from our lab identifying IRF5-protein interactions in Ramos B cells by immunoprecipitation (anti-IgG control or anti-IRF5 antibodies) and mass spectrometry analysis (**Supp. Table 3**). Among the most significantly enriched for IRF5 interacting partners, as compared to Ig control, were the 60S and 40S ribosomal proteins (RPLs), and translation initiating factors EIF4A2 and EIF6. Despite these studies being performed in B cells, IRF5 interaction with proteins involved in ribosomal assembly and biogenesis (RPLs, EIF6) and translation (EIF4A2) imply a conserved role for cytoplasmic IRF5 in the regulation of the translational apparatus, findings supported by prior independent studies (*40*).

### *Irf5^−/−^* mice are protected from Experimental Autoimmune Encephalomyelitis

Although inhibition or loss of IRF5 is protective in autoimmune and inflammatory diseases (*4, 35, 41–43*), apart from *Leishmania donovani* and inflammatory bowel disease (IBD) models, the contribution(s) of IRF5 to T cell-mediated disease pathogenesis has remained largely unexplored. Using scRNAseq, we found downregulation of proinflammatory factors previously implicated in EAE disease pathogenesis including *Il18r1*, *Irf8*, and *Mif*, and increased expression of *Rora* in the KO Th complex subset (*28, 32, 44, 45*). In addition, we detected a significant reduction in *Sp110* and *Sp140* expression, gene loci associated with risk of MS (*33*). Thus, we next expanded our studies to examine if loss of *Irf5* impacts the clinical progression of experimental autoimmune encephalomyelitis (EAE) using MOG_35-55_/PTX injections (*46*). Using a common clinical rating scale (*46*), we found that KO mice had both decreased and delayed incidence and onset of disease as well as significantly attenuated disease progression compared to WT littermate mice, indicating a protective role for loss of *Irf5* in T cell-mediated EAE (**Fig. 3A, B**).

**Fig. 3.**
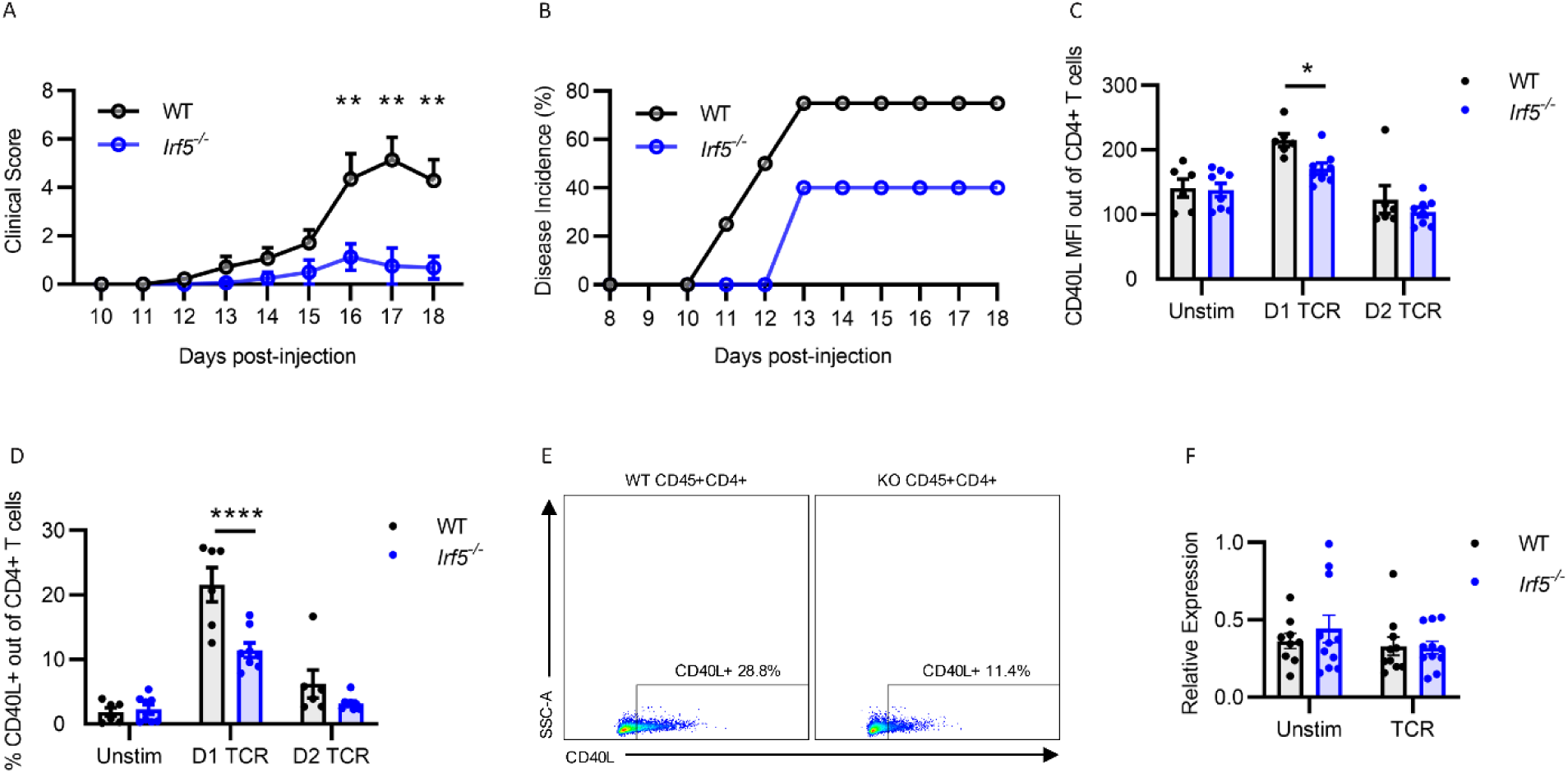
*Irf5* KO mice are protected from Experimental Autoimmune Encephalomyelitis. (**A**) Composite means +/- SEM of clinical scores. Minimum of 12 biological replicates per experiment. Data pooled from two independent experiments. *P* value calculated by 2-way ANOVA with Holm-Sidak correction for multiple comparisons. (**B**) Disease incidence. Minimum of four biological replicates per genotype. (**C, D**) WT and *Irf5^−/−^* total splenocytes were sorted for CD3^+^CD4^+^ T cells, then activated for 24- or 48-hours *in vitro* with anti-CD3/CD28 Dynabeads (TCR). Live CD3^+^CD4^+^ T cells were gated for flow cytometric marker analysis. (**E)** Representative CD40L flow cytometry gating in WT and KO CD4 T cells. *P* value calculated by 2-way ANOVA with Holm-Sidak correction for multiple comparisons. (**F**) *Cd40l* transcript expression normalized to *β-Actin* following 6-hours *in vitro* anti-CD3/CD28 (TCR) stimulation. Data pooled from two to three independent experiments, with each point representing an independent biological replicate. *P* value calculated by two-way unpaired *t* test. \**P* < 0.05, \*\**P* < 0.01. Bar graphs show means +/- SEMs.

One of the T cell-mediated adaptive immune signaling pathways dysregulated in MS involves the expression and binding of T cell CD40L to its receptor, CD40, on B cells (*47, 48*). Given the observed reductions in PB generation by coculture of WT B cells with KO T cells and the stark protection of *Irf5* KO mice from T cell-mediated EAE disease onset and progression, we next examined if loss of *Irf5* dysregulated the CD40L/CD40 pathway. Following stimulation of WT and KO total splenocytes with either anti-CD3/CD28 Dynabeads or anti-IgM (B cell receptor stimulation), CD40L and CD40 expression was quantified on Live CD3^+^CD4^+^ T cells and Live CD45^+^CD19^+^B220^+^ B cells, respectively. Anti-CD3/CD28-stimulated KO T cells demonstrated a significant decrease in both CD40L surface expression, as quantified by mean fluorescence intensity (MFI), and CD40L expressing CD3^+^CD4^+^ T cell populations 24 hours following stimulation (**Fig. 3C-E**). There were slight, albeit significant reductions in the percentage of CD40^+^ B cells and CD40 MFI on unstimulated and anti-IgM stimulated KO B cells (**Supp. 3A, B)**. Examination of basal CD40L expression revealed a slight but significant increase in the percentage of CD40L expressing KO T cells, but no significant difference in MFI (**Supp. 3C, D**). *Cd40l* transcript expression was not significantly detected via scRNAseq, thus we evaluated *Cd40l* transcript expression from sorted CD3^+^CD4^+^ T cells (purity > 95%) from WT and KO mice following 6-hour *in vitro* TCR stimulation. There was no significant difference in expression between genotypes (**Fig. 3F**). This was unsurprising as previous studies have indicated that *Cd40l* expression is extensively regulated post-transcriptionally (*49, 50*). Notably, flow cytometric examination revealed, like that seen in NP-CGG CFA immunized KO mice, a significant decrease in Tfh cells in KO spleens following EAE induction (**Supp. 3E, F**). Together, aberrant CD40L protein expression on KO CD4 T cells and reductions in Tfh cells supports a role for IRF5 in T cell support of the adaptive immune response and in post-transcriptional or translational regulation of CD40L following TCR signaling.

### IRF5 regulates protein translation in T cells via mTORC1

Given our findings, we next sought to examine the function of one of the most highly conserved mediators of translation, metabolism, and post-transcriptional regulation, whose aberrant function also has been implicated in EAE pathogenesis, the mammalian target of rapamycin 1 (mTORC1) signaling complex. mTORC1 is a key regulator of T cell proliferation, differentiation, and effector function (*51–53*). Despite current and prior studies demonstrating KO T cell defects in activation and proliferation, the regulatory mechanisms underlying these dysfunctions have yet to be fully elucidated (*4, 17, 42, 43*). Considering our findings that suggest a role for IRF5 in translational regulation, we next determined if mTORC1 activity was altered in KO T cells by examining total ribosomal S6 protein (RPS6) and phosphorylated RPS6 (phosphoRPS6) levels, both of which are canonical downstream effectors of mTORC1 signaling. Flow cytometric analysis revealed a significant defect in the phosphorylation of RPS6 in TCR stimulated KO T cells, indicating decreased mTORC1 activity (**Figs. 4A, B**). No significant difference in total RPS6 expression was detected (**Supp. 4A**). Examination of phospho(Thr389)-p70 S6 kinase (P70S6K), the kinase responsible for RPS6 phosphorylation, showed decreased phosphorylation, albeit insignificant (**Supp. 4B**). Additional studies revealed similar total mTOR protein expression in KO and WT CD4 T cells basally and following TCR stimulation, further supporting that IRF5 regulates mTORC1 activity rather than mTOR expression (**Supp. 4C**).

**Fig. 4.**
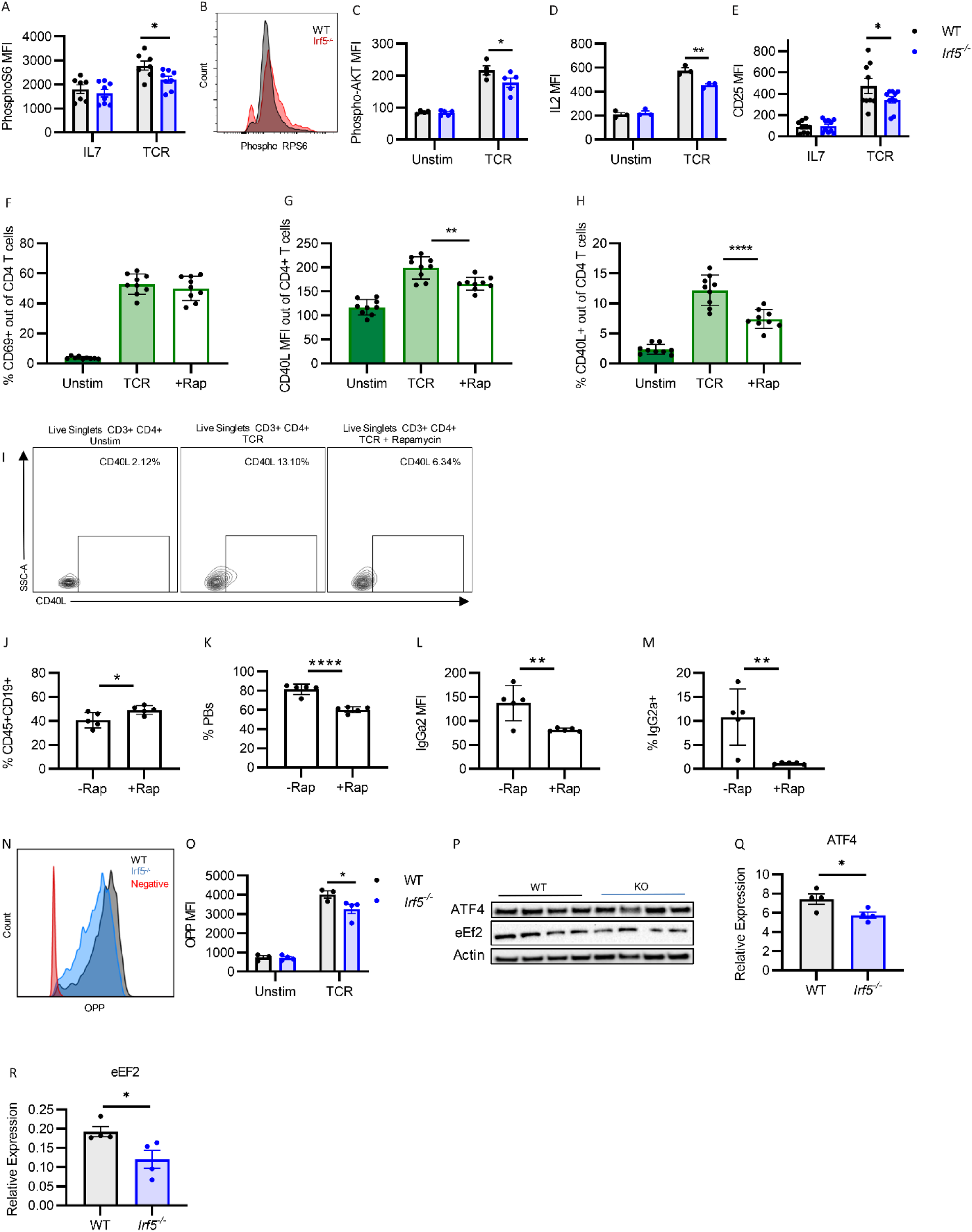
IRF5 regulates protein translation, mTORC1 and Akt signaling. (**A**) Phosphorylated RPS6 quantification in CD3^+^CD4^+^ T cells sorted from WT and *Irf5^−/−^*splenocytes following 24-hours *in vitro* stimulation with anti-CD3/CD28 (TCR) or without (IL7). Data pooled from three independent experiments. *P* value calculated by 2-way ANOVA with Holm-Sidak correction for multiple comparisons. (**B**) Representative histogram of phosphorylated RPS6 quantification from (**A**). (**C**-**E**) Summary quantification of (**C**) phospho-AKT, (**D**) IL2 and (**E**) CD25 expression in WT and *Irf5^−/−^*CD3^+^CD4^+^ T cells gated from total splenocytes following 24-hours *in vitro* stimulation with anti-CD3/CD28 (TCR) or without (Unstim). Three to five biological replicates, representative of two independent experiments. *P* value calculated by 2-way ANOVA with Holm-Sidak correction for multiple comparisons. (**F**-**H**) Summary quantification of (**F**) CD69, (**G**) CD40L mean fluorescence intensity (MFI) and (**H**) CD40L% expression in Live CD3^+^CD4^+^ T cells gated from total splenocytes following 24-hours *in vitro* stimulation in the presence (TCR + Rap) or absence (TCR) of rapamycin. Data pooled from three independent experiments. *P* value calculated by Ordinary 1-way ANOVA with Tukey correction for multiple comparisons. (**I**) Representative gating for CD40L expression in CD3^+^CD4^+^ gated on Live Singlets in the absence (TCR) or presence (+Rap) of rapamycin. (**J**-**M**) Summary quantification of (**J**) CD19^+^ B cells, (**K**) plasma blasts (PBs) (CD45 ^+^CD19^+^CD138^+^IgD^low^) and (**L, M**) IgG2a production. Five biological replicates. *P* value calculated by two-way unpaired *t* tests. (**N**) CD3^+^CD4^+^ T cells gated from total splenocytes were activated for 24-hours *in vitro* with anti-CD3/CD28 (TCR) and puromycin incorporation measured via O-propargyl-puromycin assay. Representative histogram of puromycin incorporation is shown. (**O**) Quantification of puromycin incorporation from three to four biological replicates. *P* value calculated by 2-way ANOVA with Holm-Sidak correction for multiple comparisons. (**P**-**R**) Quantification and representative immunoblot of (**P, Q**) ATF4 and (**P, R**) eEF2 in sorted CD4 T cells following anti-CD3/CD28 stimulation. N = 4 biological replicates. *P* value calculated by two-way unpaired t tests. Bar graphs show means +/- SEMs. *p < 0.05, **p < 0.01, ***p < 0.001. Each point represents an independent biological replicate.

One of the main regulators of mTORC1 is the serine/threonine protein kinase, Akt (*54*). Akt activation promotes many of the pathways regulated by mTOR, including proliferation, survival, and metabolism (*55*). Analysis of Akt activity in KO CD4 T cells following 24-hour TCR stimulation revealed a significant reduction in Akt (Ser473) phosphorylation (**Fig. 4C**). This supports previous findings of Akt dysregulation in *Irf5*-deficient myeloid cells (*56*). Akt in T cells can be activated via signaling from IL2 (*57*). Following TCR stimulation, KO CD4 T cells had decreased expression of both IL2 and the IL2 receptor, CD25 (**Fig. 4D, E**). Prior studies have shown that, unlike in humans, IL2 does not regulate murine CD40L expression (*58*). Nonetheless, to confirm that alterations in IL2 expression and signaling through the IL2R were not responsible for reduced CD40L expression in KO CD4 T cells, total splenocytes were harvested and stimulated with anti-CD3/anti-CD28 in the presence or absence of recombinant IL2. Results showed no significant difference in CD40L expression (**Supp. 4D**). Of interest, treatment with the Akt inhibitor, MK2206, inhibited CD40L expression, indicating that Akt activity is required for CD40L expression (**Supp. 4E**). These results provide initial evidence that IRF5 regulates translation through mTORC1 signaling, whose activity is inhibited by reduced Akt activity.

Rapamycin is a small molecule inhibitor of mTORC1 that binds to cytosolic FKBP12 and inhibits mTOR S2448 phosphorylation and subsequent activation (*59*). Given the reduction in phosphoRPS6 in KO CD4 T cells, we next investigated if mTORC1 regulates the expression of CD40L. We stimulated WT CD4 T cells in the presence or absence of rapamycin for 24 hours, then analyzed T cell activation (CD4^+^CD69^+^) and CD40L expression. Although there was no change in the activation of CD4 T cells with rapamycin treatment (**Fig. 4F**), there was a significant decrease in CD40L expression that replicated levels seen in KO T cells (**Fig. 4G-I**, **Fig. 3C-E**). There was no difference in viability following rapamycin treatment (**Supp. 4F**). Next, to examine if T cell intrinsic mTORC1 inhibition was sufficient to inhibit the B cell adaptive response, we sorted and pretreated WT CD4 T cells with rapamycin (+Rap) or PBS (-Rap) then cocultured T cells with B cells as previously described. Quite strikingly, we observed a significant decrease in CD45^+^CD19^+^CD138^+^IgD^low^ PB generation and IgG2a production despite the relative increase in CD45^+^CD19^+^ B cells when B cells were cocultured with rapamycin-treated CD4 T cells. Taken together, these findings further support that mTORC1 signaling in CD4 T cells is required for T cell support of B cell adaptive immune responses **(Fig. 4J-M, Supp 4G)**.

Notably, the mTORC1 signaling axis is a key regulator of protein synthesis. With the previous findings demonstrating defects in T cell activation, proliferation, and CD40L expression in the context of mTORC1 dysfunction, we next examined rates of protein translation in unstimulated and TCR stimulated KO T cells by measuring incorporation of an alkyne analog using chemoselective fluorochrome ligation with the OPP assay. We found a significant decrease in the rate of protein translation in TCR stimulated KO T cells, indicating a role for IRF5 in the positive regulation of protein synthesis (**Fig. 4N, O**). Recent studies exploring mechanisms by which mTORC1 regulates translation have revealed ATF4 as a metabolic effector of mTORC1. When expressed, ATF4 promotes protein synthesis and stimulates the uptake of various amino acids. Immunoblot analysis of ATF4 expression in KO T cells following TCR stimulation revealed significantly reduced expression in KO T cells (**Fig. 4P, Q**). We also examined the expression of eukaryotic elongation factor 2 (eEF2), a global mediator of protein translation. Prior studies have demonstrated that NF-kB activating stimuli can activate eEF2 by repressing transcription of the inhibitory calcium/calmodulin dependent eEF2 kinase (eEF2K) (*60*). Other studies have shown that eEF2K inhibitory phosphorylation of eEF2 is regulated through the rapamycin-sensitive mTOR pathway, shown here to be dysregulated in KO T cells (*61, 62*). We thus examined eEF2 levels following TCR stimulation of purified CD4 T cells and detected a significant reduction in expression within KO T cells (**Fig. 4R, S**). Altogether, these findings support a role for IRF5 in the regulation of protein translation at the transcriptional and post-transcriptional level.

### Untargeted metabolomics reveals global metabolic shifts in *Irf5^−/−^* CD4 T cells

mTORC1 activity is extensively regulated by and responsive to changes in cellular energy levels and metabolites, changes that are mediated in part through ATF4 signaling (*63*). In T cells, metabolites modulate survival, proliferation, and effector fate decision and function (*64*). Results from scRNAseq analysis of KO T cells revealed a significant reduction in the enrichment of genes involved in oxidative phosphorylation (**Supp. 2K**). Prior studies in *Irf5^−/−^* macrophages have also reported reduced oxidative phosphorylation capacity (*56, 65, 66*). Although a role for IRF5 in T cell metabolic regulation has yet to be investigated, the observed alterations in KO T cell function and mTOR signaling provided compelling initial evidence to support this. As such, we performed unbiased LC-MS metabolomics analysis on unstimulated and TCR stimulated WT and KO purified naïve CD4 T cells (**Supp. Table 4**). T cell purity and activation was confirmed by flow cytometric analysis (**Supp. 5A)**. Basally and following TCR stimulation, the metabolic landscapes of WT and KO T cells had dramatic differences (**Supp 5B**). KO T cells significantly downregulating 21 metabolites relative to WT T cells (p < 0.05, FC > 1.5) (**Fig. 5A, B**). Following 24 hours of anti-CD3/CD28 stimulation, WT and KO T cells continued to demonstrate distinct metabolic profiles (**Fig. 5A-D**). Anti-CD3/CD28 stimulated KO T cells had 20 significantly upregulated and 4 significantly downregulated metabolites compared to unstimulated KO T cells (**Fig. 5C**), while anti-CD3/CD28 stimulated WT T cells had 28 significantly upregulated and 14 significantly downregulated metabolites compared to unstimulated WT T cells (**Fig. 5D**). Of those metabolites, TCR stimulated WT T cells revealed 10 unique ones that were upregulated while KO T cells had only 2 uniquely upregulated. Conversely, WT T cells showed no uniquely downregulated metabolites while KO T cells had 23 downregulated (p < 0.05, FC > 2.0) (**Fig. 5E, F, Supp. Table 5**). Subsequent metabolite enrichment pathway analysis demonstrated regulatory perturbations in some of the most highly enriched metabolite sets following TCR stimulation (**Supp. 5C**). Of particular interest was the significant enrichment in Malate-Aspartate Shuttle (MAS) metabolites: malate, aspartate, glutamate, and α-ketoglutaramate (α-KGM) in KO T cells (**Supp. 5C, Fig. 5G-K)**. We next quantified transcript levels of the cytoplasmic Glutamic-oxaloacetic transaminase 1 (*Got1*) and mitochondrial Glutamic-oxaloacetic transaminase 2 (*Got2*) MAS transaminases. Following anti-CD3/CD28 stimulation, both *Got1* and *Got2* transcript levels were increased in KO compared to WT T cells (**Supp. 5D, E**). Finally, to confirm that the increased levels of MAS metabolites and transaminases were reflective of increased utilization of the MAS in KO T cells, we treated WT and KO T cells with the MAS inhibitor aminooxyacetate acid (AOAA). Following 24 hours of AOAA treatment, anti-CD3/CD28 stimulated KO T cells had significantly increased rates of apoptosis compared to WT T cells (**Supp. 5F**). Together, these findings indicate global shifts in T cell metabolism with loss of *Irf5*, and increased reliance of KO T cells on the malate-aspartate shuttle following T cell receptor stimulation.

**Fig. 5.**
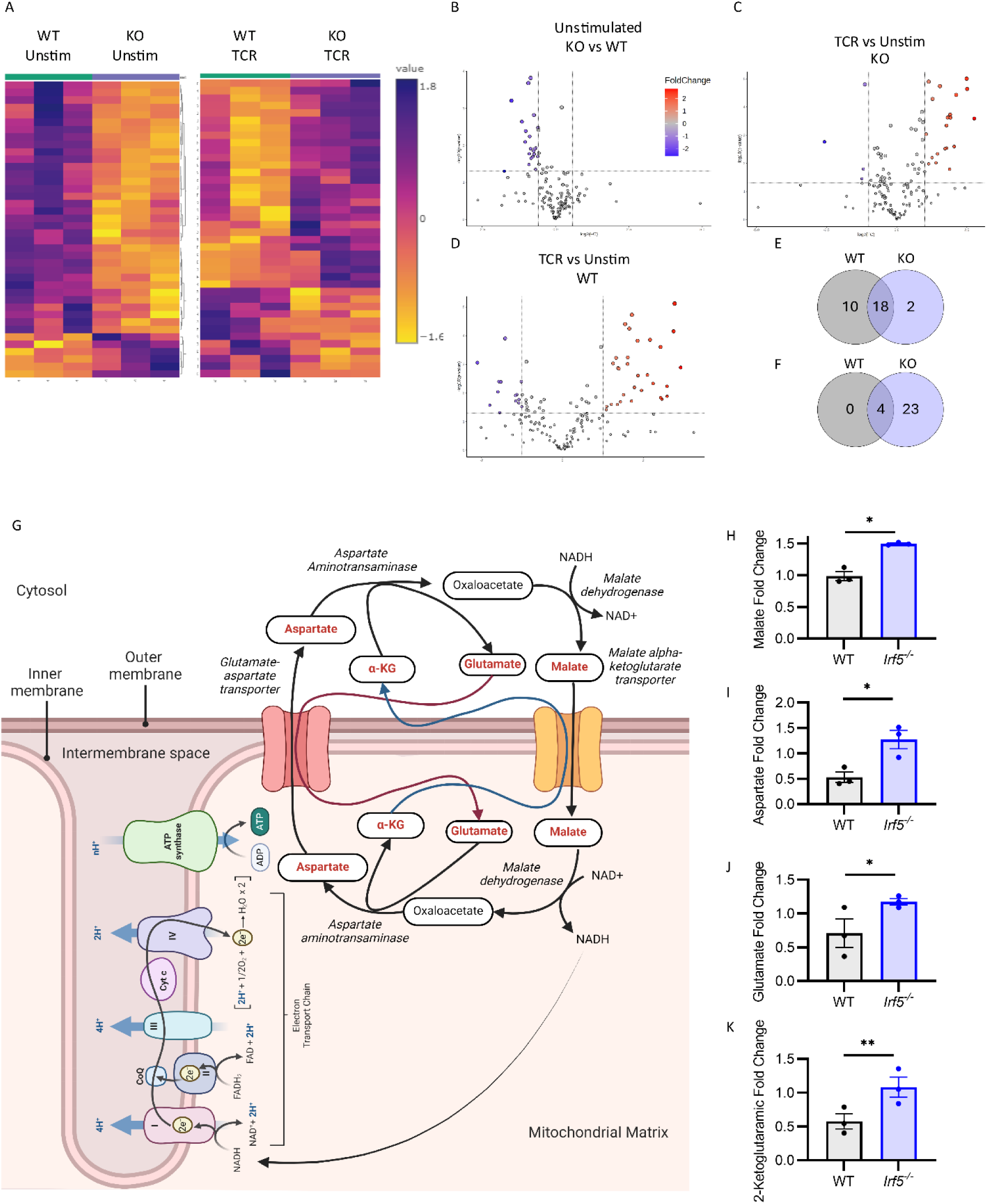
IRF5 regulates CD4 T cell metabolism. (**A**) Purified naive CD4 T cells from 3 WT and 3 *Irf5^−/−^* mouse spleens were analyzed using untargeted LC-MS approach following either 24-hours *in vitro* stimulation with anti-CD3/CD28 (TCR) or IL7 (Unstim). Heatmap of top 40 differentially expressed metabolites in WT and KO CD4 T cells generated using Metaboanalyst 5.0. (**B**-**D**) Volcano plots of significantly altered metabolites in (**B**) IL7 unstimulated (Unstim) KO compared to WT, (**C**) TCR vs Unstim KO, and (**D**) TCR vs Unstim WT. Significance determined by *P* value < 0.05 and log2FC >1.5. (**E, F**) Venn diagrams showing shared and unique (**E**) upregulated and (**F**) downregulated metabolites following TCR stimulation in WT and KO T cells. Significance determined by *P* value < 0.05 and log2FC >2.0. (**G**) Schematic of the malate-aspartate shuttle. Metabolites significantly enriched in KO T cells are written in red. (**H**-**K**) Summary graphs of the following normalized metabolite levels detected in LC-MS analysis of WT and KO T cells (**H**) malate, (**I**) aspartate, (**J**) glutamate, (**K**) alpha-ketoglutaramate. *P* value calculated by two-way unpaired T test. *p < 0.05, **p < 0.01.

### IRF5 and glutamine metabolites regulate glutamine transporter protein expression

As previously described, the *in vitro* metabolomics analysis revealed increased levels of glutamate in KO T cells (**Fig. 5J**). However, previous studies demonstrated that inhibition rather than enrichment in glutamine metabolism is a molecular mechanism to inhibit Th1 and Th17 effector functions, drive Treg generation, and inhibit T cell activation and proliferation, phenotypes that have been described by us and others in KO T cells (**Supp. 1**) (*67–70*). Thus, given the results of our *in vitro* metabolic studies that demonstrated increased glutamine metabolites in KO T cells (**Fig. 5**), we next interrogated other factors involved in glutamine transport metabolism in CD4 T cells. Two key murine glutamine transporters expressed in CD4 T cells are ASCT2 and SNAT2. Following TCR stimulation, ASCT2 and SNAT2 expression were downregulated in KO T cells (**Fig. 6A-C**). However, there was no significant difference in *Slc1a5* (ASCT2) or *Slc38a2* (SNAT2) transcript expression (**Fig. 6D, E**). Further evaluation of key glutamine metabolic enzymes, Glutamate dehydrogenase 1 (*Glud1*) and Glutaminase 2 (*Gls2*) revealed no significant differences in transcript expression (**Supp. 6A, B**). Interestingly, although there was no change in GLUD1 expression (**Supp. 6C, D**), there was a dramatic, albeit insignificant reduction in GLS2 protein expression in KO T cells following anti-CD3/CD28 stimulation (**Supp. 6E, F**). Together, these results indicate that the increased levels of glutamine metabolites in the KO T cells are more likely due to contributions from upregulation of the MAS rather than an upregulation in glutamine metabolism.

**Fig. 6.**
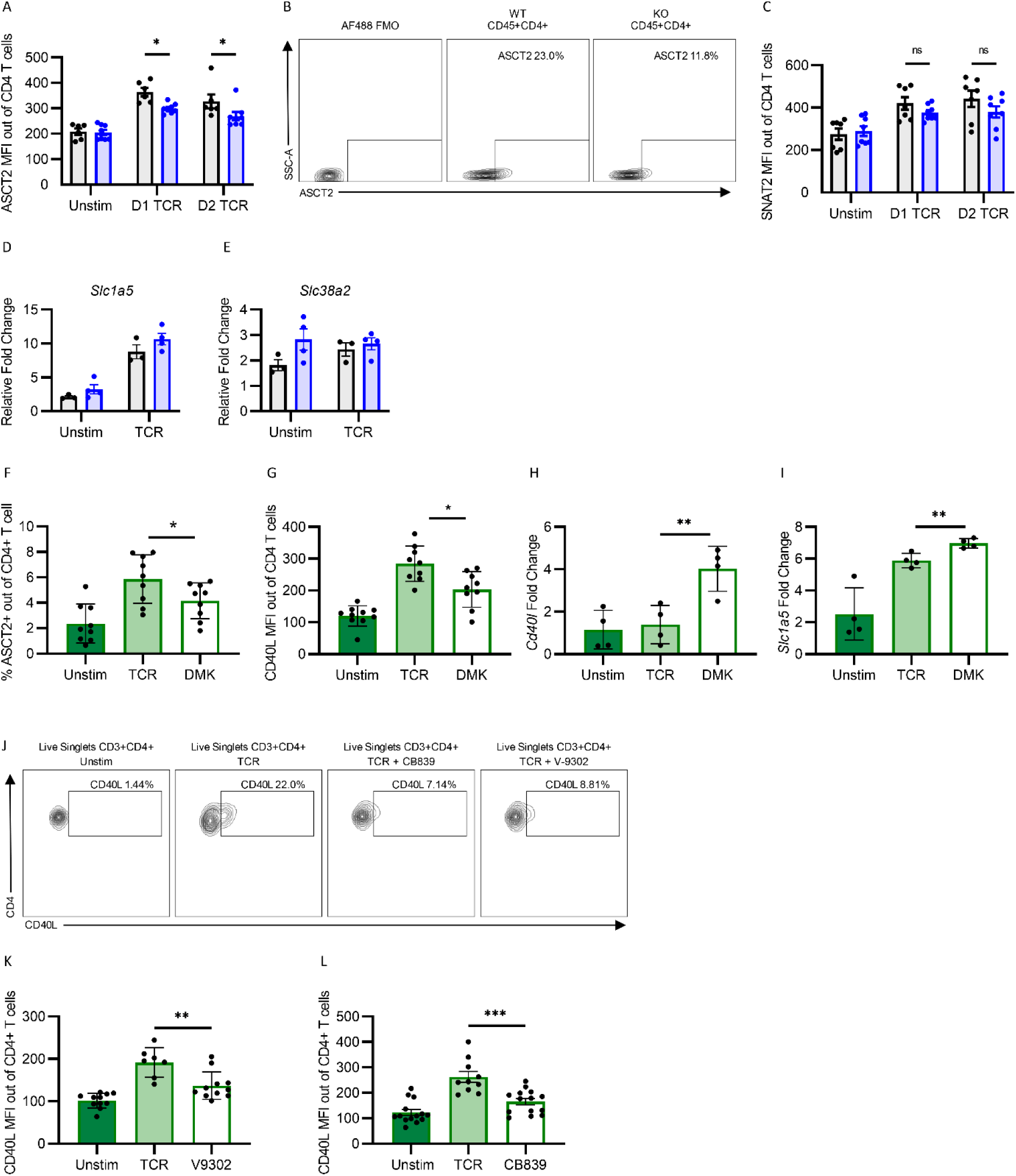
IRF5 regulation of glutamine metabolism modulates effector protein expression. (**A**) WT and *Irf5^−/−^* total splenocytes were activated for 24- (D1) and 48-hours (D2) *in vitro* with anti-CD3/CD28 (TCR). Summary graphs of ASCT2 expression pooled from two independent experiments. *P* value calculated by 2-way ANOVA with Holm-Sidak correction for multiple comparisons. (**B**) ASCT2 representative gating strategy in Live CD45^+^CD4^+^ T cells gated from WT and *Irf5^-/^* total splenocytes. (**C**) Summary graph of SNAT2 expression pooled from two independent experiments. Sorted CD3^+^CD4^+^ T cells WT and *Irf5^−/−^* splenocytes were activated for 6 hours *in vitro* with anti-CD3/CD28 (TCR). (**D**) *Slc1a5* and (**E**) *Slc38a2* transcript expression normalized to *β-Actin*. (**F, G**) WT and *Irf5^−/−^* total splenocytes were activated for 24-hours *in vitro* with anti-CD3/CD28 in the presence (DMK) or absence (TCR) of dimethyl ketoglutarate (DMK). Summary graphs of (**F**) ASCT2 and (**G**) CD40L expressing CD3^+^CD4^+^ T cells pooled from three independent experiments. (**H, I**) WT sorted CD3^+^CD4^+^ T cells were activated for 6-hours *in vitro* with anti-CD3/CD28 (TCR) in the presence (DMK) or absence (TCR) of dimethyl-ketoglutarate. Summary graphs of (**H**) *CD40l* expression and (**I**) *Slc1a5* transcript levels normalized to *B-Actin.* Four biological replicates. (**J**) Representative flow gating strategy for CD40L expression in Live CD3^+^CD4^+^ T cells under indicated treatment conditions. WT total splenocytes were activated for 24-hours *in vitro* with anti-CD3/CD28 (TCR) in the presence of ASCT2 inhibitor (V-9302) or absence (TCR). Summary graph of (**K**) CD40L expression in CD3^+^CD4^+^ T cells. Data pooled from three independent experiments. (**L**) WT total splenocytes were activated for 24-hours *in vitro* with anti-CD3/CD28 (TCR) in the presence (CB839) or absence (TCR) of CB839. Summary graphs of CD40L expression in activated CD3^+^CD4^+^ T cells. Data pooled from three independent experiments. *P* value calculated by one-way analysis of variance (ANOVA) with Tukey’s post-hoc test for multiple comparisons unless otherwise noted. Bar graphs show means +/- SEMs. *p < 0.05, **p < 0.01, ***p < 0.001, ns = not significant. Each point represents an independent biological replicate.

Given the observed increase in the intermediary metabolite of α-ketoglutarate (α-KG), α-KGM, in KO T cells (**Fig. 5J**), we next examined if ASCT2 expression was influenced by the upregulation of this epigenetic regulator of immune responses (*68, 71–74*). Following treatment of WT CD4 T cells with the cell permeable α-KG analog, dimethyl-ketoglutarate (DMK), we observed a significant reduction in ASCT2 expression (**Fig. 6F**). As previously discussed, glutamine metabolism regulates T helper cell function. Thus, we examined if alterations in metabolites contribute to CD40L dysregulation observed in KO T cells. Following WT T cell DMK treatment, we found a significant downregulation of CD40L expression (**Fig. 6G**). As previously reported, treatment with DMK did not inhibit S6 phosphorylation, indicating that DMK modulation of CD40L expression signals through an mTORC1-independent pathway (**Supp. 6G**) (*75–77*). In support of the previously established regulatory role of α-KG in demethylation via the ten-eleven translocation (Tet) α-KG dependent dioxygenases, increasing levels of DMK resulted in a significant increase in transcript expression for both *Cd40l* and *Slc1a5* despite the significant reductions in CD40L and ASCT2 protein expression (**Fig. 6H, I**) (*77*). These findings indicate a more complex role for α-KG regulation at a post-epigenetic level.

To further elucidate how the reductions in ASCT2 expression mechanistically contribute to defects in KO CD4 T cell function, we sought to mimic the reduction in ASCT2 expression by using the small molecule ASCT2 inhibitor, V-9302. Treatment of WT CD4 T cells with V-9302 dramatically inhibited CD40L expression (**Fig. 6J, K**). Further, inhibition of glutaminase, the enzyme responsible for converting glutamine into glutamate, with CB-839 (Telaglenastat) also inhibited CD40L expression in WT CD4 T cells (**Fig. 6J, L**). Together, our findings indicate that increased intracellular glutamine metabolites act as negative regulators of glutamine transporter and enzyme expression, which in turn are required for the positive regulation of CD40L.

### T cell conditional *Irf5^−/−^* mice are protected from EAE

mTORC1 signaling, ASCT2 expression, and Speckled proteins have been linked as either risk factors for or protective against multiple sclerosis (*7, 13, 70*). Thus, to determine the specificity of our findings, we generated *Irf5^fl/fl^-Lck-Cre^+^* T cell-specific KO mice (cKO) and examined if loss of *Irf5* in T cells was sufficient to confer protection from EAE as previously observed in our whole-body KO studies (**Fig. 3**).

cKO mice had significantly attenuated disease progression with a marked improvement in disease scores compared to the *Irf5*^fl/fl^-*Cre^-^* littermate controls (**Fig. 7A**). Histologic analysis of spinal cord sections from cKO mice showed an intact myelin sheath and reduced inflammatory foci (**Supp. 7A.**), whereas typical demyelination and inflammation were observed in *Irf5*^fl/fl^-*Cre^-^*littermate mice. To examine the role of IRF5 in disease onset as well as disease progression, we harvested spleens at symptom onset (D13) and peak symptoms (D20). cKO spleen sizes showed trends towards increased size compared to *Irf5*^fl/fl^-*Cre^-^* mice at D20 (**Fig. 7B, Supp. 7B**). Despite this, flow cytometric analysis of total splenocytes revealed a significant decrease in the overall percentage of CD4 T cells in the spleens of cKO mice at both timepoints (**Fig. 7C**).

**Fig. 7.**
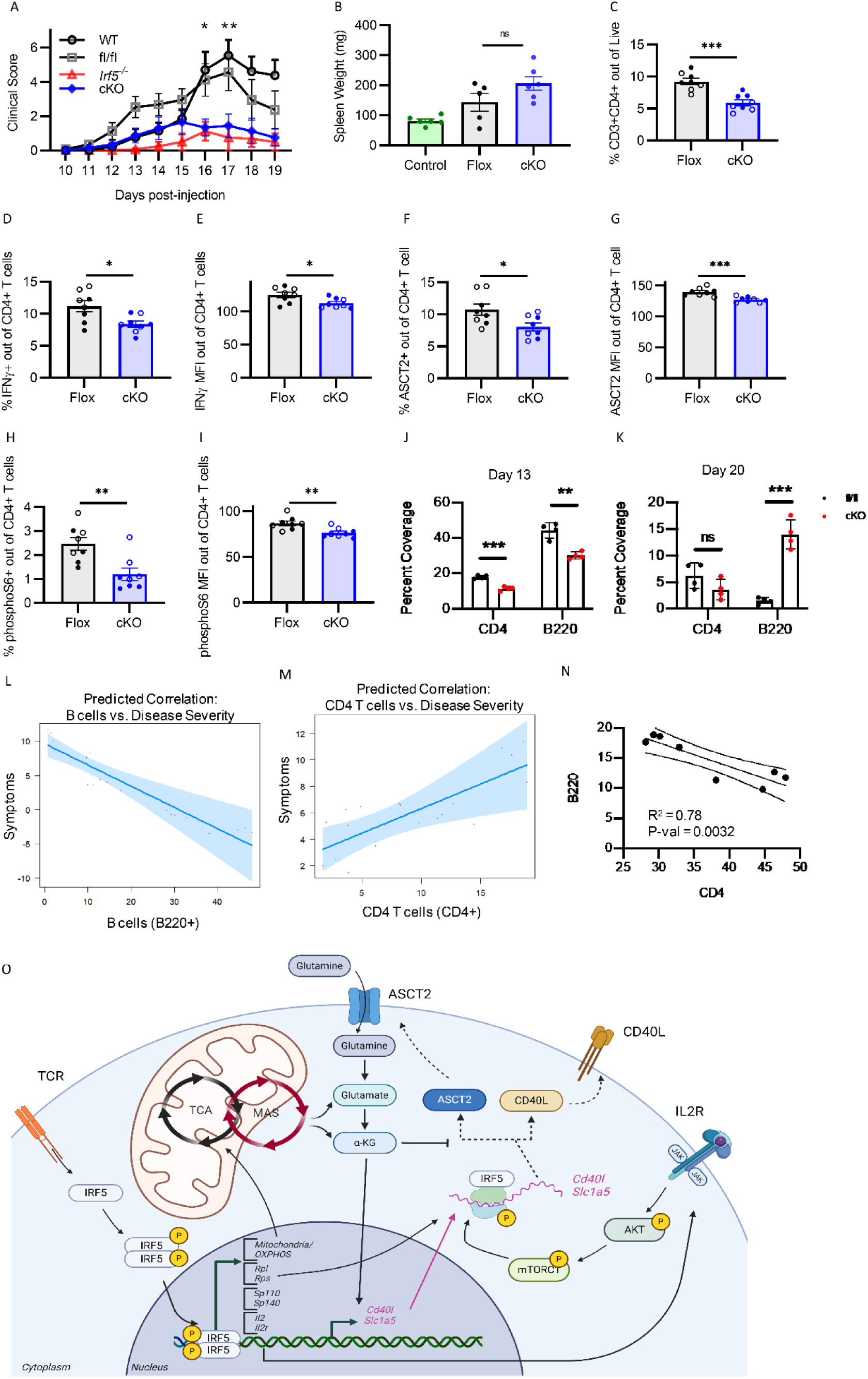
T cell conditional *Irf5^−/−^* mice are protected from EAE. (A) Composite means +/- SEM of clinical scores. Minimum of 10 biological replicates per experiment. Pooled data from three independent experiments comparing clinical progression of *Irf5^fl/fl^-Lck-Cre^+^* (cKO), *Irf5^fl/fl^-Lck-Cre^-^* (fl/fl), *Irf5^−/−^* (KO), and WT mice following EAE induction. (B) Spleen weights from WT control, cKO and fl/fl (Flox) mice at peak disease. Five-six biological replicates pooled from two independent experiments. *P* value calculated by one-way analysis of variance (ANOVA) with Tukey’s post-hoc test for multiple comparisons. (**C**-**I**) Summary graphs of flow cytometric analyses from day 13 post immunization (clear circles) and day 20 post immunization (colored circles). (**C**) CD3^+^CD4^+^ T cells gated from live cells, (**D, E**) IFNγ production from live CD3+CD4+ T cells, (**F, G**) ASCT2 expression, and (**H, I**) S6 phosphorylation (phosphoRPS6). *P* value calculated by multiple T tests with Holm-Sidak correction for multiple comparisons. (**J, K**) Quantification of splenic CD4^+^ and B220^+^ immunohistochemistry staining at (**J**) day 13 post-immunization and (**K**) day 20 post-immunization. Data represents four biological replicates. (**L, M**) Predicted correlation between (**L**) B220 B cells and disease severity and (**M**) CD4 T cells and disease severity. Partial residual plot generated using visreg analysis package (*115*). Shaded area represents 95% confidence interval. (**N**) Linear regression analysis of B220 and CD4 IHC coverage in spleens harvested from mice at Day 13 and Day 20 timepoints post EAE induction. Outer lines represent 95% confidence interval. Bar graphs show means +/- SEMs. *p < 0.05, **p < 0.01, ***p < 0.001, ns = not significant. Each point represents an independent biological replicate. (**O**) Summary schematic of the transcriptional, translational and metabolic regulatory roles for IRF5.

We next investigated if effector T cells were impacted by loss of *Irf5*. We observed a significant reduction in splenic IFNγ production by CD4 cKO T cells (**Fig. 7D, E**). There were no differences in IL17 or IL4 expression in cKO CD4 T cells (**Supp. 7C-F**). Despite the protective phenotype in cKO mice, there was no significant alteration in Tfh (CD4^+^BCL6^+^CXCR5^+^), Treg (CD4^+^FoxP3^+^) or CD40L expression (**Supp. 7G-J**). As previously discussed, ASCT2 expression and mTORC1 activity have been implicated in EAE. Examination of the expression and activation of these molecules, respectively, showed significant reductions in ASCT2 expression (**Fig. 7F, G**) and RPS6 phosphorylation (**Fig. 7H, I**) in cKO CD4 T cells following EAE induction, supporting a vital role for the expression of these molecules in EAE disease pathogenesis, as well as their regulation by T cell intrinsic IRF5.

Our previous studies in NP-CGG CFA immunized mice revealed defects in KO splenic follicles (**Fig. 1A**). Thus, we next examined the spleen at both D13 and D20 in cKO and *Irf5*^fl/fl^- *Cre^-^* mice by IHC. D13 IHC analyses revealed decreased CD4 and B220 cells in cKO spleens, while D20 IHC analyses revealed no significant differences in CD4, but a dramatic increase in B220 coverage (**Figs. 7J, K, Supp. 7K, L**). To better elucidate how these early histologic findings correlated with disease progression, spleens from *Irf5*^fl/fl^-*Cre^-^* and cKO mice were stratified by disease score (advanced score > 10, moderate score 6 to 8, minimal score 1 - 2) and correlations performed between B or T cell coverage and disease score. Our findings revealed that increasing EAE disease scores inversely correlated with B220 coverage (**Fig. 7L**) and positively correlated with CD4 coverage (**Fig. 7M**), with the coverage of CD4 thus inversely correlated with that of B220 (R^2^ = 0.78, p-val< 0.005) (**Fig. 7N**).

Notably, Sp110 and Sp140 dysregulation has been implicated in the pathogenesis of MS (*33*). Thus, to further assess the link between loss of *Irf5* expression and *Sp* expression and validate our scRNAseq findings, we performed qPCR on sorted CD4 T cells and B cells from cKO mice following anti-CD3/CD28 stimulation, as previously described. Our findings revealed a select decrease in *Sp110* and *Sp140* in cKO CD4 T cells (**Supp. 7M-O, Fig. 2**). Taken together, these data demonstrate a clear relationship between the presence and function of CD4 T cells and B cells in modulating EAE disease pathogenesis, identifies regulatory pathways by which inhibition of T cell intrinsic IRF5 may offer protection from EAE disease onset and severity, and reveals novel IRF5 translational and transcriptional targets (**Fig. 7O**).

## DISCUSSION

Recent studies have demonstrated the vital importance of metabolic and translational regulatory pathways in T cell function (*78–81*). IRF5 was previously implicated as a mediator of T helper cell-intrinsic cytokine/chemokine expression and migration. However, prior to this study, a thorough functional examination of T cell intrinsic IRF5 beyond its canonical role as a transcriptional regulator of proinflammatory cytokines had yet to be conducted (*4, 43, 82*).

Our mechanistic studies using immunophenotyping, targeted inhibitor assays, metabolomics, co-immunoprecipitation and scRNAseq analyses revealed dysregulation of ribosomes, protein translation and metabolism in *Irf5^−/−^* T cells. Specifically, we found an overall reduction in *Irf5^−/−^* T cell translational capacity and mTORC1 activity. *Uba52*, ATF4, and eEF2 were identified as candidate regulatory targets of IRF5 that may contribute to the observed reduction in protein translation. UBA52, a ubiquitin-ribosomal fusion protein, is an understudied master regulator of translation that functions through roles in both the assembly of the ribosome translational complex and as a major supplier of ubiquitin. Inhibition of global protein synthesis and decreased proliferation in KO T cells with decreased *Uba52* expression closely replicates findings from prior studies, indicating a likely role for IRF5-dependent UBA52 expression in the regulation of CD4 T cells (*83*). Analysis of IRF5 interacting partners in Ramos B cells revealed significant enrichment for EIF6 and EIF4A2. Both of these factors are of particular interest as EIF6 plays a key role in ribosome biogenesis (*84*) while EIF4A2 is a RNA helicase involved in translational repression via the miRNA degradation pathway (*85, 86*). Thus, it stands to reason that loss of *Irf5* may alter the function and/or expression of EIF6 and EIF4A2 and hence reduce protein translation. Lastly, as previously discussed, ATF4 is a key regulator of protein translation and metabolism, whose expression we reveal as dysregulated with loss of *Irf5*. Further validation of these candidate regulatory targets and their impacts on T cell function will be a focus of future study.

Beyond this, rapamycin inhibition assays provided evidence that mTORC1 signaling is crucial for CD4 T cell support of B cell adaptive immunity through regulation of CD40L expression. Prior studies demonstrated correlations between reduced CD40L expression on T cells and impaired Th1 polarization, recapitulating our findings in KO CD4 T cells (*87, 88*). However, the T cell intrinsic regulatory mechanisms governing CD40L expression have yet to be fully elucidated. Here, we provide compelling evidence that mTORC1 activity and glutamine metabolism are molecular regulators of CD40L expression. We further demonstrate that elevated levels of the metabolite α-KG were sufficient to reduce both glutamine transporter expression and CD40L through post-transcriptional mechanisms, thereby curtailing the adaptive immune response. Interestingly, prior studies showed increased α-KG levels produced from glutamine metabolism drives M2-like macrophage polarization, while other studies have demonstrated the promotion of alternative (M2-like) anti-inflammatory macrophages from M1 occurs with loss of *Irf5* (*89–91*). Our data suggest that IRF5-mediated alterations in α-KG may contribute to its role in macrophage polarization. Beyond glutamine metabolism, another metabolite of interest that was uniquely downregulated in KO but not WT T cells following stimulation was glycerolphosphorylethanolamine (GPE), levels of which directly correlate with phosphatidyl ethanolamine (PE) (**Supp. Table 4, 5)** (*92*). Recent studies have identified PE as a key metabolite in Tfh differentiation and support of humoral immunity. It is tempting to speculate that dysregulation of these metabolic pathways in KO mice may contribute to our findings of reduced Tfh cells (*93*). Lastly, our findings demonstrate increased reliance of KO CD4 T cells on the malate aspartate shuttle. The MAS is an alternative method by which cells can generate NADH and provide electrons to the electron transport chain. Our current findings, along with prior work demonstrating defects in oxidative phosphorylation with loss of *Irf5* (*56, 65*), suggest that mechanisms exist by which KO T cells compensate for decreased oxidative phosphorylation. Taken together, we propose that loss of *Irf5* in CD4 T cells drives aberrant effector function through a combination of transcriptional, translational, and metabolic reprograming (**Fig. 7O**).

Metabolic intervention to combat disease has been met with success in both preclinical and clinical studies. Increasing our understanding of how immune cells respond to metabolic dysregulation is crucial for continued advancement. Through unbiased metabolomics analysis, we reveal global alterations in T cell metabolism with loss of *Irf5*. Prior studies have shown that reducing glutamine in media drives Tregs while inhibition of glutamate conversion to α-KG with the MAS inhibitor, AOAA, drives Th17 inflammatory cells through 2-hydroxyglutarate dependent methylation of the *FoxP3* promoter. Here, our studies reveal a shift towards a Treg phenotype with loss of *Irf5* and an increase in intracellular α-KG and glutamate, supporting a role for IRF5 in the metabolic regulatory axis governing Th17 and Treg fate decision. Of further interest, α-KGM is an intermediary molecule in the conversion of glutamate to α-KG through the understudied glutaminase II pathway (*94*). Unlike the conversion of glutamate to α-KG by glutamate dehydrogenase, conversion of α-KGM to α-KG results in the production of NH_4_^+^. High ammonia levels inhibit T cell activation, proliferation and contribute to T cell exhaustion (*95*). Further studies elucidating the role of T cell intrinsic ammonia production in T effector functions, and by extension how increasing levels of α-KGM might contribute to the dysregulation of *Irf5^−/−^* T cells, remains to be performed.

IRF5 has canonically been described as a transcription factor. Our scRNAseq analyses highlighted the differentiating potential and transcriptional regulation of T cell subsets, particularly the Th complex, by IRF5. DEG analysis in WT CD4 T cells revealed Th complex cells to express chemokines and chemokine receptors in addition to proinflammatory and inhibitory cytokines, transcription factors and proteins. These factors describe a migratory CD4 T cell population poised to rapidly respond to stimuli with the potential to gain proinflammatory or inhibitory effector functions. Notably, we found the Th complex subset to be significantly enriched in *Irf5^−/−^*mice. Interestingly, analysis of the transcriptional profiles of *Irf5^−/−^*Th complex subset showed them poised for an anti-inflammatory response with reduced translational capacity compared to their WT counterparts. It is tempting to speculate that these transcriptional differences contribute to the protection observed in EAE cKO mice. Of note, a recent paper revealed that CXCR4-dependent MOG autoreactive T cell migration into the bone marrow was required for CCL5-dependent aberrant inflammatory myelopoiesis that escalates the CNS demyelination characteristic of EAE (*96*). These same pathways were downregulated in *Irf5^−/−^* Th complex cells.

Our scRNAseq analysis additionally revealed two autoimmune risk genes under IRF5 transcriptional regulation: *Sp110* and *Sp140*. SP110 and SP140 are nuclear body proteins hypothesized to regulate gene transcription, ribosome biogenesis and apoptosis (*97, 98*). Of interest, *SP110* missense mutations and deletions are associated with Hepatic Venoocclusive Disease with Immunodeficiency (VODI), which clinically presents with severe hypogammaglobulinemia, a T-cell immunodeficiency, absent lymph node germinal centers (GCs) and absent tissue plasma cells; findings remarkably similar to those reported in *Irf5^−/−^* mice (*99*). SP140 has been identified as a modulator of chromatin accessibility in macrophages (*100*). Previous ATAC-seq studies performed in *Irf5^−/−^*T cells revealed significant alterations in chromatin accessibility. This finding was not attributed to IRF5’s role as a transcription factor (*4*). Furthermore, analysis of the *SP110* and *SP140* promoter region with Genome.UCSC revealed predicted binding sites for IRF5. We propose that *Sp110* and *Sp140* are IRF5-specific regulatory targets. Examination of this novel IRF5 regulatory axis in other immune cells will be the focus of future studies.

Prior work described alterations in SP110 expression, glutamine transporter expression and aberrant mTORC1 activity as significant risk factors for MS. Here, we show each of these risk factors to be regulated by IRF5 (*101, 102*). cKO protection from EAE further demonstrated the preclinical relevance of IRF5 expression in CD4 T cells. The alterations in secondary lymphoid architecture and reduced B cell egress from the spleens provide additional compelling evidence that the expression of IRF5 in CD4 T cells is crucial for the B cell adaptive response. In the context of MS, mounting evidence demonstrates the crucial role for pathogenic B cell maturation and antibody secretion in driving demyelination (*103*). Treatment with rituximab to deplete B cells is gaining traction as an effective therapy for treatment resistant disease (*104, 105*). However, this fails to counter existing plasma cells which lack CD20 expression (*106*). Thus, inhibiting the generation of pathogenic autoantibody secreting B cells has remained a challenge due to a lack of understanding of modulators of disease pathogenesis (*106–109*). Based on the results of our studies, we propose that inhibiting B cell maturation and autoantibody production by targeting IRF5 activity in CD4 T cells through recently developed IRF5 specific inhibitors is a potential therapeutic strategy for the treatment of MS (*35, 41*).

In summary, our findings highlight novel regulatory roles for IRF5 in mTORC signaling, as well as in the metabolic, transcriptional, and translational regulation of CD4 T cells. We elucidate how subtle shifts in metabolism have a profound impact on the adaptive immune response with clinical implications. Altogether, these data significantly increase our knowledge of IRF5 regulatory pathways in CD4 T cells at a transcriptional, translational and metabolic level, expands our understanding of metabolic regulation of signaling pathway molecules, broadens our knowledge of the KO CD4 single cell landscape, and highlights IRF5 expression in CD4 T cells as a potential therapeutic target in the treatment of multiple sclerosis.

In the context of metabolic dysregulation, previous studies established that disruption of metabolites drives transcriptional alterations and epigenetic remodeling, which in turn reprograms immune cells. Thus, a limitation of this study is an inability to definitively determine the primary perturbation driving dysregulation in these highly interlinked transcription-translation-metabolic feedback loops. We attempted to address this using small molecule inhibitors in unperturbed WT systems. However, our findings only provide partial insight into the complex interplay between metabolism and T cell function. In addition, in the cKO EAE model, we did not observe reductions in CD40L. Review of literature shows that CD40L has historically been difficult to study directly due to the highly transient and early expression of this signaling molecule. Further studies investigating the kinetics of CD40L expression in EAE models are of interest.

## MATERIALS AND METHODS

### Study design

This study was designed to understand the molecular mechanisms by which IRF5 modulates T cell effector function in response to TCR activation *in vitro* and in the *in vivo* EAE disease model. Endpoint analyses for *in vitro* and *in vivo* studies included flow cytometry, quantitative polymerase chain reaction (qPCR), scRNAseq, and unbiased metabolomics. All *in vitro* studies used randomly assigned mice without investigator blinding. We compared the clinical response to EAE induction following MOG_35-55_/PTX injections of 8-12 weeks-old C57BL/6 *Irf5^+/+^* (WT) and *Irf5^−/−^* (KO) littermate mice, and *Irf5^fl/fl^-Lck-Cre^-^* and *Irf5^fl/fl^-Lck-Cre^+^*littermate mice. Mechanisms of immunopathology were investigated during early onset and peak disease as determined by a common clinical scoring method. Clinical scoring was performed with investigator blinding. All data points and *n* values reflect biological replicates. The specific numbers and genotypes of the mice and the statistics performed are included in the figure legends such as the two-way unpaired *t* test, 1-way analysis of variance (ANOVA) with Tukey’s post-hoc test for multiple comparisons, and 2-way ANOVA with Holm-Sidak correction for multiple comparisons as indicated. Experiments were repeated in multiple settings using complementary methods of molecular interrogation and laboratory techniques to validate findings. All data points from the studies were included unless methodologic errors occurred resulting in decreased cell viability, poor activation, sample contamination or low cell numbers.

### Mice

C57BL/6J *Irf5^−/−^* mice were originally obtained from Taniguchi via the Rifkin Lab and backcrossed 11 generations (*110*). C57BL/6J *Irf5^+/+^*WT littermate mice were used as controls. *Irf5^fl/fl^-Lck-Cre^+^* mice were a kind gift from Simona Stager’s lab (1INRS-Institut Armand-Frappier). C57BL/6J *Irf4^−/−^*mice were a kind gift from Dr. Alessandra Pernis (Hospital for Special Surgery). Genotype was confirmed using PCR. Mice used in experiments were between 6 weeks to 4 months of age and of both genders. Experiments were performed in agreement with our Institutional Care and Use Committee and according to National Institutes of Health guidelines. All mice were bred in house at the Feinstein Institutes for Medical Research. *Irf5*^fl/fl^ cage-matched littermates (*Irf5^fl/fl^-Lck-Cre^-^*) were used as wild-type controls in experiments utilizing *Irf5^fl/fl^-Lck-Cre^+^* (cKO) mice. For all experiments, mice were age- and sex-matched.

### Naïve CD4 T cell isolation

Freshly isolated splenocytes from WT and KO mice were homogenized in 4 mL PBS. Red blood cell (RBC) lysis was performed in 8 mL of RBC lysis buffer (Biolegend) for 5 minutes on ice and quenched with excess PBS + 2% FBS (vol/vol). Naïve CD4 T cells were isolated using Miltenyi T cell Isolation Kit following manufacturer’s protocol (Cat#: 130-104-453). Magnetic separation was performed to achieve a >95% enriched population of naïve CD4 T cells. For T cell activation, cells were cultured in RPMI 1640 supplemented with 10% FBS and 1X Pen/Strep in the presence of either 1 ng/mL Recombinant murine IL-7 (PeproTech, Cat#217-17) or stimulated with anti-CD3/CD28 Dynabeads at a 4:1 bead:cell ratio (Fisher Scientific, cat#: 11-452-D).

### Total B cell and CD4 T cell isolation

Total splenocytes were isolated from mice and RBCs lysed as previously described. Cells were stained with B220R PerCP and CD4 BV510 in FACS buffer for 30 minutes at room temperature, briefly washed in FACS buffer, and sorted using BDFacsAria.

### In vitro CD4 T cell proliferation

CD4 T cells were sorted then stained with CellTrace™ Violet Proliferation Dye as per manufacturer protocols (Invitrogen™ Cat#:C34557). Labeled CD4 T cells were activated with anti-CD3/CD28 Dynabeads and cultured for four days before being analyzed by flow cytometry.

### Immunizations

Intraperitoneal injections were performed using 200 µl/mouse. Sterile PBS solution was used to reconstitute 4-Hydroxy-3-nitrophenylacetyl hapten conjugated Chicken Gamma Globulin (NP-CGG, Ratio 29:20) to a final concentration of 2 mg/mL (Biosearch Technologies, Cat#: N-5055C-5). Complete Freund’s Adjuvant (CFA) was purchased from Invivogen (Cat#: vac-cfa-60). Intraperitoneal injections of NP-CGG-CFA were conducted using 100 µg/100 µL NP-CGG emulsified in 100 µL CFA.

### Flow cytometry

Samples were harvested, washed in PBS + 2% FBS (vol/vol), blocked with anti-CD16/CD32, and stained with a Live/Dead Fixable Dead Cell stain prior to surface and intracellular stains. For a detailed list of antibodies used in staining panels, see Supplemental Methods. Intracellular staining was performed per manufacturer’s protocol using Transcription Factor Staining Buffer Set (Cat#: 00-5523-00, eBioscience^TM^). Phospho-flow cytometry was performed using the two-step fixation/methanol protocol (Protocol C: Two-step Protocol for Fixation/Methanol, ThermoFisher).

### In vitro CD4 T cell activation and cocultures

Sorted CD4^+^ T cells or total splenocytes were cultured in 24-well flat-bottom plates at a density of 2 × 10^6^ cells/500 µL media (Fischer Scientific, cat# 3473). For T cell activation, cells were cultured in RPMI 1640 supplemented with 10% FBS (v/v) and 1X Penn/Strep and stimulated with anti-CD3/CD28 Dynabeads as per manufacturer’s protocol (Fisher Scientific, cat#: 11-452-D). For cocultures, 2 x 10^6^ sorted CD4^+^ T cells and 2 x 10^6^ sorted B cells were co-cultured for 4 – 5 days in the presence of Dynabeads, 10 ng/mL CpG-B (ODN 2006) (Fischer Scientific, Cat#: HC4039) and 10 µg/mL anti-IgM (Southern Biotech, Cat#: 1021-01). For rapamycin pretreatment, CD4^+^ T cells were sorted and plated in RPMI 1640 supplemented with 10% FBS and 1X Penn/Strep and then incubated at 37°C with either 100 nM of rapamycin or equal volume of PBS (control) for 2 hours (Selleck Chemicals, Cat#: S1039). Media was removed and cells were washed three times with excess volume of PBS prior to being plated with sorted B cells. Cocultures were stimulated following rapamycin treatment as previously described.

### In vitro protein synthesis quantification

OPP staining was performed using Click-iT plus OPP Alexa-555 protein synthesis kit per manufacturer’s protocol (ThermoFischer, Cat#: C10456). Briefly, 2 x 10^6^ cells were incubated at 25°C for 30 minutes with 1:400 dilution of the Click-iT OPP reagent. Cells were washed three times with PBS and then fixed in PBS with 4% formaldehyde (v/v). Following fixation, cells were permeabilized in PBS supplemented with 2.0% saponin and 3.0 % FBS (v/v) for 15 minutes. For the Click-iT reaction, cells were incubated in the dark at room temperature for 30 minutes in the Click-iT reaction cocktail. Samples were then washed twice with PBS supplemented with 2% FBS and immediately analyzed by flow cytometry.

### qPCR

RNA was isolated from sorted CD4^+^ T cells following specified stimulation and treatments using the Qiagen RNeasy Mini Kit (Cat#: 74106). cDNA synthesis was performed with the GoScript reverse transcription system (Cat #: A5001). qPCR was performed in triplicate for each sample using the PowerUp SYBR green real-time PCR master mix with 5–10 ng input cDNA (cat#: A25776). Threshold values (*C*_T_) were averaged across sample replicates, followed by normalization *via* the ΔΔ*C*_T_ method to *β-actin*. For primer sets, see **Supp. Table 6**.

### Apoptosis analysis

Total splenocytes were cultured for 24 hours as previously described. All stains were performed protected from light. Cells were washed twice with prewarmed FACS buffer and surface stained as previously described. Cells were then washed once with prewarmed FACS buffer and Annexin V binding buffer. Samples were resuspended in 100 µL binding buffer supplemented with 5 µL of Annexin V-FITC (Cat#: 10040-02, SouthernBiotech) and stained for 20 minutes at room temperature. Cells were washed once with Annexin V binding buffer (Cat#: 10045-01, SouthernBiotech) then resuspended in 200 µL of binding buffer and stained with 5 µL of 7-AAD (Cat#: 420403, Biolegend) for 15 min at room temperature. Samples were placed on ice and analyzed using Fortessa.

### Immunoblot analysis

Briefly, whole cell lysates were prepared using NP-40 lysis buffer (50 mM Tris-HCl (pH 7.4), 150 mM NaCl, 1% NP-40 and 5 mM EDTA) (Thermo Scientific, J60766-AP) supplemented with Halt Protease Inhibitor Cocktail (Thermo Scientific, 87786) and PhosStop Phosphatase Inhibitor Cocktail (Roche, 4906845001). Sample protein concentrations were quantified using the DC protein assay (Bio-Rad, 5000112). 20 μg of protein per sample were separated by SDS-PAGE using the Bolt Bis-Tris system (Invitrogen). Proteins were transferred to 0.45 μm nitrocellulose membranes (MDI, SCNX8402XXXX101) using a wet tank transfer system. The membranes were blocked for 1 hour at room temperature with 5% bovine serum albumin (BSA) in TBST and incubated overnight at 4°C with the primary antibody diluted in the blocking buffer. The membranes were washed three times for 5 min each with TBST and incubated with the secondary antibody diluted in the blocking buffer for 1 hour at room temperature. The membranes were washed five times for 5 min each with TBST and incubated with 1 mL of chemiluminescent detection reagent (Cytiva, RPN2232) for 3 min before image acquisition using a ChemiDoc MP Imaging System (Bio-Rad Laboratories). Horseradish peroxidase (HRP)-conjugated antibodies against β-actin (Cell Signaling, 12620, 1:5000) were used as loading controls for protein normalization. Densitometric analysis was performed using the Image Lab software (Bio-Rad Laboratories).

### Metabolomics analysis

For each sample, approximately 3 × 10^6^ naïve splenic CD4^+^ T cells were purified and stimulated as previously described. LC-MS/MS analysis was performed by The Metabolomics Innovation Centre (Alberta, Canada). For detailed protocol, see Supplemental Methods.

### Metabolomics Pathways Interpretation

Biological interpretation was performed by MetaboAnalyst 5.0 using the metabolite set library *Homo sapiens* based on normal human metabolic pathways from the Kyoto Encyclopedia of Genes and Genomes (KEGG) database and the Human Metabolome Database (HMDB).

### scRNA-Seq sample preparation

FACS-sorted WT and KO CD4^+^ T cells were purified and stimulated as previously described, then directly processed for scRNA-seq with 10X Genomics 3′ kit (10X Genomics, Pleasanton, CA) following the manufacturer’s instructions. 10,000 CD4 T cells were used to construct single-cell libraries with the Chromium Single Cell 3′ Reagent Kits (v2 Chemistry) according to manufacturer’s instructions. Libraries were sequenced using the Illumina platform.

### scRNA-Seq data analysis

scRNA-seq data were aligned to mm10 using CellRanger v.3.1.0 and downstream processing was performed using Seurat v3.1.1 (*111*). Briefly, cells with fewer than 200 features, higher than 1% mitochondrial gene content, or transcripts expressed by fewer than 3 cells were removed prior to log normalization. Principal component analysis (PCA) was performed, and the subsequent Uniform Manifold Approximation and Projection (UMAP) analysis was conducted using the first 30 principal components. Cluster-specific genes were determined using the FindMarkers algorithm in the Seurat suite. Clustering was performed by calculating a shared nearest neighbor graph with a resolution of 0.5. The FindAllMarkers function determined differentially expressed genes (DEG) based on the non-parametric Wilcoxon rank sum test for each subset. Subsetting was performed using previously published markers of T cell subsets. Trajectory analysis was performed using Slingshot (*112*).

### Histological analysis

For histological analysis, spleens, right inguinal lymph nodes, and spinal cords were harvested as indicated and fixed in 10% Neutral Buffer Formalin (NBF) overnight before being transferred to 70% EtOH. For spinal cord harvest, mice were CO_2_ euthanized, then immediately perfused with 10 mL PBS followed by 10 mL of 10% NBF. All samples were sent to Histowiz for paraffin embedding and histological analysis (Brooklyn, NY). A minimum of three independent samples were sent for each study. Immune cell infiltration and lymphoid architecture were examined using Hematoxylin and Eosin (H&E), anti-CD4 (Cat# ab183685, Abcam) and anti-B220 (Cat#: NB100-77420, Novus Biologicals). Demyelination was analyzed using Luxol Fast Blue. IHC quantification was performed using ImageJ.

### EAE immunization

8-12 weeks-old male and female mice, minimum of 3 per gender and genotype, were subcutaneously injected at three sites with 200 µg of MOG peptide 35–55 emulsified in Complete Freund’s Adjuvant (CFA) containing 400 µg of Mycobacterium (Cat#: EK-0111, Hooke Laboratories). On day 0 (D0) and 1 day after (D1) immunization, mice were intraperitoneally injected with 200 ng of pertussis toxin (Cat#: 180, List Biological Laboratories). All mice were examined and graded daily for neurological signs in a blinded manner as previously described. For the tail: 0, no disease; 1, half paralyzed; 2, full paralysis. For each hind and forelimb assessed separately: 0, no disease; 1, weak or altered gait; 2, paresis; 3, limb paralysis; and 5, moribund state (*46*). Average clinical scores were calculated daily for each group of mice and plotted. Immunological studies were performed on the onset (13 days after immunization) or peak (20 days after immunization) of disease. 4 female mice were chosen for each genotype for further molecular analyses according to typical and representative clinical symptoms.

### Statistical analysis

GraphPad Prism v.9.2 was used for statistical analysis. Statistical analysis was performed using. *P* value calculated by two-way unpaired *t* tests, one-way ANOVA with Tukey’s post-hoc test for multiple comparisons, or 2-way ANOVA with Holm-Sidak correction for multiple comparisons as indicated. Differences were considered statistically significant when p < 0.05.

## Supplementary Materials and Methods

### Detailed metabolomics analysis, immunoprecipitation and LC/MS/MS, flow cytometry and immunoblot analysis

**Supplementary Figures**

Supp. F1 – Supp. F7

**Supplementary Tables**

Supp. Table 1. scRNA-Seq enrichment analysis comparing Treg0/Treg1 and Tfh0/Tfh1 clusters

Supp. Table 2. scRNA-Seq gene set enrichment analysis using KEGG_Hallmark_C5GO

Supp. Table 3. LC/MS/MS analysis of IRF5 interacting partners from immunoprecipitation in Ramos B cells

Supp. Table 4. Tier 1 and Tier 2 metabolites from Mass Spectrometry Analysis

Supp. Table 5. Fold-change of metabolites from TCR stimulated vs. unstimulated WT and KO T cells

Supp. Table 6. Primers for qPCR analysis

**Database Files**

**Data file S1:** scRNAseq files will be made publicly available at GSE267271.

## Acknowledgements

We thank the Feinstein Flow Cytometry Core Facility for flow cytometry technical assistance, the CCP animal facility for assistance with breeding, The Metabolomics Innovation Centre Canada for metabolomics, and the Center for Advanced Proteomics Research, New Jersey Medical School Cancer Research Center (Rutgers). We thank Dr. Lionel Blanc for OPP reagents and techniques and Amy Pitler and Hong Li at Rutgers University for assistance in the LC/MS/MS analysis of IRF5 interacting partners by immunoprecipitation.

## Funding

Lupus Foundation of America Gina M. Finzi Fellowship (ZB)

National Institutes of health grant 1R01AR076242 (BJB)

National Institutes of health grant R03TR004623 (BJB)

Department of Defense (DoD) CDMRP Lupus Research Program grant W81XWH-18-1-0674 (BJB)

The Lupus Research Alliance (BJ.B)

## Author contributions

Conceptualization: ZB, BJB

Methodology: ZB, AL, BM, MM, BJB

Investigation: ZB, AL, AH, LB

Visualization: ZB, BJB

Funding acquisition: ZB, BJB

Project administration: BJB

Supervision: BJB

Writing - original draft: ZB

Writing – review and editing: ZB, BJB

## Data and materials availability

Raw and processed bulk scRNA-seq files are available from Gene Expression Omnibus (GEO) under accession number GSE267271. All other data are available in the main text or the supplementary materials.

## Supplementary Materials and Methods

### Detailed Metabolomics Analysis

#### Chemicals and Reagents

All chemicals and reagents were purchased from Sigma-Aldrich (Markham, ON, Canada), except those specifically stated. LC-MS grade water, acetonitrile, methanol and formic acid were purchased from Canadian Life Sciences (Peterborough, ON, Canada). Metabolome quantification kit for sample normalization and chemical isotope labeling kits for sample labeling were purchased from Nova Medical Testing Inc. (Edmonton, AB, Canada).

#### Normalization

Samples were randomized before any procedures to eliminate batch variations in sample analysis. Sample normalization was conducted by measuring the total metabolite concentration in each sample usingNova Medical Testing Inc (Product Number: NMT-6001-KT). According to quantification results, different volumes of sample were taken and adjusted to a final concentration of 1.2 mM. The samples were stored in an -80°C freezer and used for the following preparations and analyses.

#### LC-MS Analysis

Individual samples were labeled with ^12^C-reagents and a pooled sample, which was generated by mixing an aliquot from each individual sample, was labeled with ^13^C-reagents. The ^12^C_2_-labeled individual sample was mixed with ^13^C_2_-labeled reference sample in equal volumes. The mixture was injected into LC-MS for analysis. Prior to analysis, quality control (QC) samples were prepared by equal volume mix of a ^12^C-labeled and a ^13^C-labeled pooled sample. QC samples were run at an interval of one QC injection after 10 sample injections. The peak intensity ratio between ^12^C-peak and ^13^C-peak represents the relative quantification result for a specific metabolite in an individual sample. All LC-MS analyses were conducted on an Agilent 1290 LC linked to Bruker Impact II QTOF Mass Spectrometer. The column used was Agilent eclipse plus reversed-phase C18 column (150 × 2.1 mm, 1.8 µm particle size) and the column oven temperature was 40°C. Mobile phase A was 0.1% (v/v) formic acid in water and mobile phase B was 0.1% (v/v) formic acid in acetonitrile. The gradient setting was: t = 0 min, 25% B; t = 10 min, 99% B; t = 15 min, 99% B; t = 15.1 min, 25% B; t = 18 min, 25% B. The flow rate was 400 µL/min. Mass spectral acquisition rate was 1 Hz, with an m/z range from 220 to 1000.

### Data Processing and Metabolite Identification

LC-MS data was first converted to .csv files using DataAnalysis 4.4 (Bruker Daltonics). Files were uploaded to IsoMS Pro 1.2.12 (Nova Medical Testing Inc.) for data processing and metabolite identification. Peak pairs that originated from blank samples and those not presented in at least 80.0% of samples in any group were removed. Data was then normalized by Ratio of Total Useful Signal, calculated as sum of all useful ^12^C-peaks over sum of all useful ^13^C-peaks. Metabolite identification was carried out using a three-tiered approach against NovaMT Metabolite Databases 2.0 (Nova Medical Testing Inc.) (*113*). In tier 1, peak pairs were searched against a labeled metabolite library (CIL Library) based on accurate mass and retention time. In tier 2, the remaining peak pairs were searched against a linked identity library (LI Library). In tier 3, any remaining peak pairs were searched based on accurate mass match against the MyCompoundID (MCID) library (www.mycompoundid.org) (*114*).

### Immunoprecipitation and LC/MS/MS

Briefly, 250μg lysate of unstimulated Ramos B cells was used for immunoprecipitation with either anti-IRF5 or anti-Ig control antibodies (*16*). Samples were processed via 1D gel and in-gel tryptic digestion resulting in peptides fractionated by reversed phase chromatography and analysis by LC/MS/MS on a LTP Orbitrap Velos mass spectrometer at the Center for Advanced Proteomics Research, New Jersey Medical School Cancer Research Center. Database search was performed using Mascot search engine against SwissPRot human protein database. Listed proteins (**Suppl. Table 3**) had at least 1 unique peptide identified with ≥95% identification probability.

### Flow cytometry

The following antibodies were used for surface staining as indicated: AF700-conjugated anti-CD3 (Clone 17A2; Biolegend); BV711-conjugated anti-CD3 (Clone 17A2; Biolegend) BV510-conjugated anti-CD4 (Clone RM4-5; Biolegend); PerCP-conjugated anti-CD45 (Clone 30-F11; Biolegend), PE/CY7-conjugated anti-CD45 (Clone 30-F11; Biolegend), PE/CY7-conjugated anti-CD25 (Clone PC61; Biolegend), PerCP-conjugated anti-CD45R(B220) (Clone RA3-6B2; Biolegend), AF647-conjugated anti-CD38(B220) (Clone 90; Biolegend), BV421-conjugated anti-CD86 (Clone GL-1; Biolegend), FITC-conjugated anti-CD40 (Clone 3/23; Biolegend), Pe-Cy7-conjugated anti-95(Fas) (Cat#: 152617; Biolegend), Pe-Cy5.5-conjugated anti-CXCR4) (Clone#: 2B11; eBioscience), APC/Cy7-conjugated anti-CD69 (Clone#: H1.2F3; Biolegend), BV421-conjugated anti-FoxP3 (Clone#: MF-14; Biolegend), PE-conjugated anti-CD40L(Clone#: 24-31; eBioscience), anti-Asct2 (V501) (Cat#: 5345S; Cell Signaling Technology), and anti-SLC38A2 (Cat#: BS-12125R; Bioss Antibodies).

The following antibodies were used for intracellular staining: PE-conjugated anti-mTOR(7C10) (Cat# 15006S; Cell Signaling Technology), anti-phospho-S6 Ribosomal Protein(Ser235/236) (Cat#: 4857S; Cell Signaling Technology), anti-phospho-Akt(Ser473) (Cat#: 9271T, Cell Signaling Technology), anti-S6 (Cat#2217S; Cell Signaling).

The following secondary antibodies were used: FITC-conjugated anti-Rabbit IgG (Cat#: 65-611-1; Invitrogen), PE-conjugated anti-Rabbit IgG, (Cat#: 12-4739-81, eBioscience).

For dead cell exclusion, either Live/Dead Fixable Yellow Dead Cell Stain (Cat#: 50-112-1528, Thermo Fischer) or Live/Dead Fixable Green Dead Cell Stain (Cat#: L23101, Thermo Fischer)) was used in each experiment.

### Immunoblot Analysis

The following antibodies were used as indicated.β-Actin HRP conjugate (Cell Signaling, Cat #: 12620, 1:5000), GLS recombinant antibody (Proteintech, Cat #: 81486-1-RR, 1:1000), GLUD1 monoclonal antibody (Cat #: 67026-1-lg, 1:1000), ATF-4 (Cell Signaling, Cat #: 11815, 1:1000), eEF2 (Cell Signaling, Cat #: 2332, 1:1000).

---

**Fig. S1.**
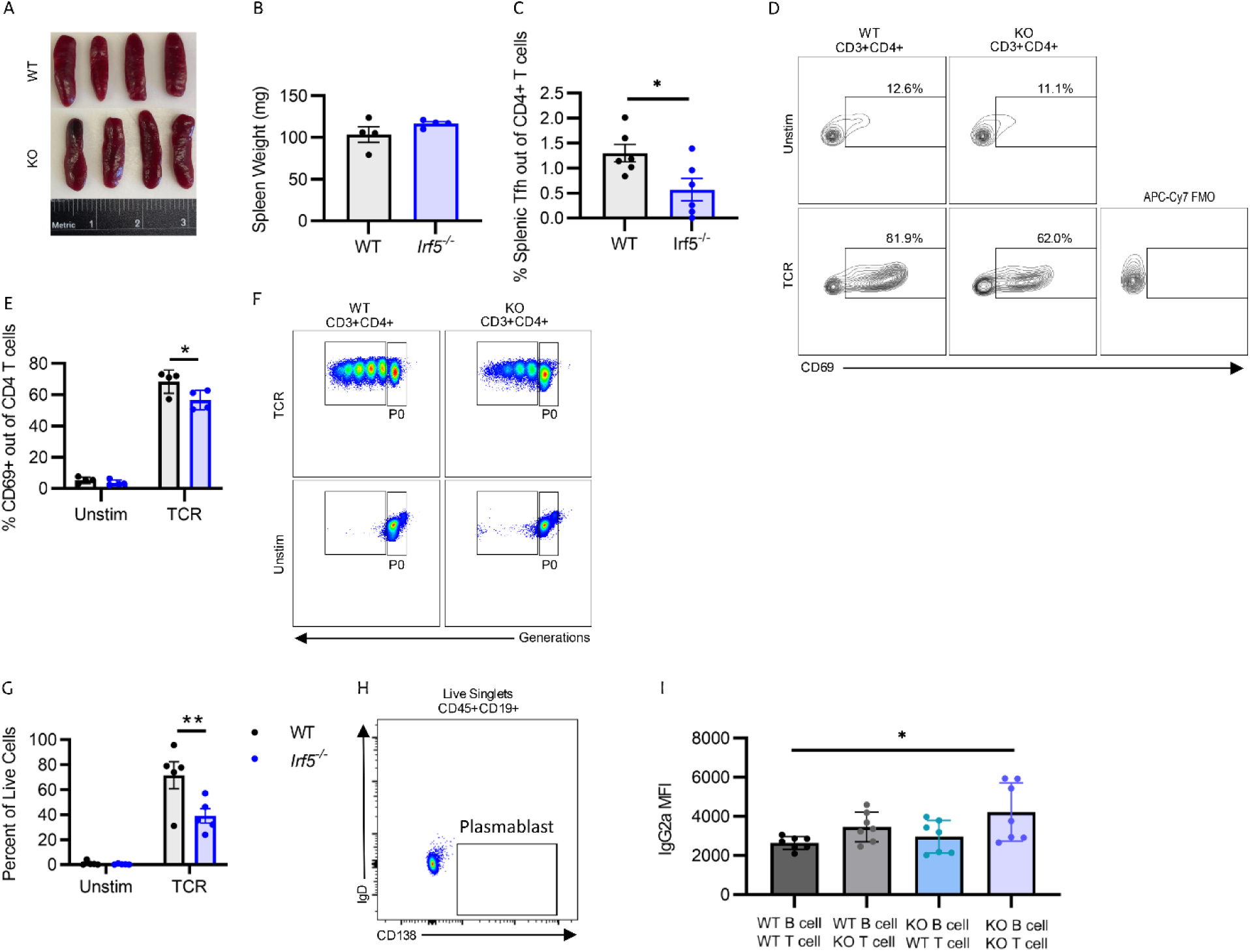
(**A**) Spleen images and (**B**) weights from age and gender matched WT and *Irf5^−/−^*mice. (**C**) Tfh percentages from splenocytes of unimmunized mice as determined by flow cytometry. Six biological replicates. *P* value calculated by two-way unpaired *t* tests. (**D-E**) Sorted CD3^+^CD4^+^ WT and *Irf5^−/−^*T cells were activated for 24-hours *in vitro* with anti-CD3/CD28 (TCR). (**D**) Representative flow plots and (**E**) summary graphs of frequency of CD69 expression out of live CD3^+^CD4^+^ T cells. Four biological replicates, representative of three independent experiments. (**F-G**) Sorted CD3^+^CD4^+^ WT and *Irf5^−/−^*T cells were activated for 24-hours *in vitro* with anti-CD3/CD28 (TCR). (**F**) Representative flow plots and (**G**) summary graphs of T cell proliferation. Five biological replicates. *P* value calculated by 2-way ANOVA with Holm-Sidak correction for multiple comparisons. (**H**) Control gating strategy for plasmablasts (CD45^+^CD19^+^CD138^+^IgD^low^). (**I**) Summary graphs of IgG2a production. *P* value calculated by 1-way ANOVA with Tukey correction for multiple comparisons. *p < 0.05, **p < 0.01. Each point represents an independent biological replicate. Bar graphs show means +/- SEMs.

**Fig. S2.**
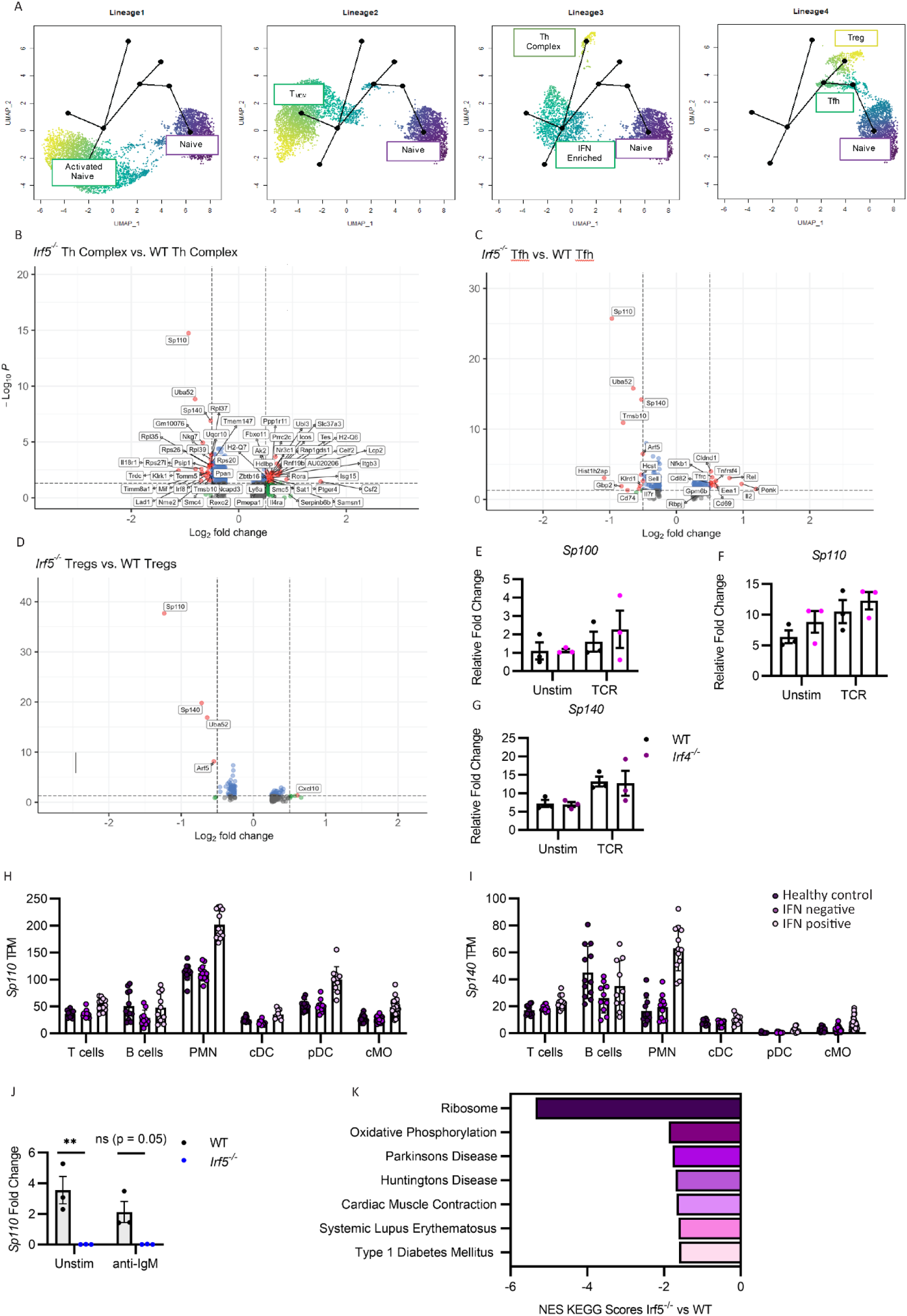
(**A**) Pseudotime trajectory analysis performed using Slingshot. Depiction of four predicted trajectories. Differential transcript expression in (**B**) *Irf5^−/−^*vs WT Th Complex cluster (**C**) *Irf5^−/−^* vs WT Tfh cluster and (**D**) *Irf5^−/−^* vs WT Treg cluster. (**E-G**) Sorted CD3^+^CD4^+^ T cells from WT and *Irf4^−/−^* splenocytes were activated for 6-hours *in vitro* with anti-CD3/CD28 (TCR). (**E**) *Sp100*, (**F**) *Sp110* and (**G**) *Sp140* transcript expression normalized to *β-Actin.* Three biological replicates. *P* value calculated by 2-way ANOVA with Holm-Sidak correction for multiple comparisons. All comparisons not significant. (**H**) *Sp110* and (**I**) *Sp140* transcripts per million (TPM) in Healthy Control, Interferon (IFN) negative and IFN positive systemic lupus erythematosus (SLE) patient peripheral blood mononuclear cells (PBMCs) analyzed from published scRNAseq dataset (PMC8015858). (**J**) *Sp110* transcript expression normalized to *β-Actin* in sorted CD45^+^B220^+^ WT and *Irf5^−/−^* B cells stimulated for 6-hours with anti-IgM. Three biological replicates per genotype. *P* value calculated by 2-way ANOVA with Holm-Sidak correction for multiple comparisons. (**K**) Pre-ranked gene set enrichment analyses (GSEA) using Kyoto Encyclopedia of Genes and Genomes (KEGG) C2 gene sets. Top 7 significantly enriched pathways sorted by Normalized Enrichment Score (NES) (p < 0.05). Representative graph of two WT and two *Irf5^−/−^* biological replicates, with one male and one female representing each genotype. Bar graphs show means +/- SEMs. **p < 0.01, ns = not significant. Each point represents an independent biological replicate.

**Fig. S3.**
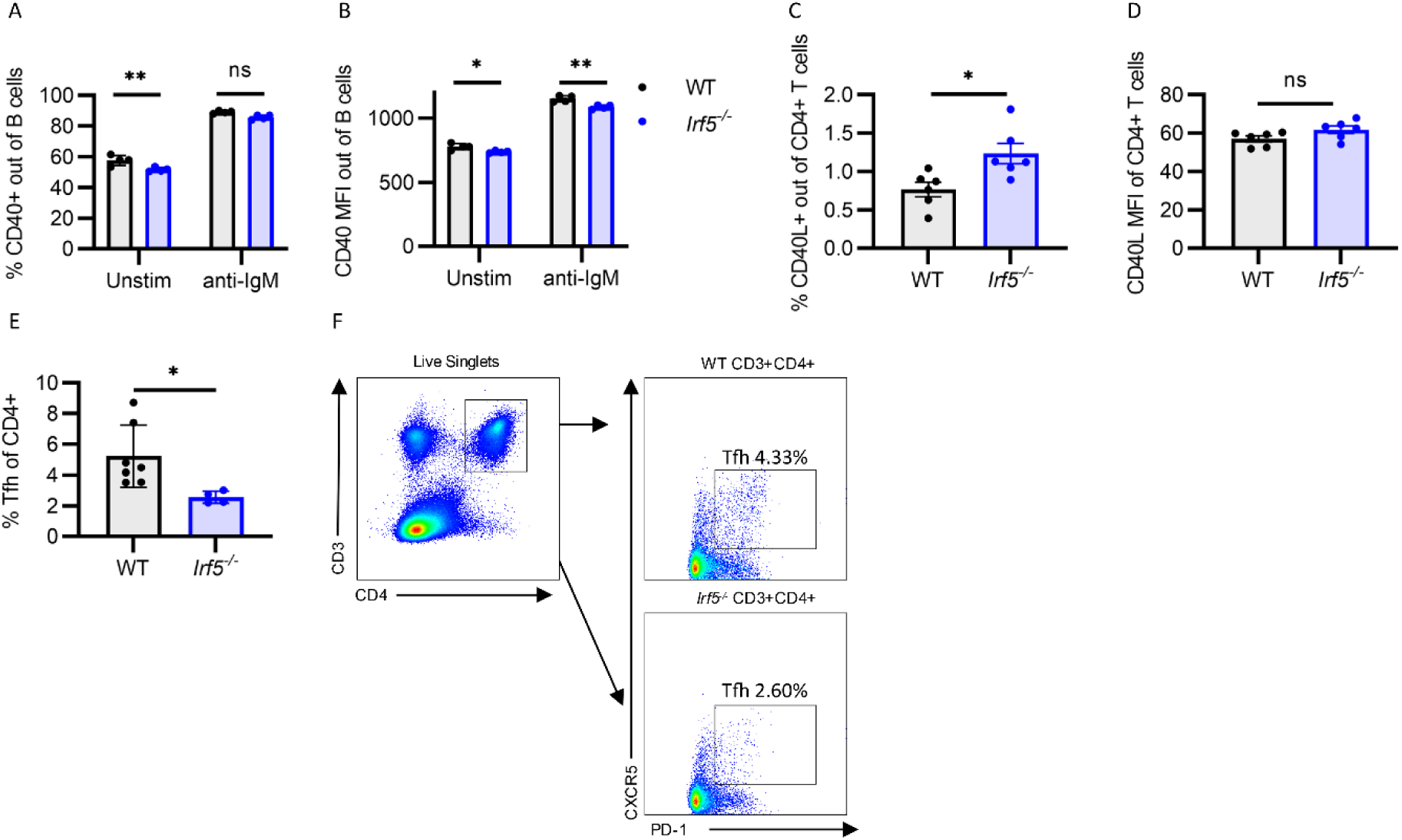
(**A,B**) WT and *Irf5^−/−^* total splenocytes were activated for 24-hours *in vitro* with anti-IgM. Summary graphs of (**A**) CD40% expression and (**B**) CD40 mean fluorescence intensity (MFI) in CD45^+^B220^+^ B cells. Four biological replicates, representative of two independent experiments. *P* value calculated by 2-way ANOVA with Holm-Sidak correction for multiple comparisons. CD40L (**C**) percent expression and (**D**) MFI in CD3^+^CD4^+^ T cells gated from WT and *Irf5^−/−^* total splenocytes. Six biological replicates. (**E**) Summary graphs of Tfh (CD3^+^CD4^+^PD1^+^CXCR5^+^) quantification and (**F**) representative flow cytometric gating. Four to six biological replicates. *P* value calculated by two-way unpaired *t* test.*p < 0.05, **p < 0.01, ns = not significant.

**Fig. S4.**
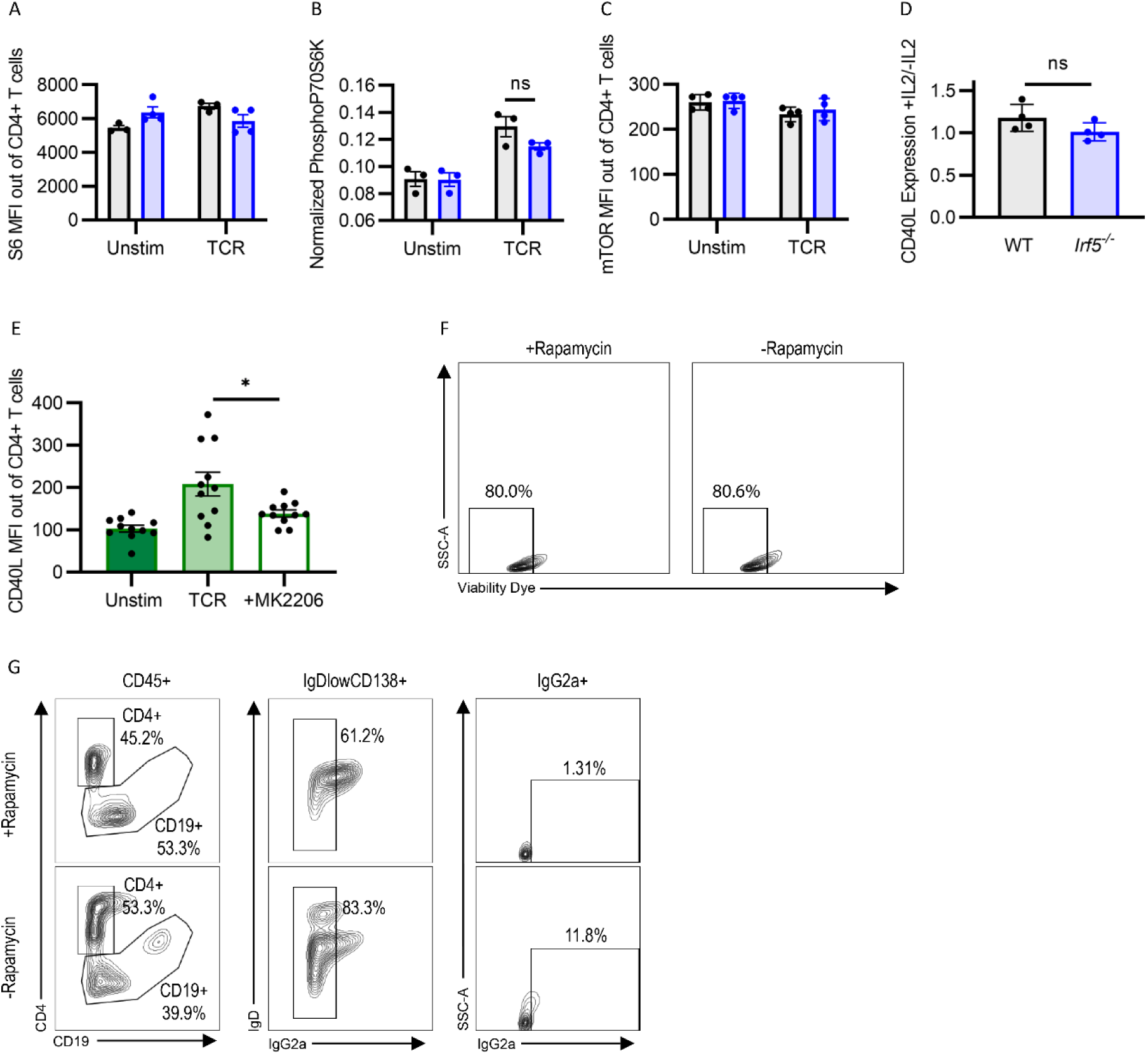
(**A-C**) WT and *Irf5^−/−^* total splenocytes were activated for 24-hours *in vitro* with anti-CD3/CD28 (TCR). Summary graphs of indicated intracellular protein expression and/or phosphorylation levels in CD3^+^CD4^+^ T cells gated from total splenocytes. Three-five biological replicates, representative of a minimum of two independent experiments. *P* value calculated by 2-way ANOVA with Holm-Sidak correction for multiple comparisons. (**D**) WT and *Irf5^−/−^* total splenocytes were activated for 24-hours *in vitro* with anti-CD3/CD28 (TCR) in the presence (+IL2) or absence (-IL2) of IL2. Summary graphs of relative CD40L expression comparing IL2/TCR (+Il2) stimulated vs TCR stimulated (-IL2) CD4^+^ T cells. *P* value calculated by two-way unpaired *t* test. (**E**) WT and *Irf5^−/−^*total splenocytes were activated for 24-hours *in vitro* with anti-CD3/CD28 (TCR) in the presence (+MK2206) or absence (TCR) of the ASCT2 inhibitor, MK2206. Summary graphs of CD40L expression on live CD3^+^CD4^+^ T cells gated from total splenocytes. Data pooled from three independent experiments. *P* value calculated by 1-way ANOVA with Tukey correction for multiple comparisons. (**F,G**) WT and *Irf5^−/−^* total splenocytes were activated for 24-hours *in vitro* with anti-CD3/CD28 (TCR) in the presence (+Rapamycin) or absence (-Rapamycin) of Rapamycin. (**F**) Representative flow cytometric gating of CD3^+^CD4^+^ T cell viability using cell permeable viability dye. (**G**) Representative gating strategies for CD45^+^CD19^+^CD138^+^IgD^low^ PBs gated from CD4 T cell:B cell cocultures with (+Rapamycin) and without (-Rapamycin) CD4 T cell Rapamycin pretreatment. *p < 0.05, ns = not significant. Each point represents an independent biological replicate. Bar graphs show means +/- SEMs.

**Fig. S5.**
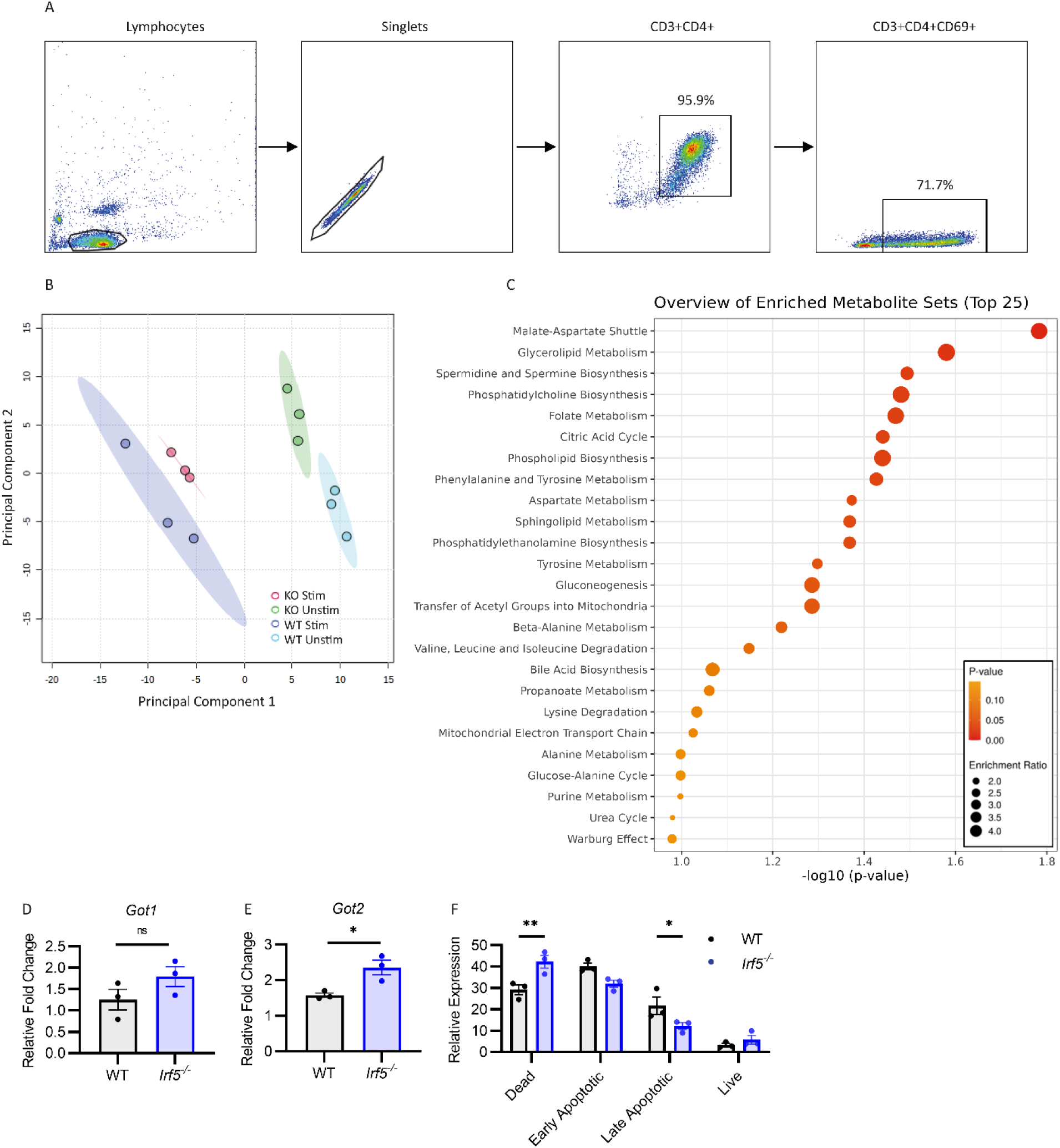
(**A**) Representative flow cytometry gating strategy confirming the purity and activation of naïve CD3^+^CD4^+^ T cells sorted from total splenocytes. (**B**) Principal component analysis from metabolomics profiling following 24-hours IL7 (Unstim) and anti-CD3/CD28 (Stim) treated CD3^+^CD4^+^ T cells sorted from total splenocytes of WT and *Irf5^−/−^* (KO) mice. Each condition is represented by three independent biological replicates. (**C**) Top 25 significantly enriched metabolite sets. (**D, E**) Sorted CD3^+^CD4^+^ T cells from WT and KO splenocytes were activated for 6-hours *in vitro* with anti-CD3/CD28 (TCR). *Got1* and *Got2* transcript expression normalized to *β-Actin*. *P* value calculated by two-way unpaired *t* test. (**F**) Summary graphs of CD4 T cell apoptosis following AOAA treatment. 3 independent biological replicates. *P* value calculated by 2-way ANOVA with Holm-Sidak correction for multiple comparisons. Bar graphs show means +/- SEMs. *p < 0.05, **p < 0.01, ns = not significant. Each point represents an independent biological replicate.

**Fig. S6.**
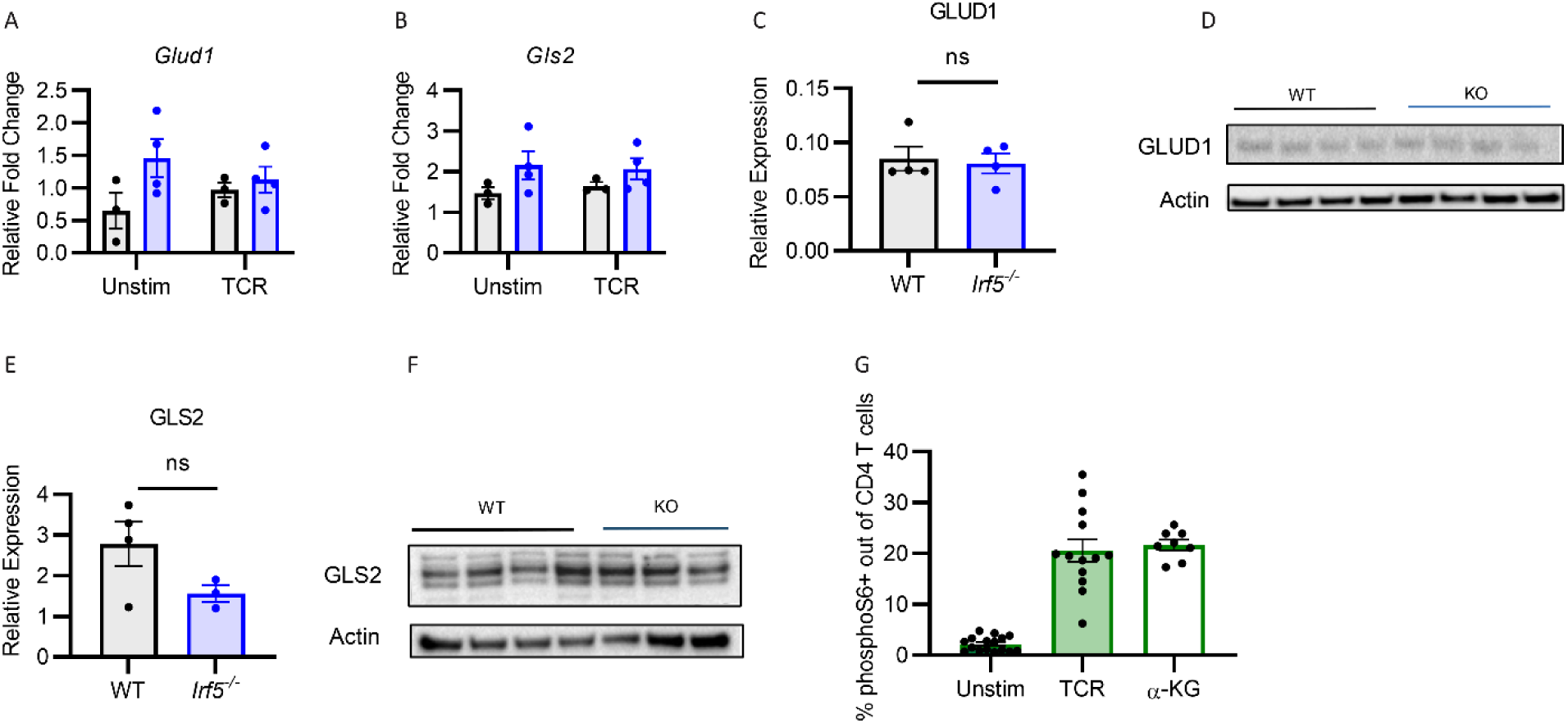
(**A,B**) Sorted CD3^+^CD4^+^ T cells from WT and *Irf5^−/−^*splenocytes were activated for 6-hours *in vitro* with anti-CD3/CD28 (TCR). (**A**) *Glud1* and (**B**) *Gls2* transcript expression normalized to *β-Actin.* Four biological replicates. *P* value calculated by 2-way ANOVA with Holm-Sidak correction for multiple comparisons. All comparisons not significant. (**C-F**) Purified CD3^+^CD4^+^ T cells were stimulated for 24 hours with anti-CD3/CD28, then analyzed with immunoblot for (**C, D**) GLUD1 (glutamate dehydrogenase) and (**E, F**) GLS (glutaminase). Values representative of 3-4 independent biological replicates normalized to Actin. *P* value calculated by two-way unpaired *t* test. (**G**) WT and *Irf5^−/−^* total splenocytes were activated for 24-hours *in vitro* with anti-CD3/CD28 (TCR) in the presence (DMK) or absence (TCR) of dimethyl ketoglutarate. Summary graphs of phosphorylated (phospho) S6 expression in CD3^+^CD4^+^ T cells. Data pooled from three independent experiments. *P* value calculated by 1-way ANOVA with Tukey correction for multiple comparisons. ns = not significant. Bar graphs show means +/- SEMs. Each point represents an independent biological replicate.

**Fig. S7.**
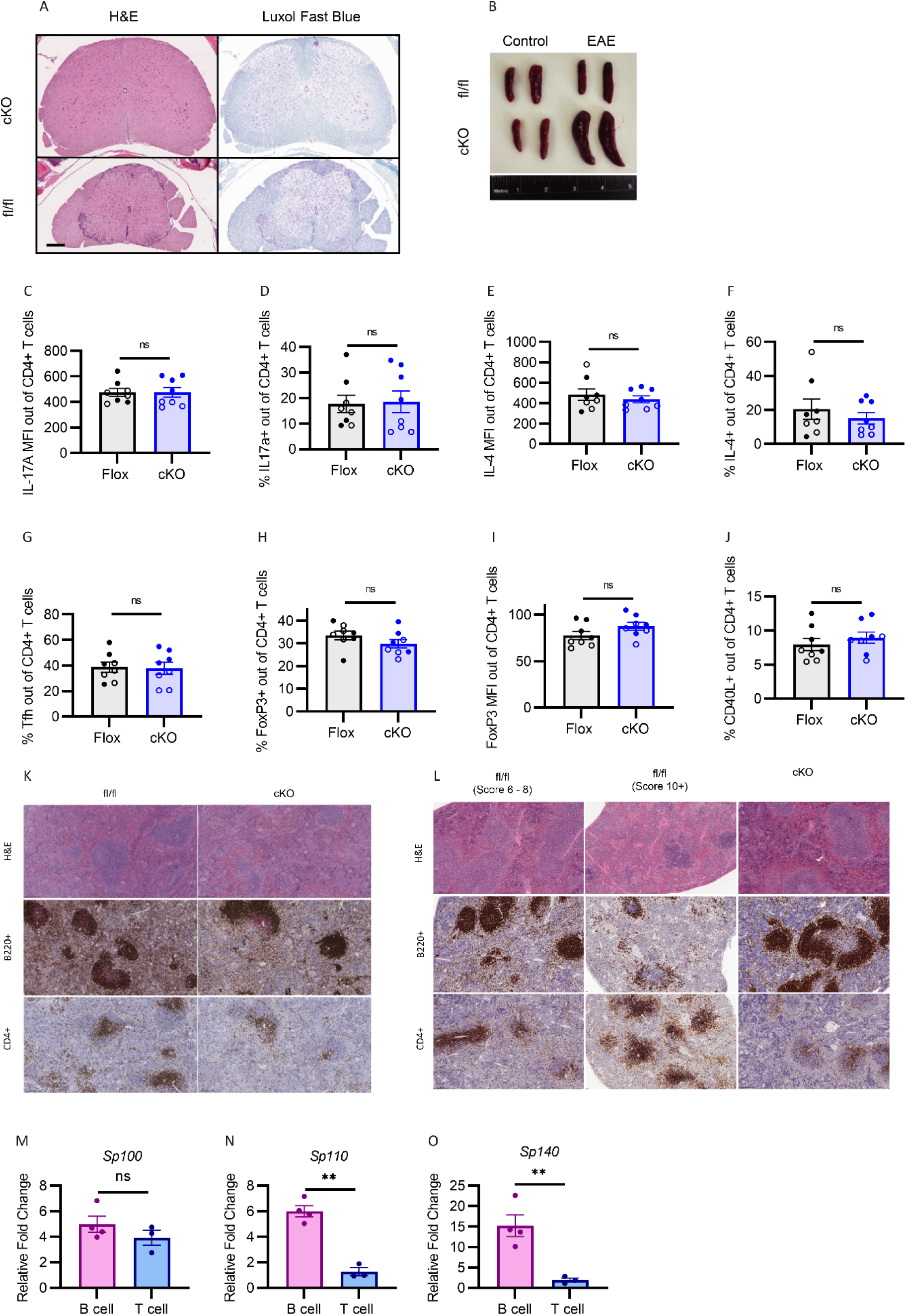
(**A**) Spinal cord cross sections stained with H&E and Luxol Fast Blue from cKO and fl/fl mice at peak disease. (**B**) Representative images of spleens from control and experimental cKO and fl/fl mice at clinically determined disease end points. Summary graphs of (**C, D**) IL-17expression, (**E, F**) IL4 expression, (**G**) CD4^+^CXCR5^+^BCL6^+^ Tfh, (**H, I**) CD4^+^FoxP3^+^ Tregs, and (**J**) CD40L expression following EAE induction. Clear circles represent day 13 post immunization, filled circles represent day 20 post immunization. (**K,L**) Representative images of spleen sections stained with H&E, anti-B220, and anti-CD4 at (**K**) day 13 and (**L**) day 20 post-immunization. On day 20, spleens were harvested from mice with varying disease severity as indicated by respective scores. (**M-O**) B cells and CD4 T cells were sorted from cKO mice and stimulated for 6 hours with anti-CD3/CD28 Dynabeads. Transcript levels of (**M**) *Sp100*, (**N**) *Sp110* and (**O**) *Sp140* normalized to *β-Actin*. Bar graphs show means +/- SEMs. *P* value calculated by two-way unpaired *t* test. **p < 0.01, ns = not significant. Each point represents an independent biological replicate.

**Supplemental Table 1.**
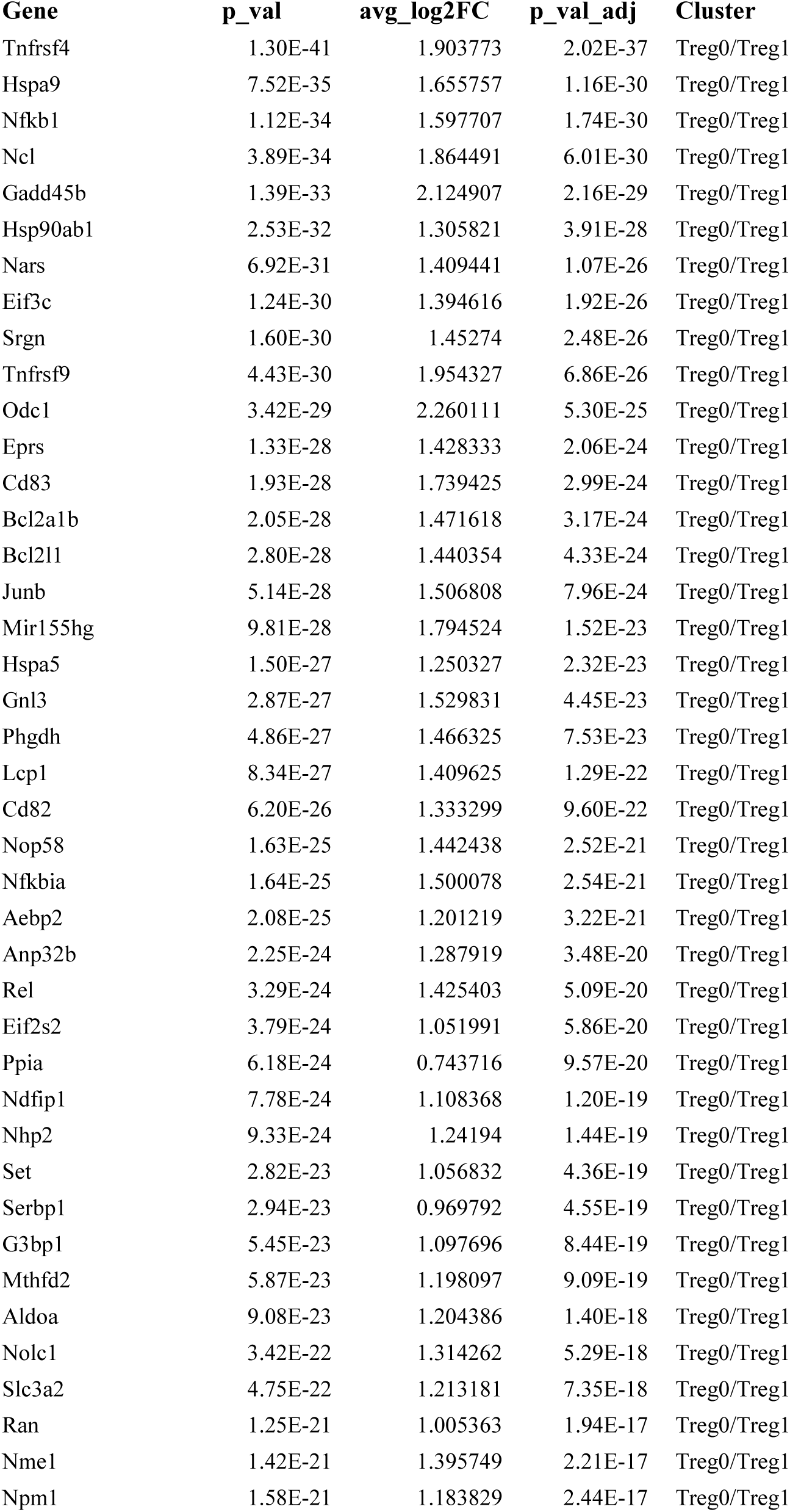
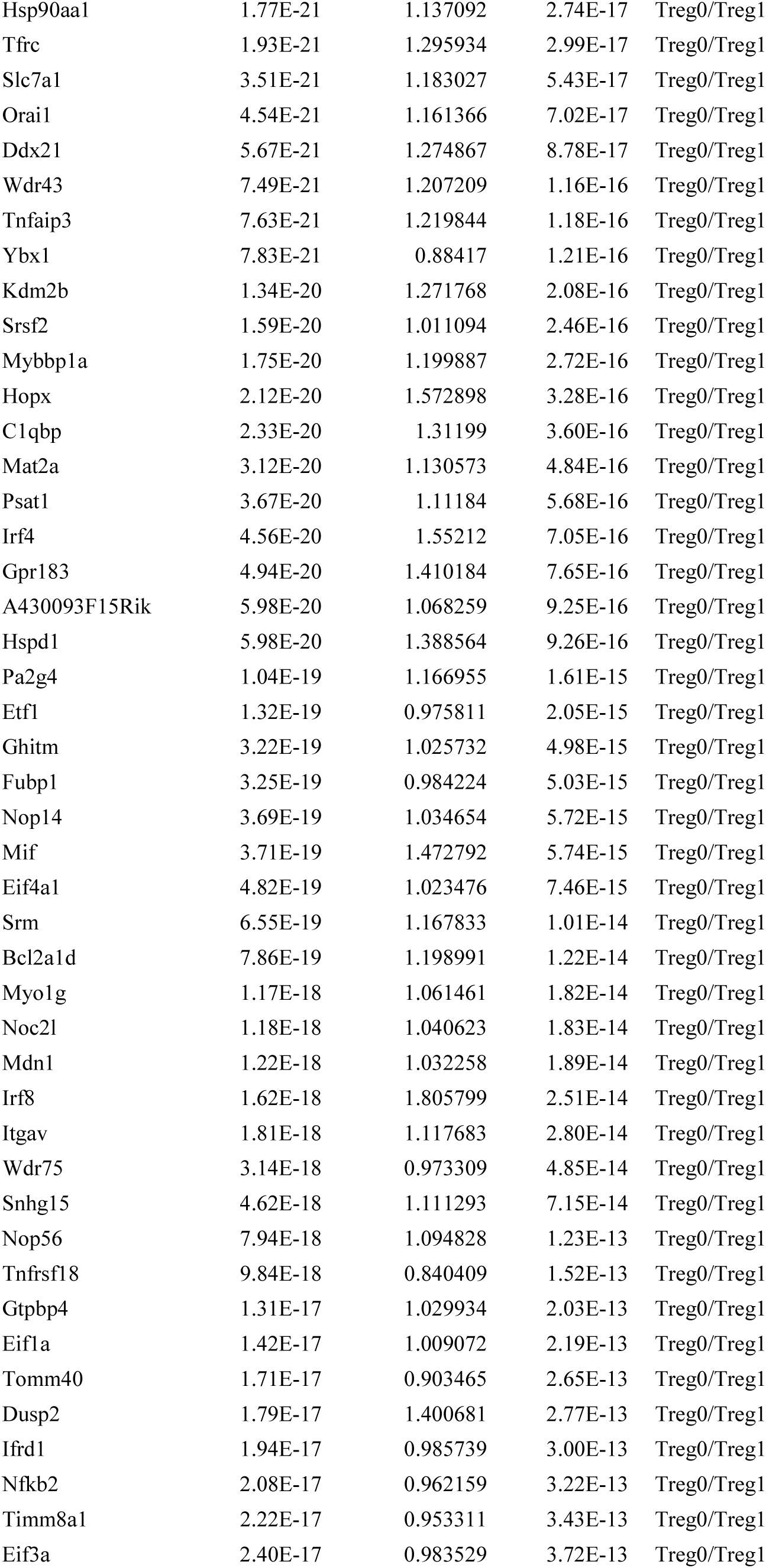
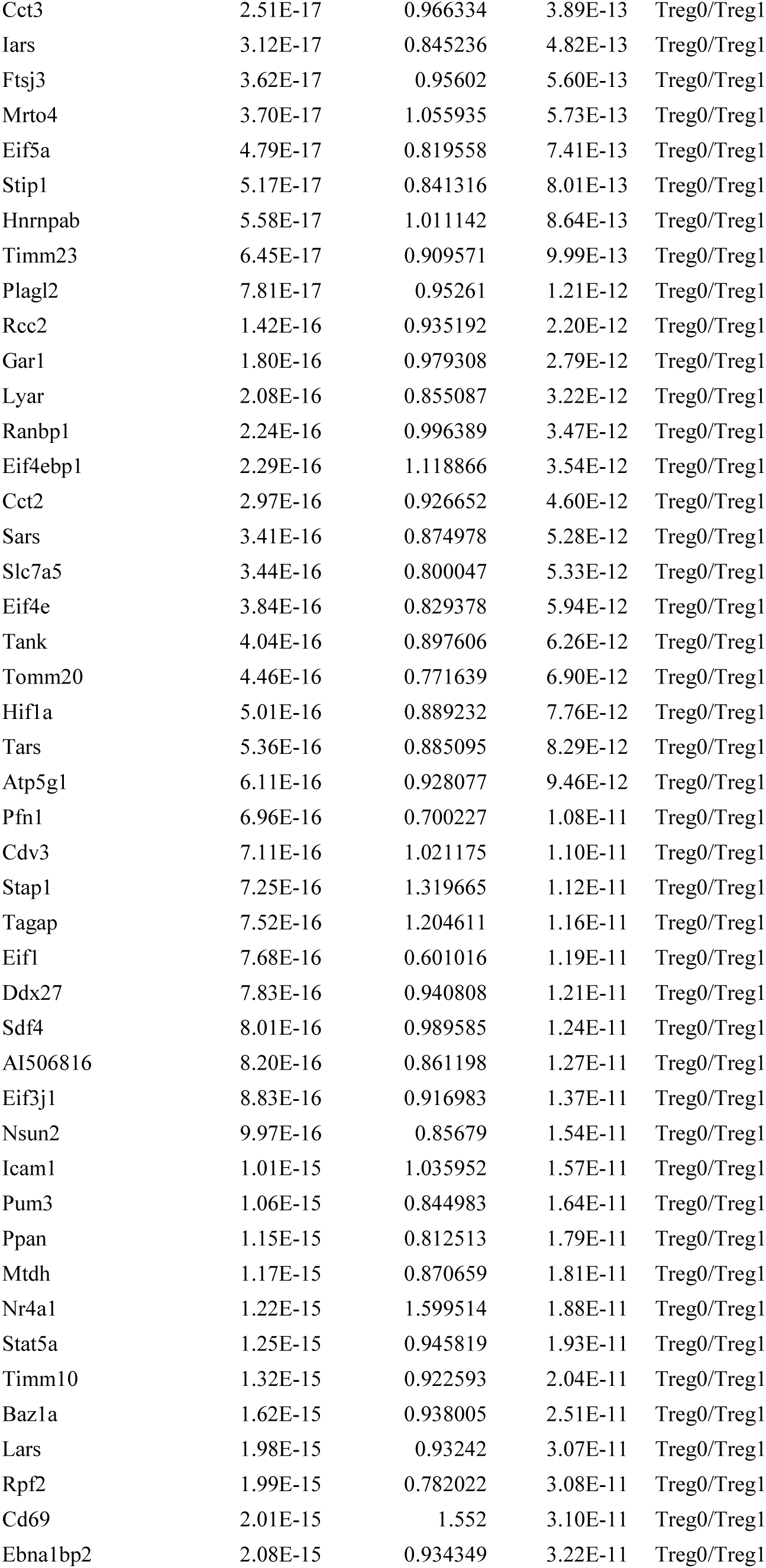
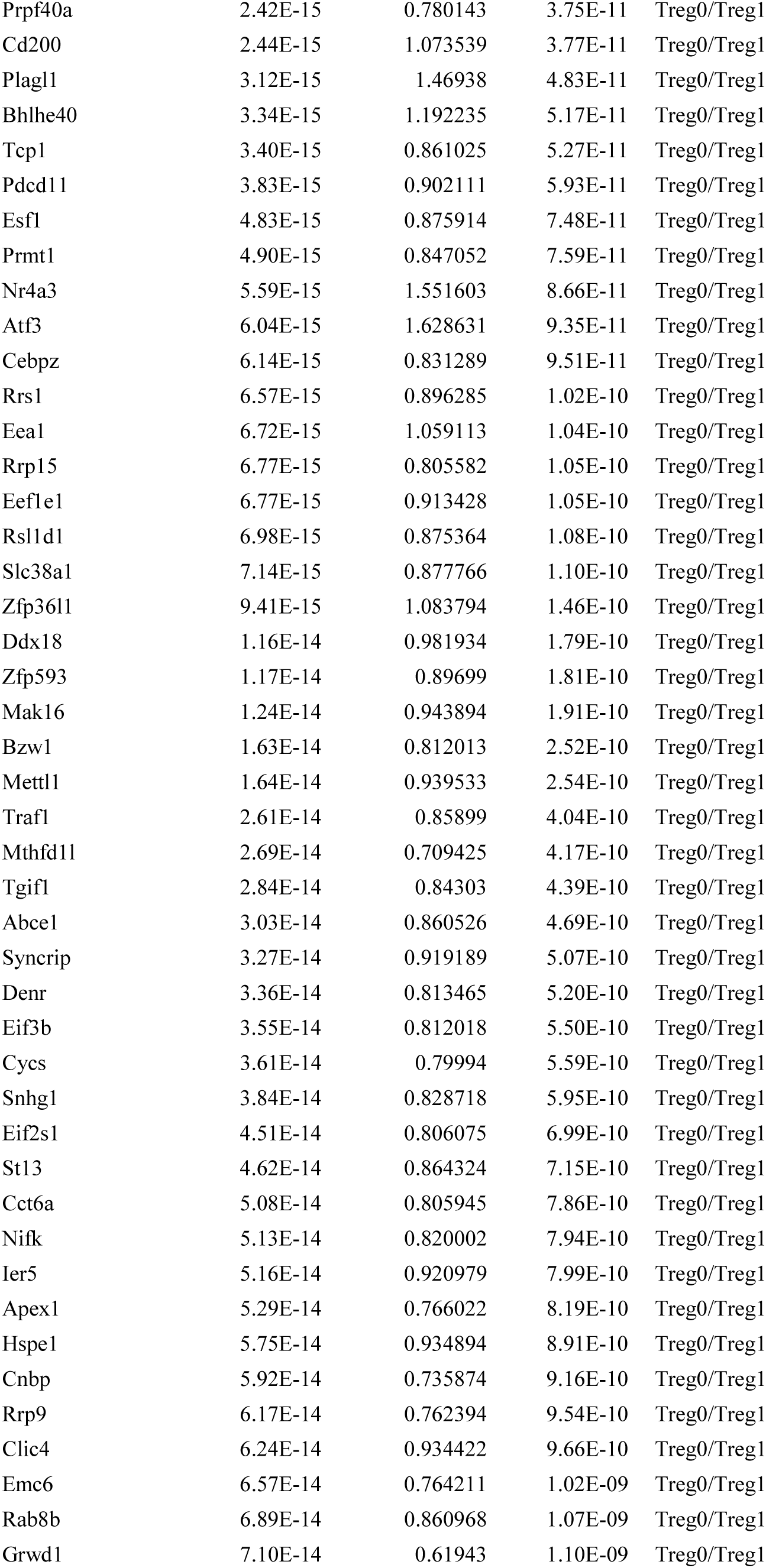
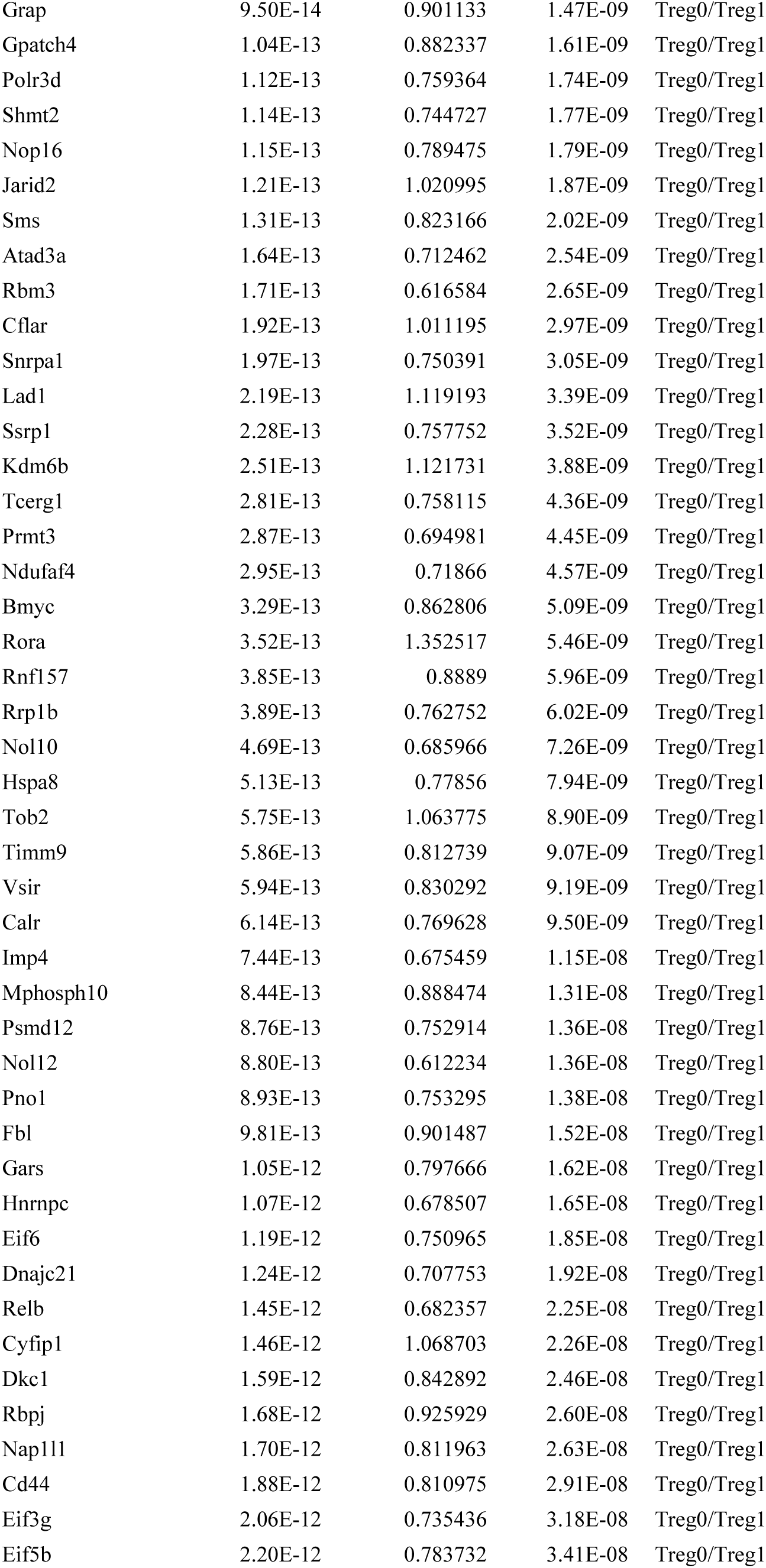
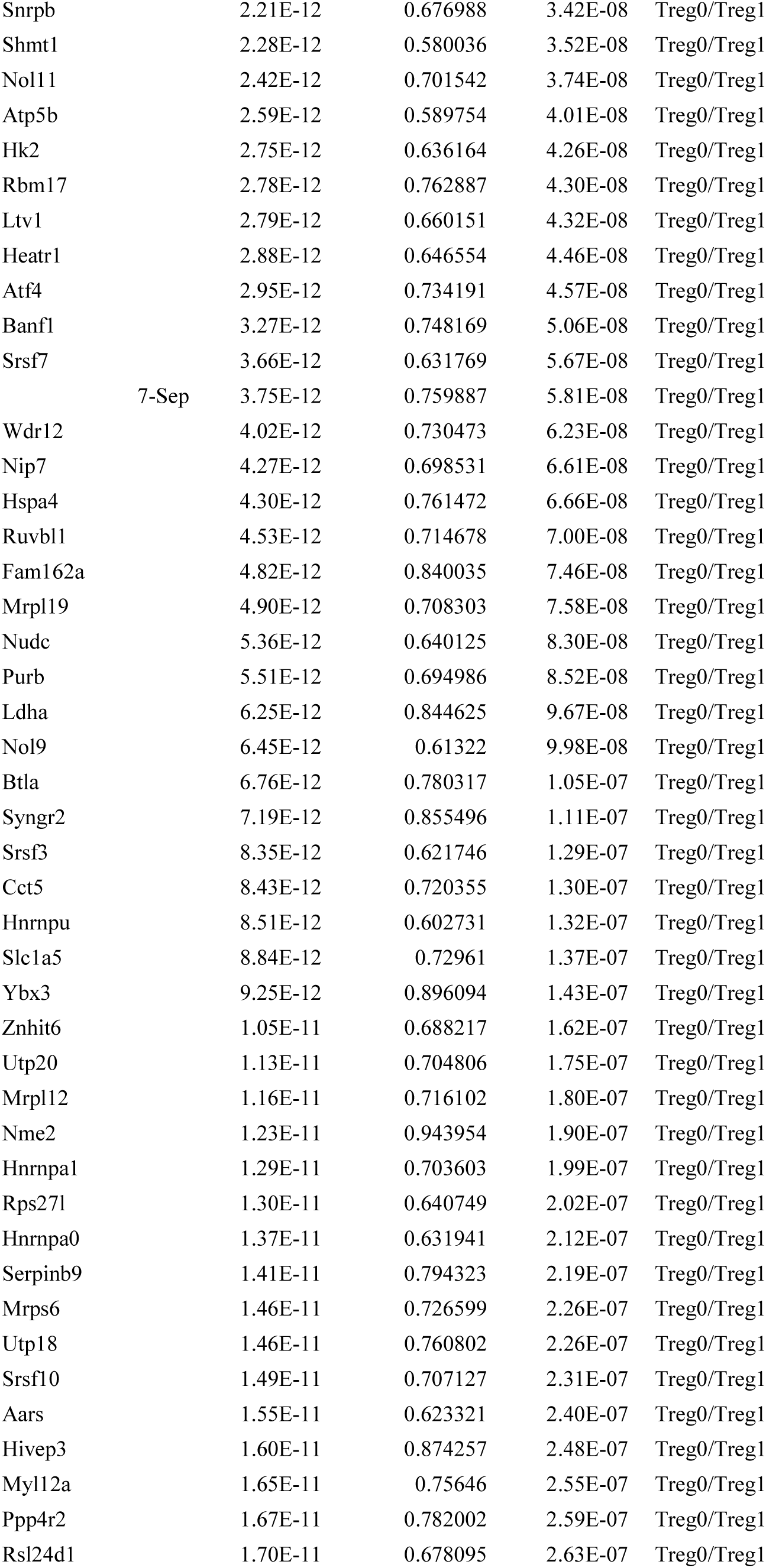
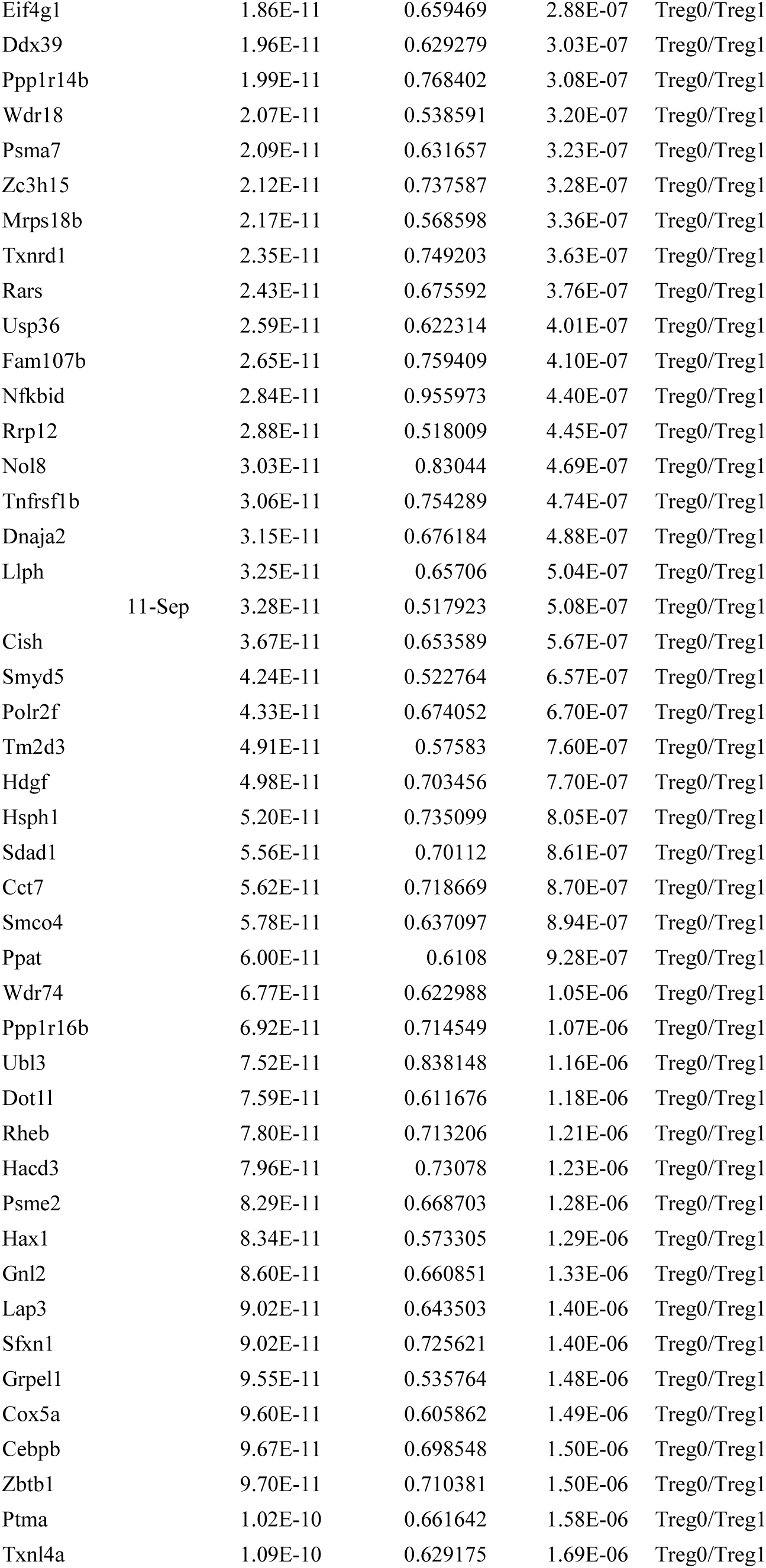
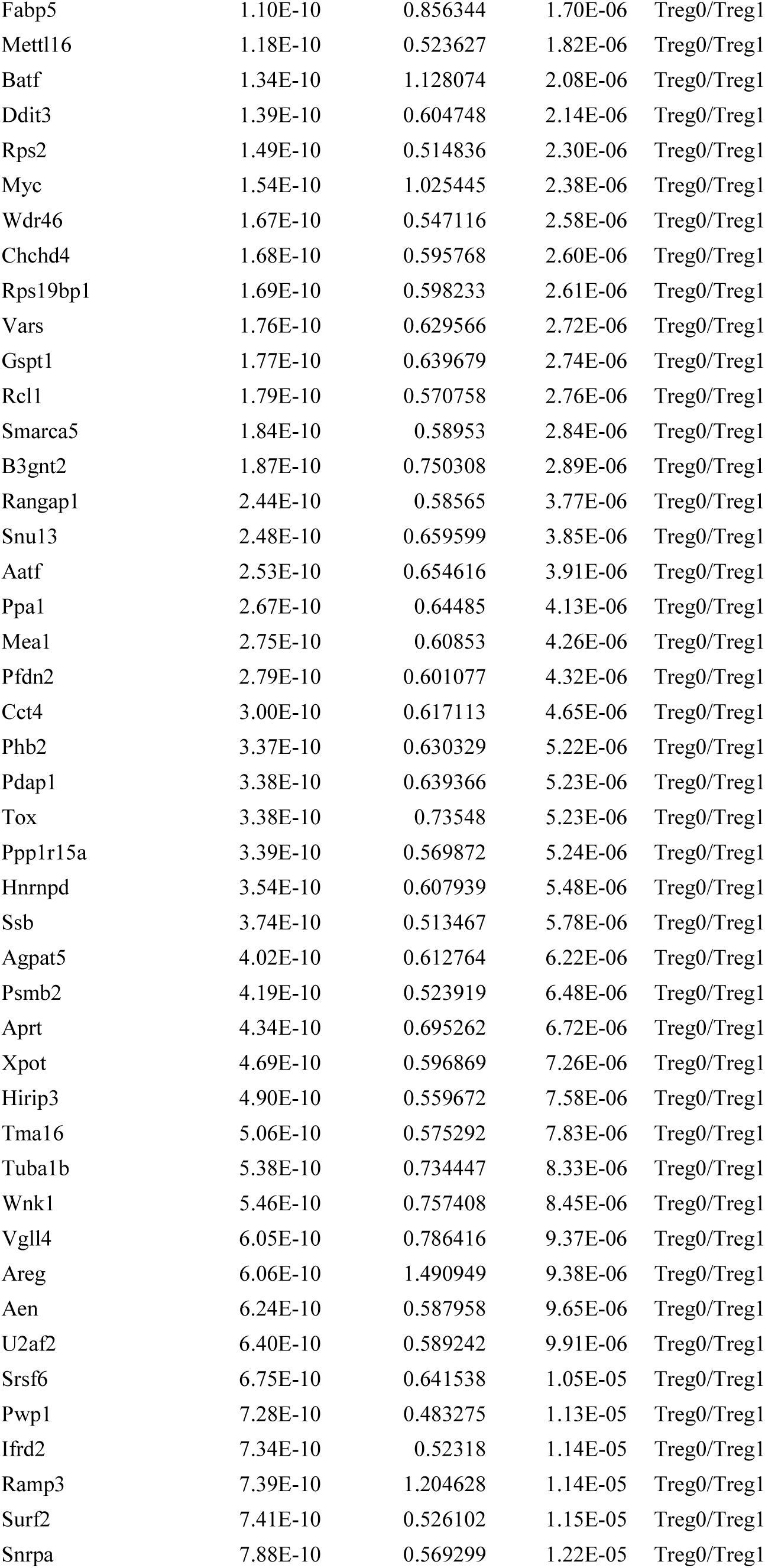
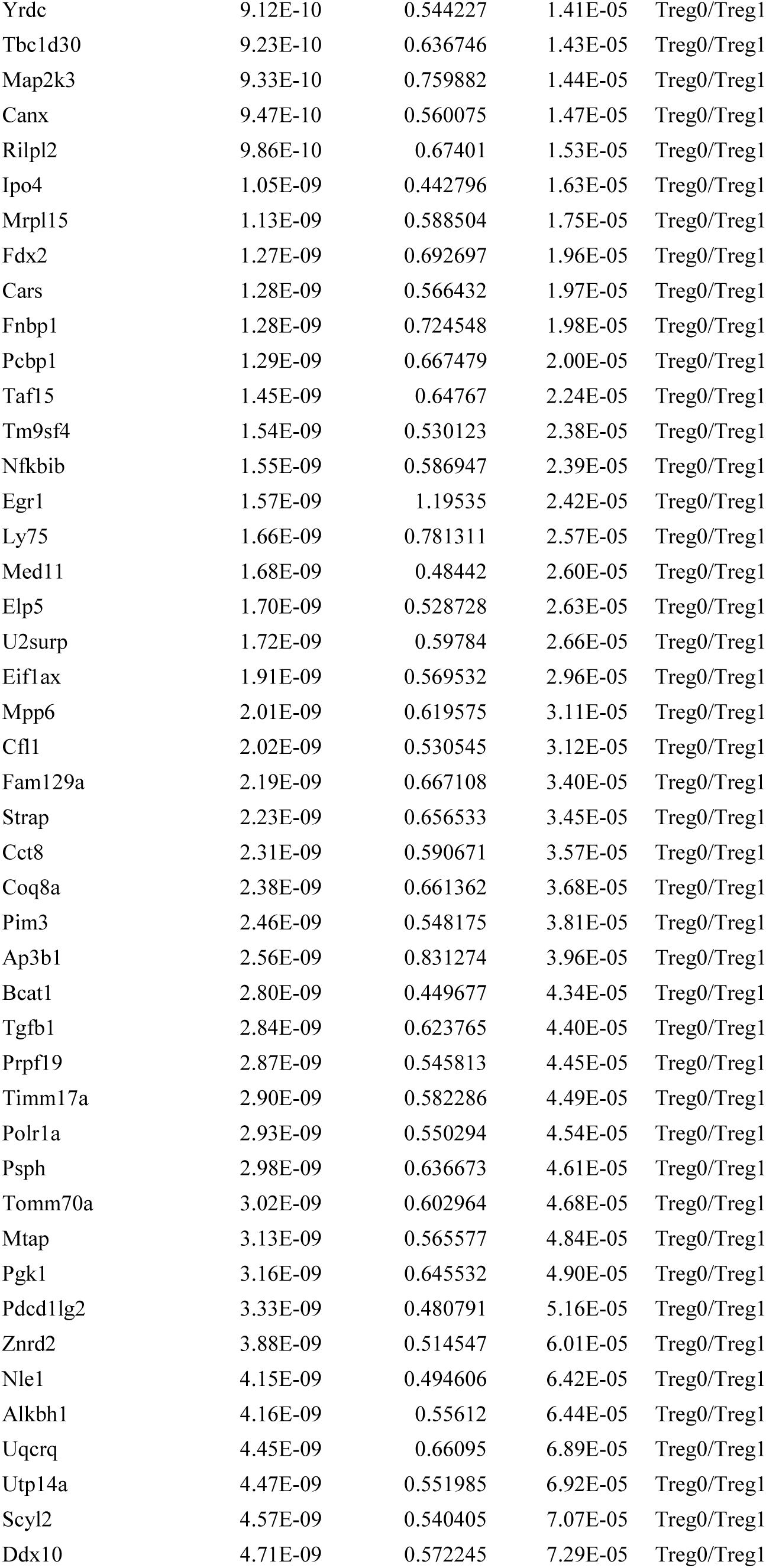
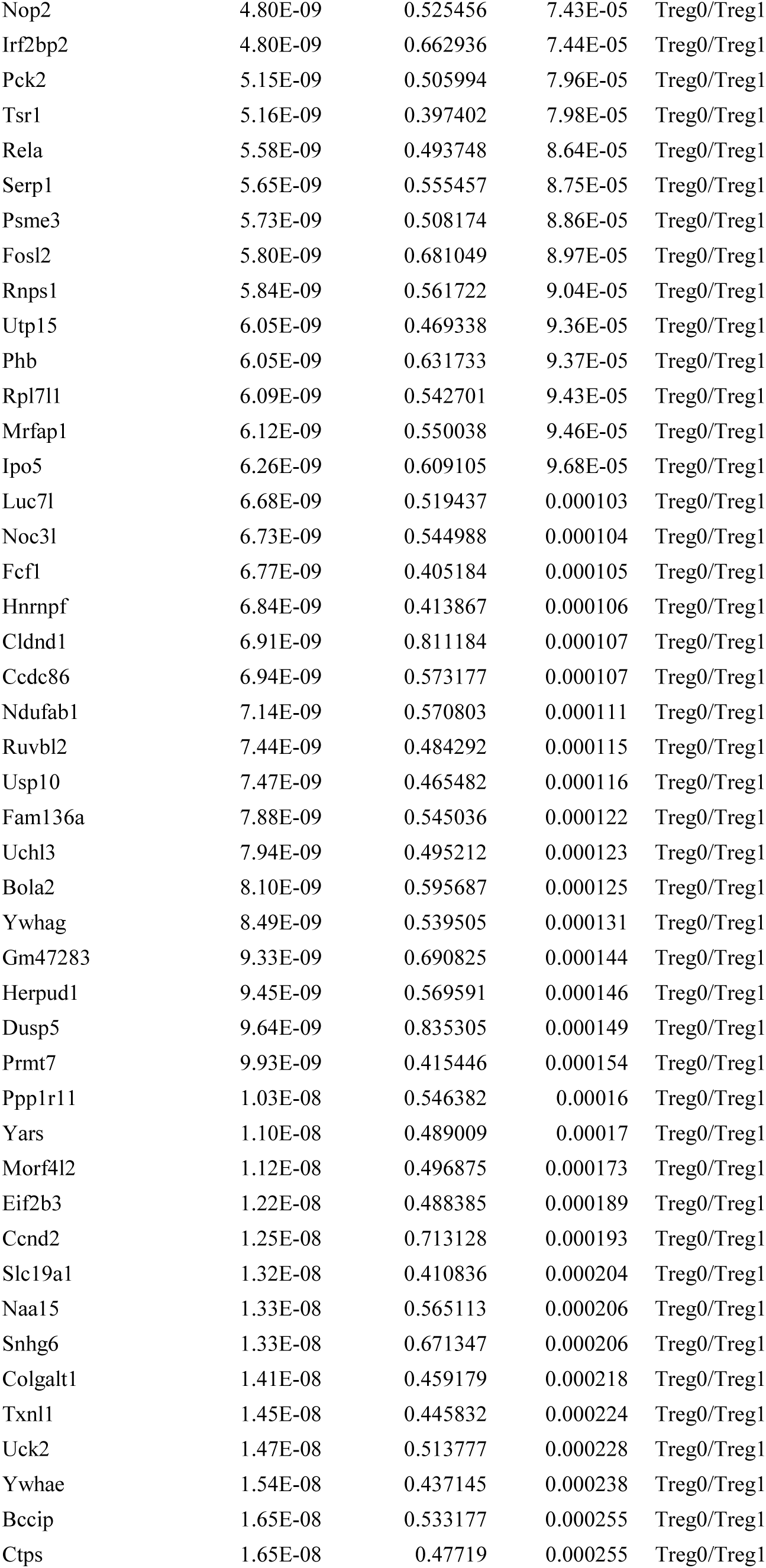
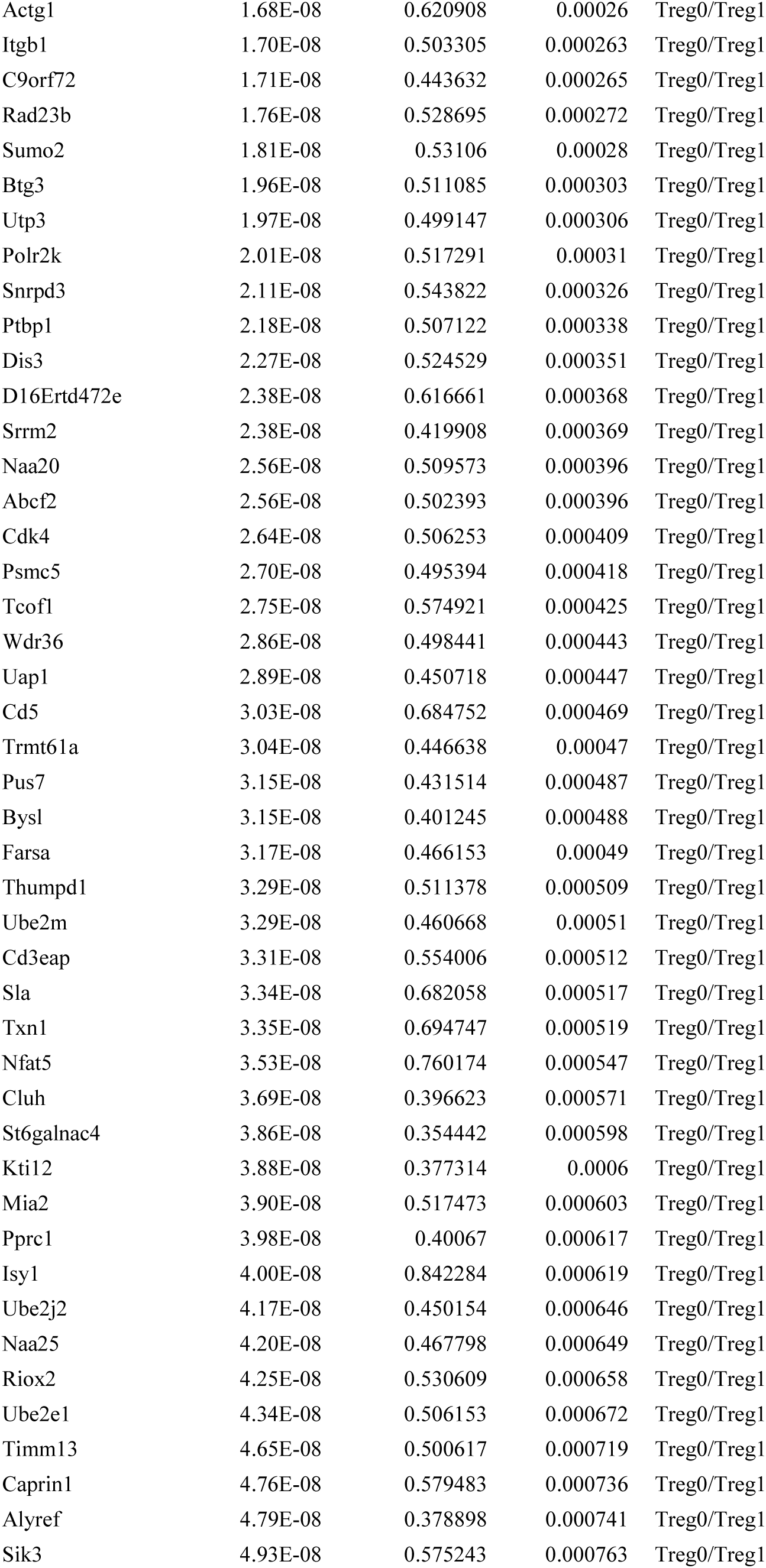
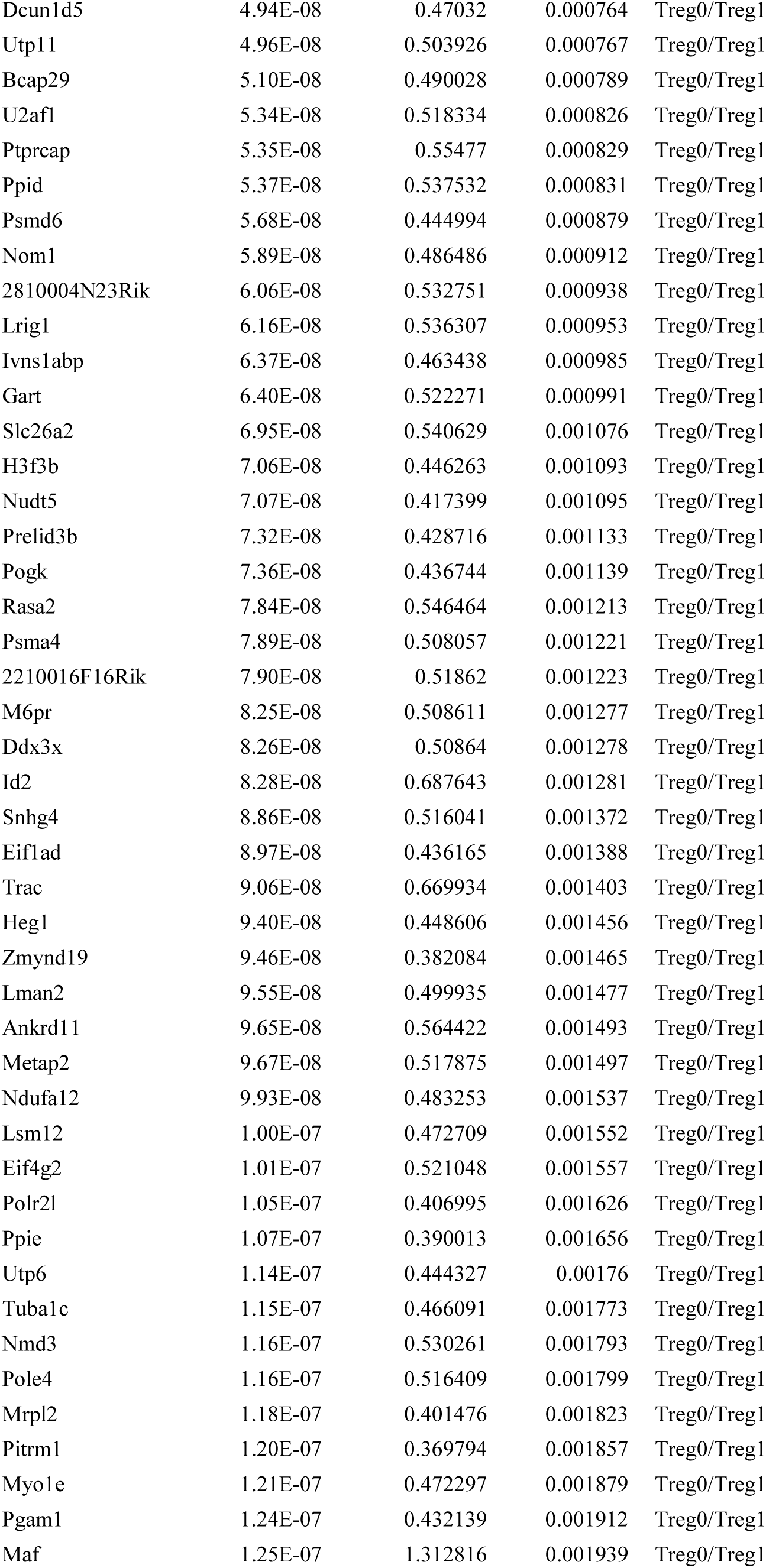
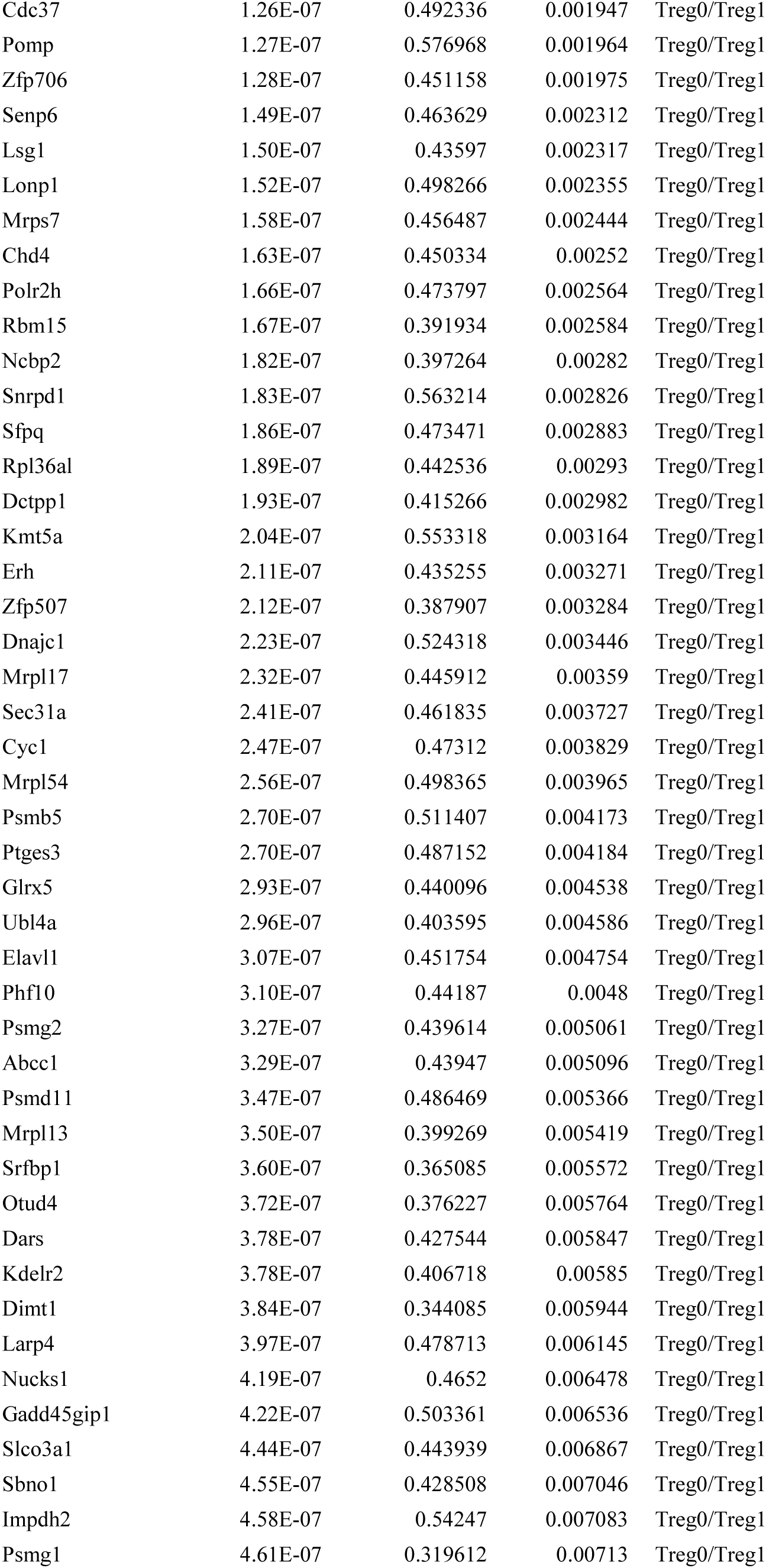
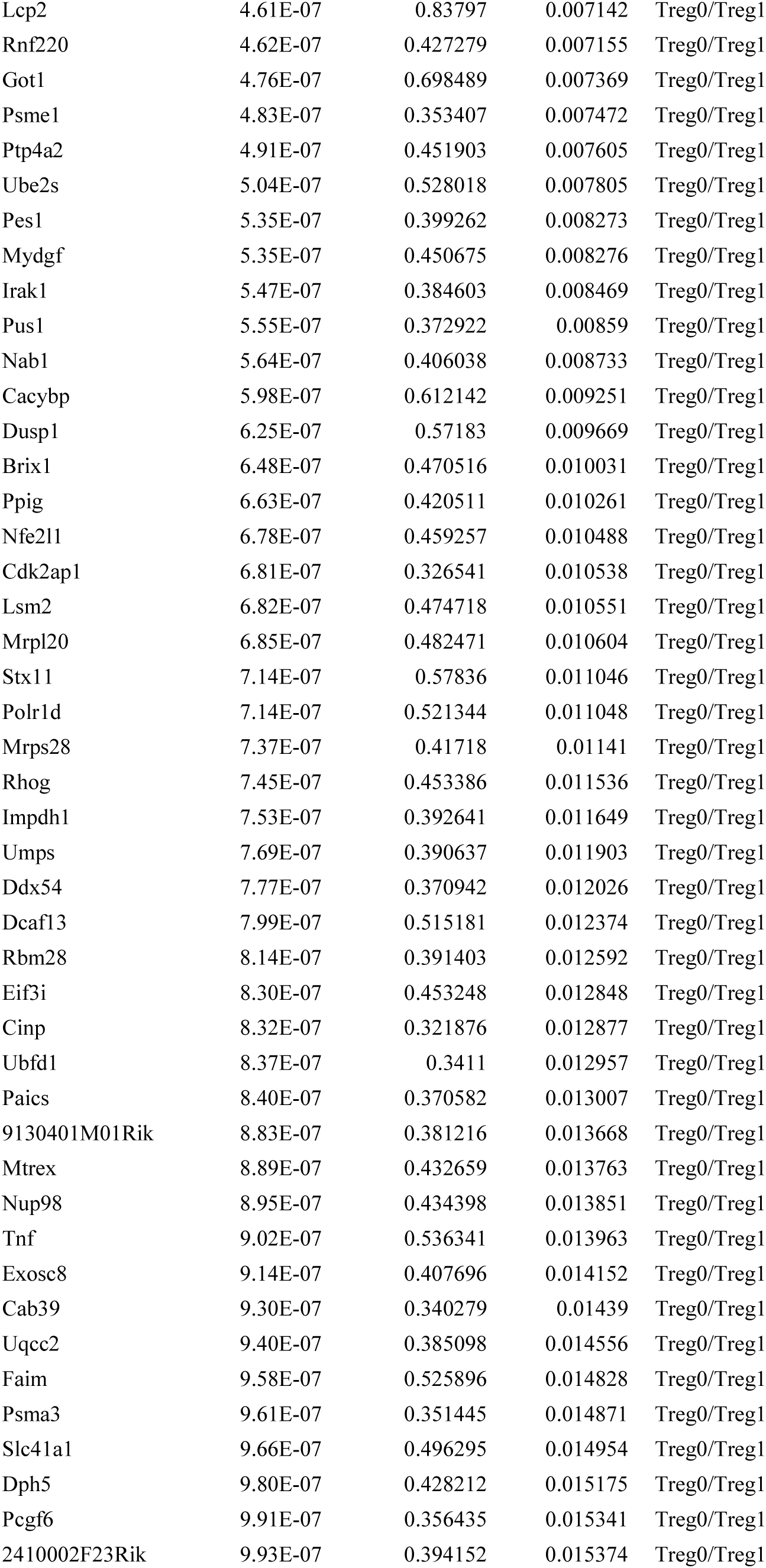
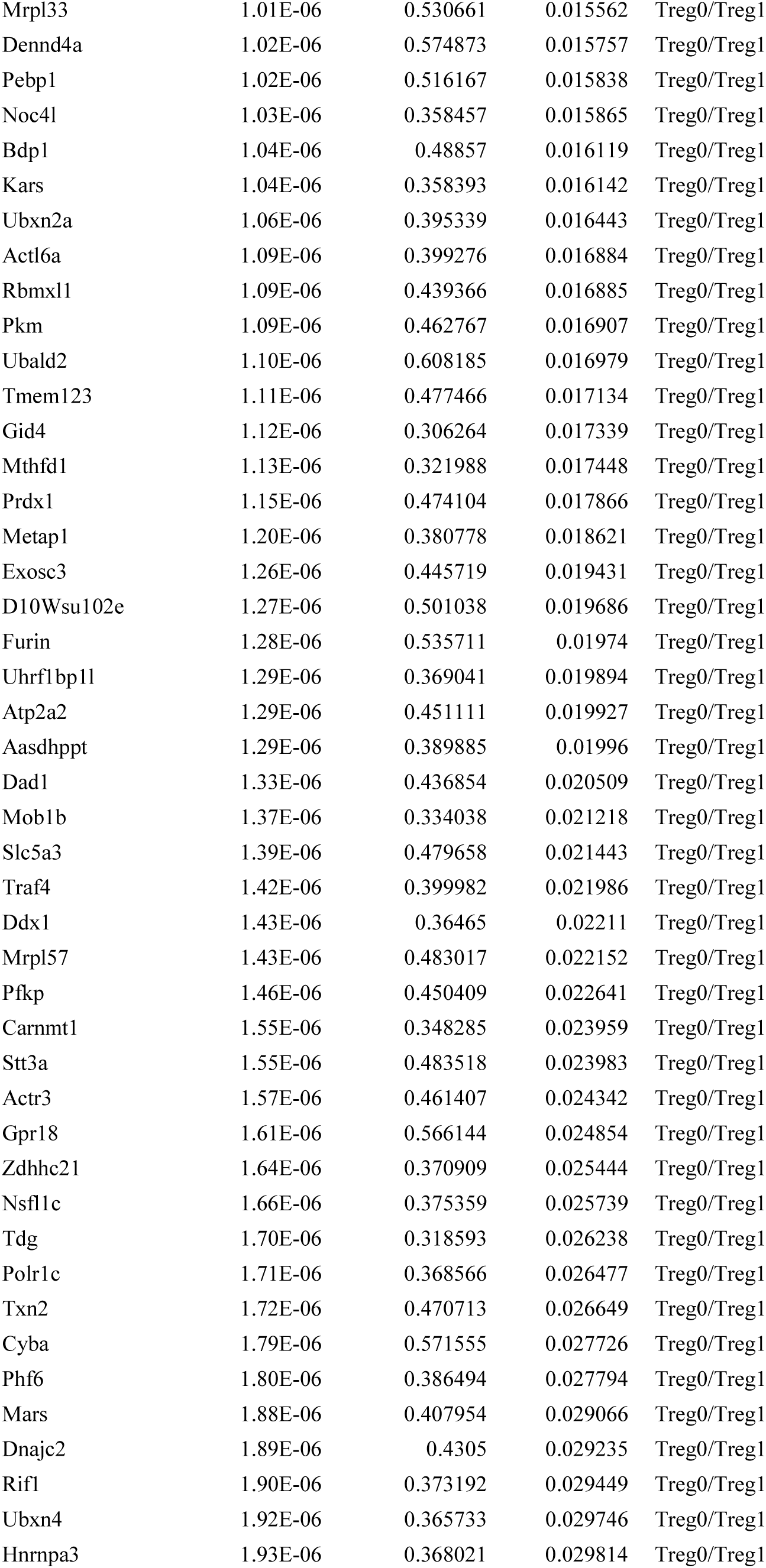
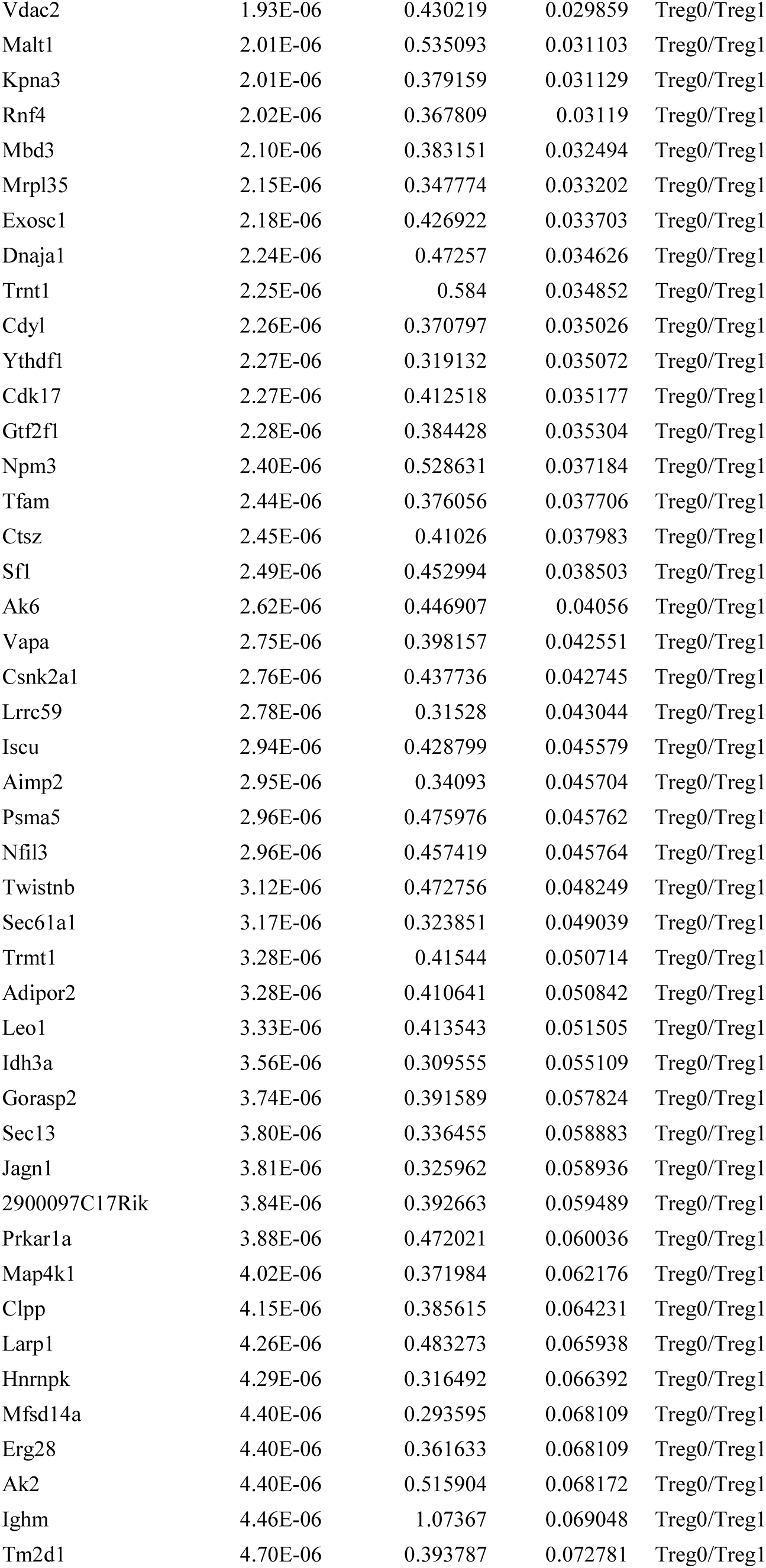
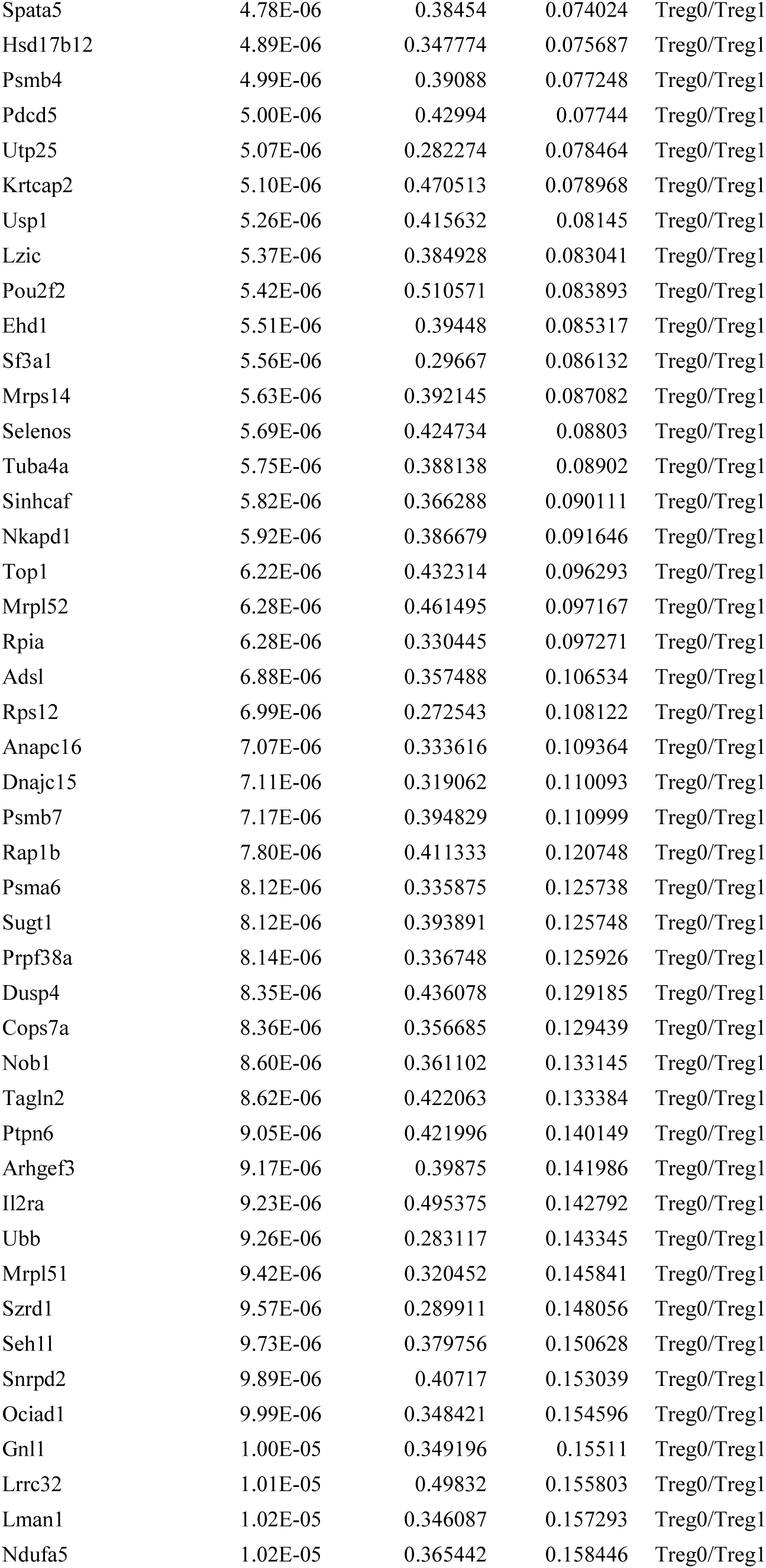
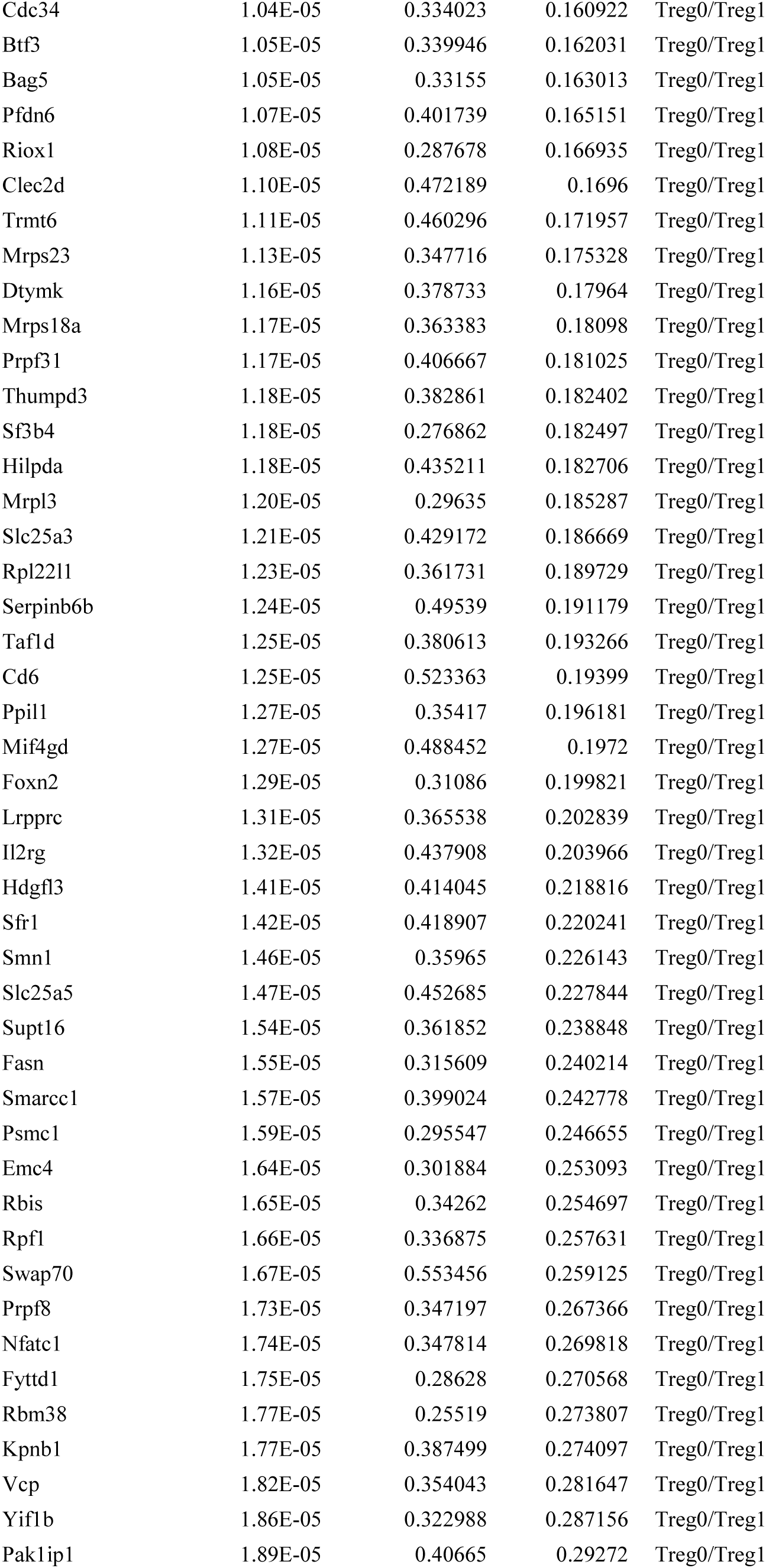
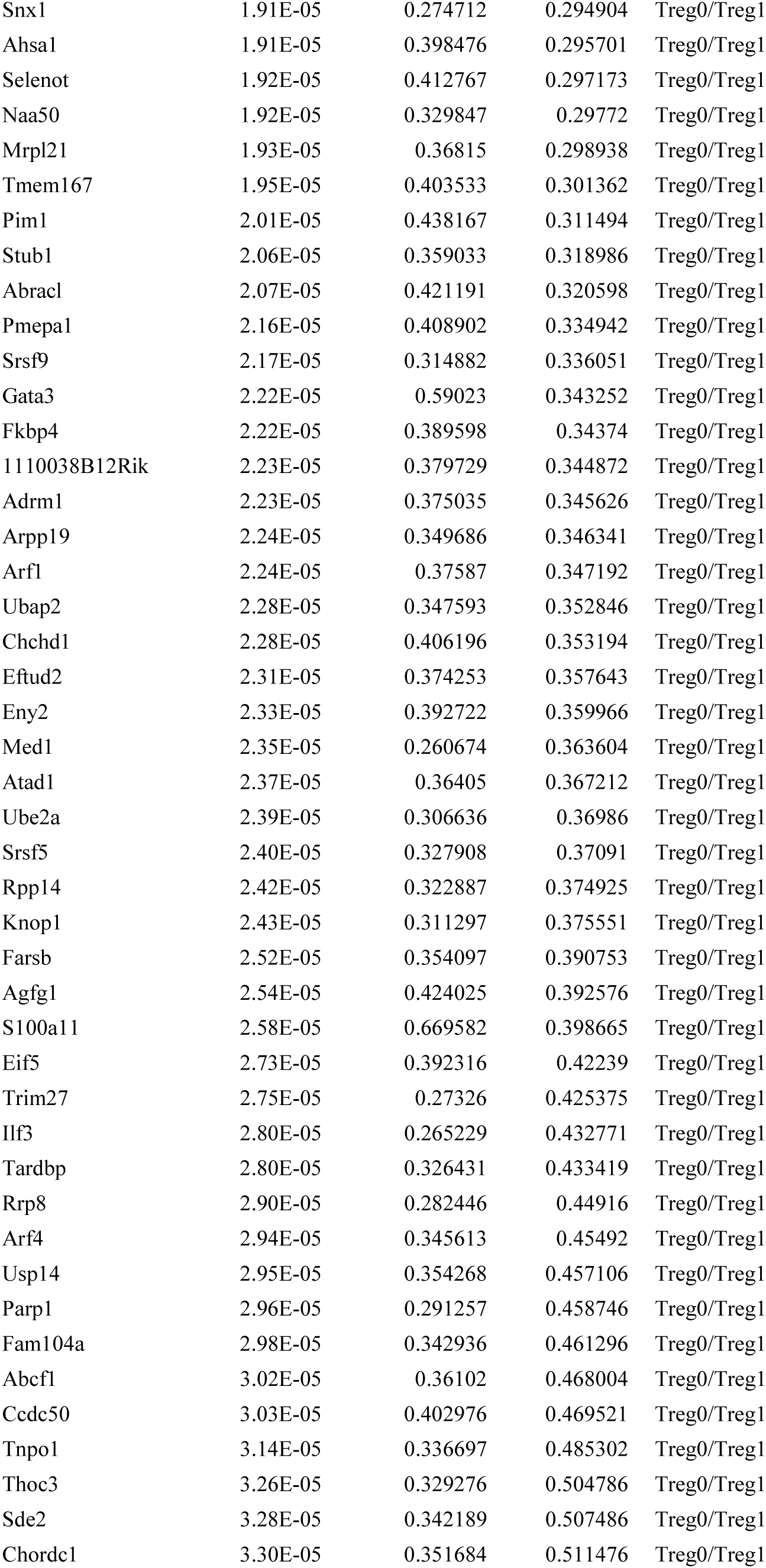
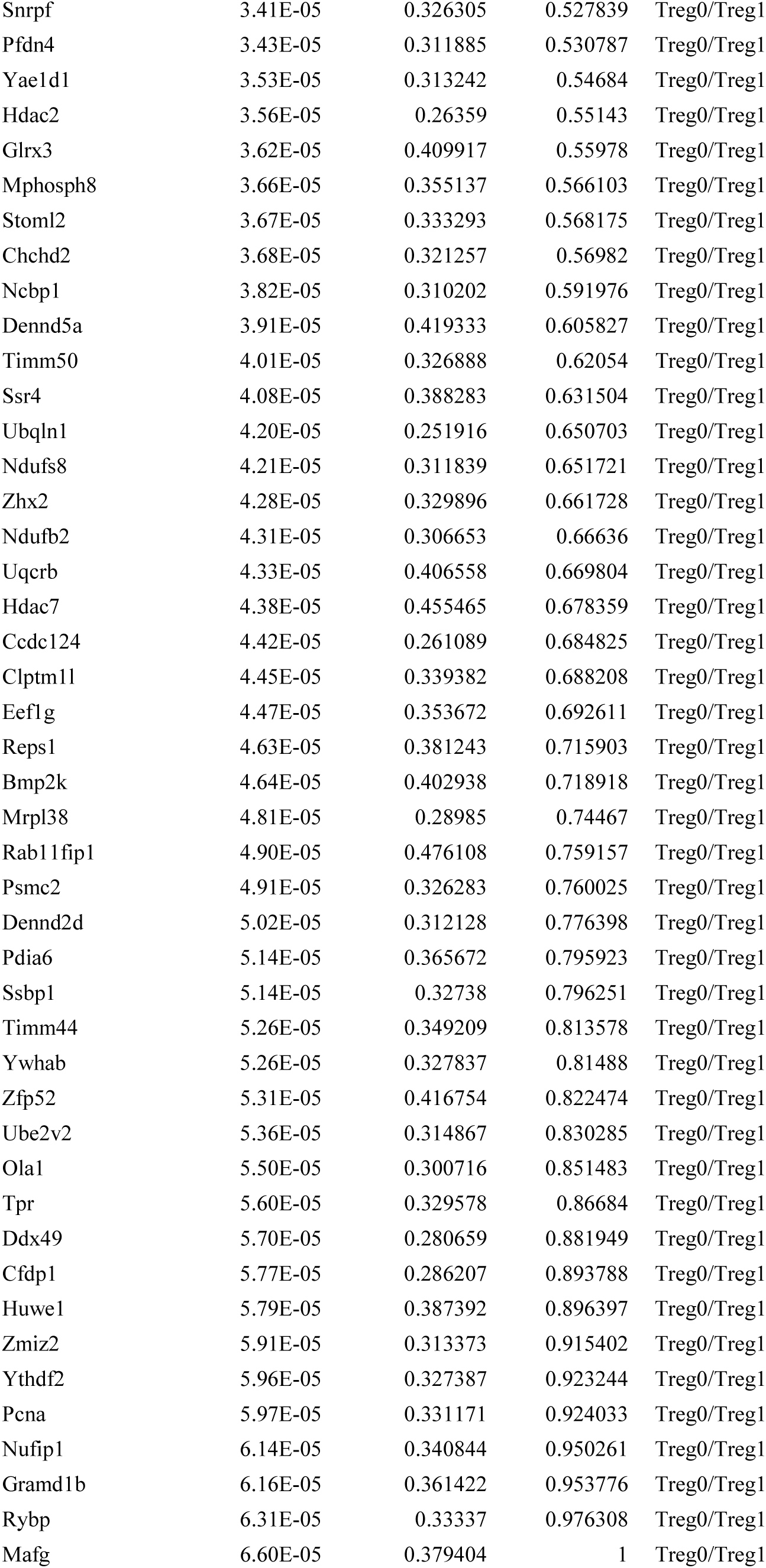
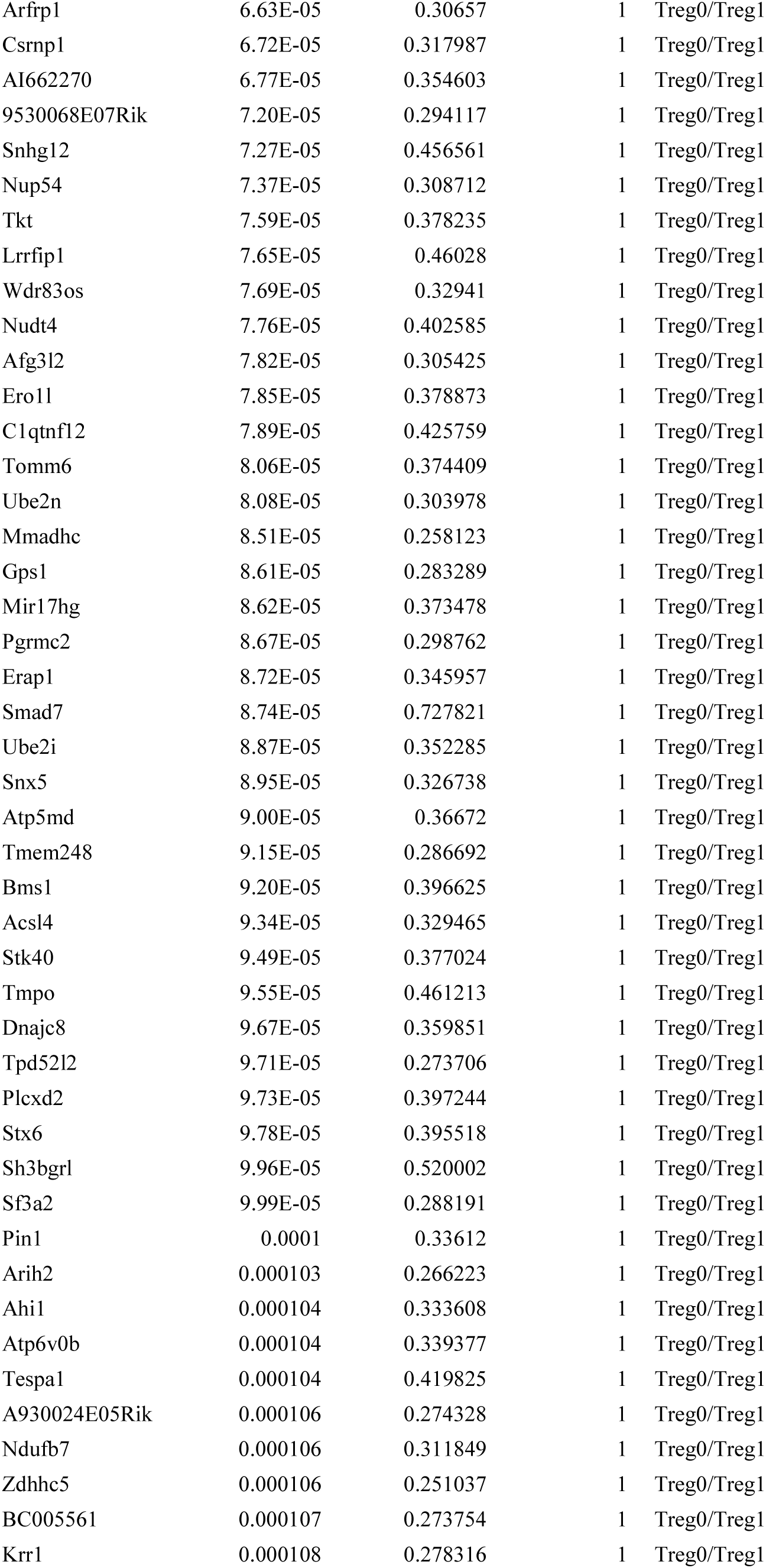
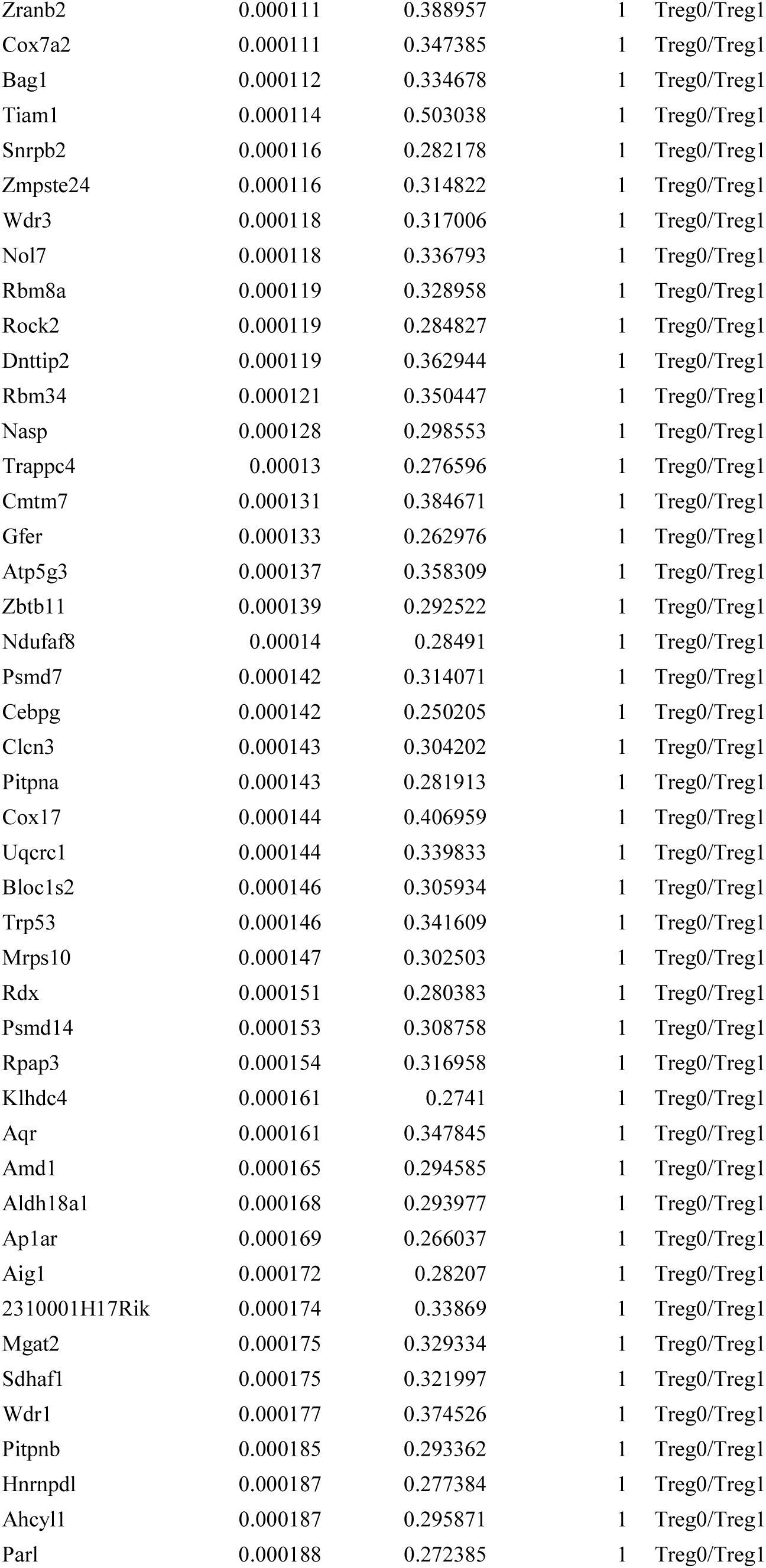
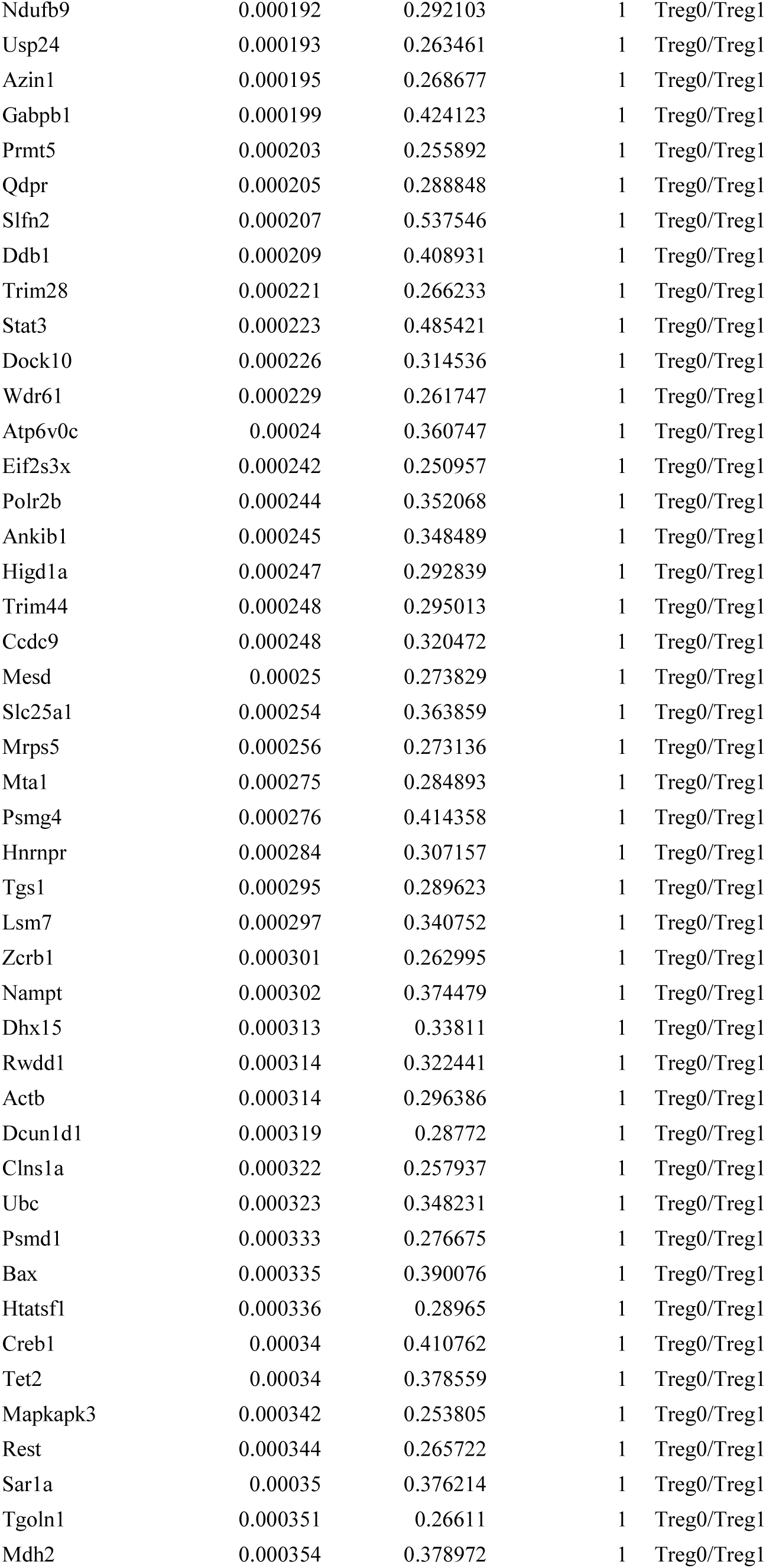
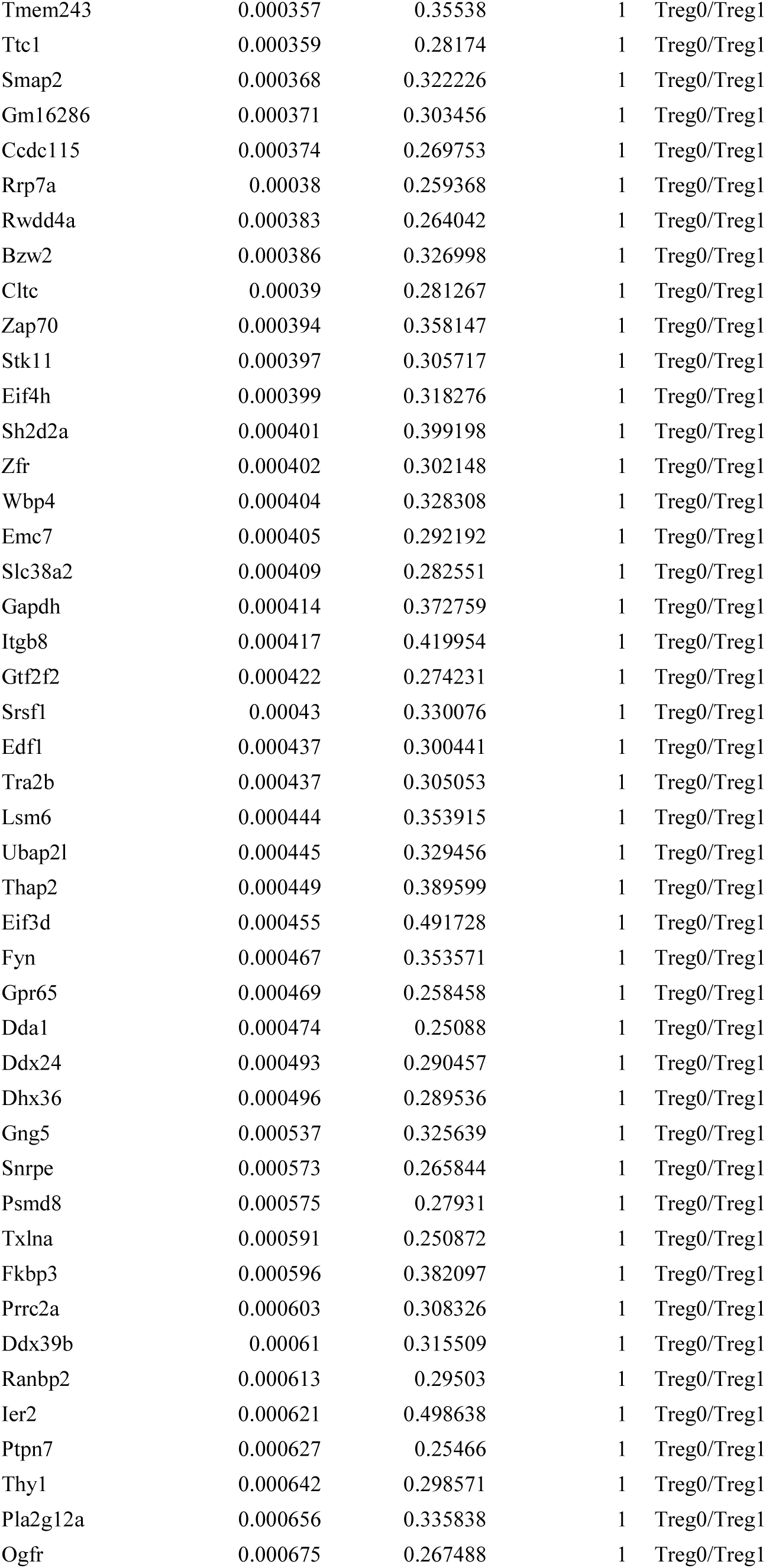
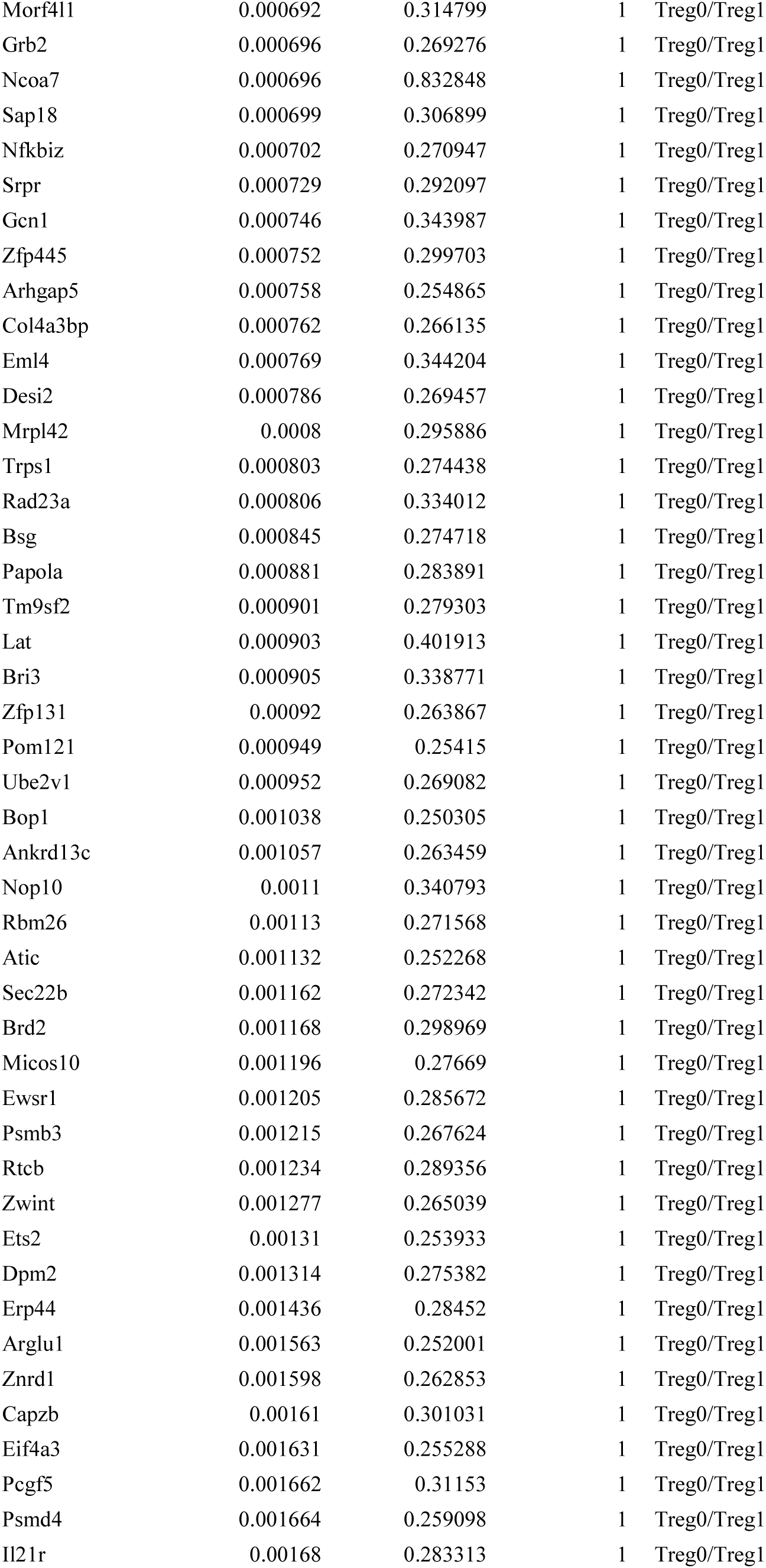
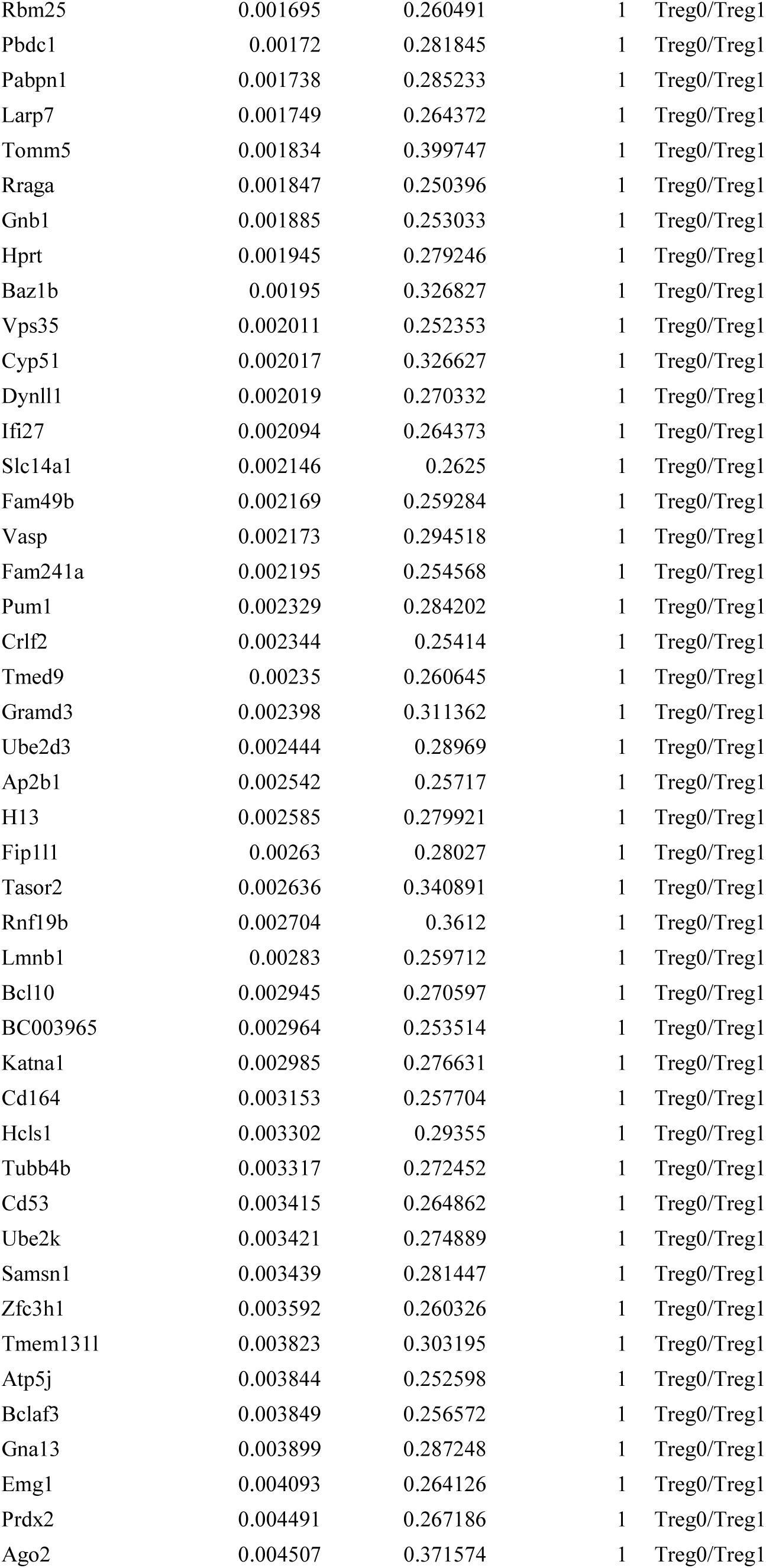
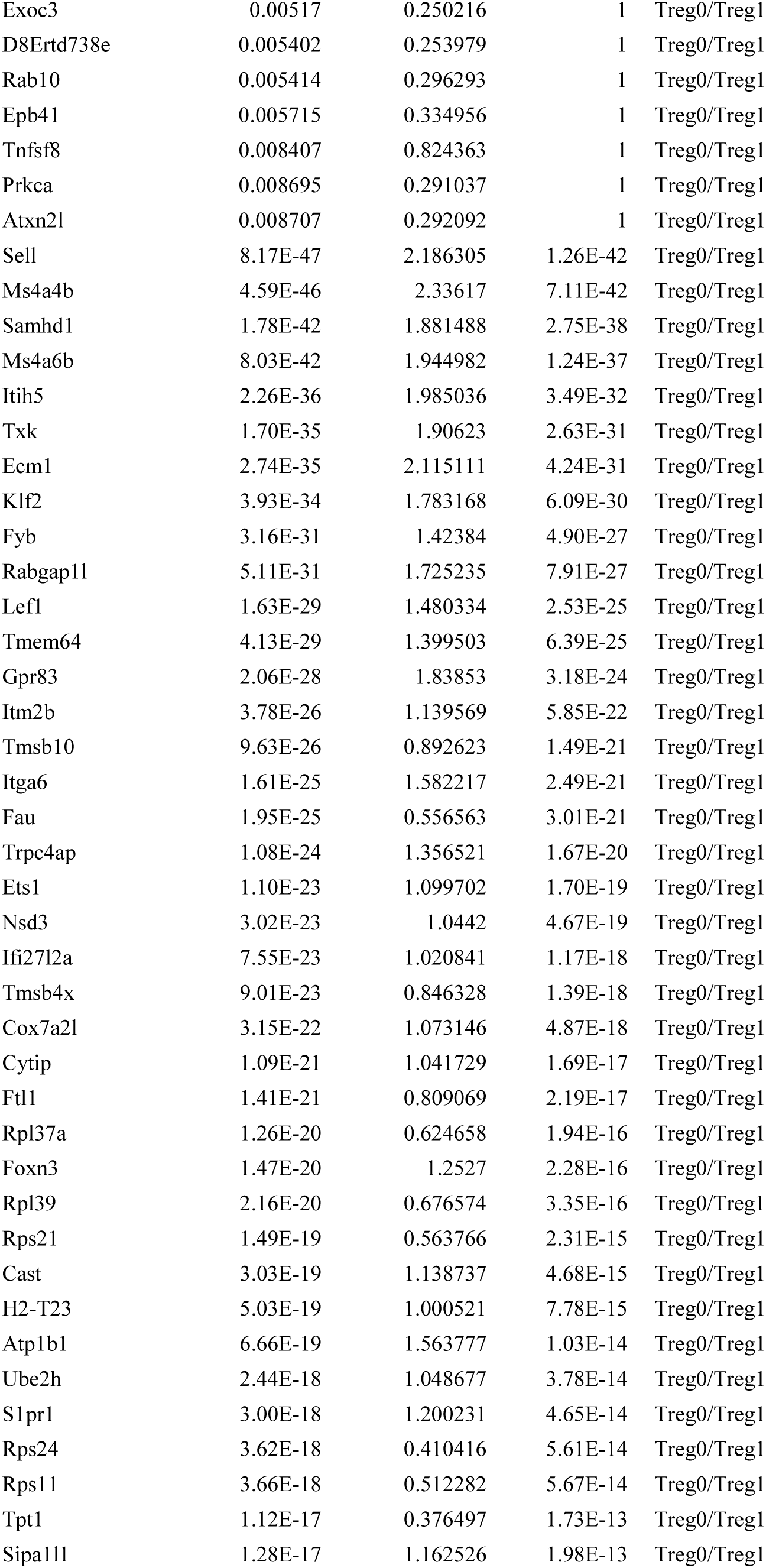
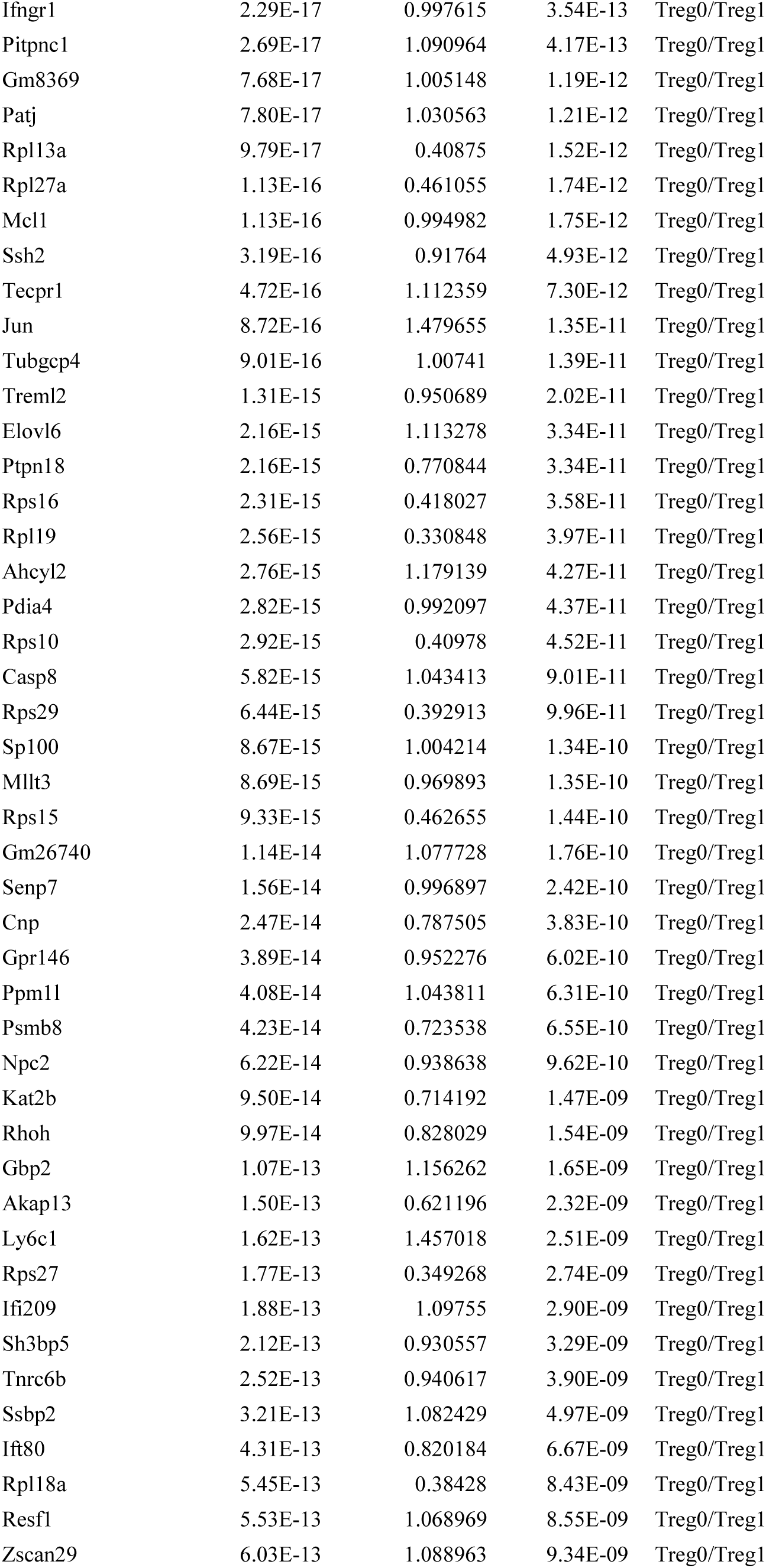
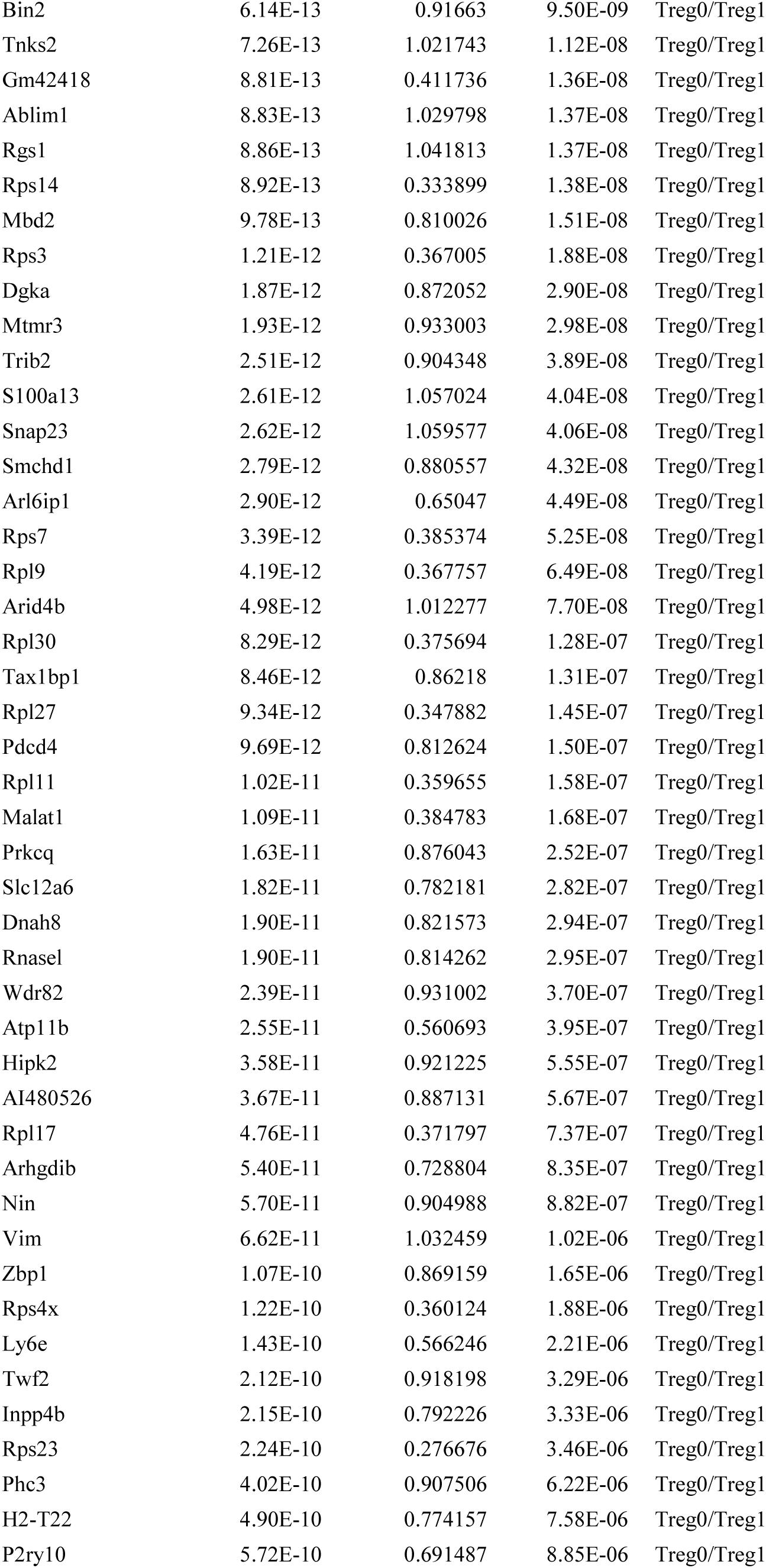
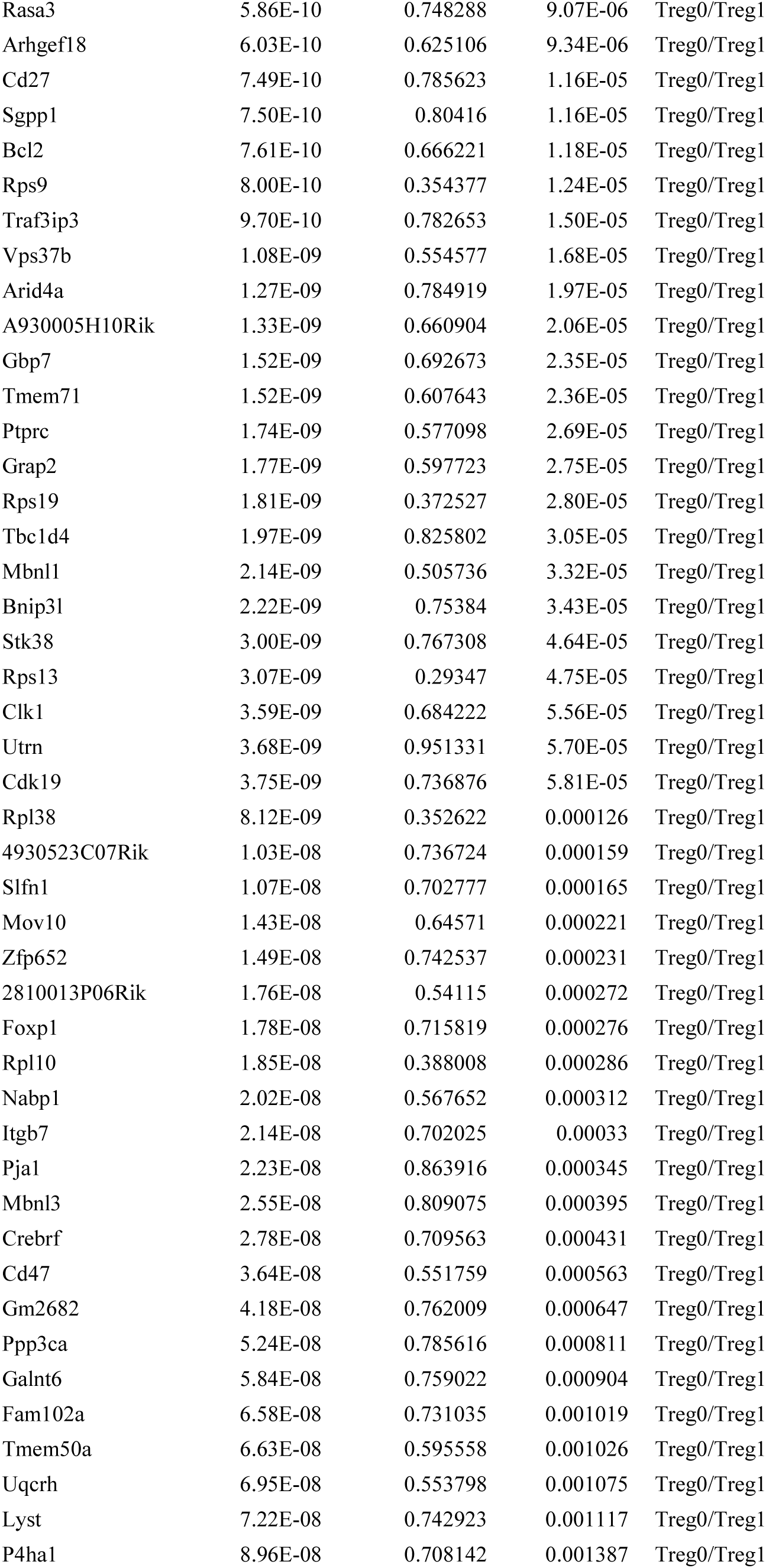
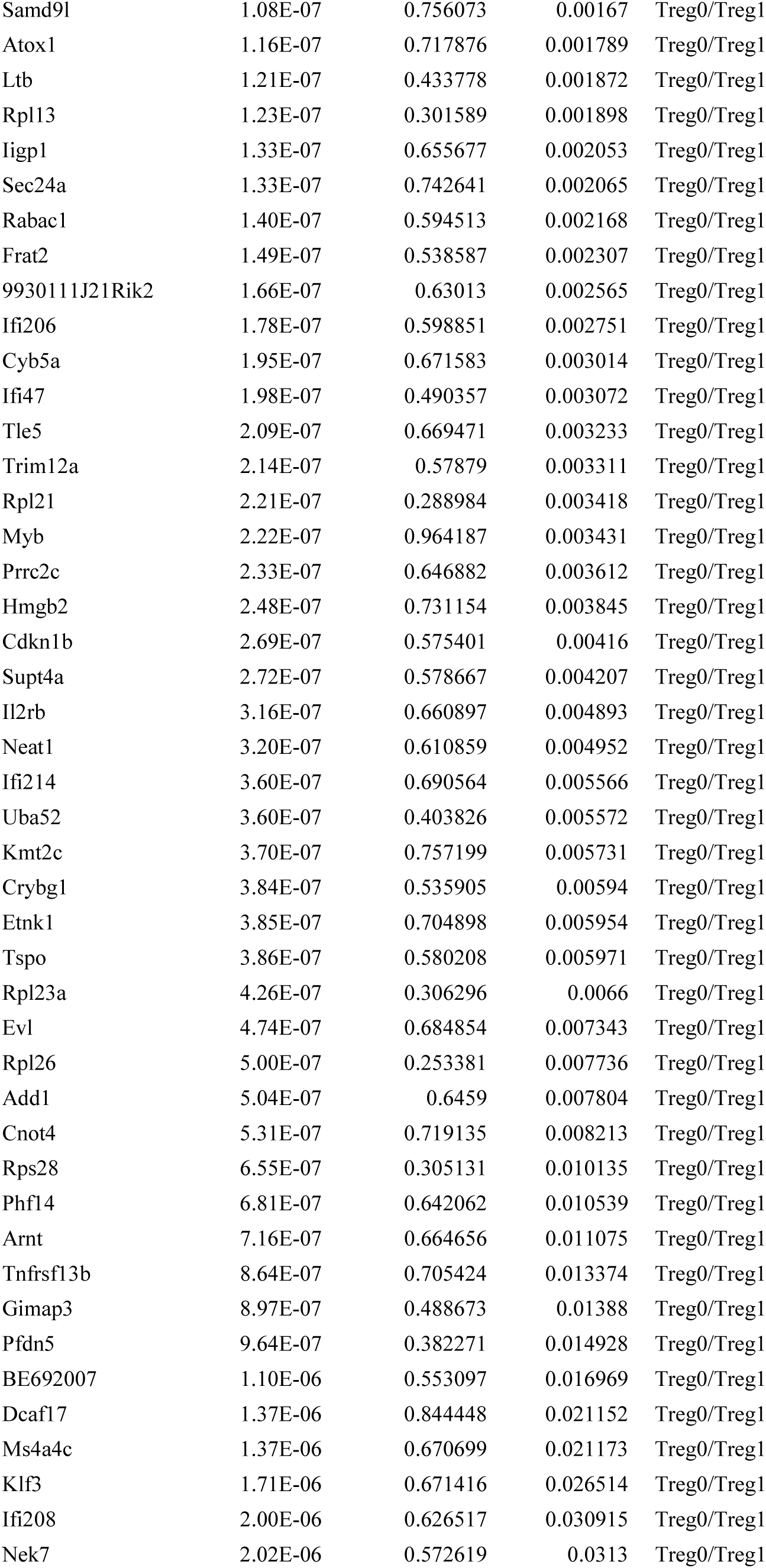
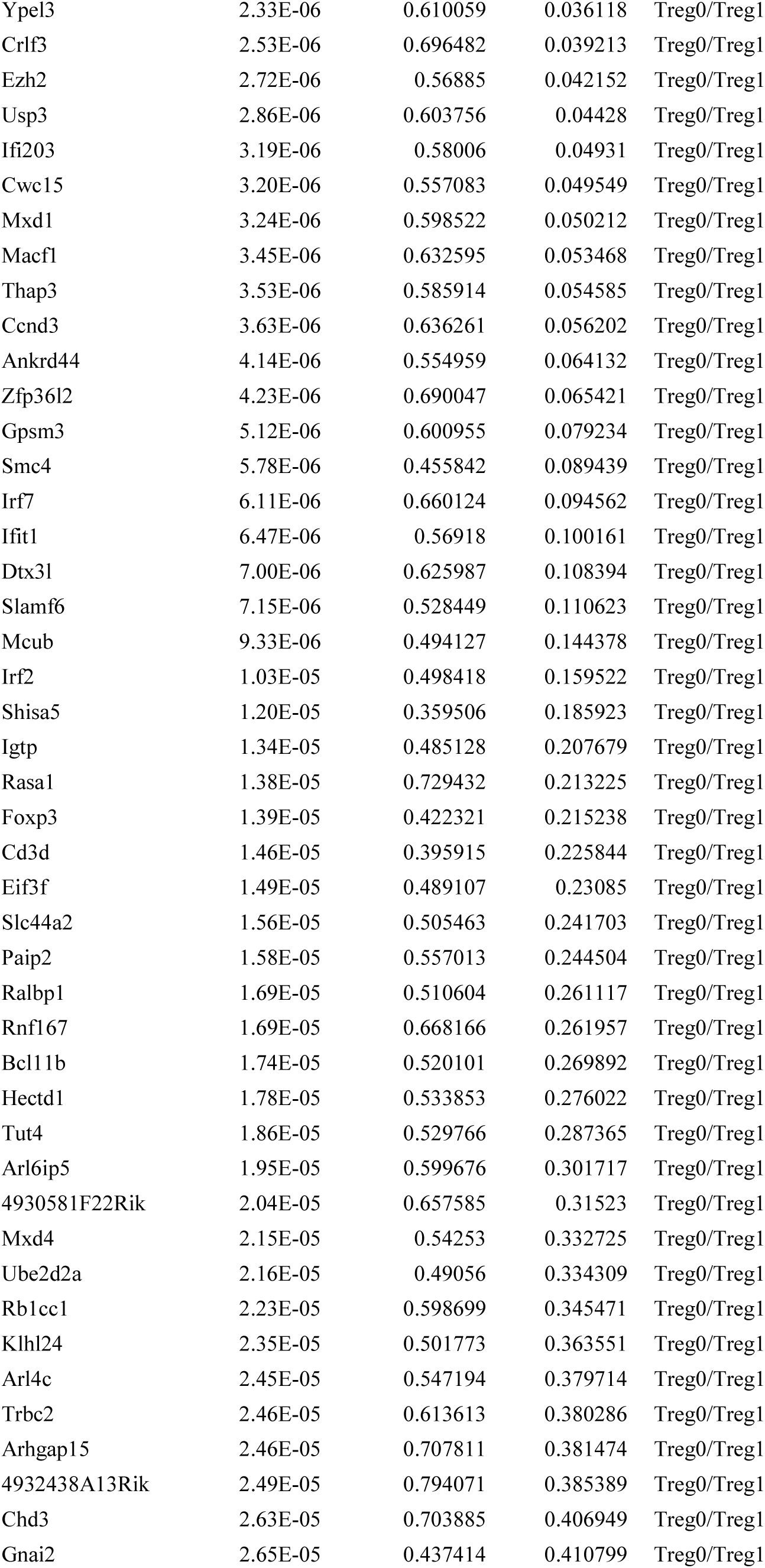
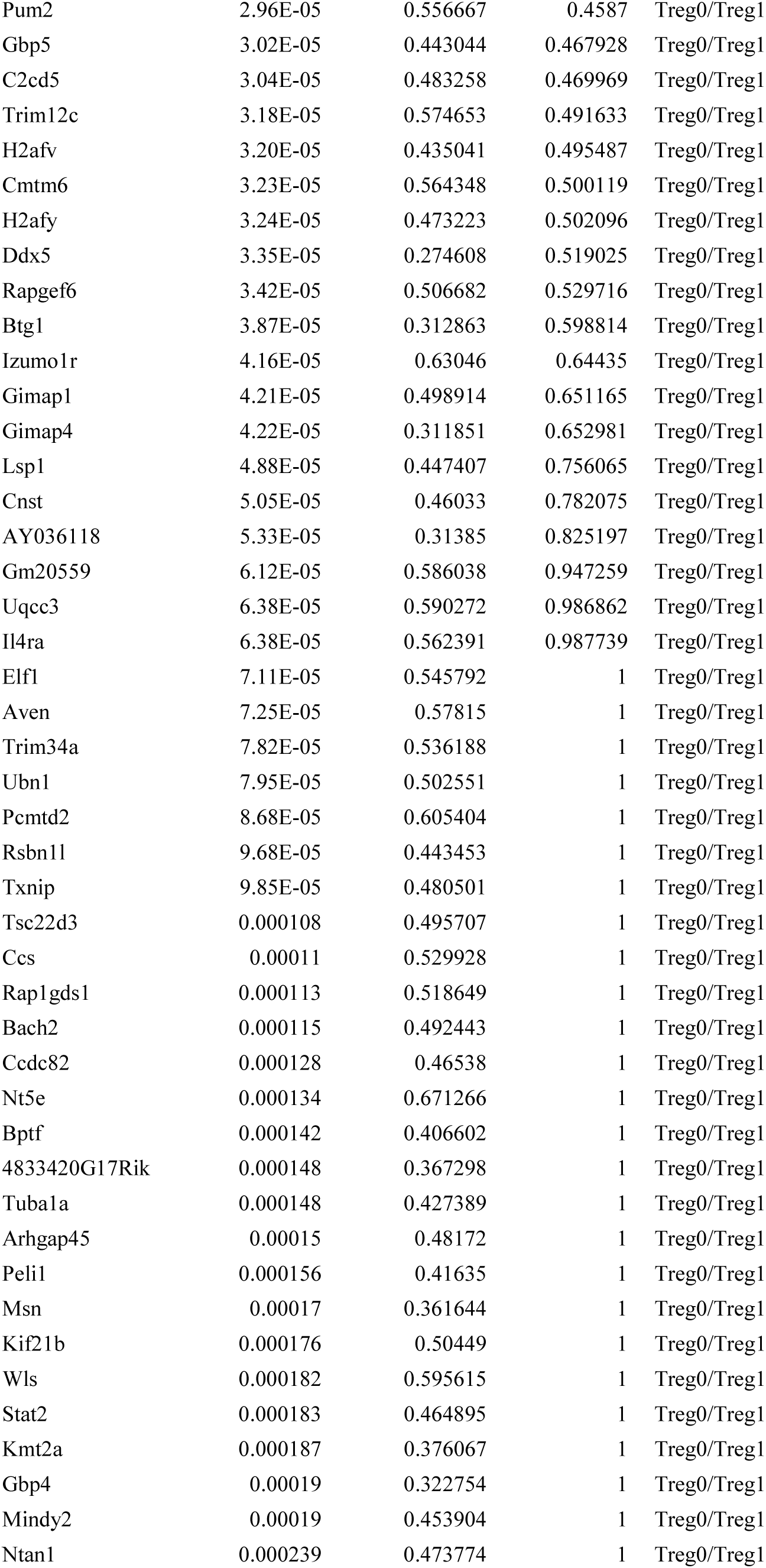
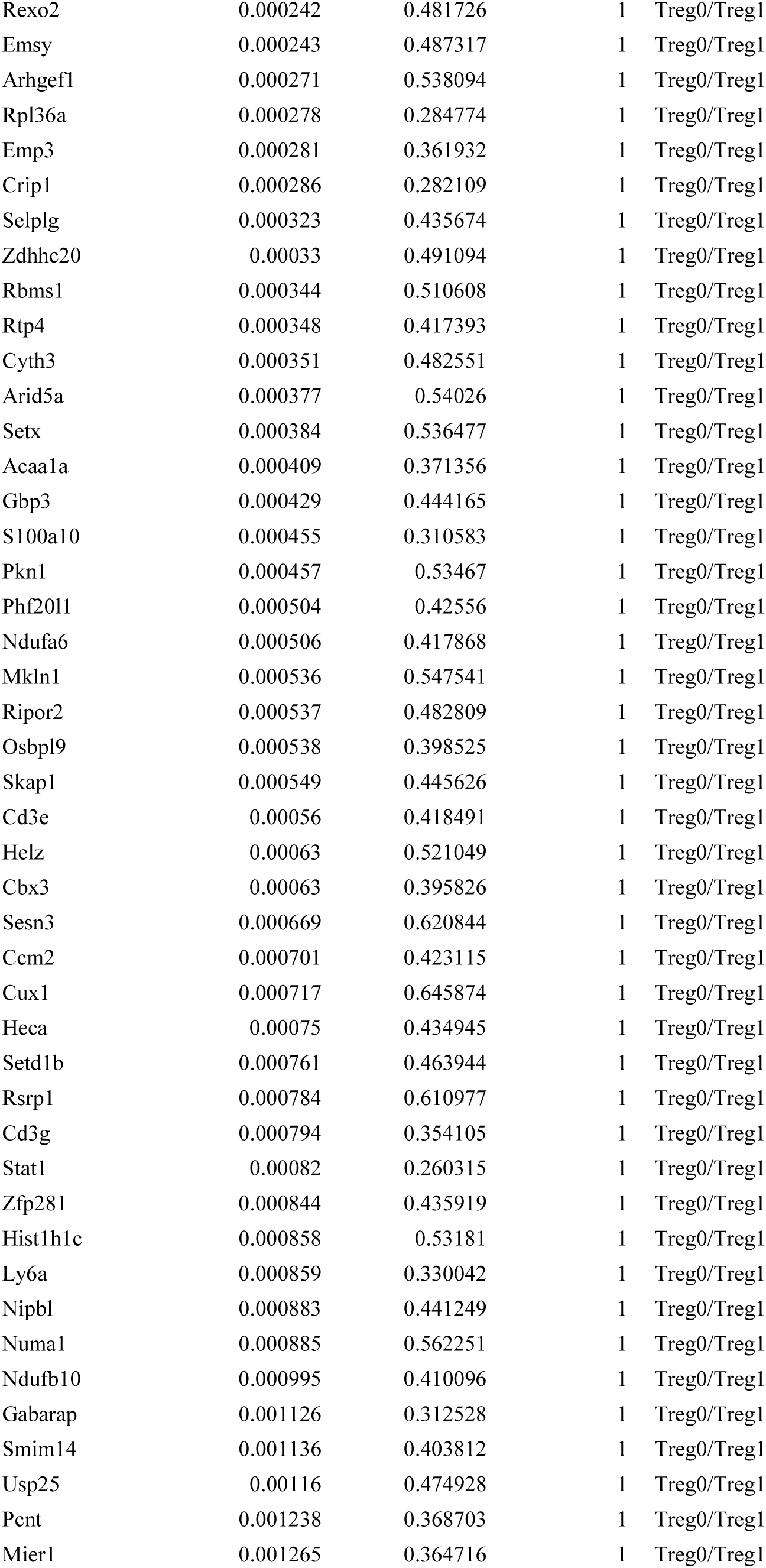
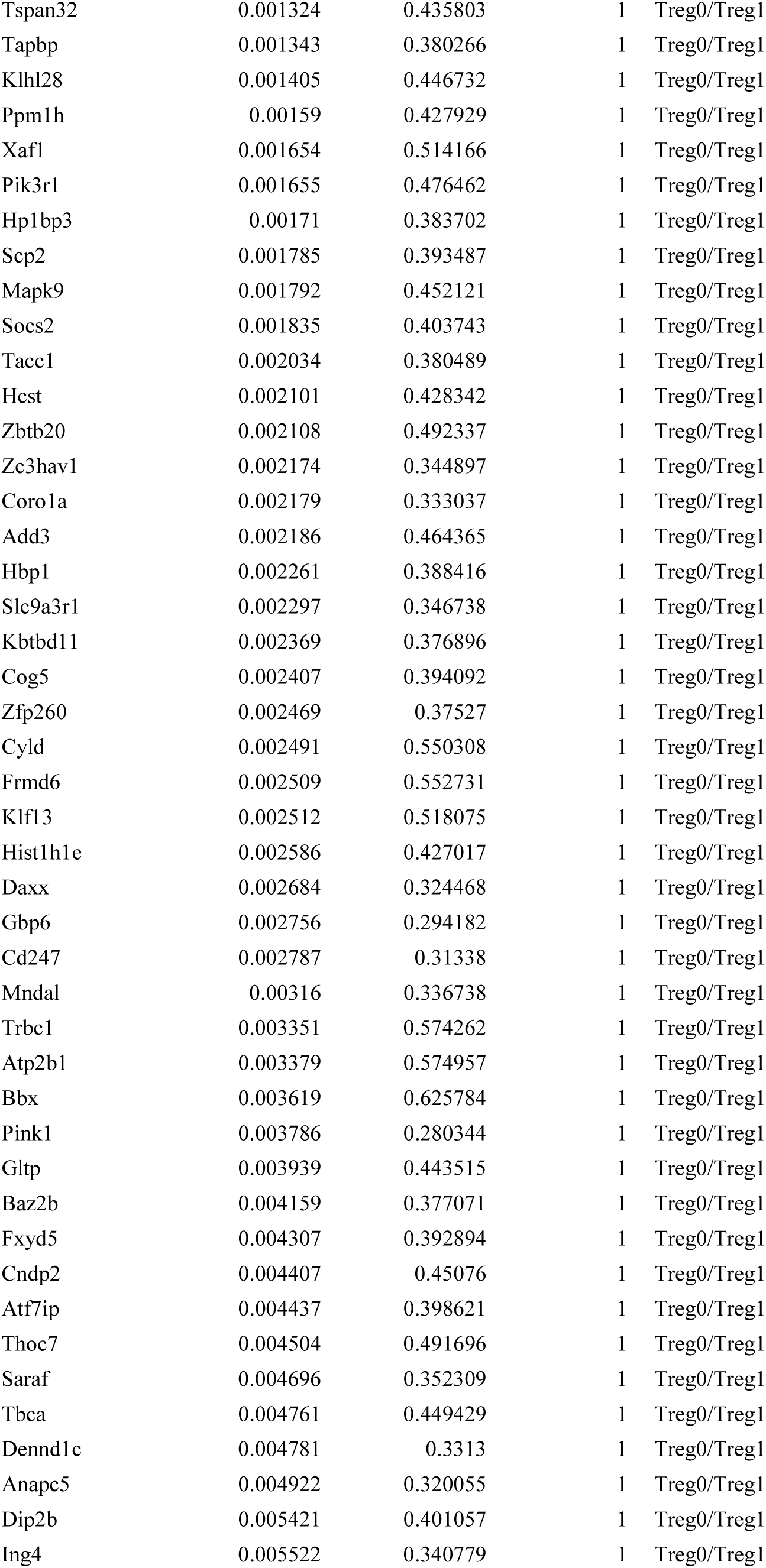
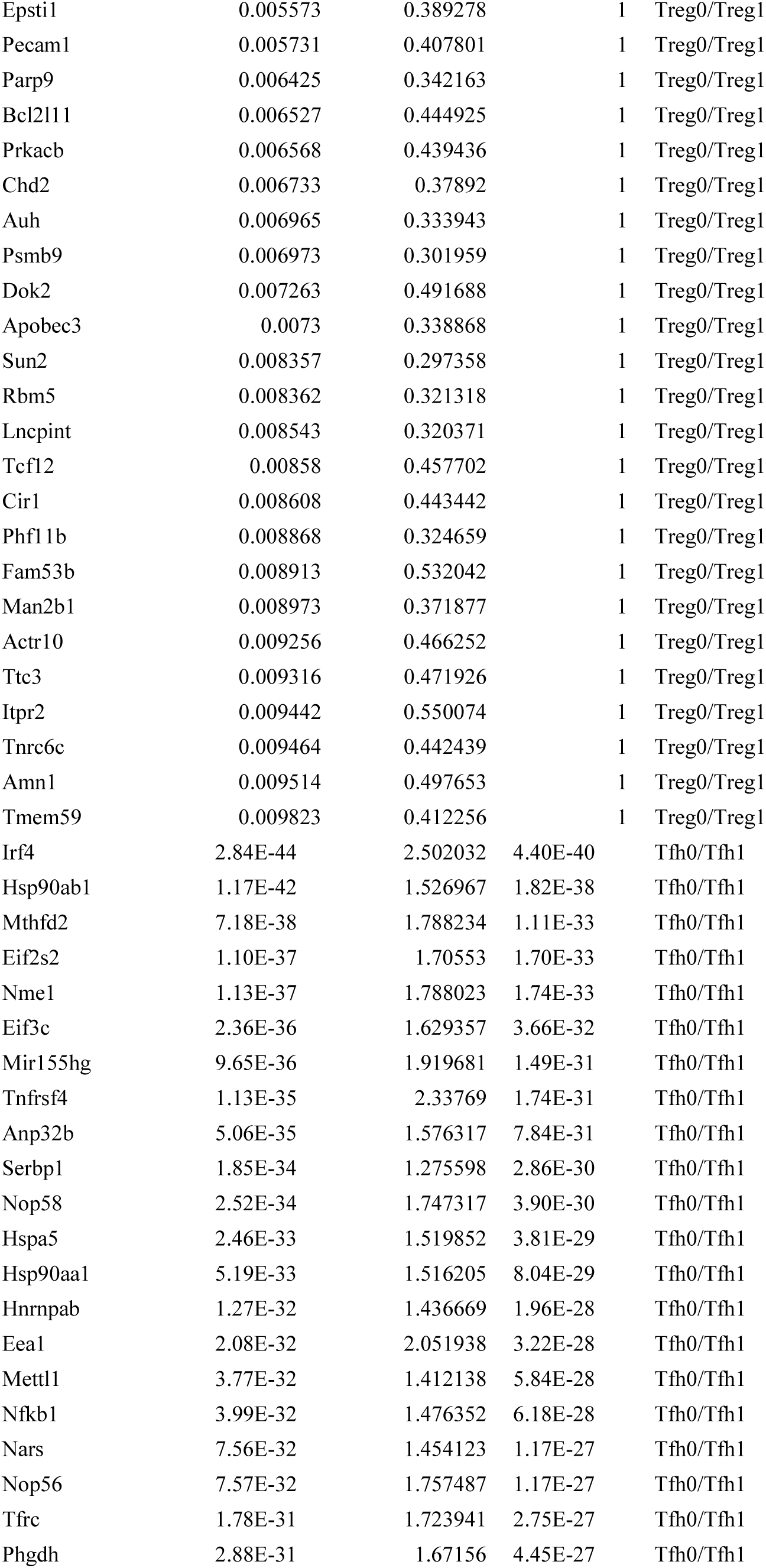
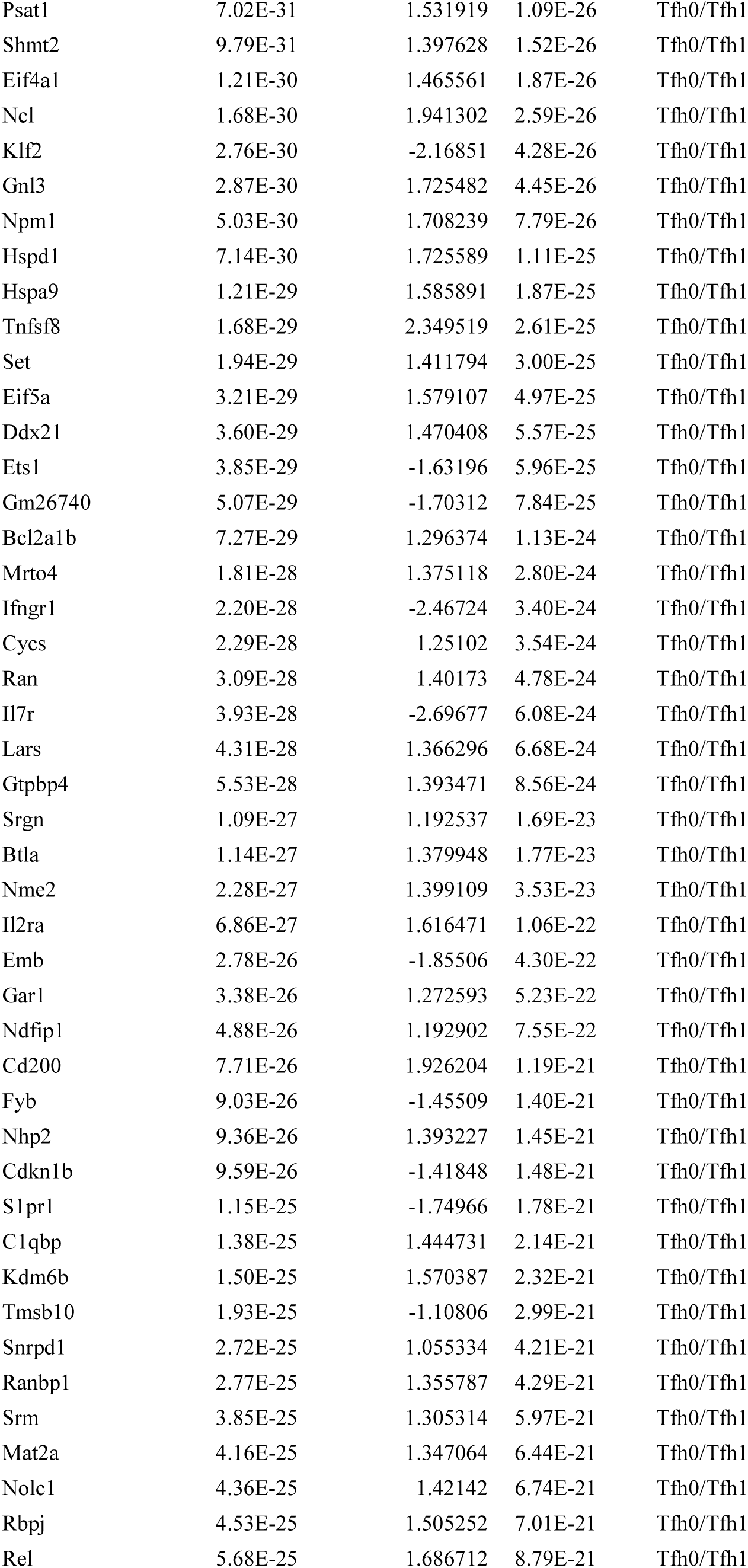
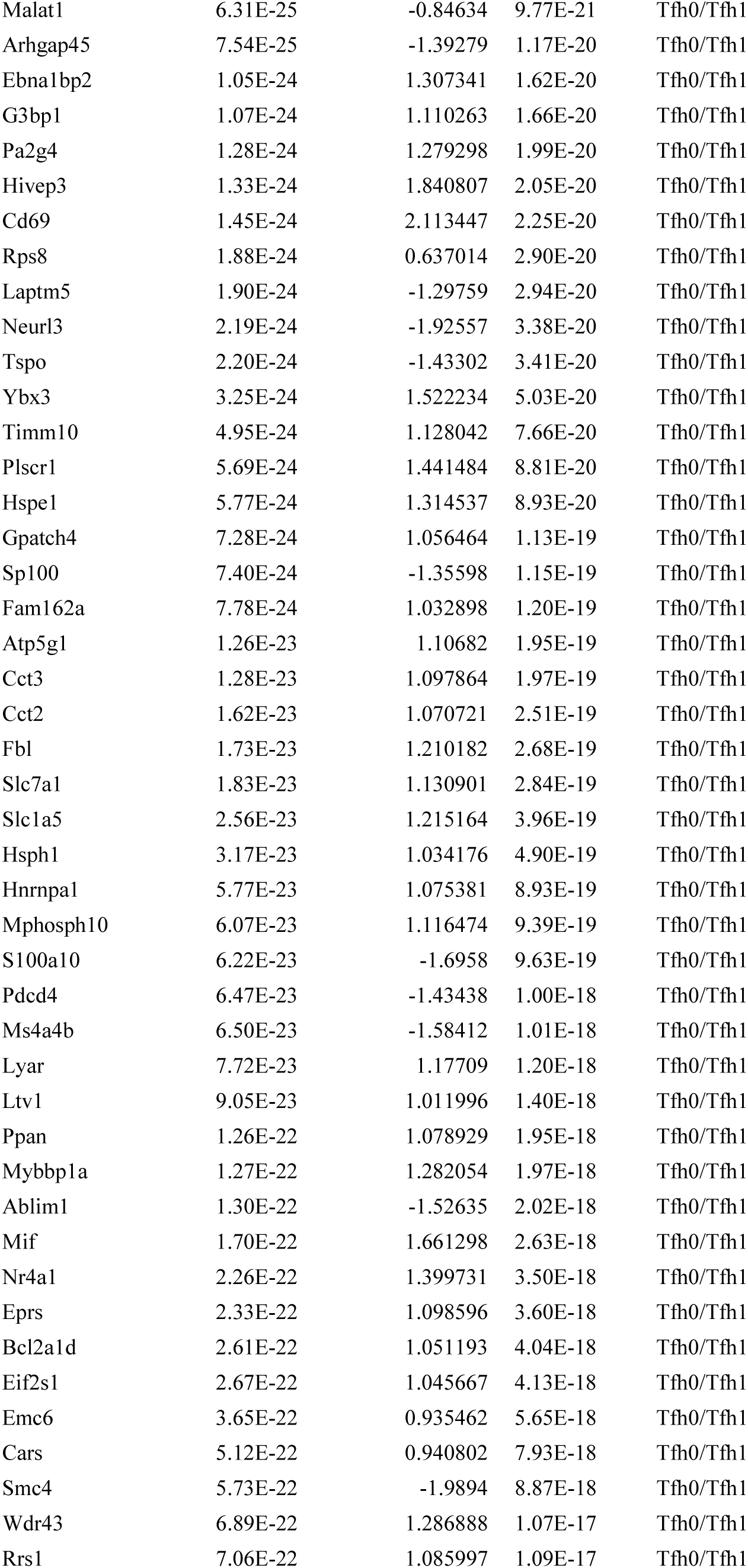
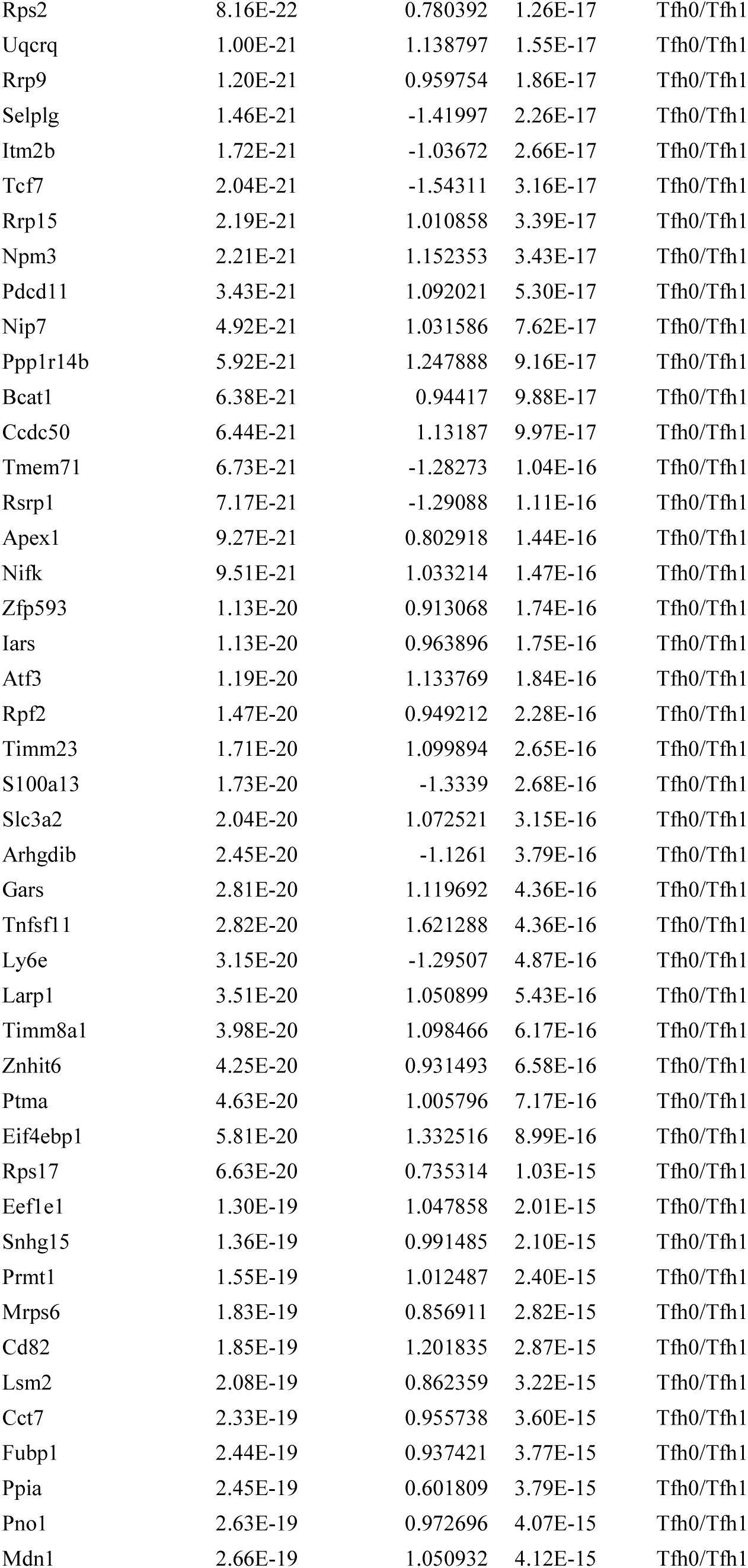
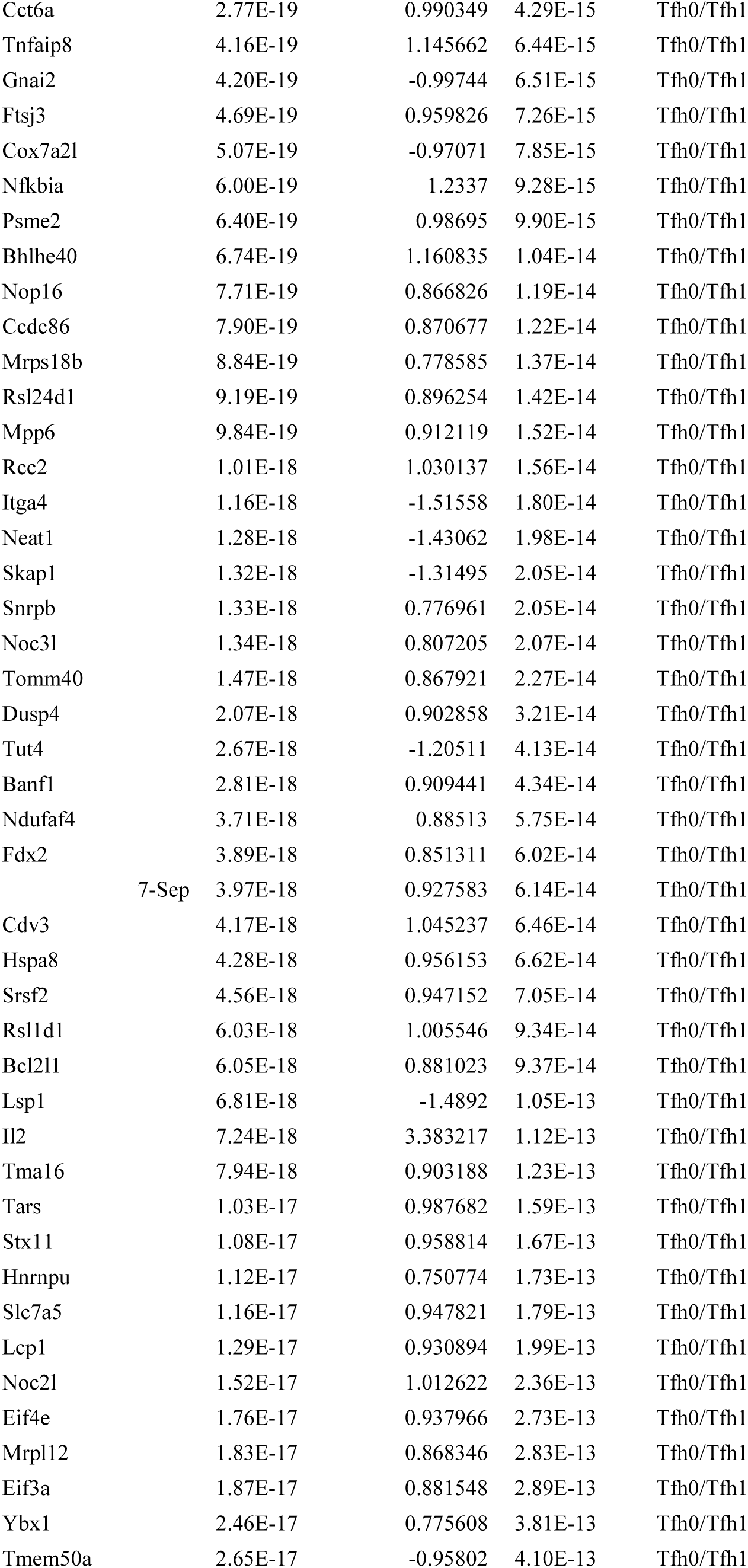
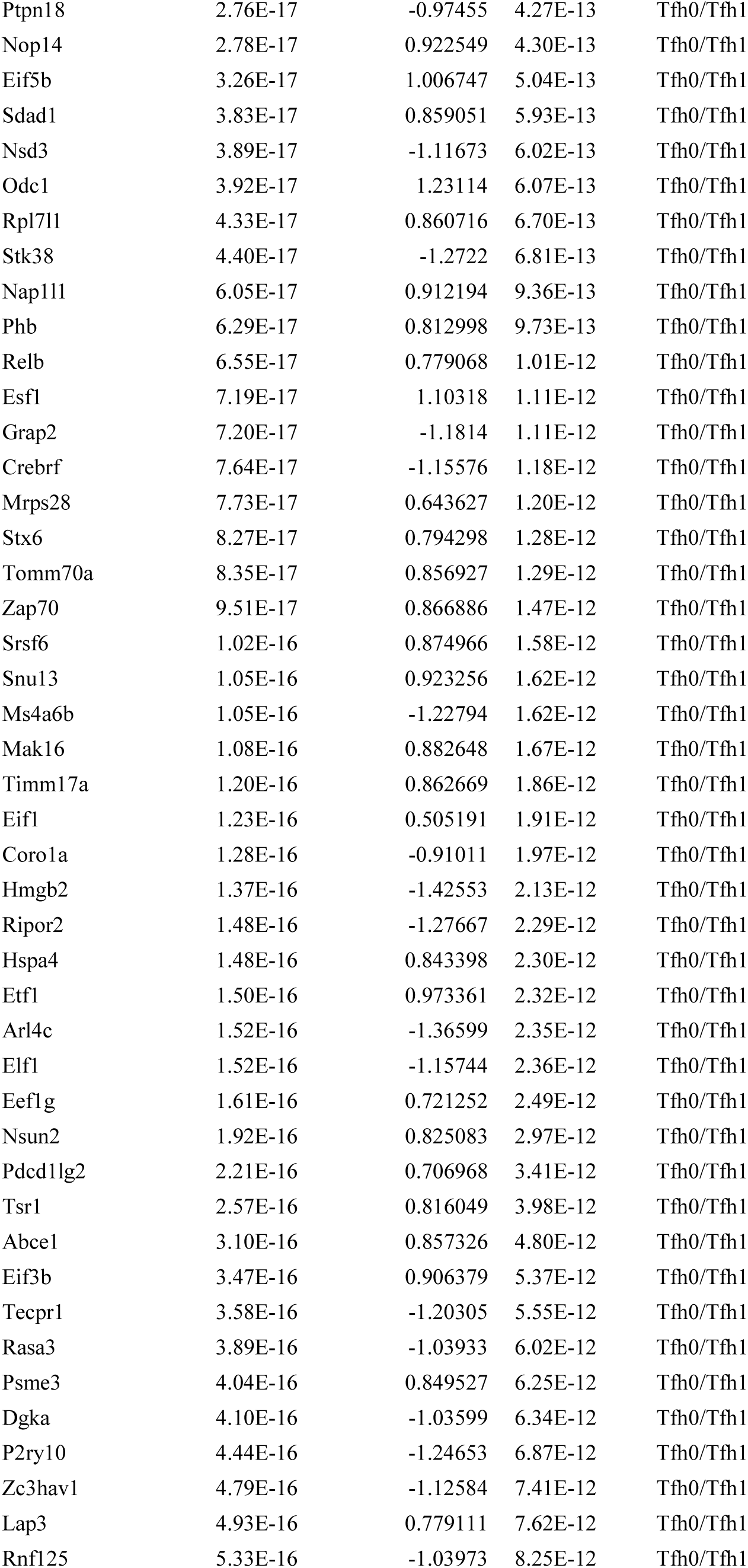
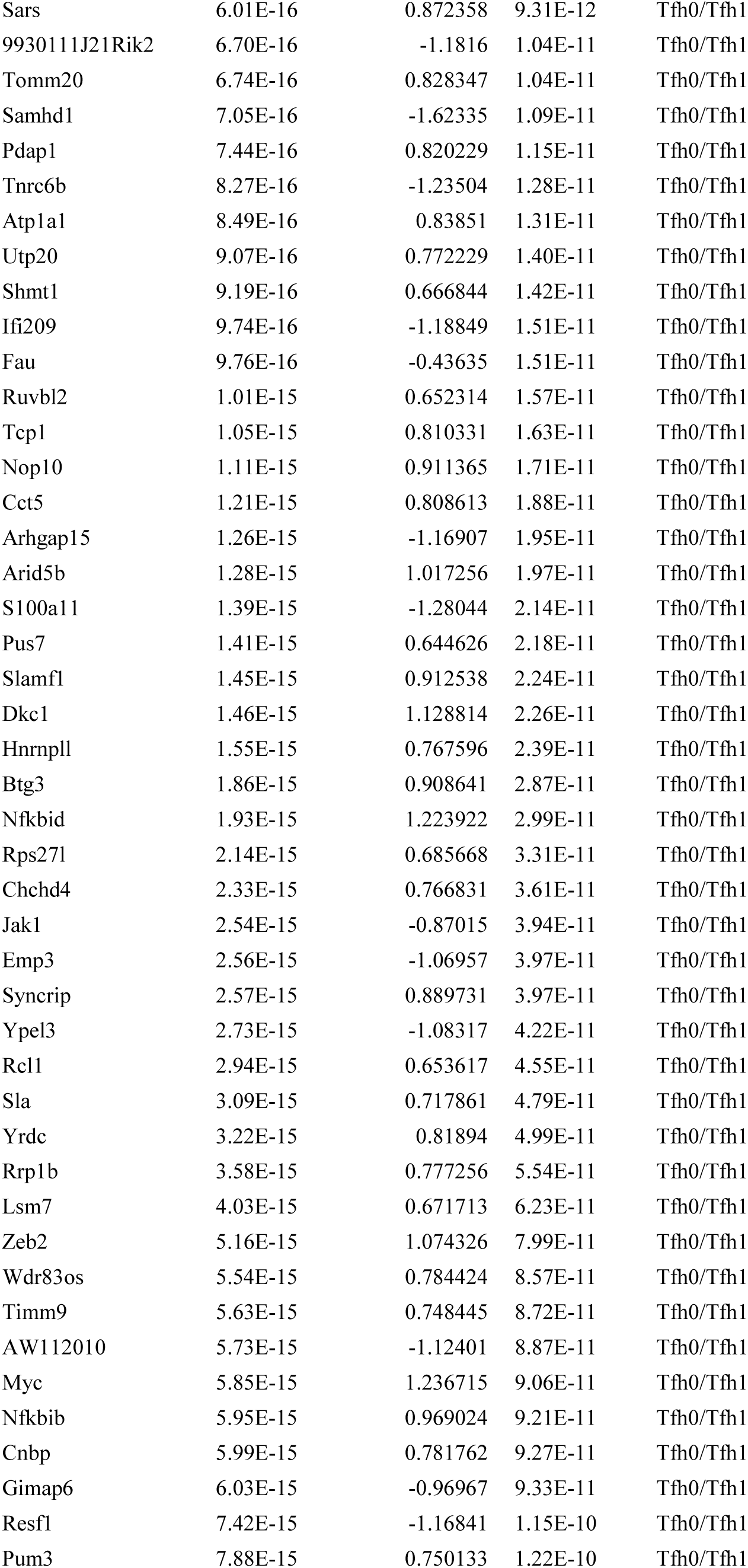
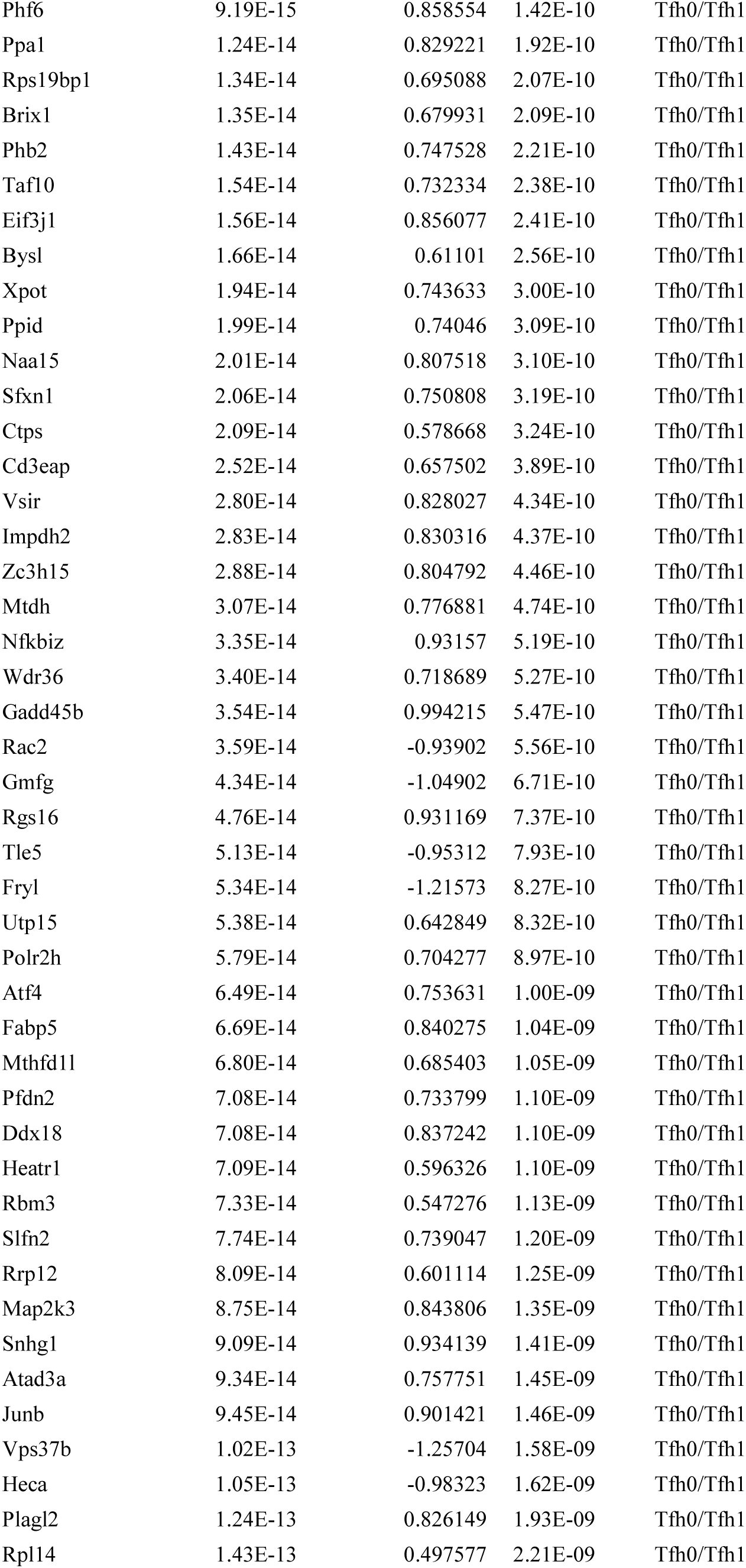
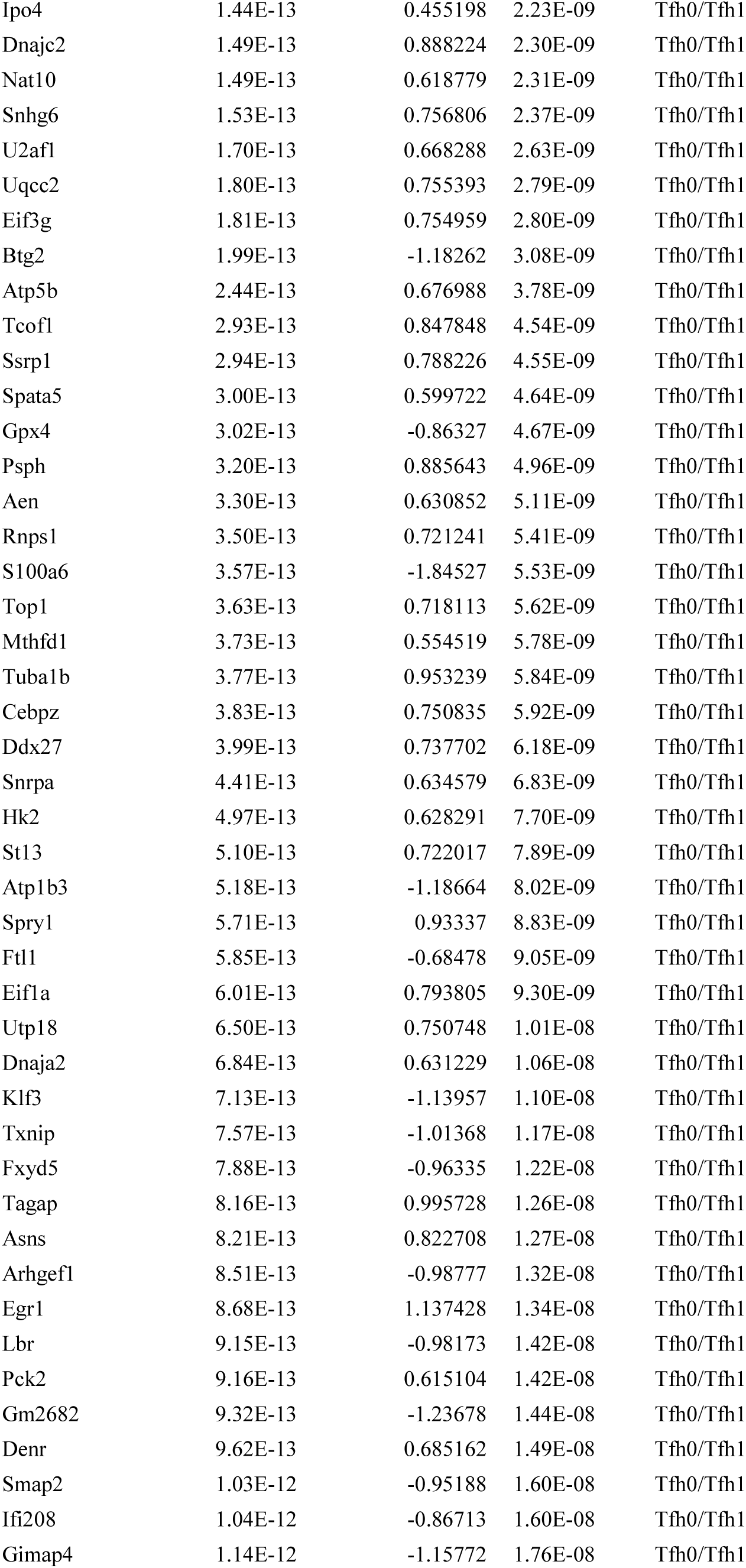
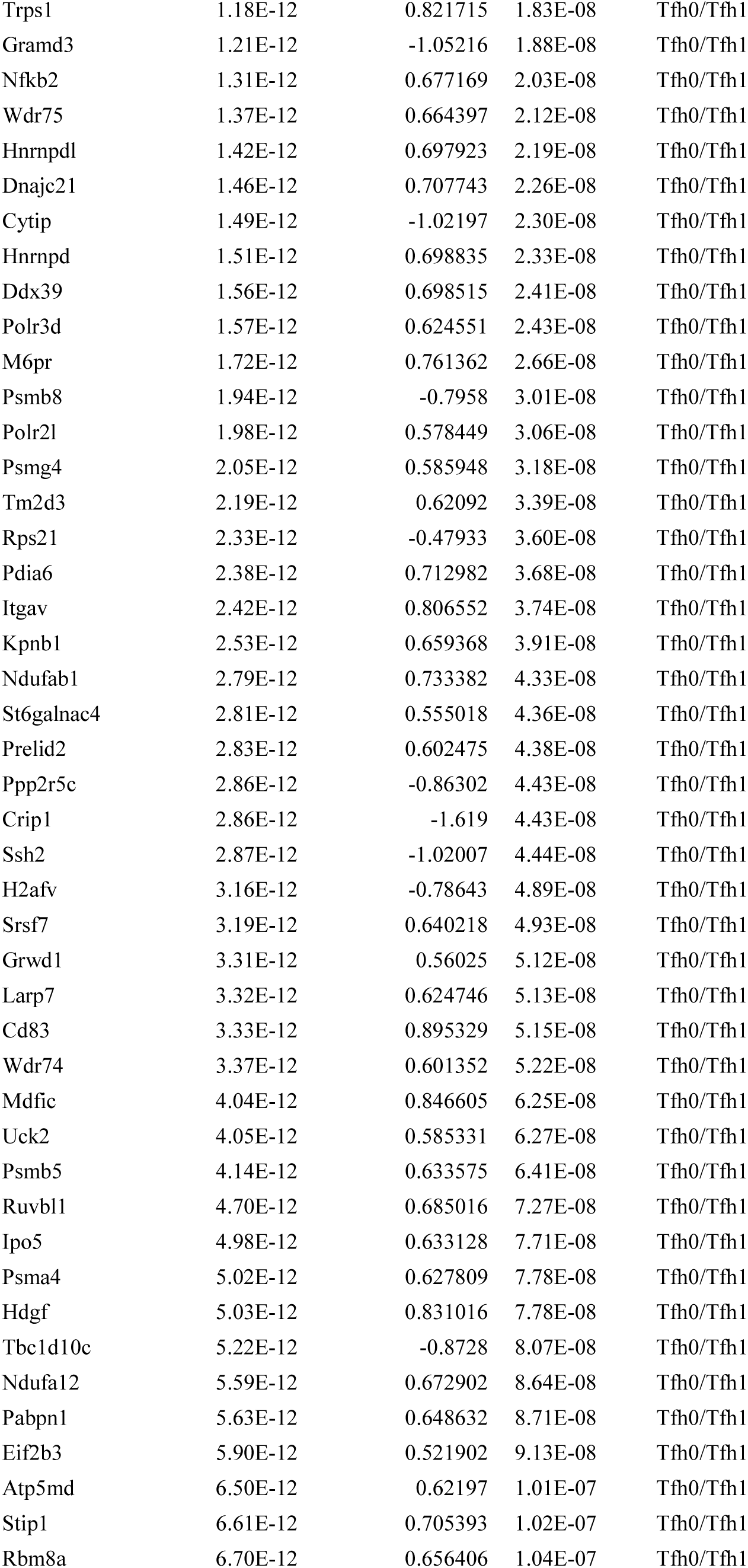
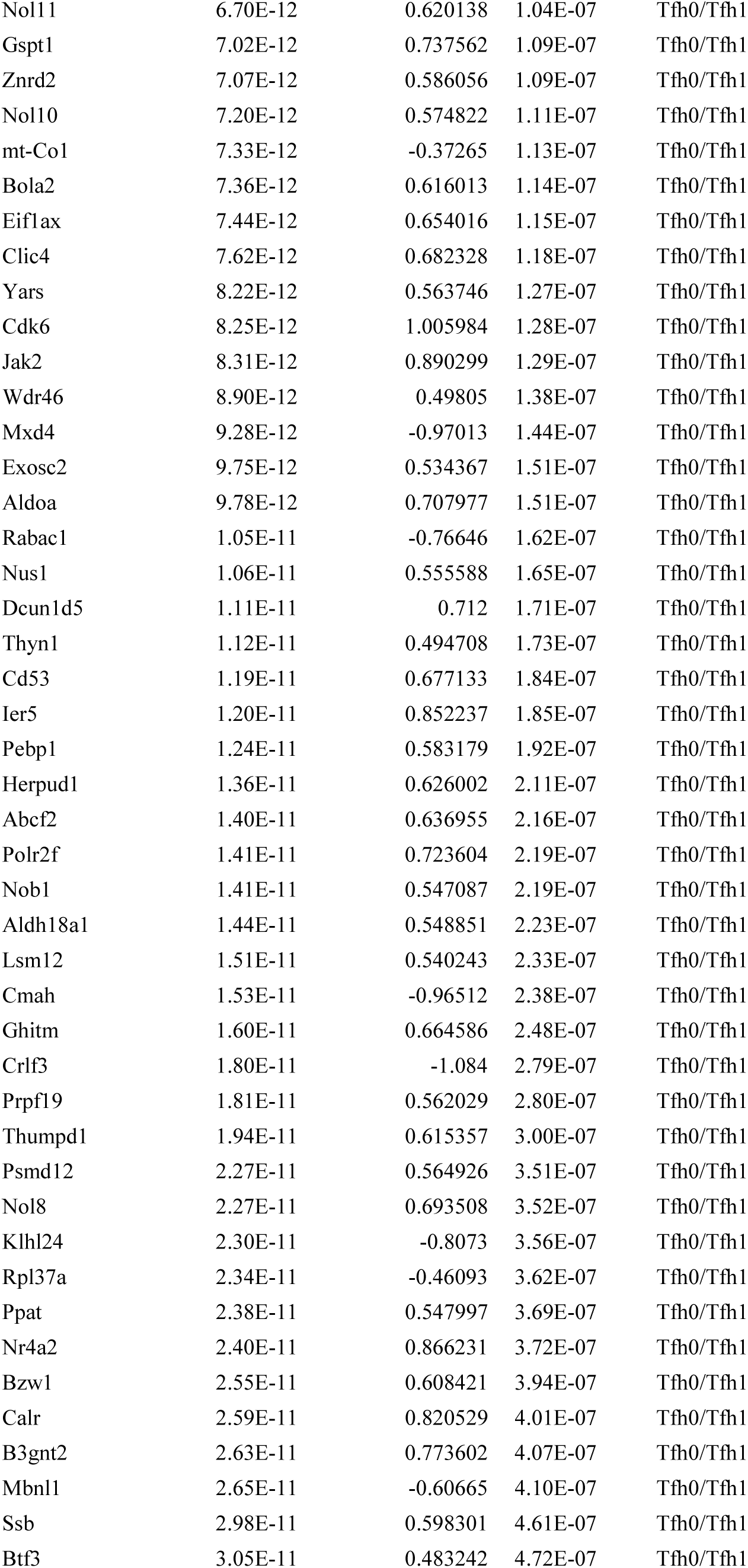
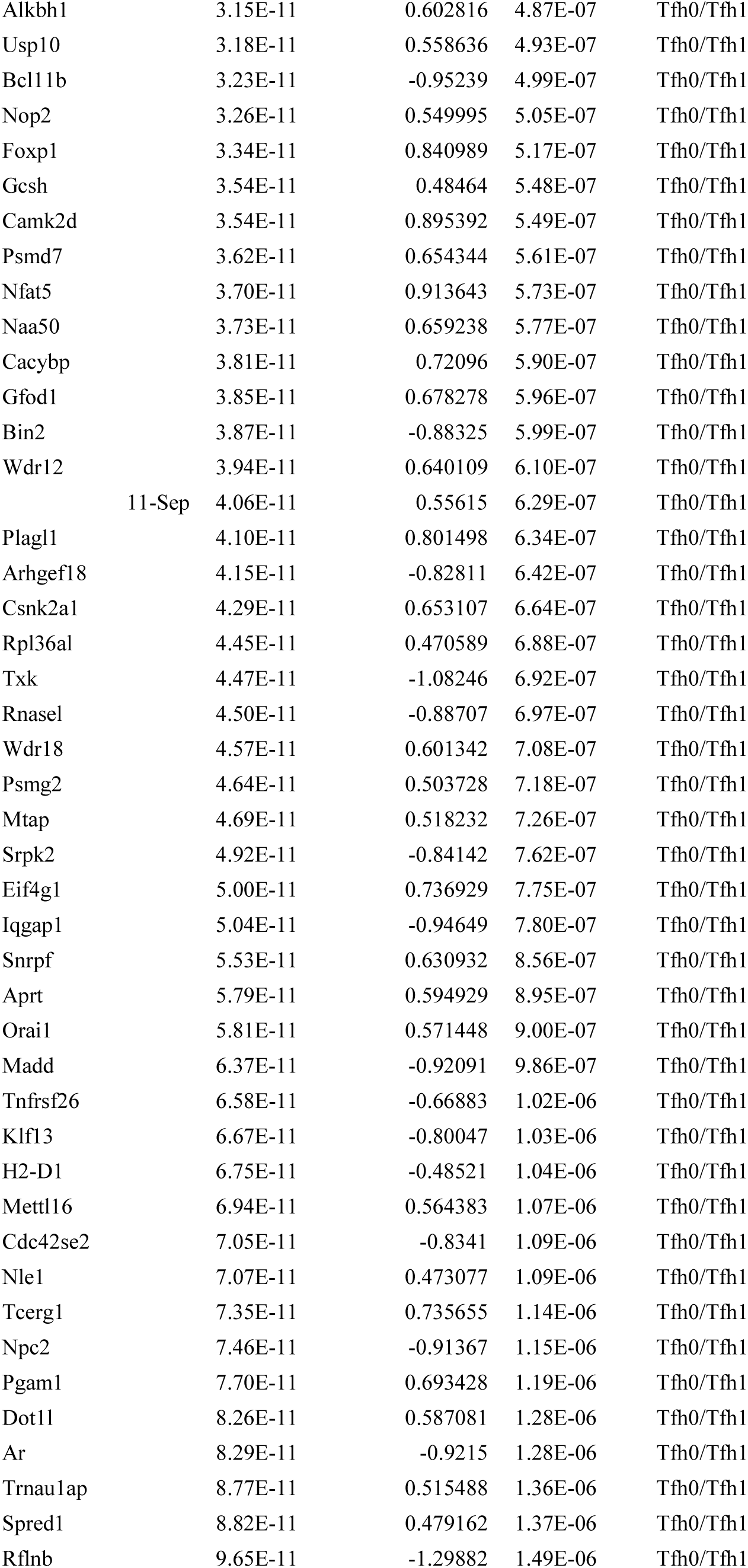
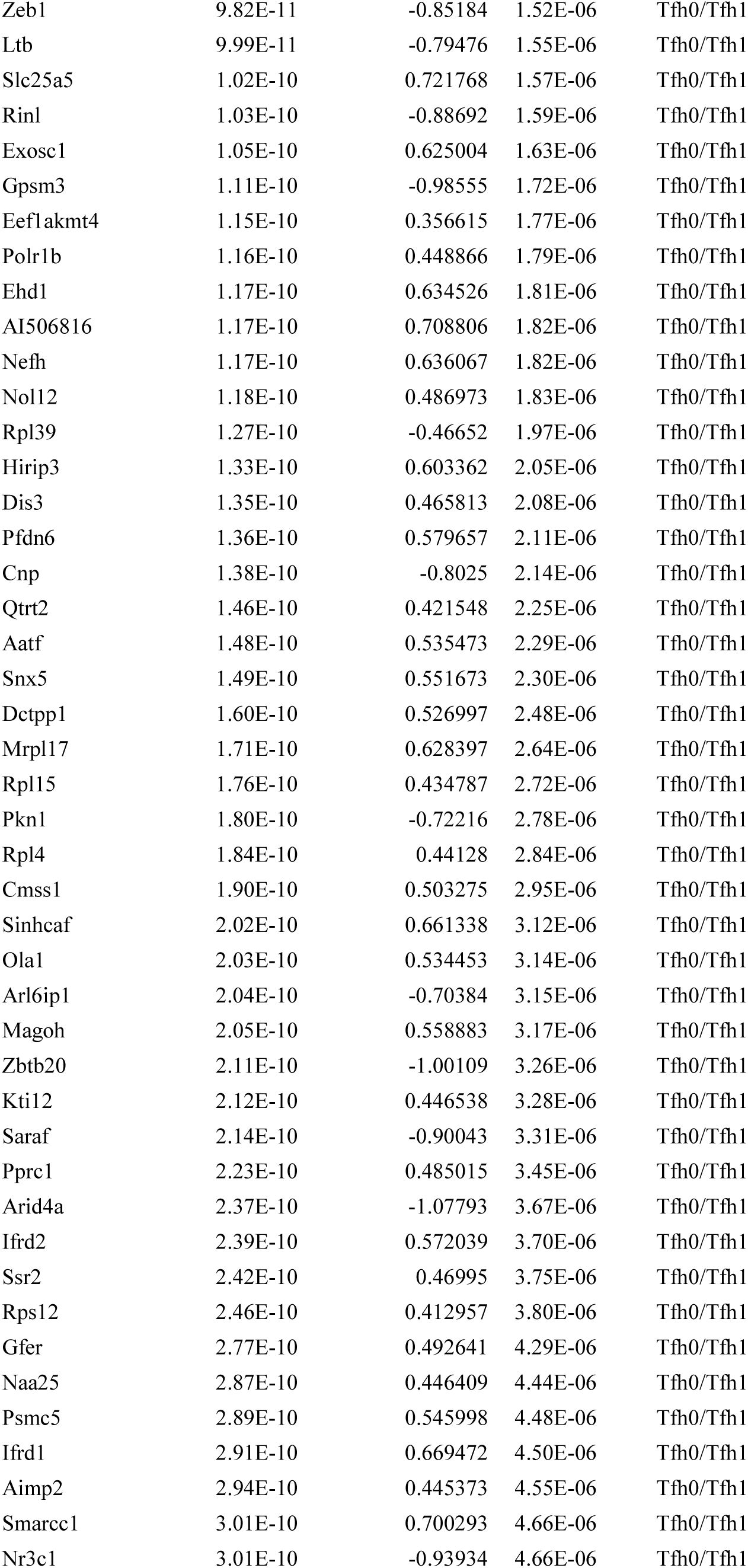
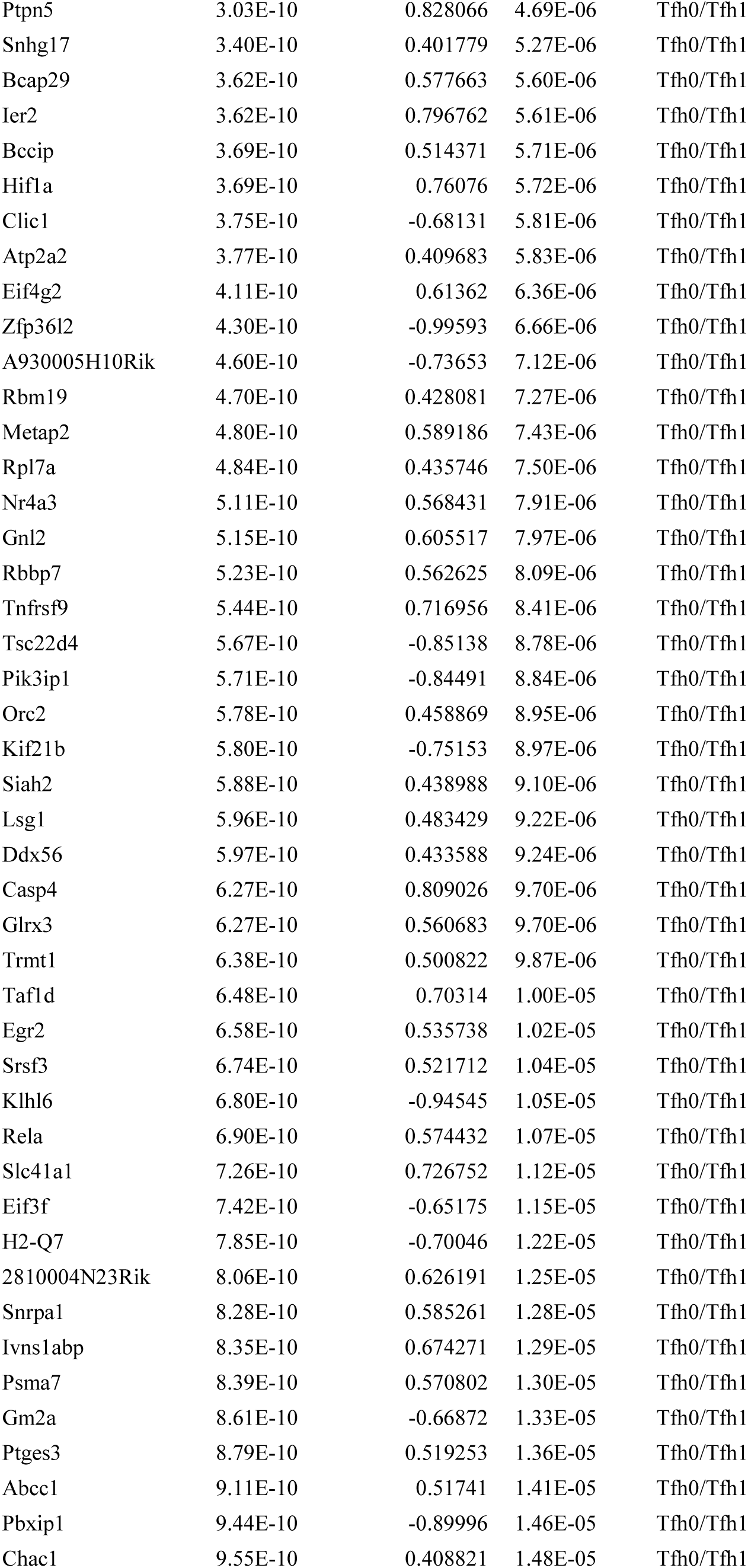
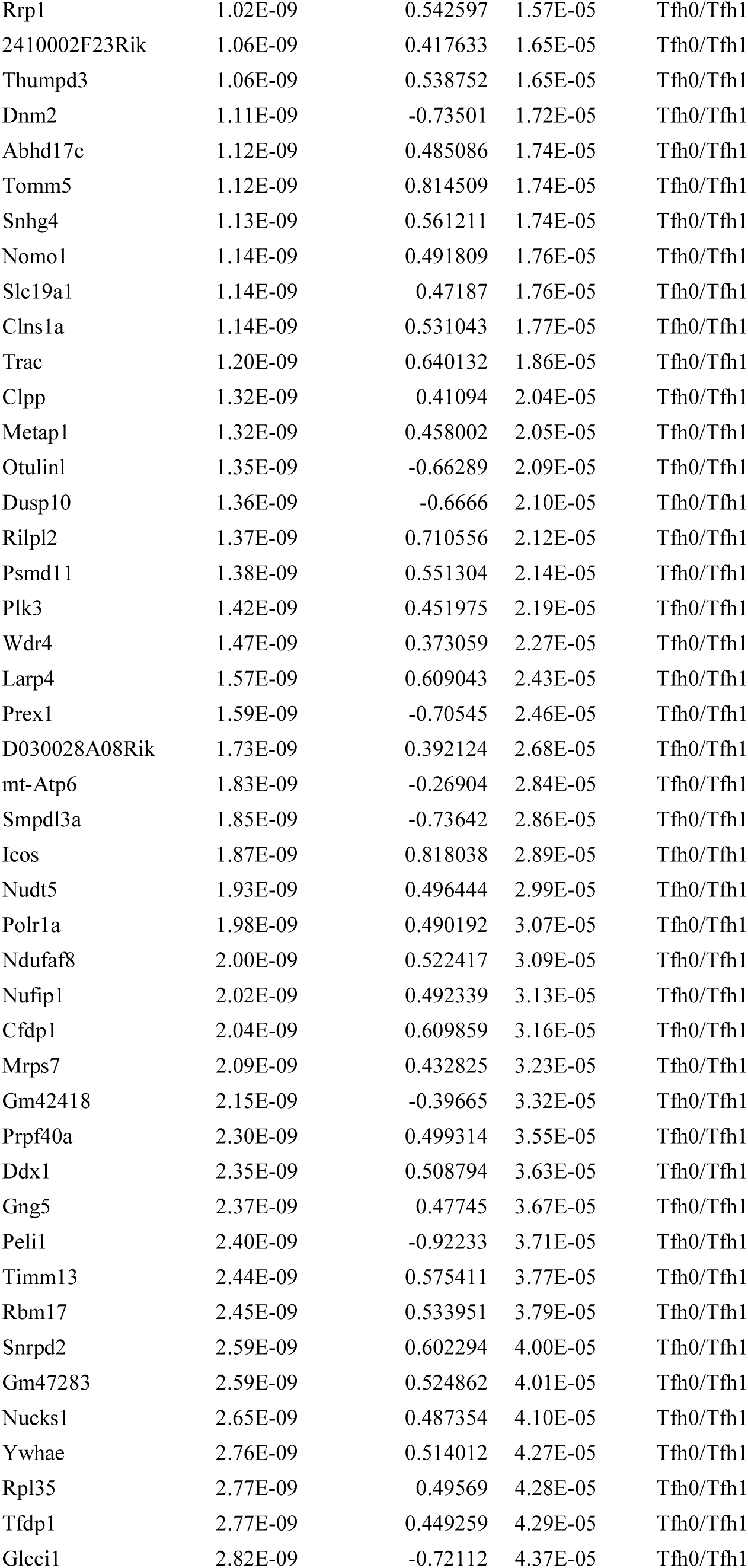
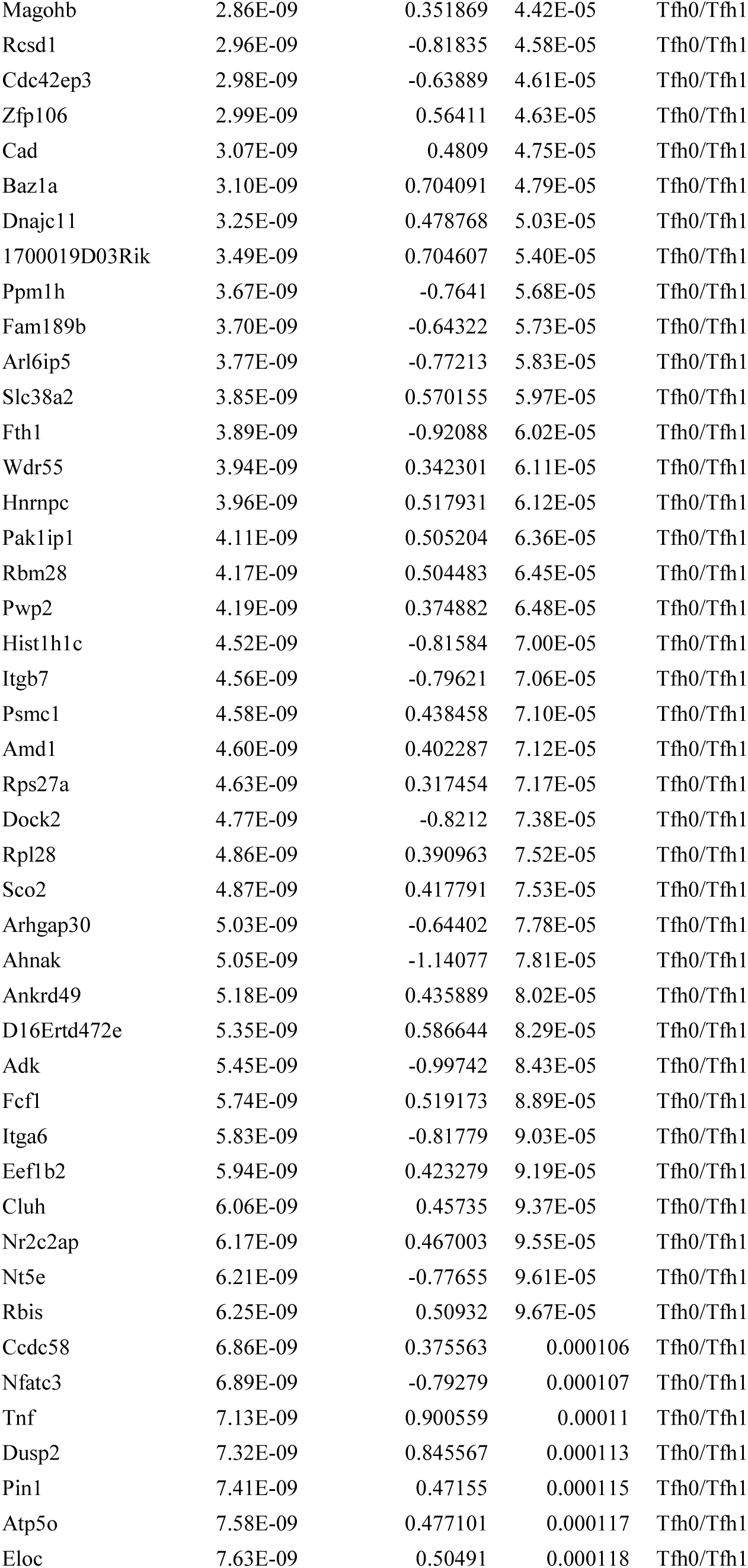
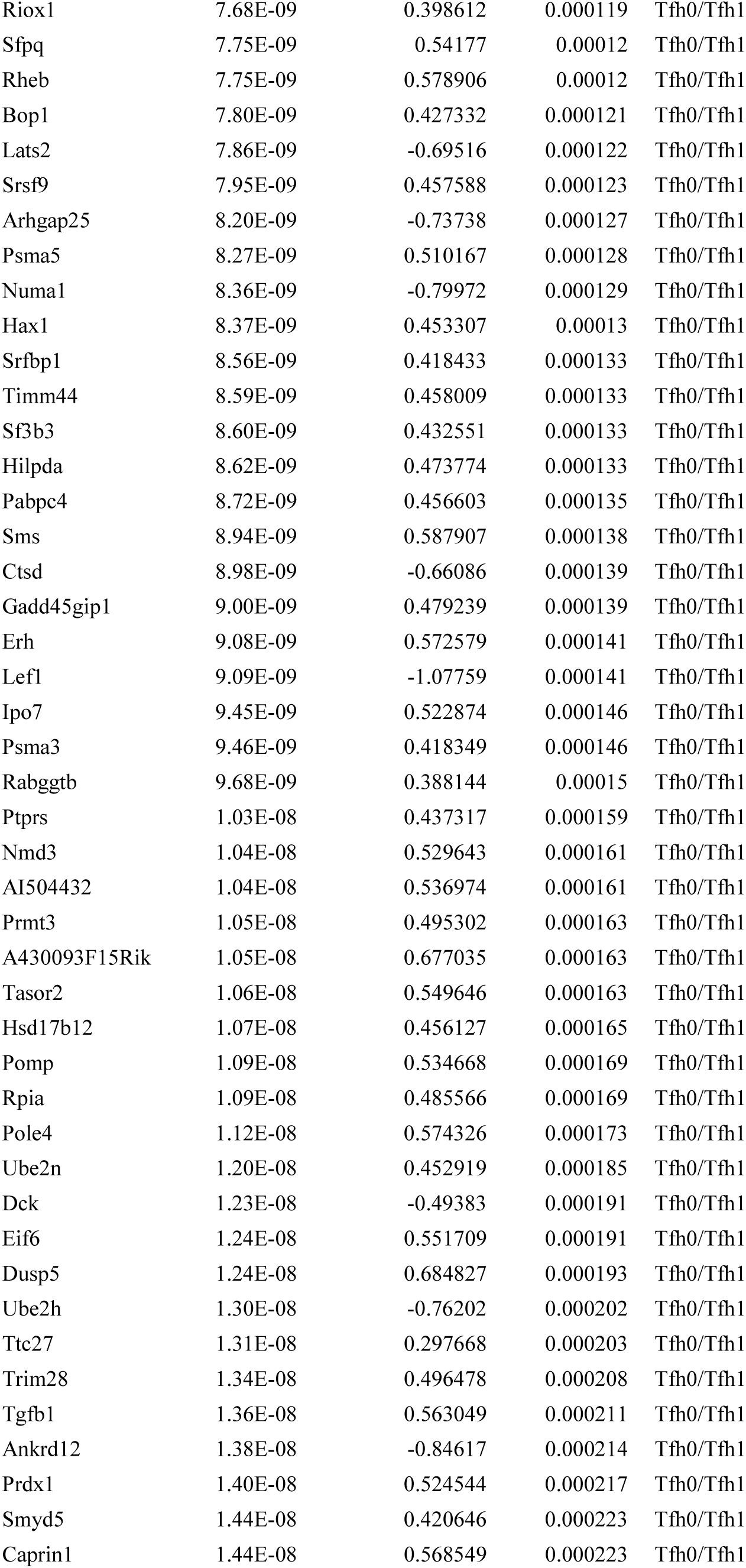
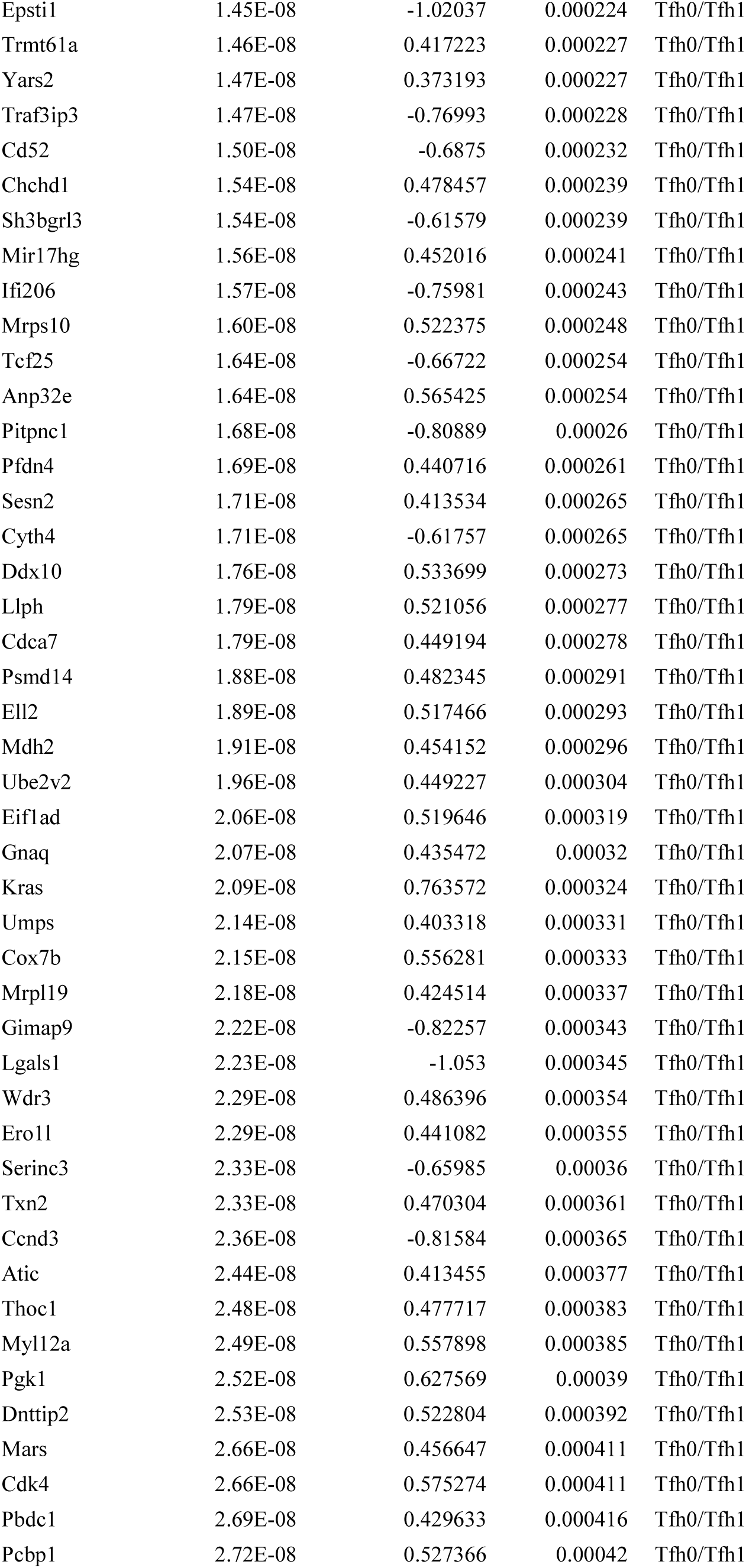
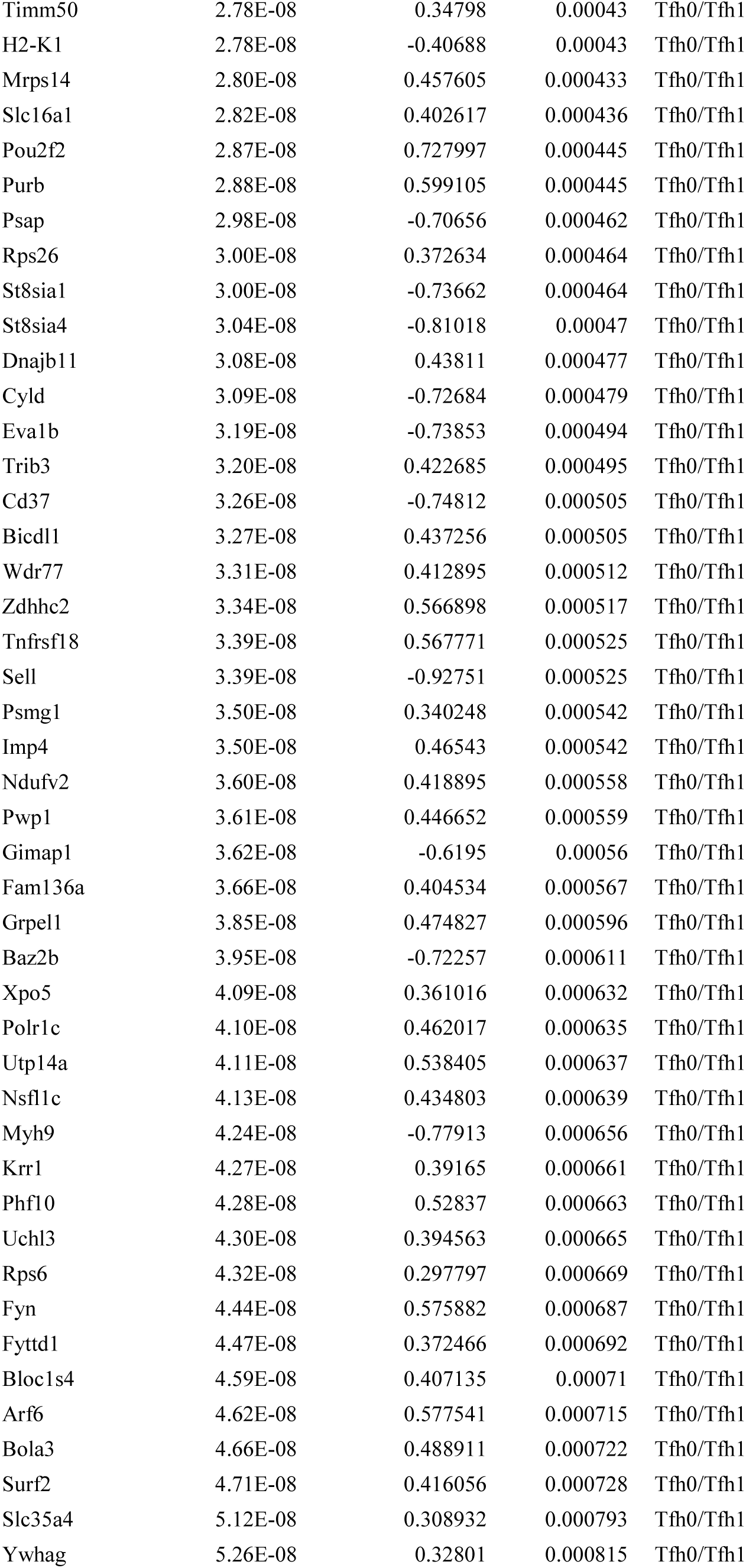
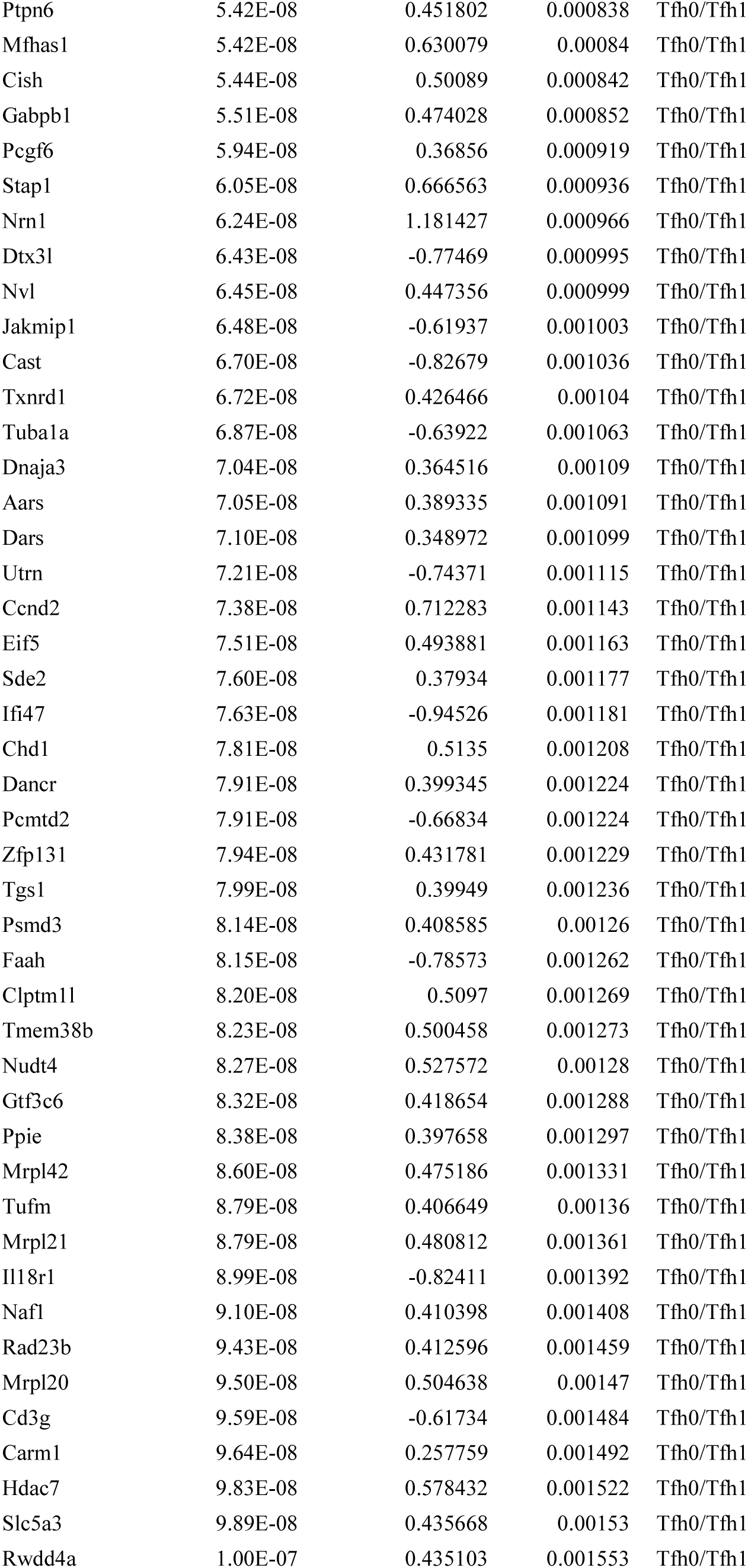
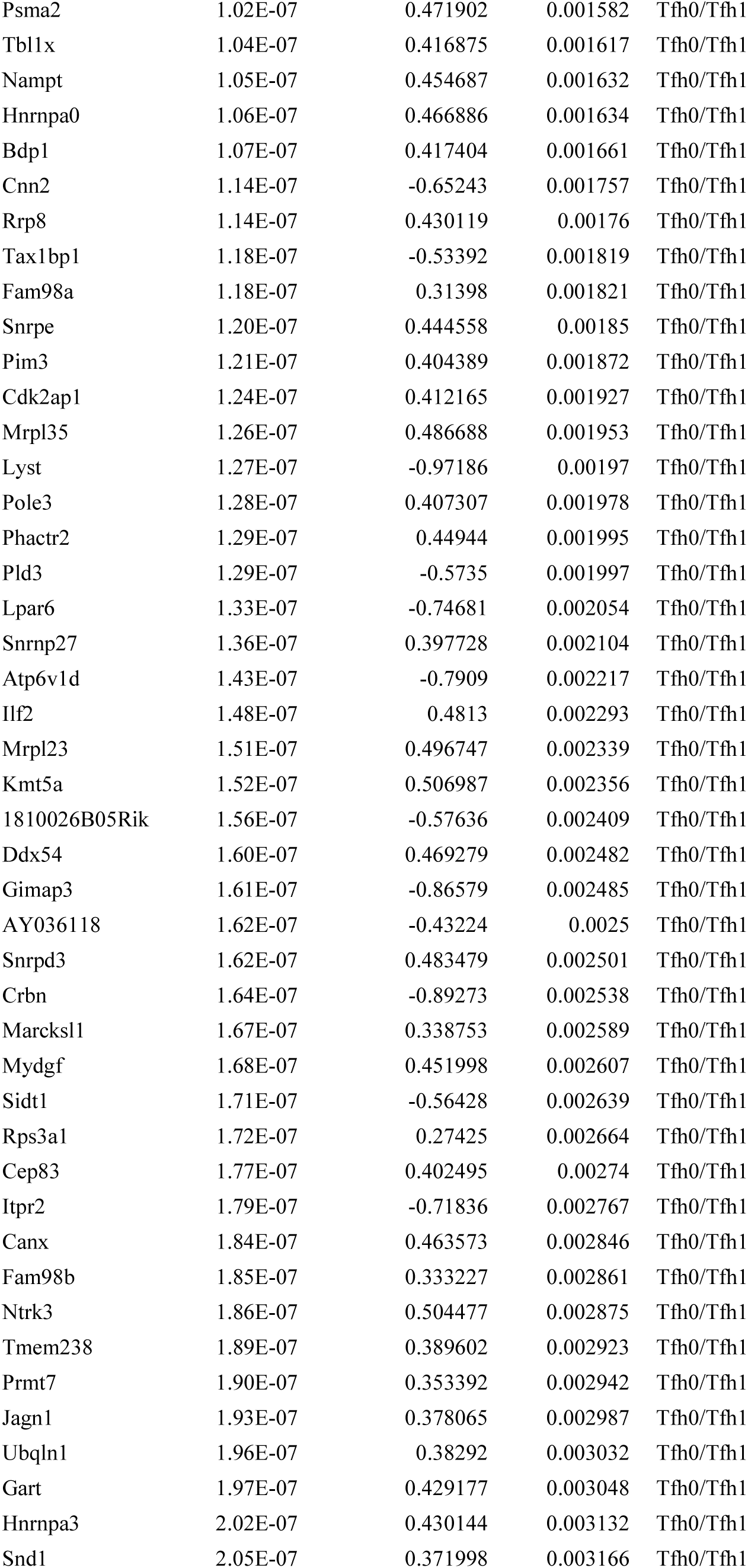
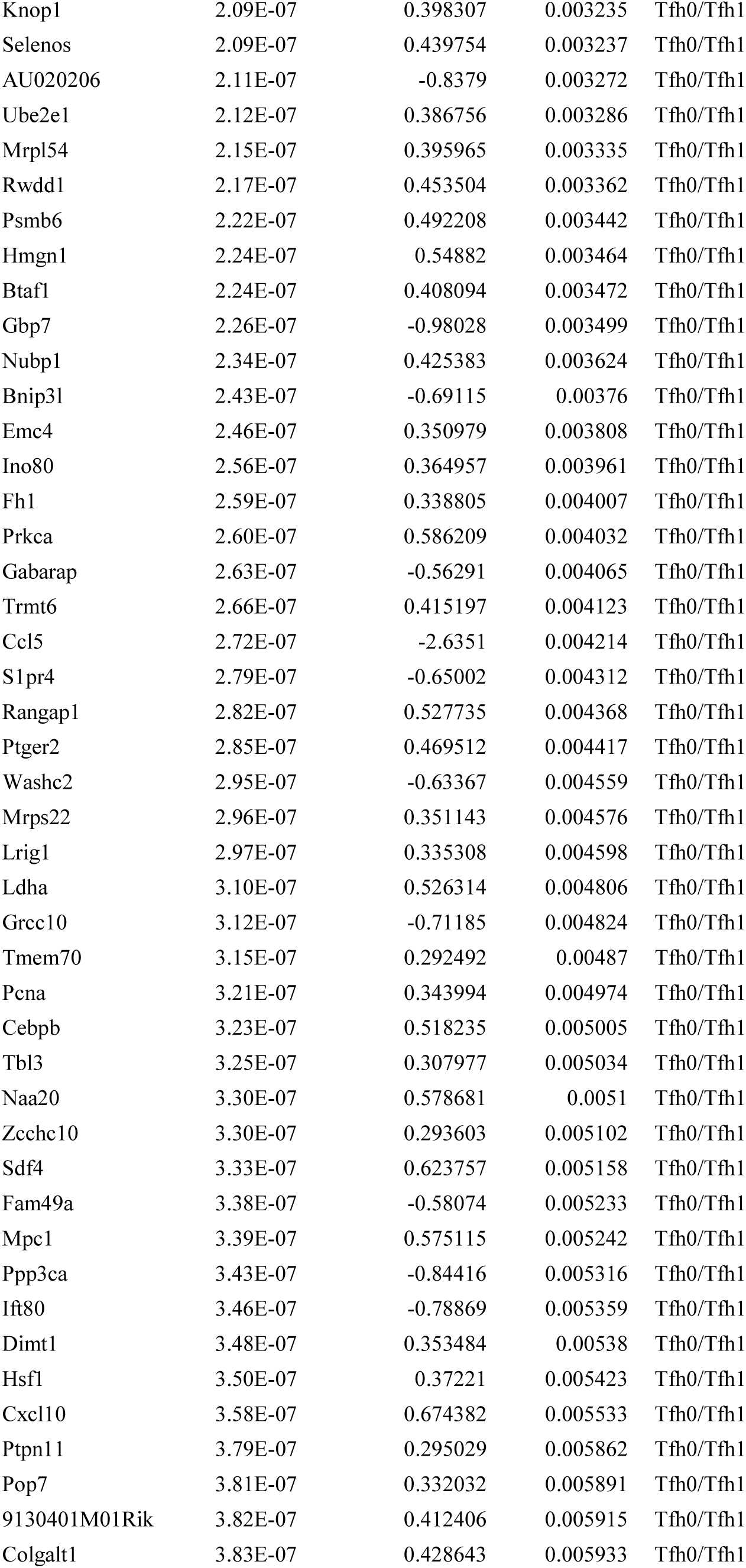
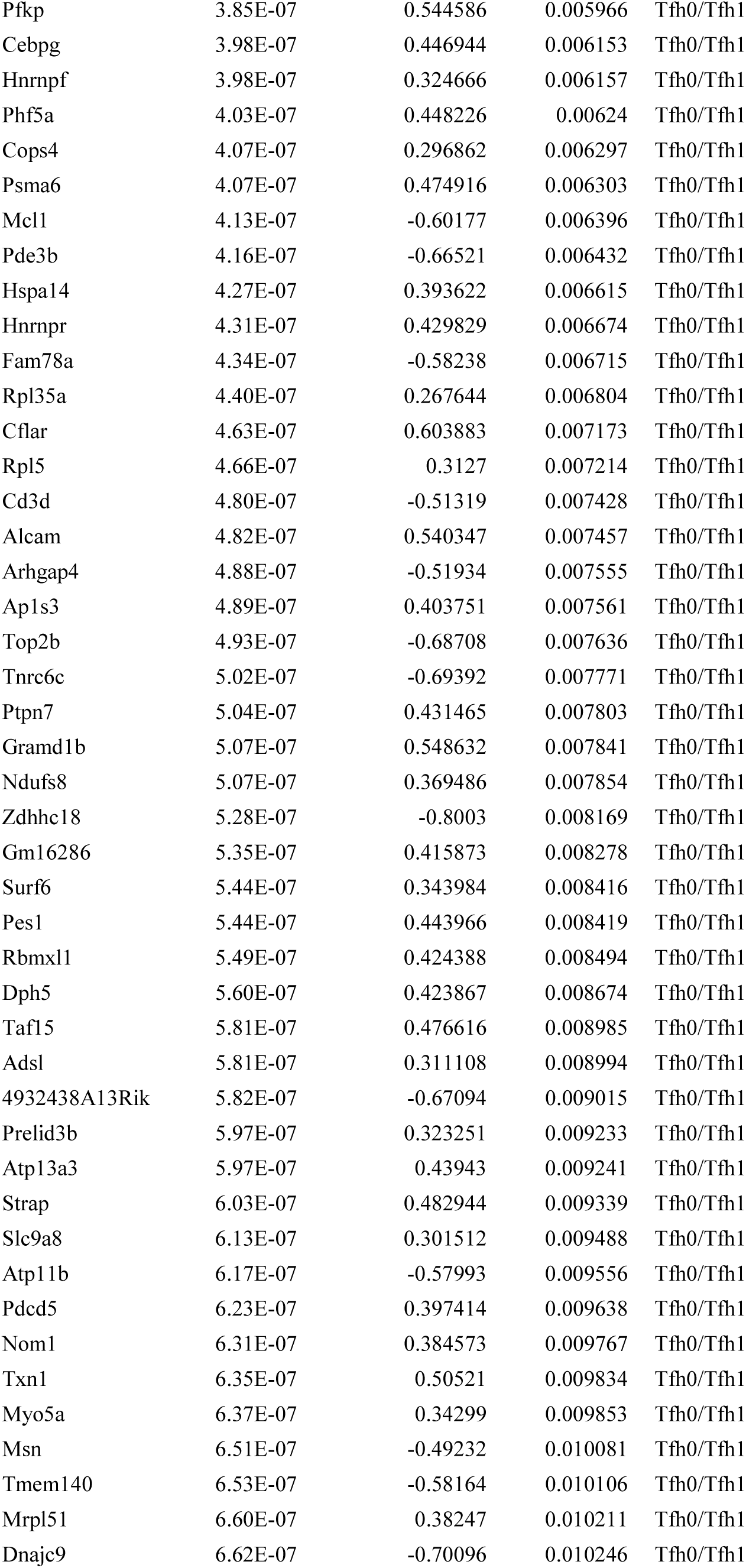
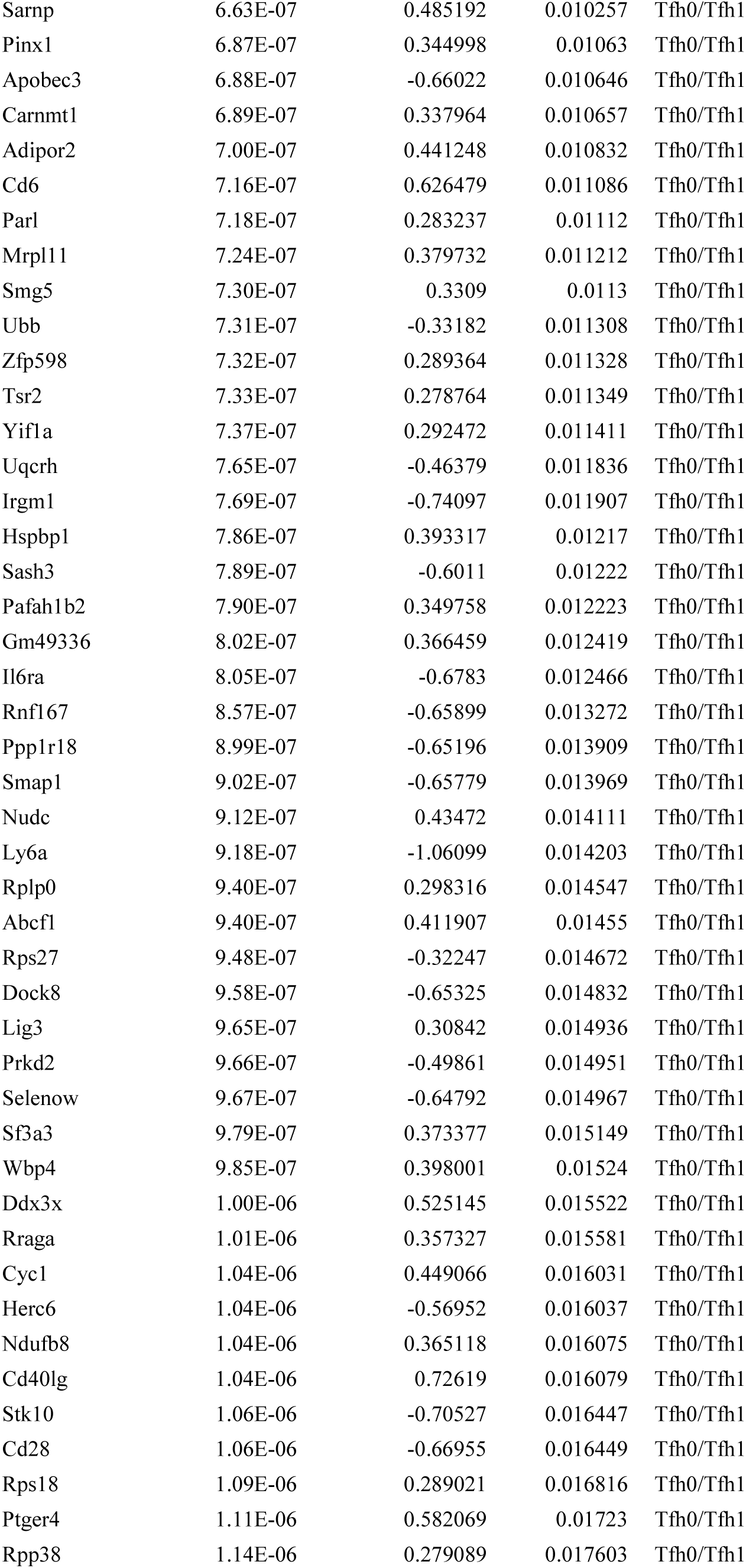
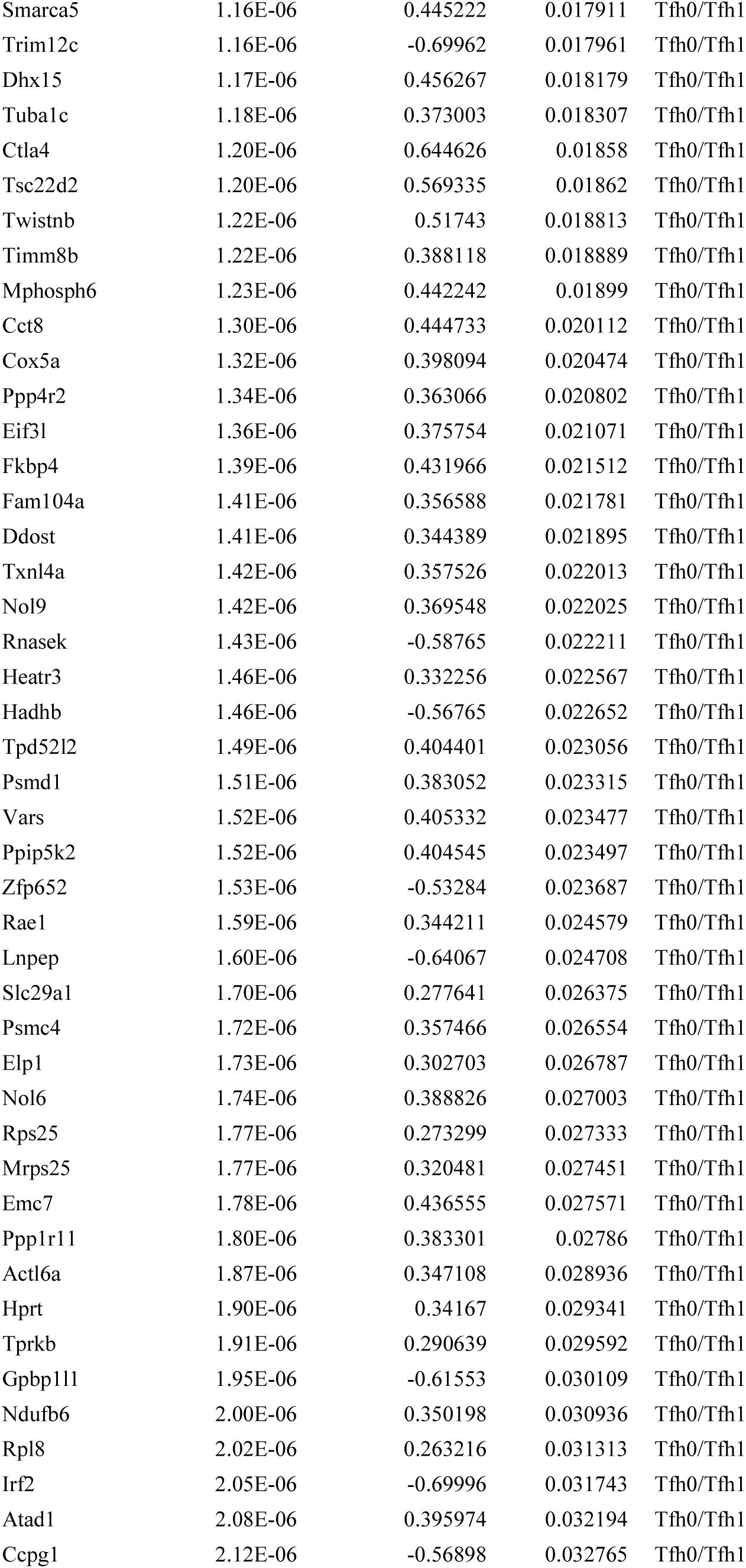
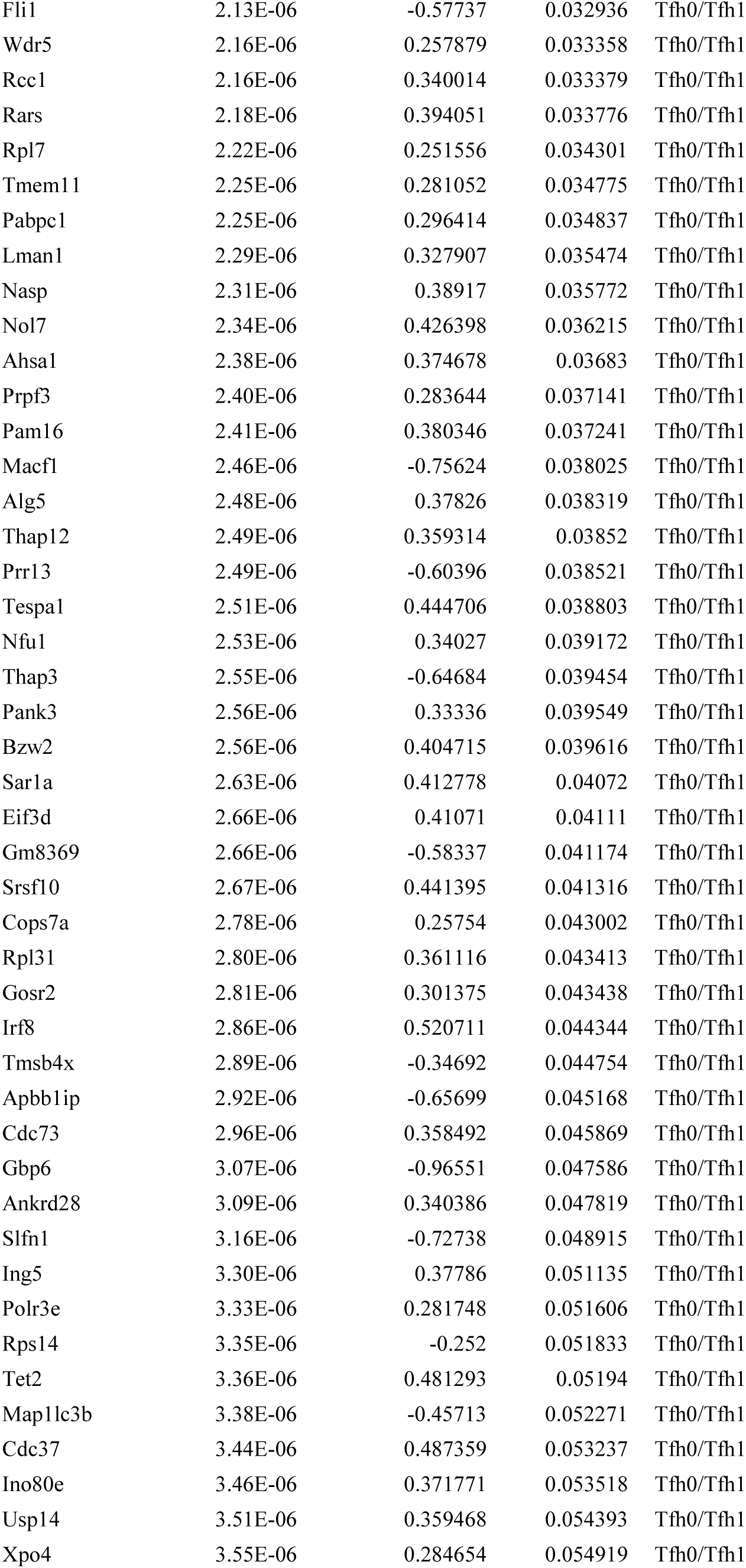
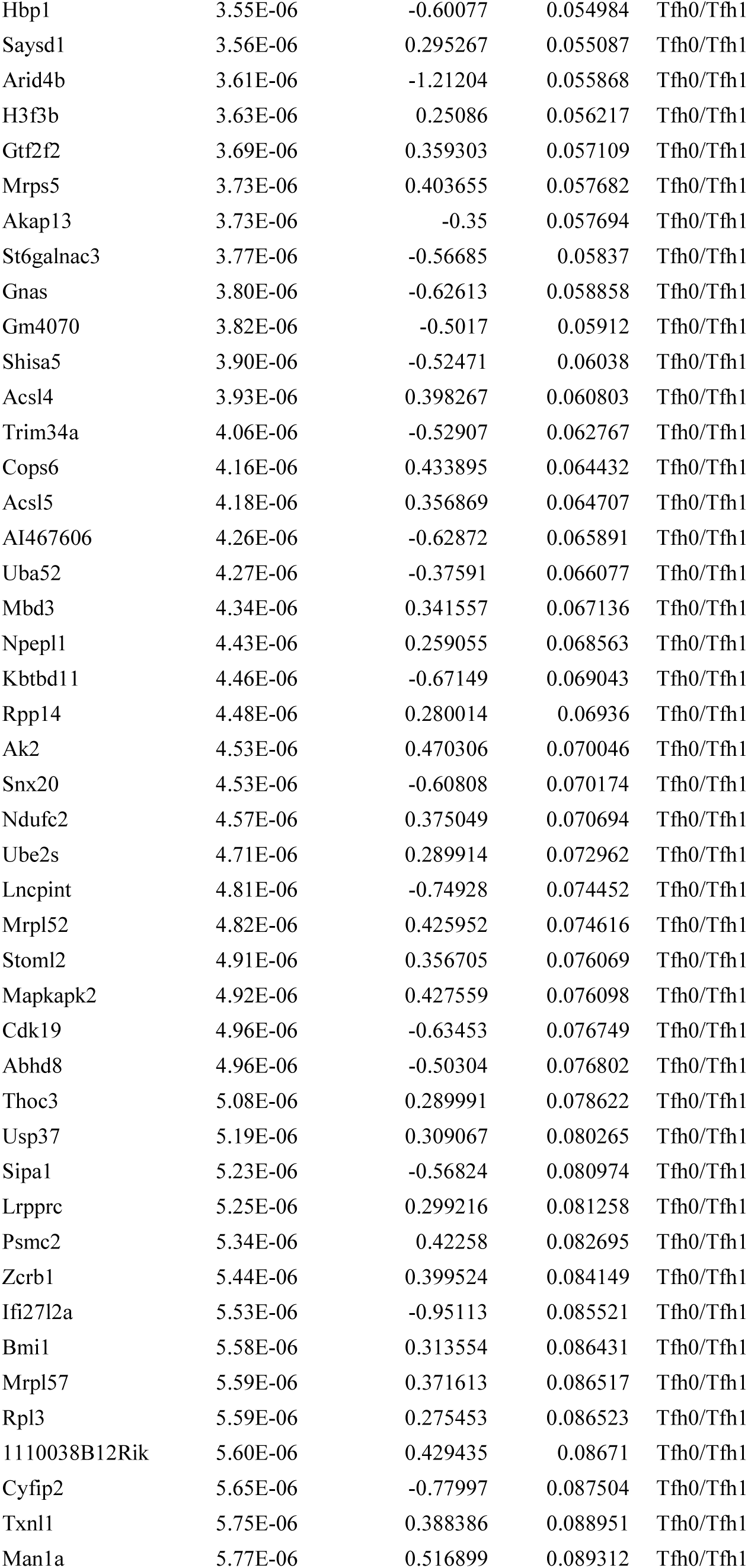
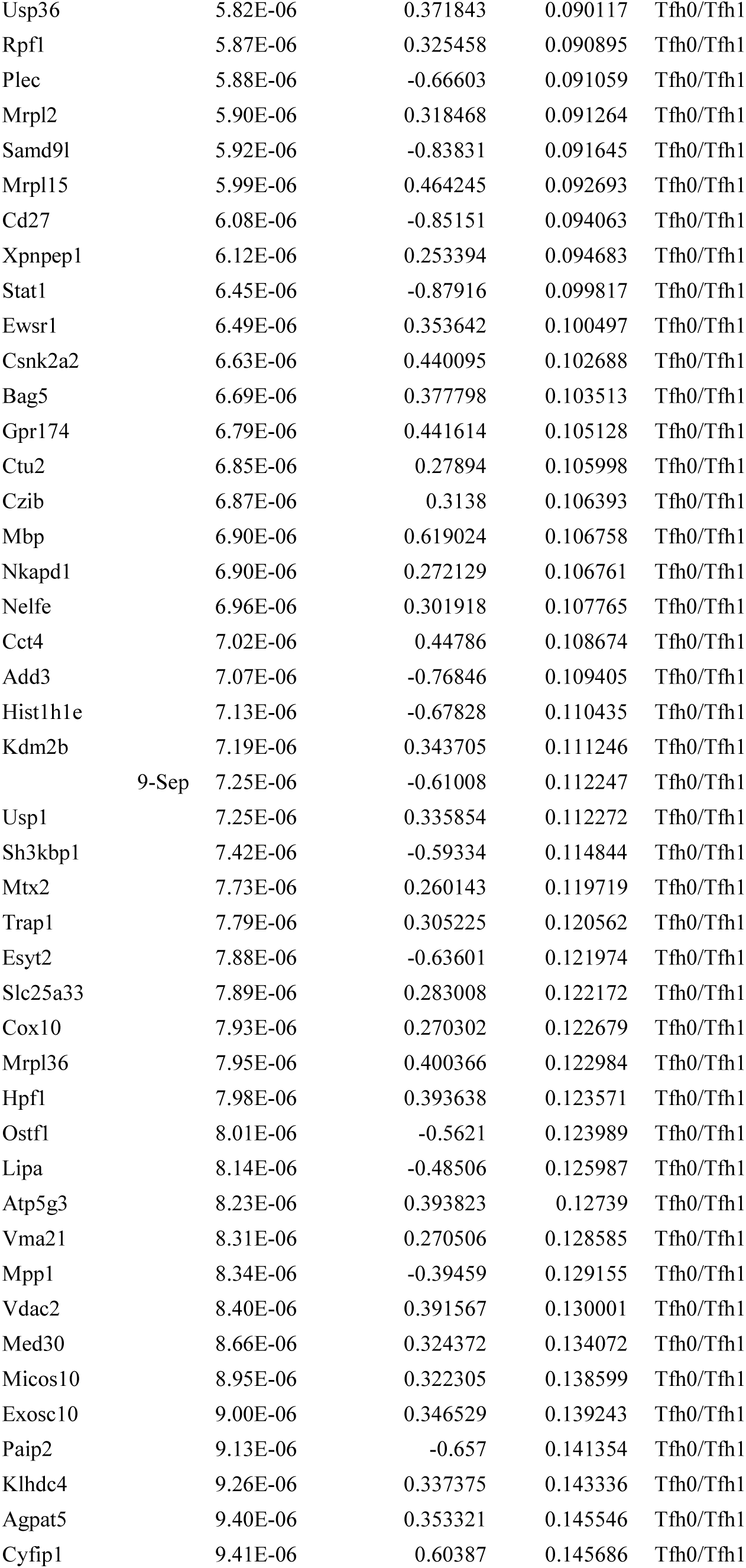
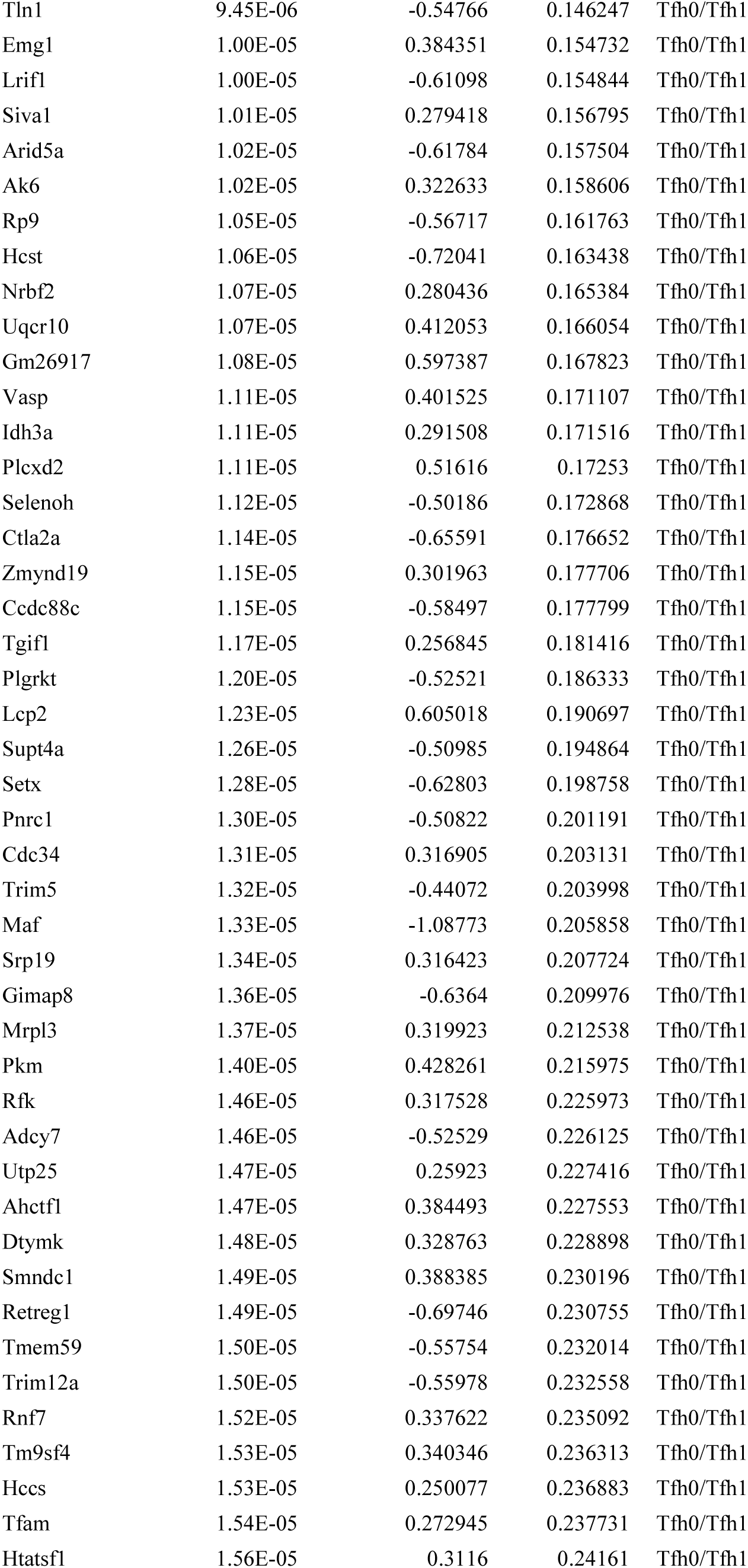
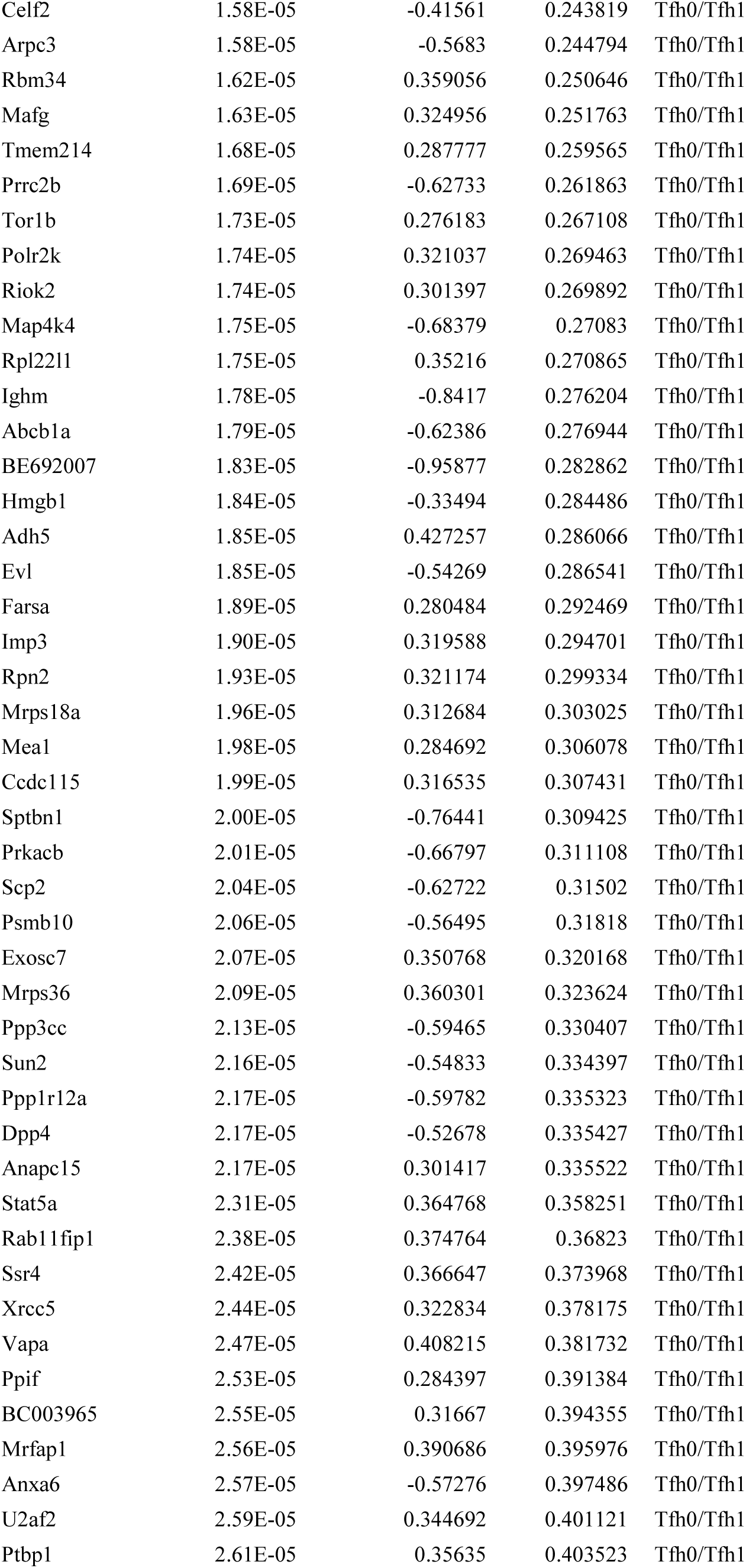
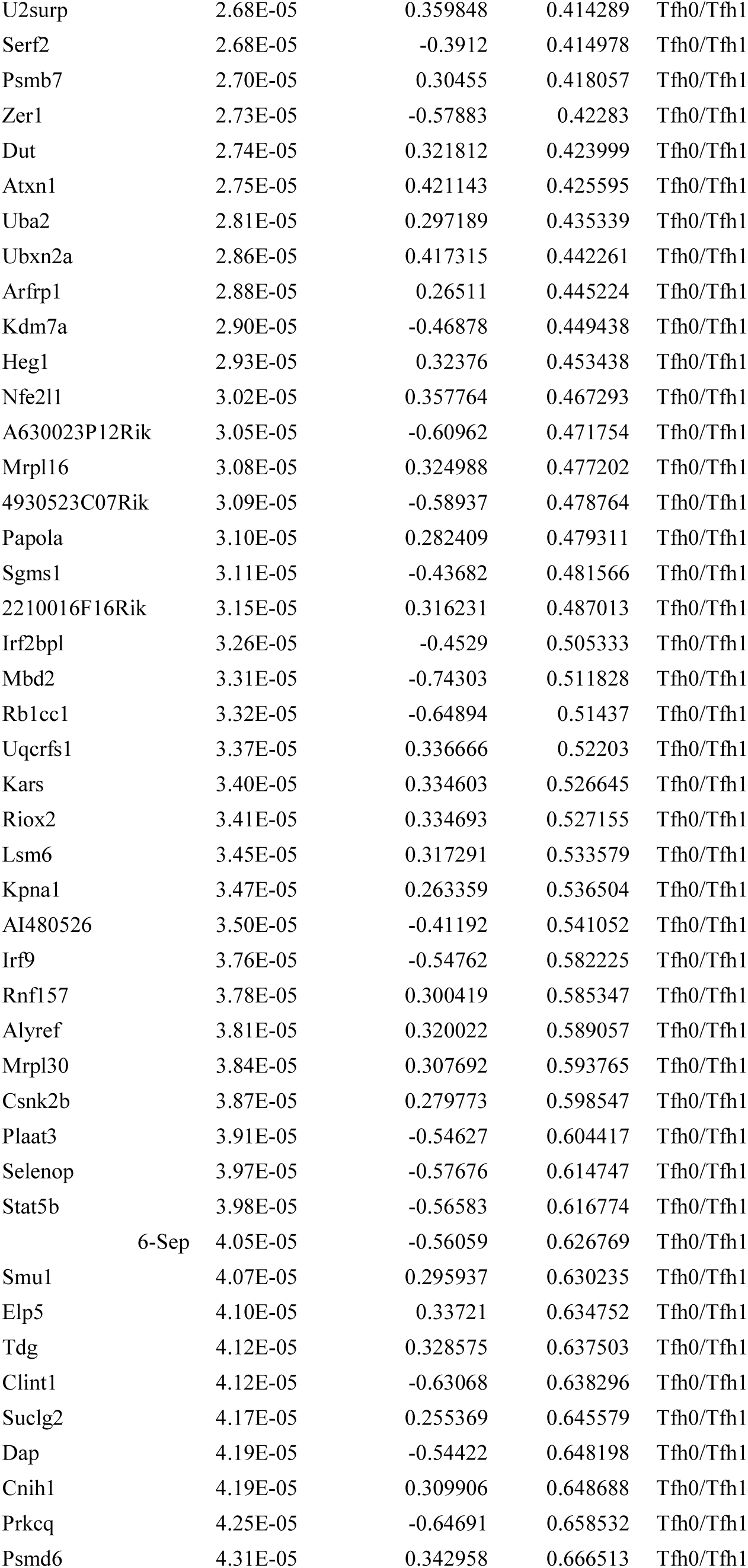
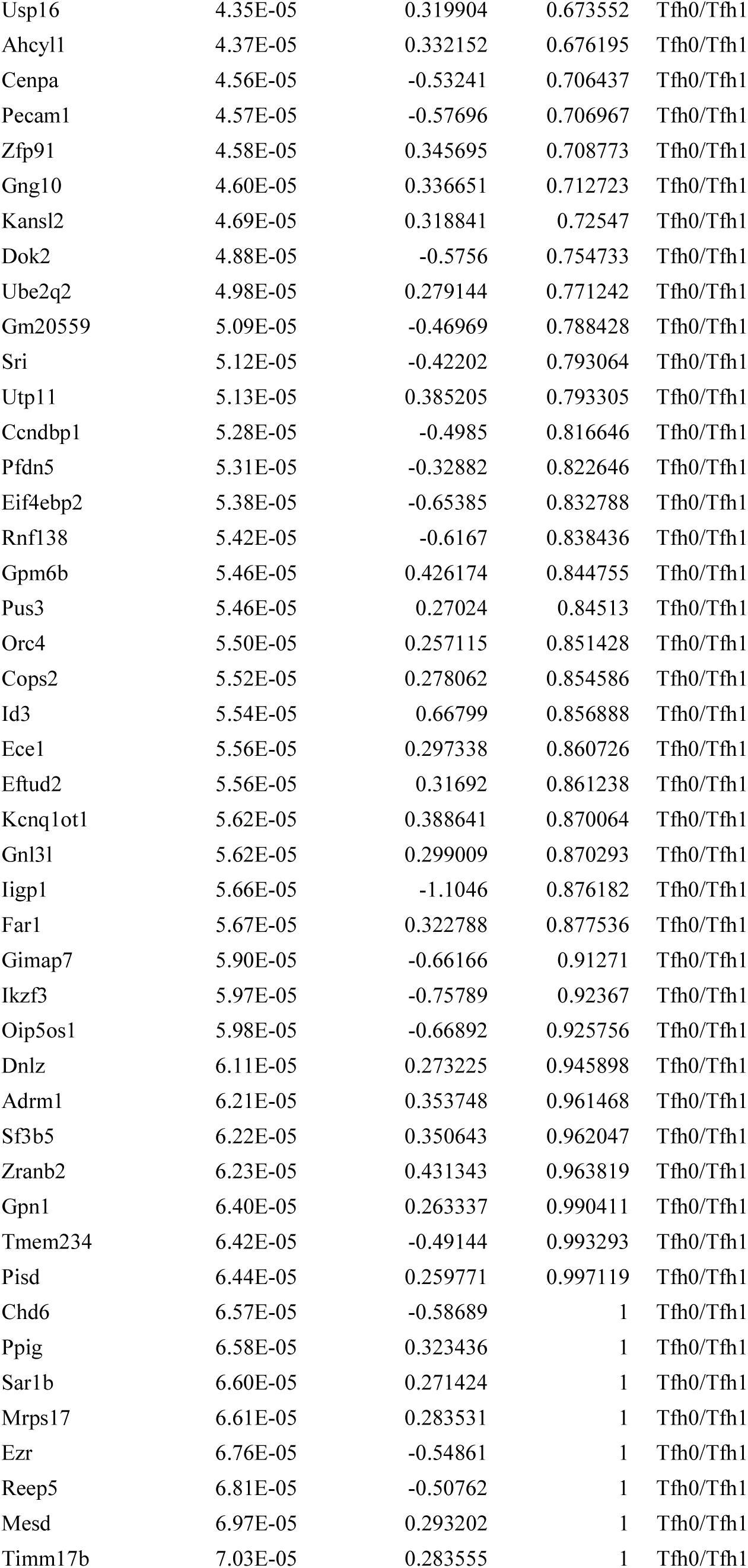
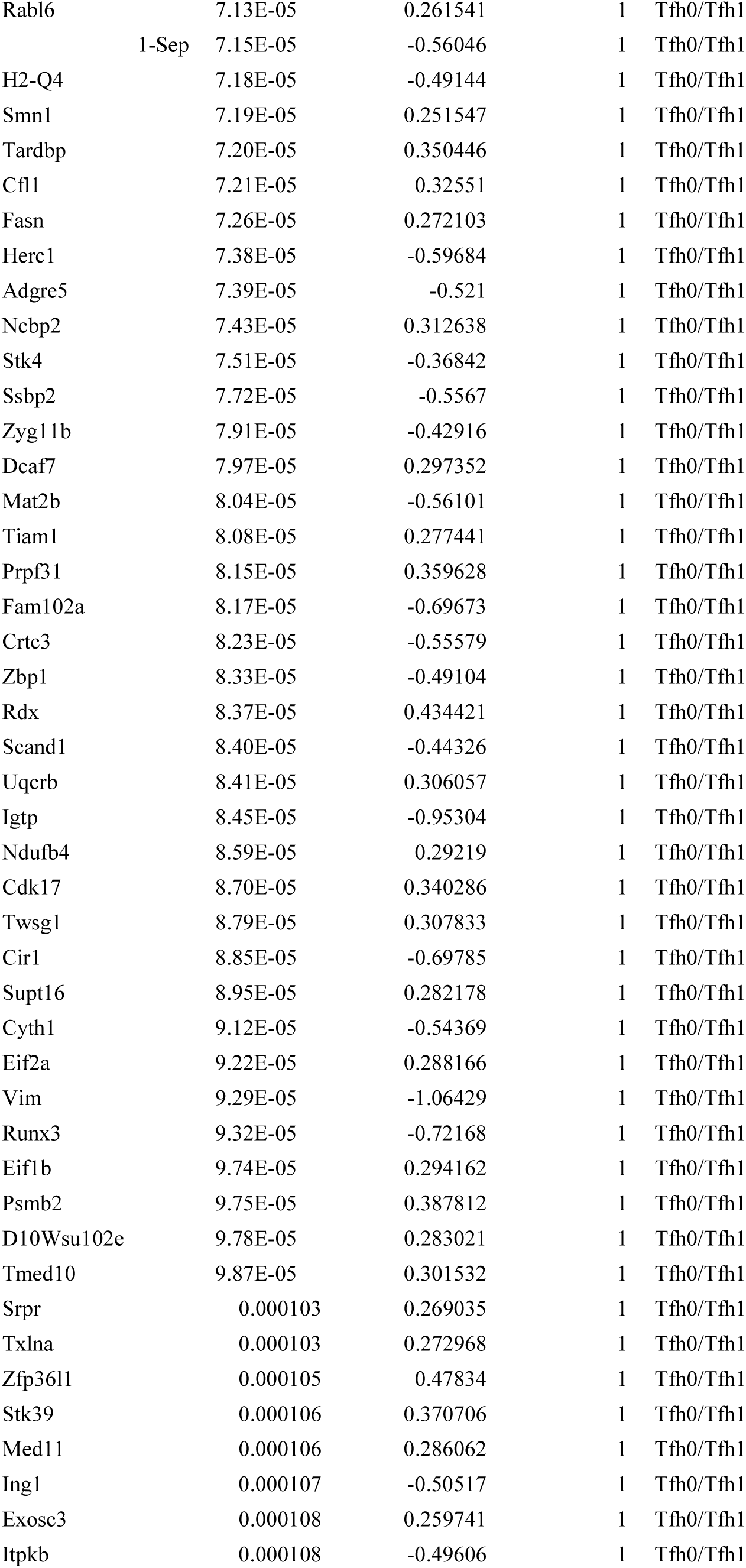
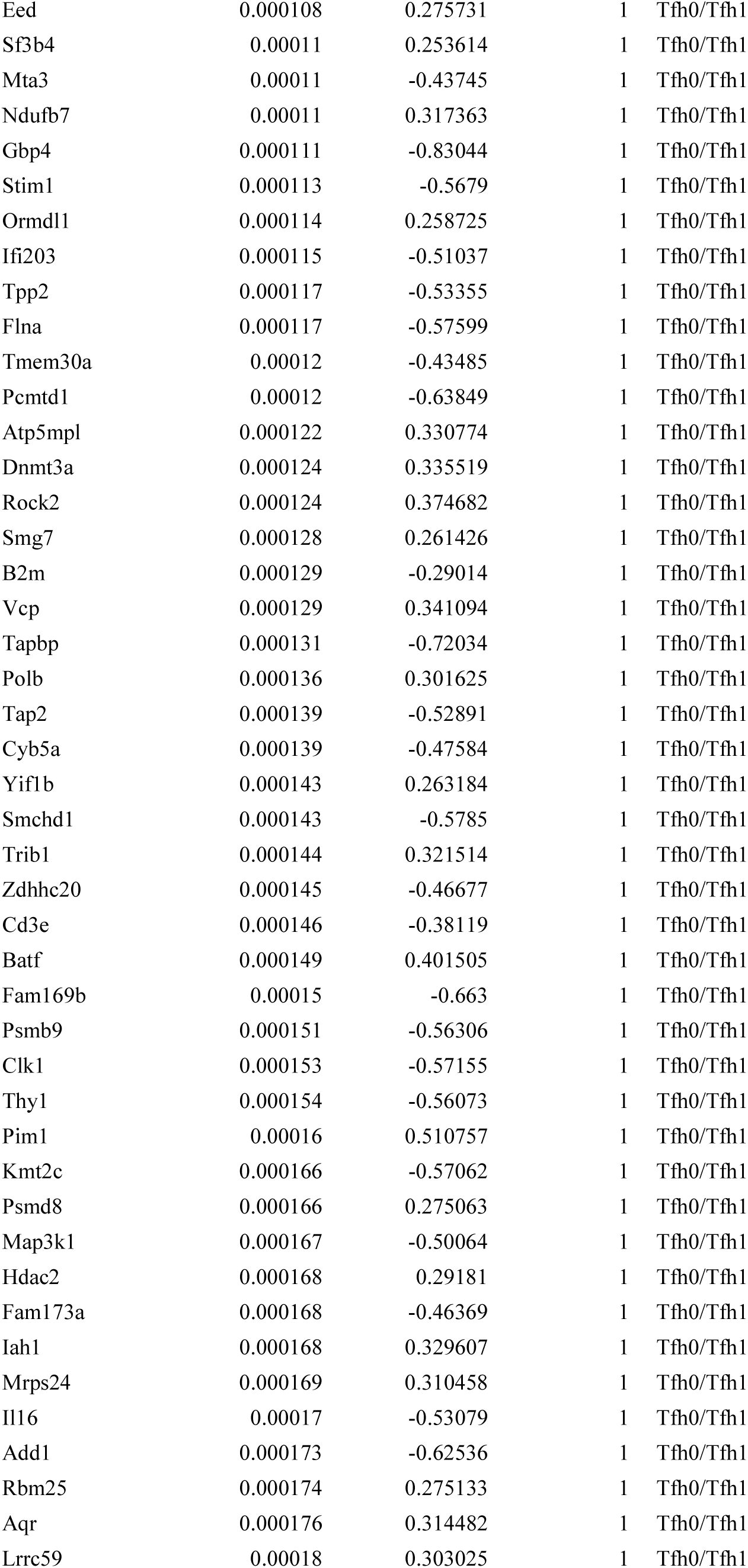
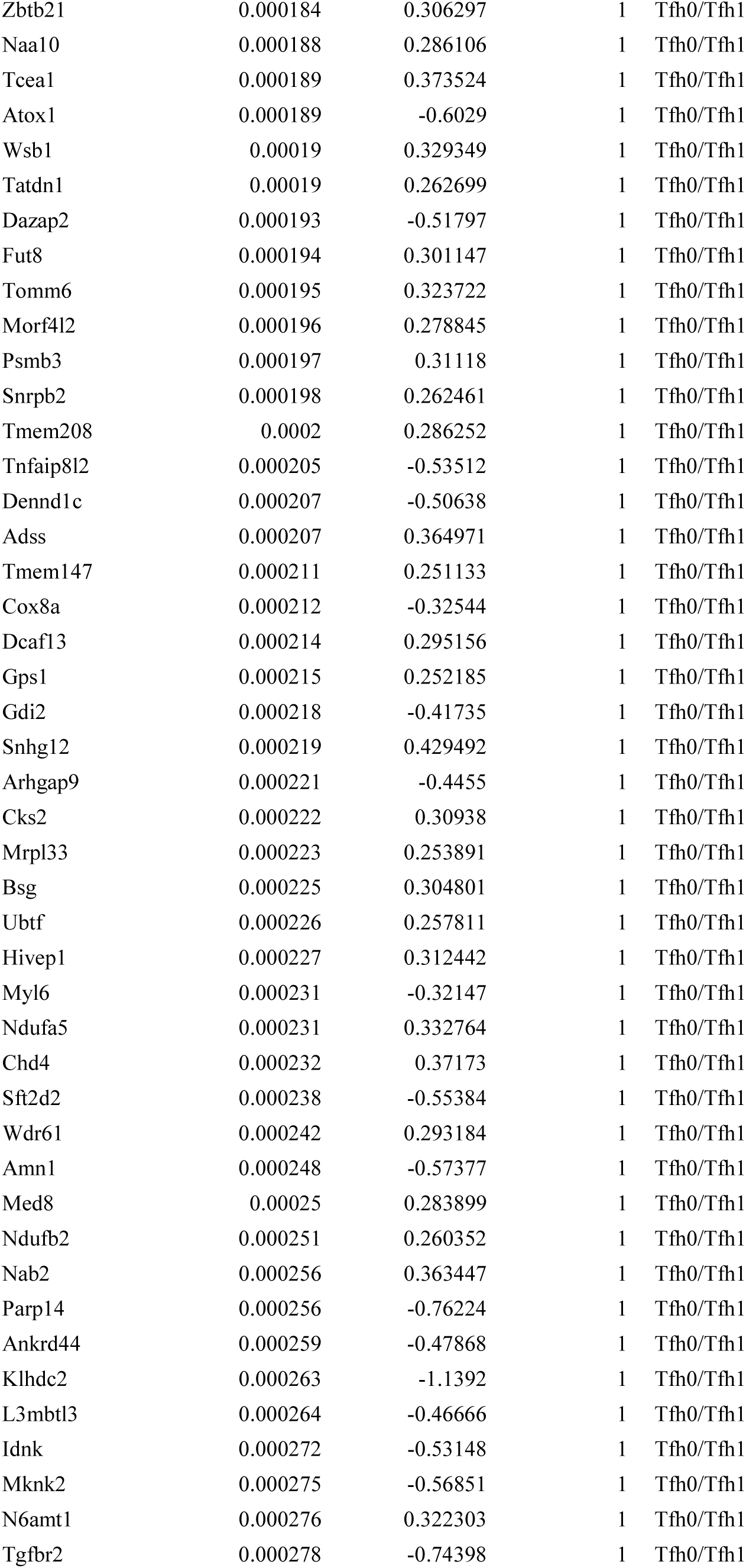
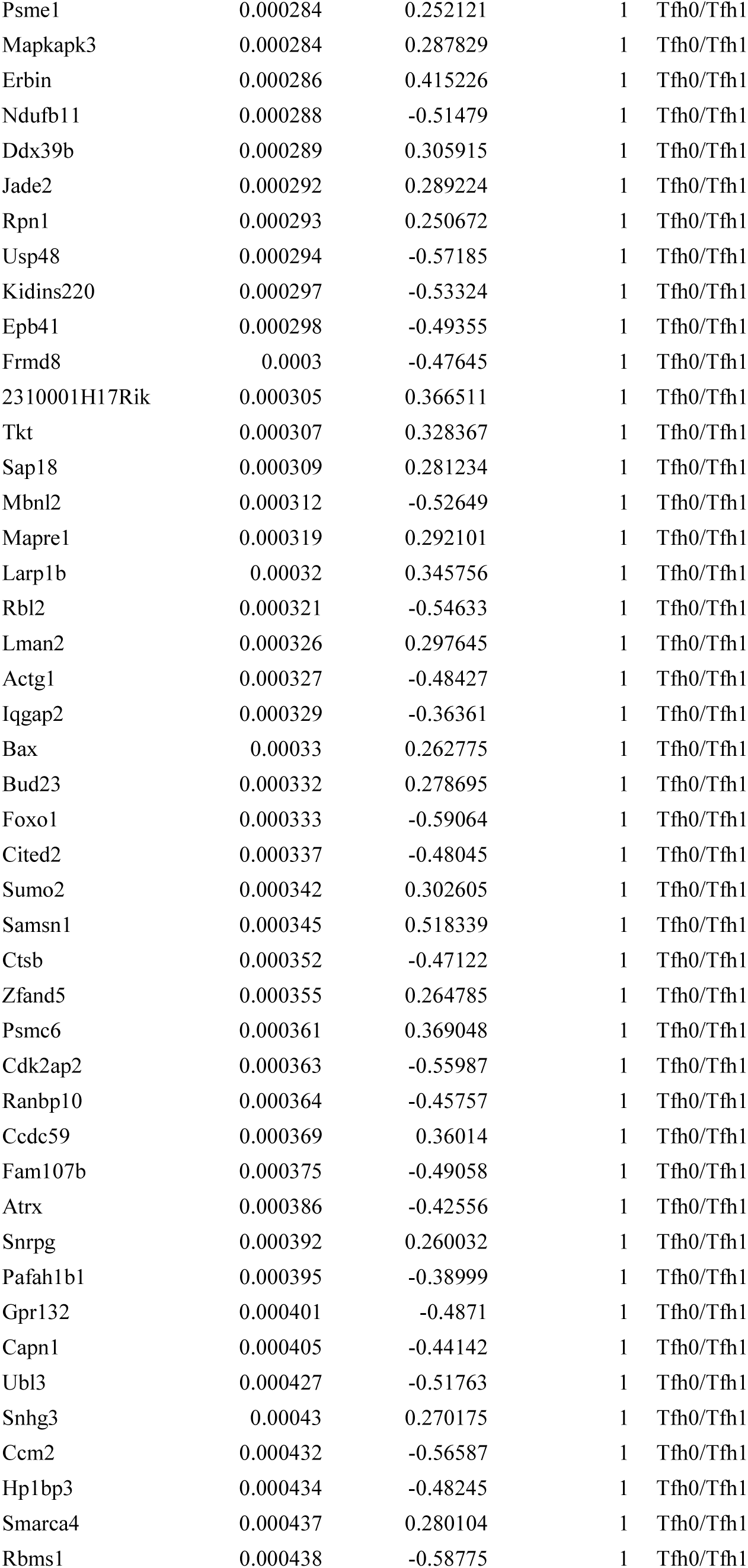
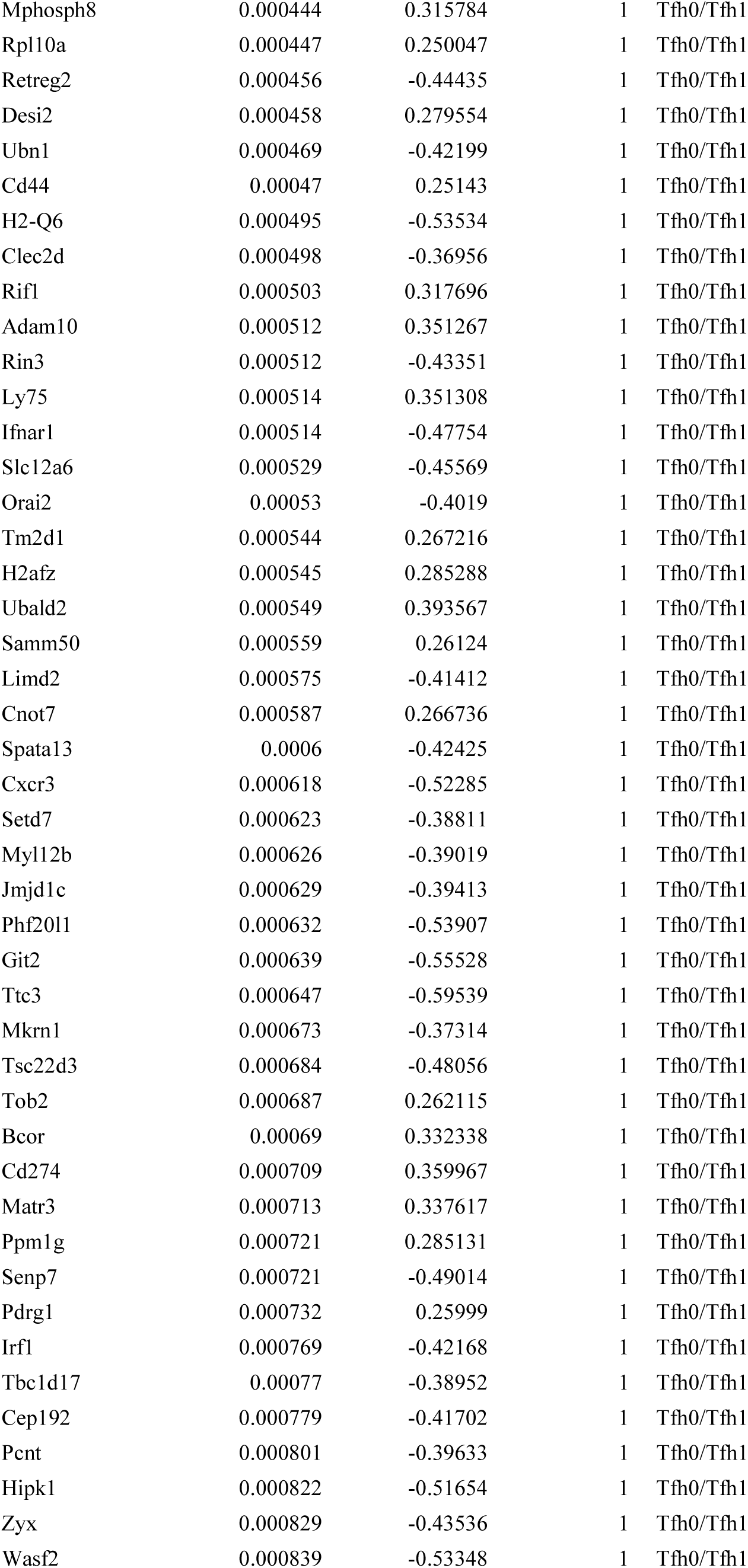
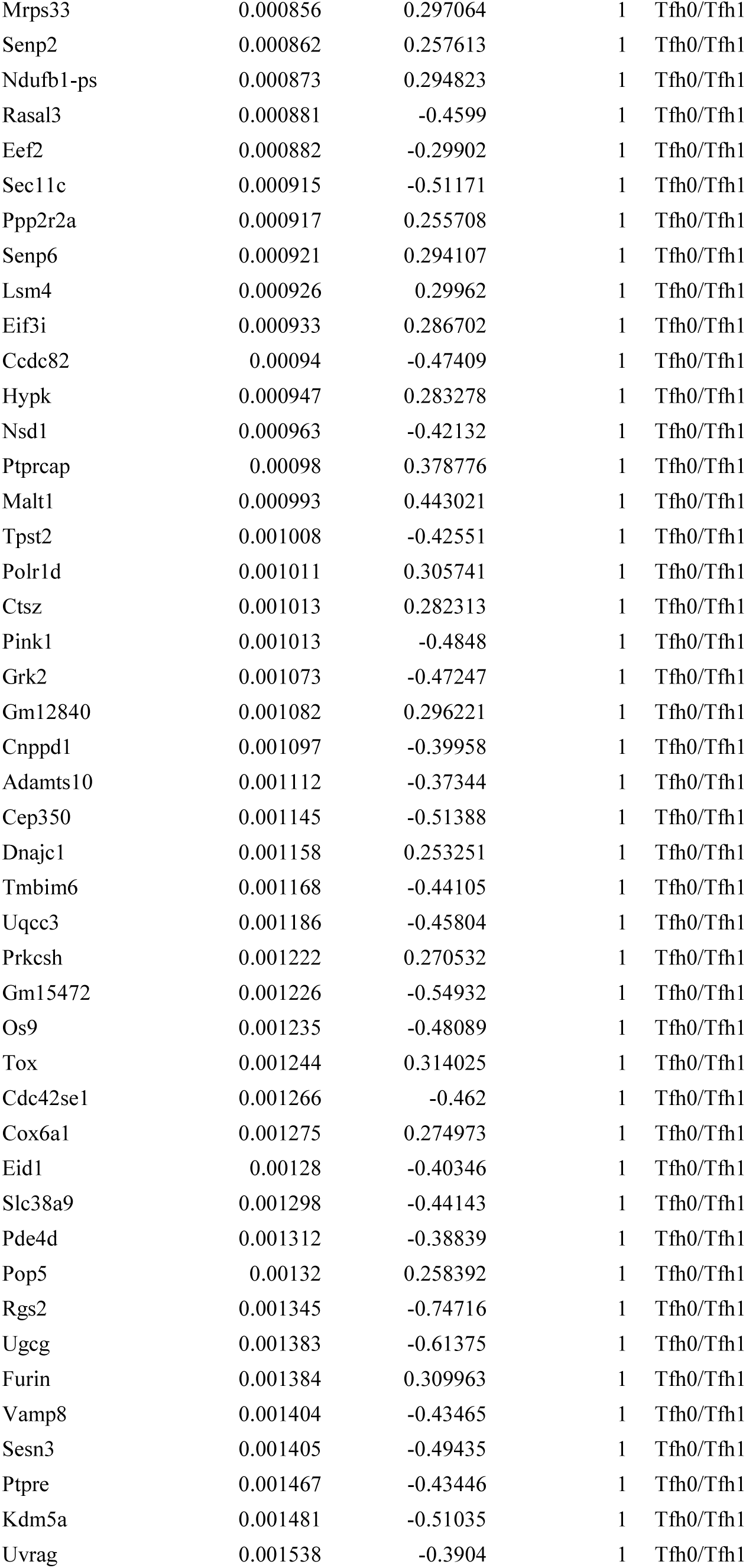
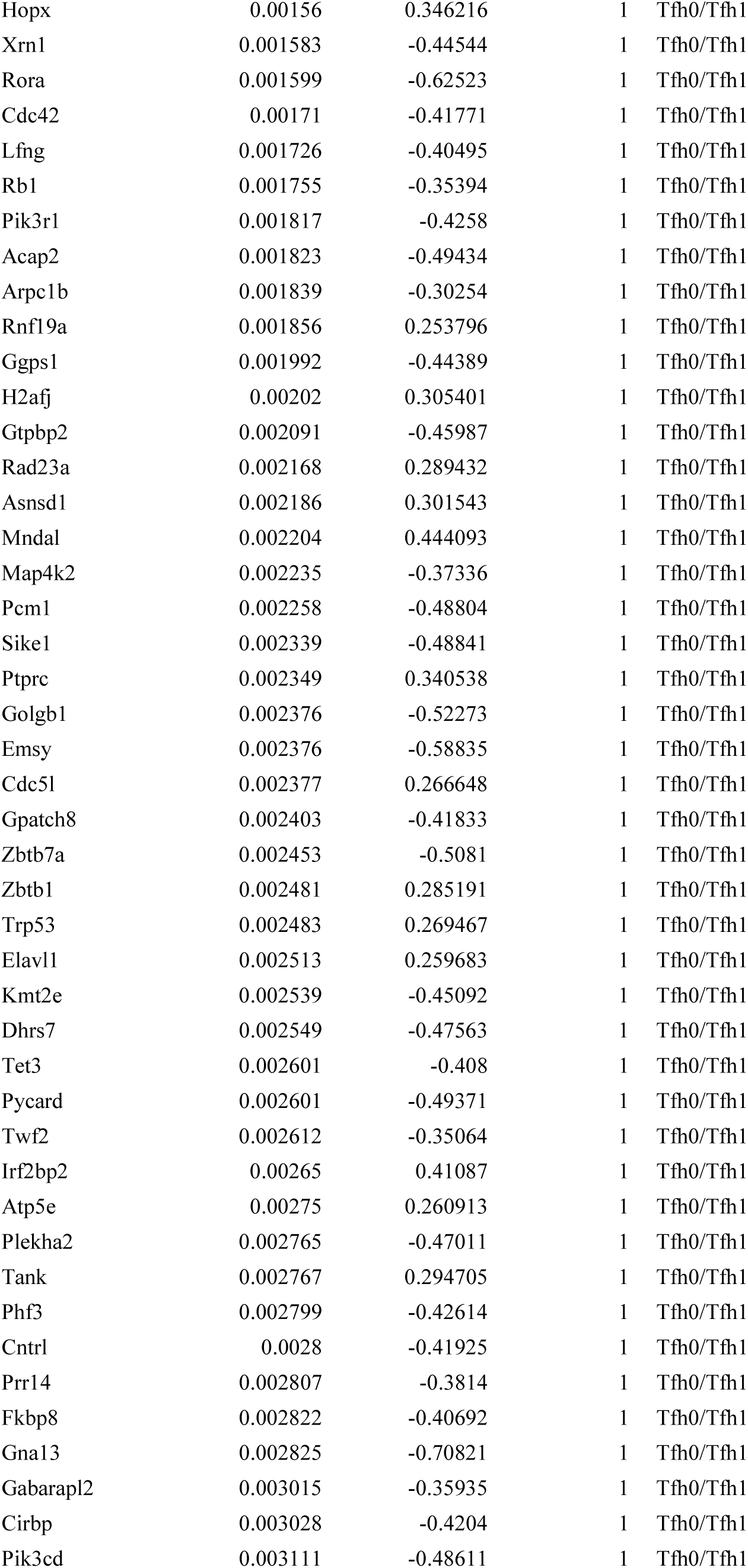
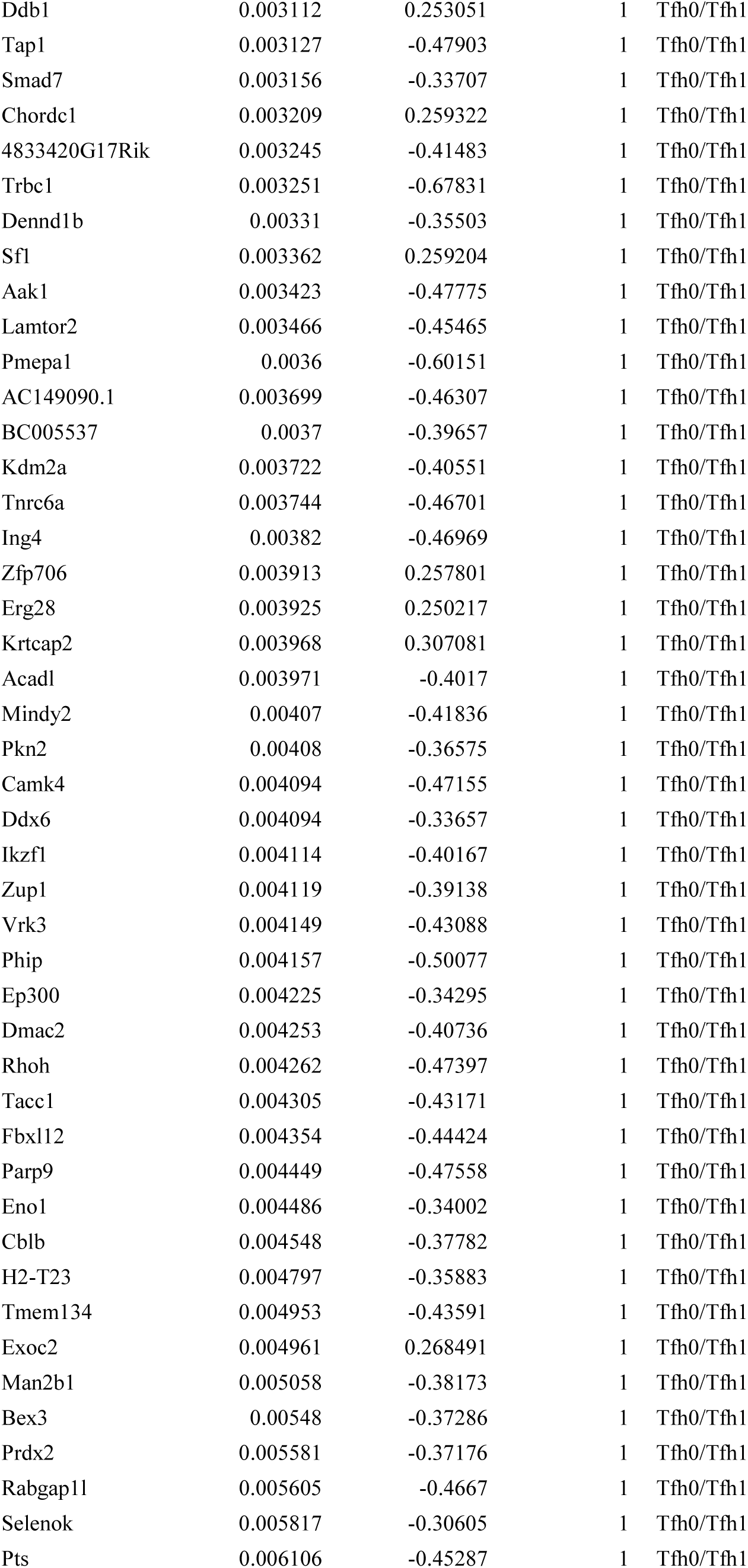
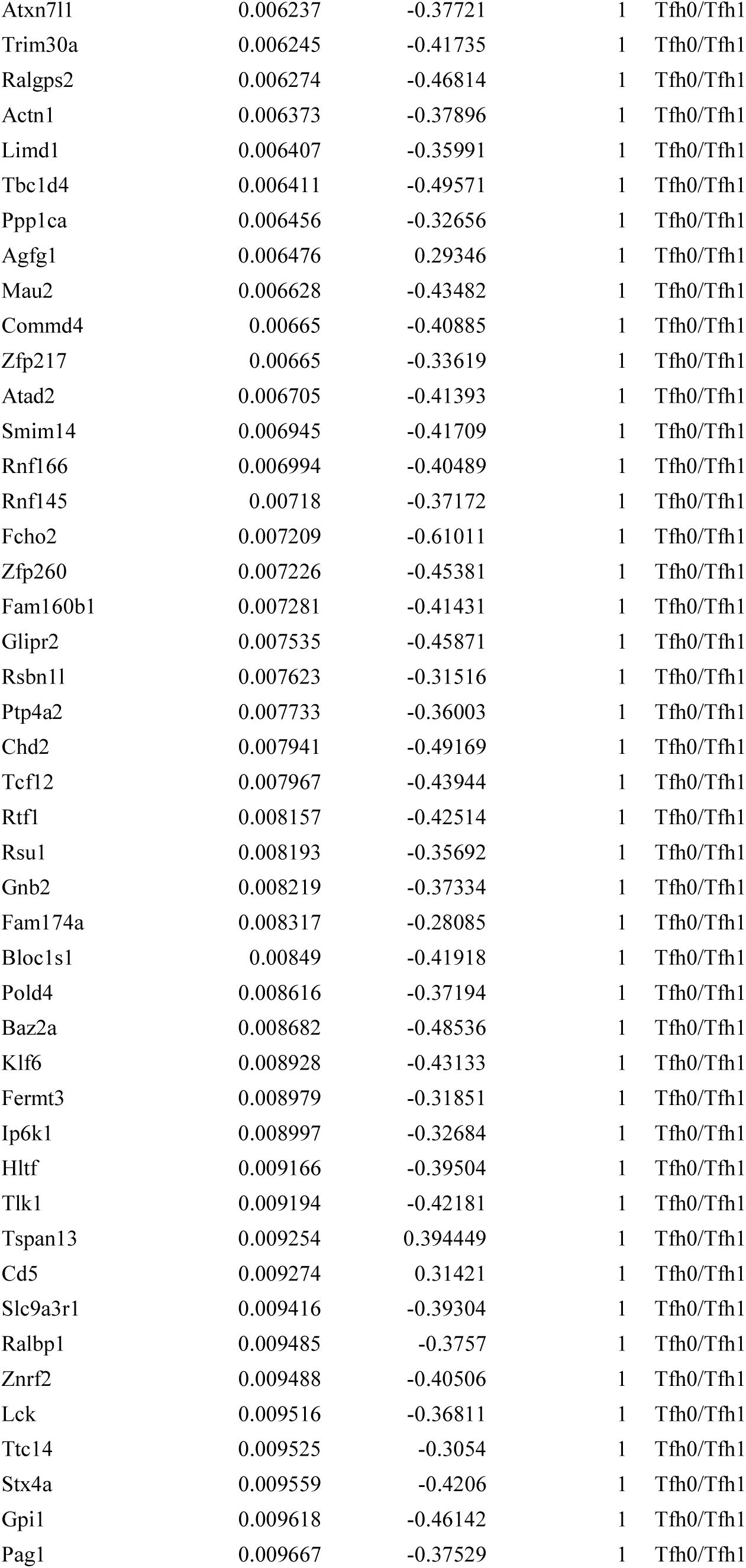
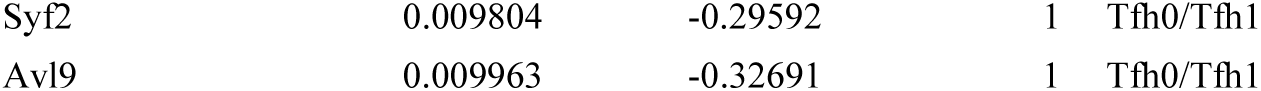

**Supplemental Table 2.**
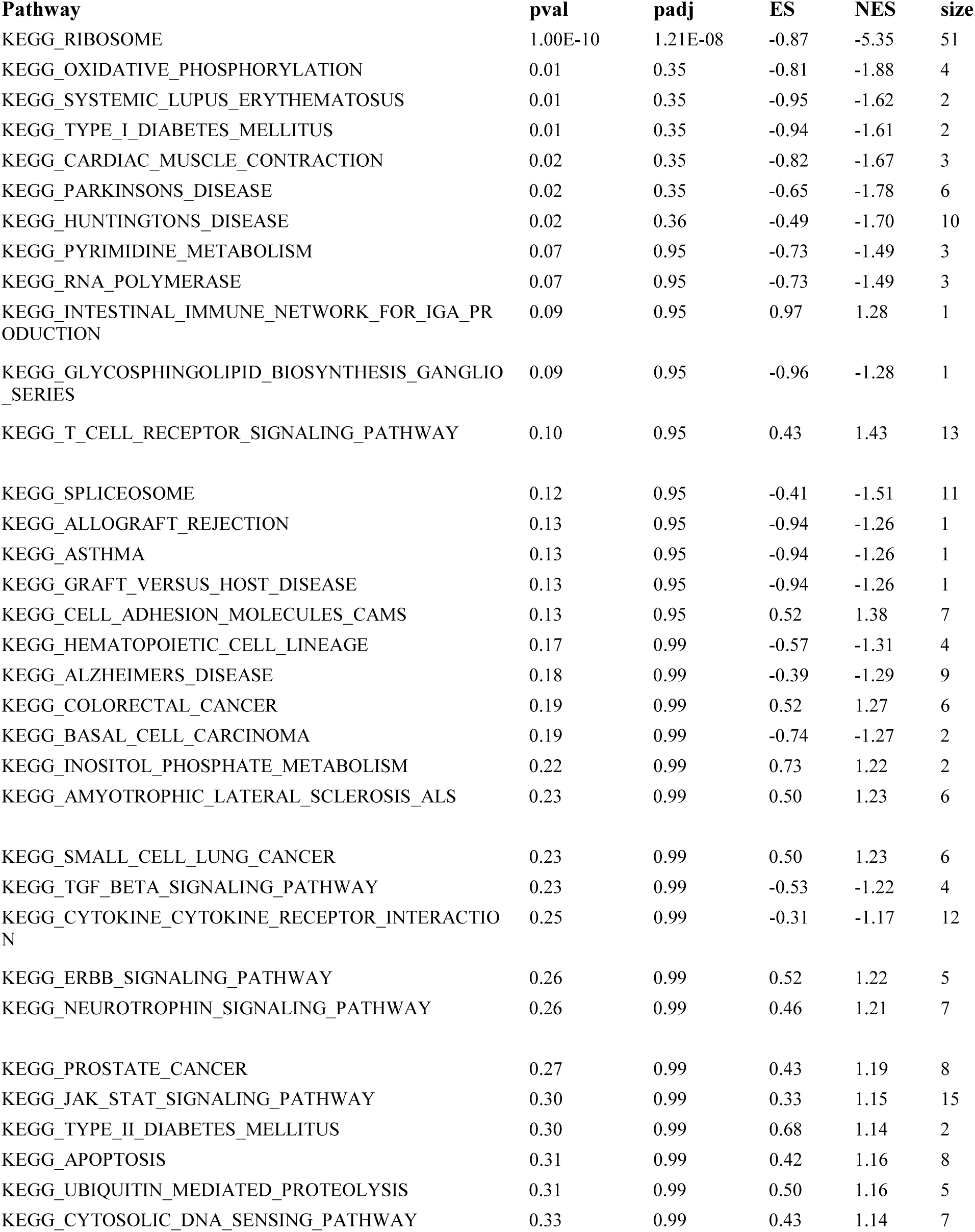
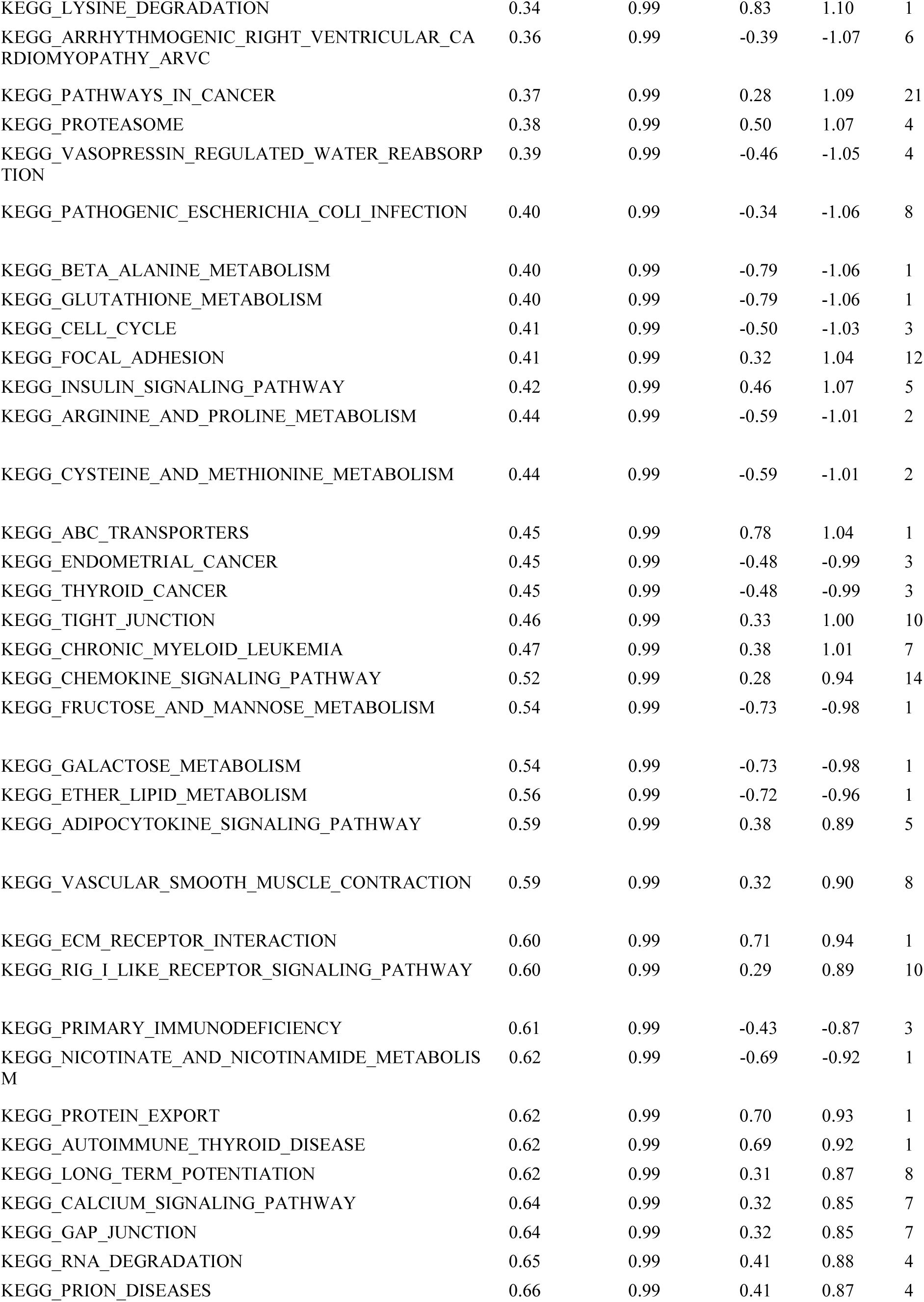
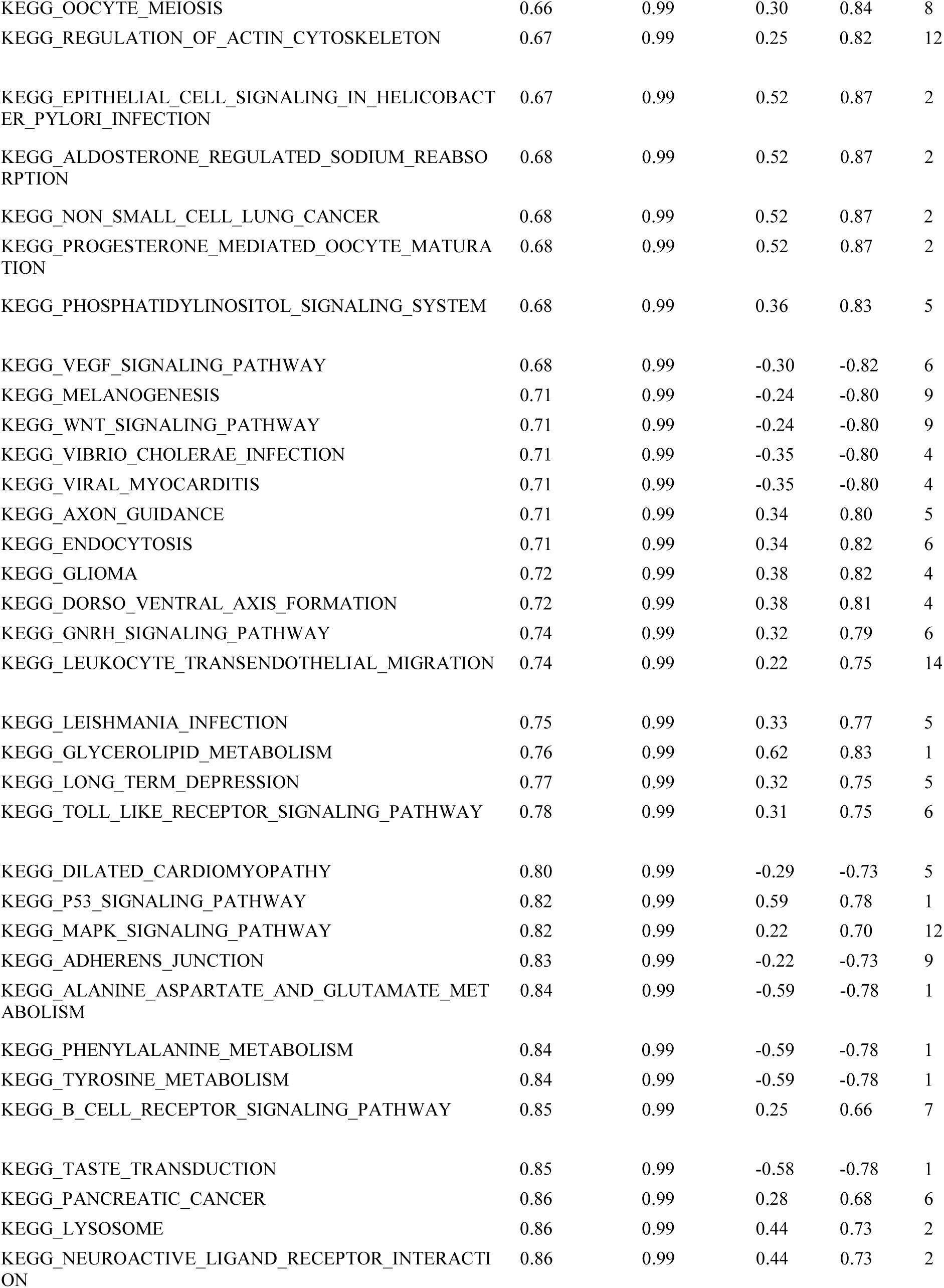
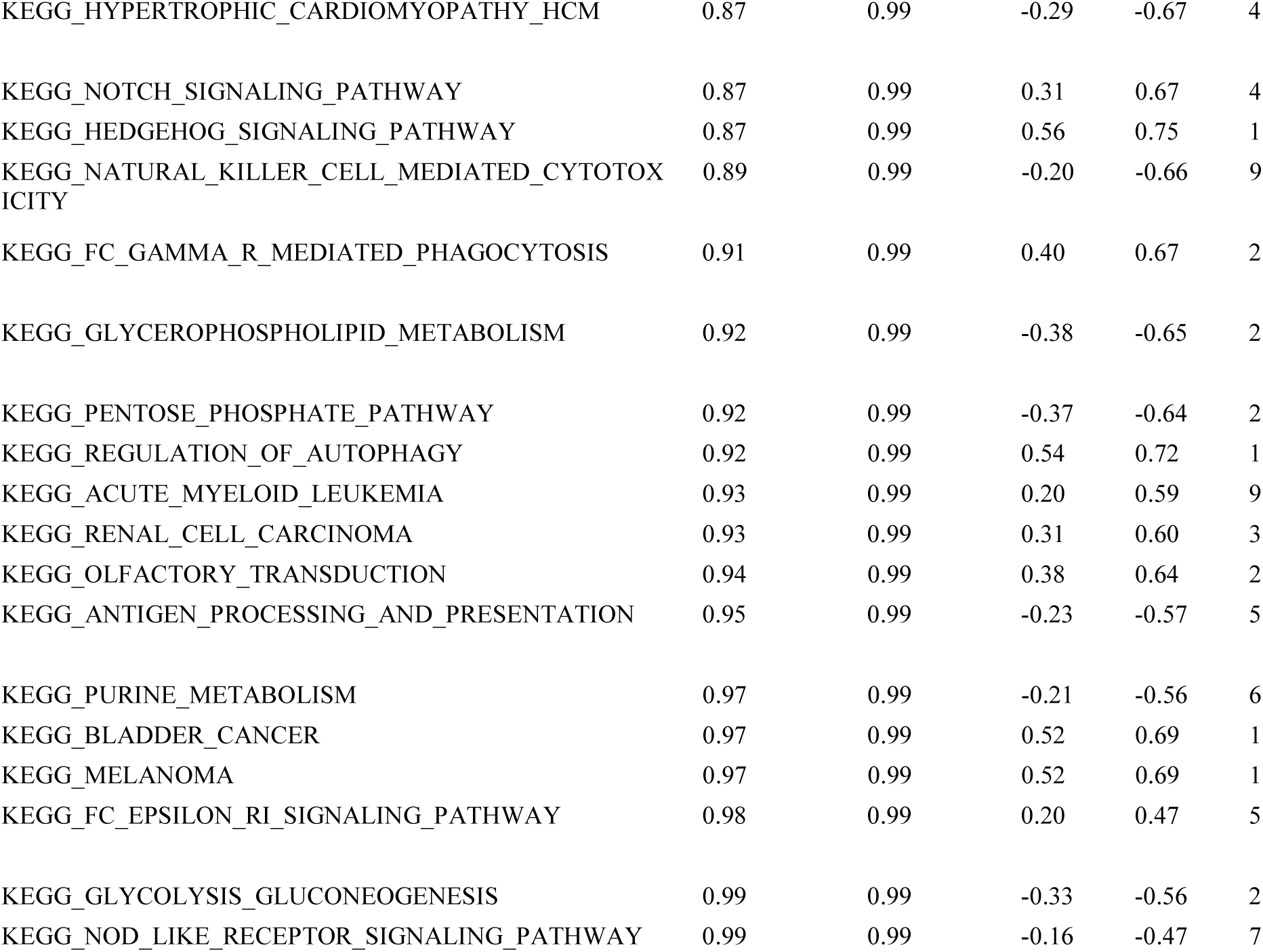

**Supplemental Table 3.**
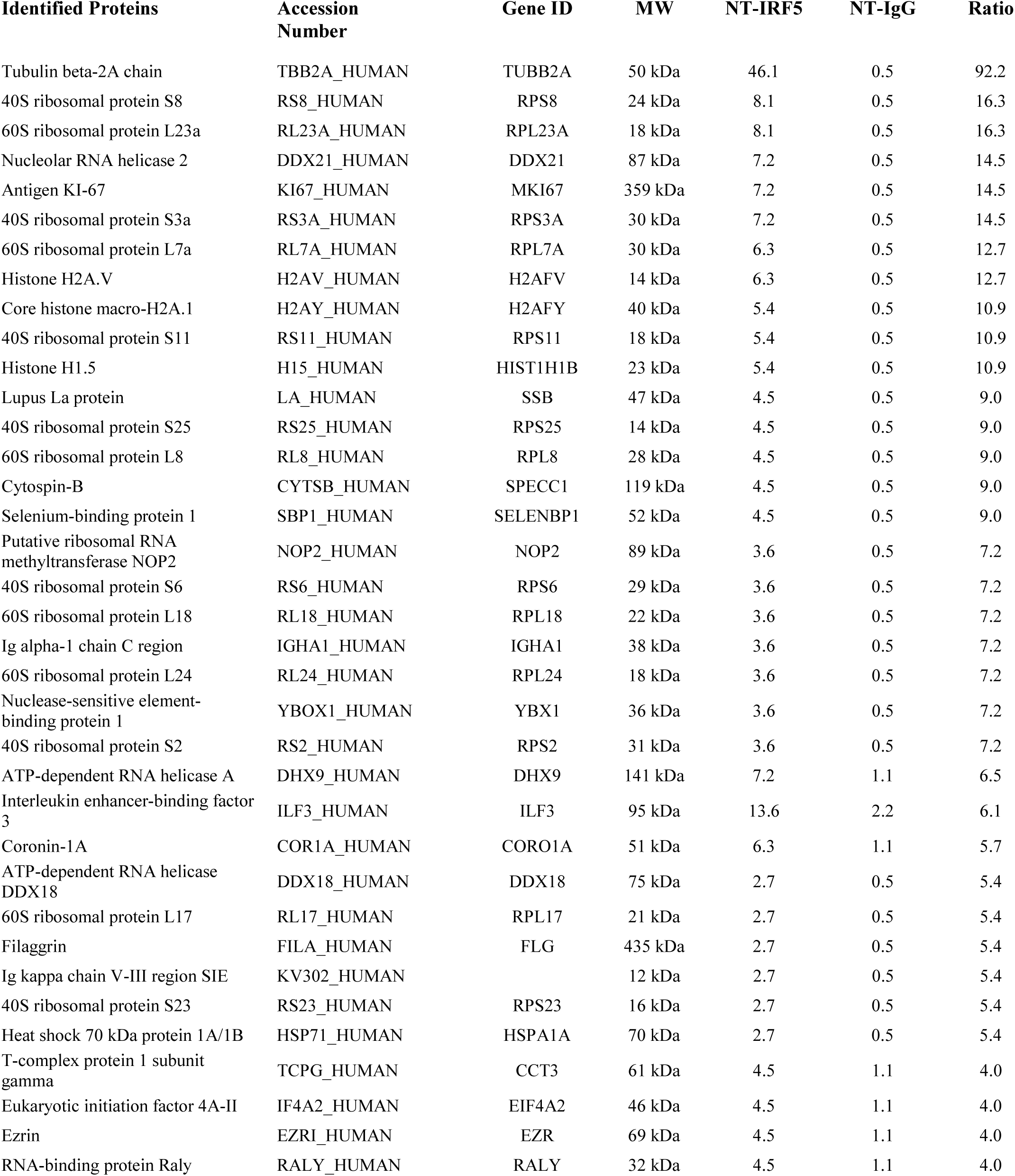
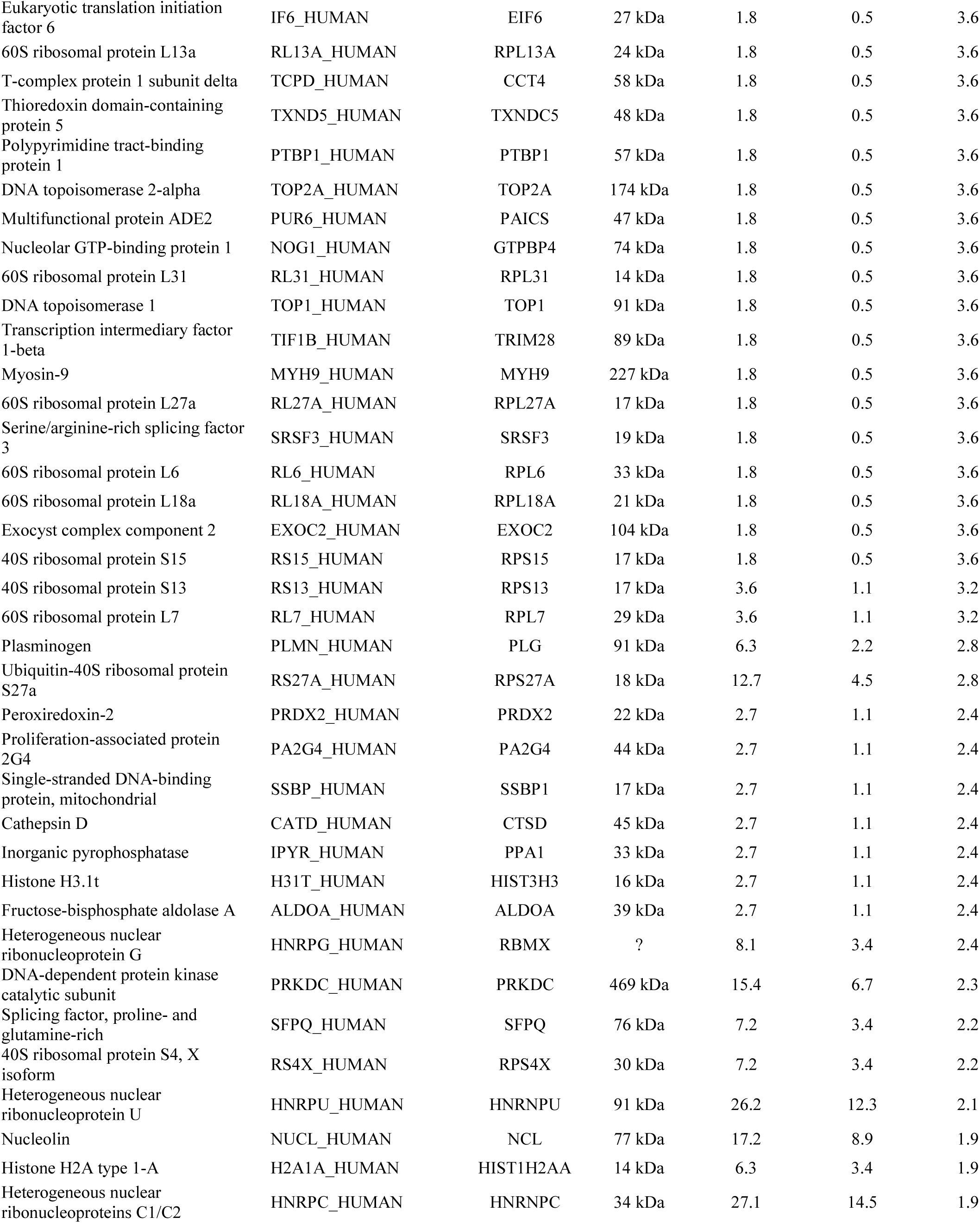
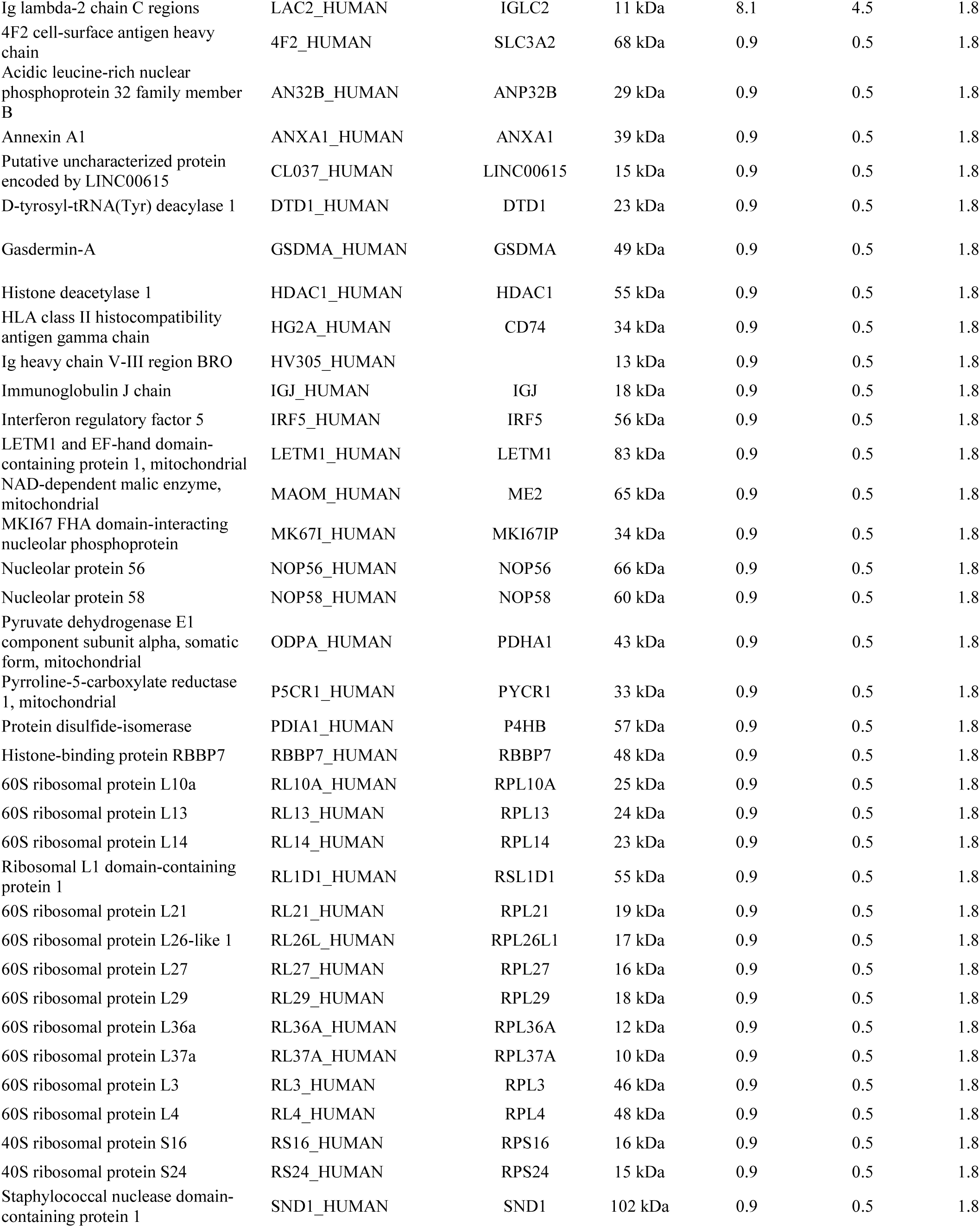
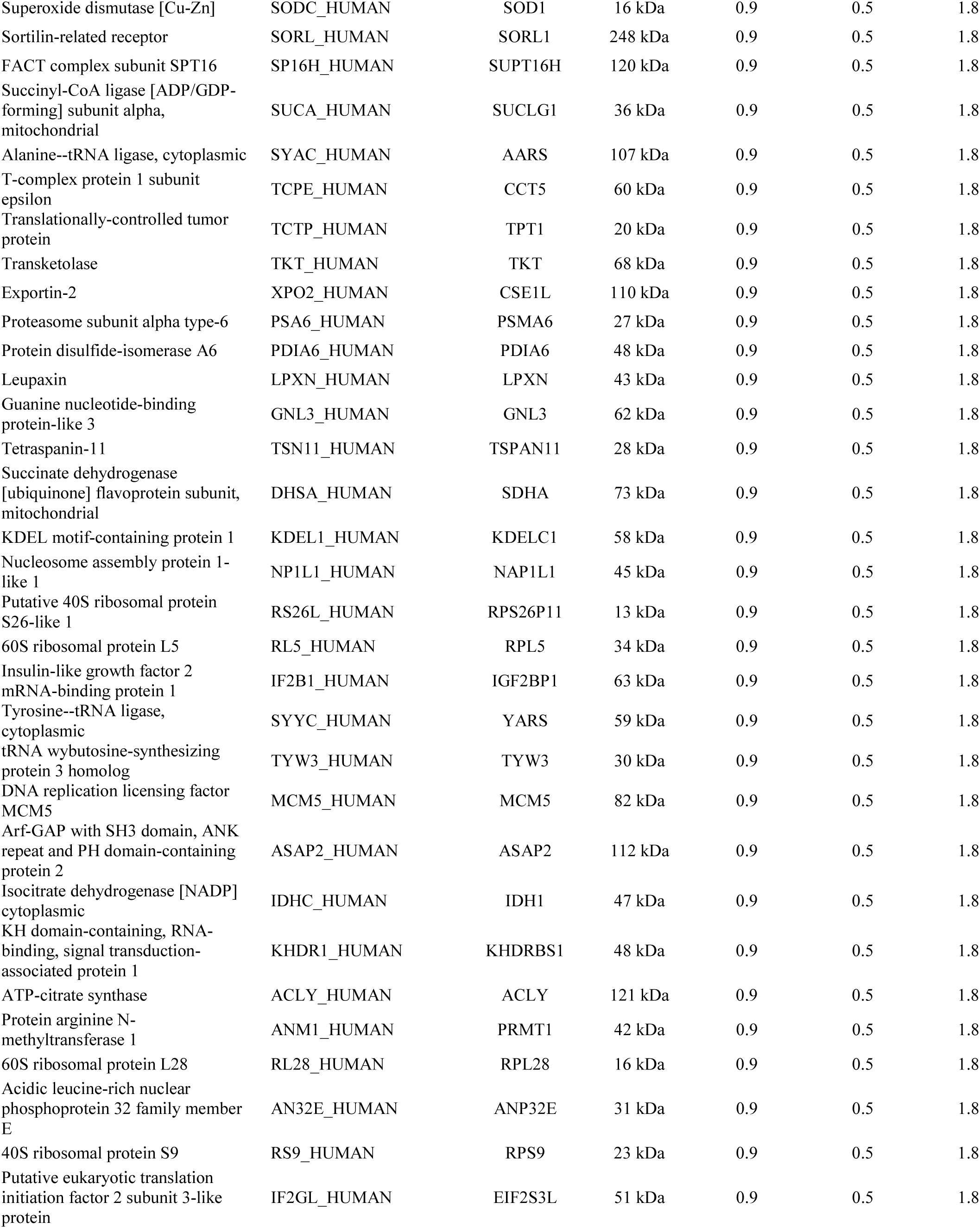
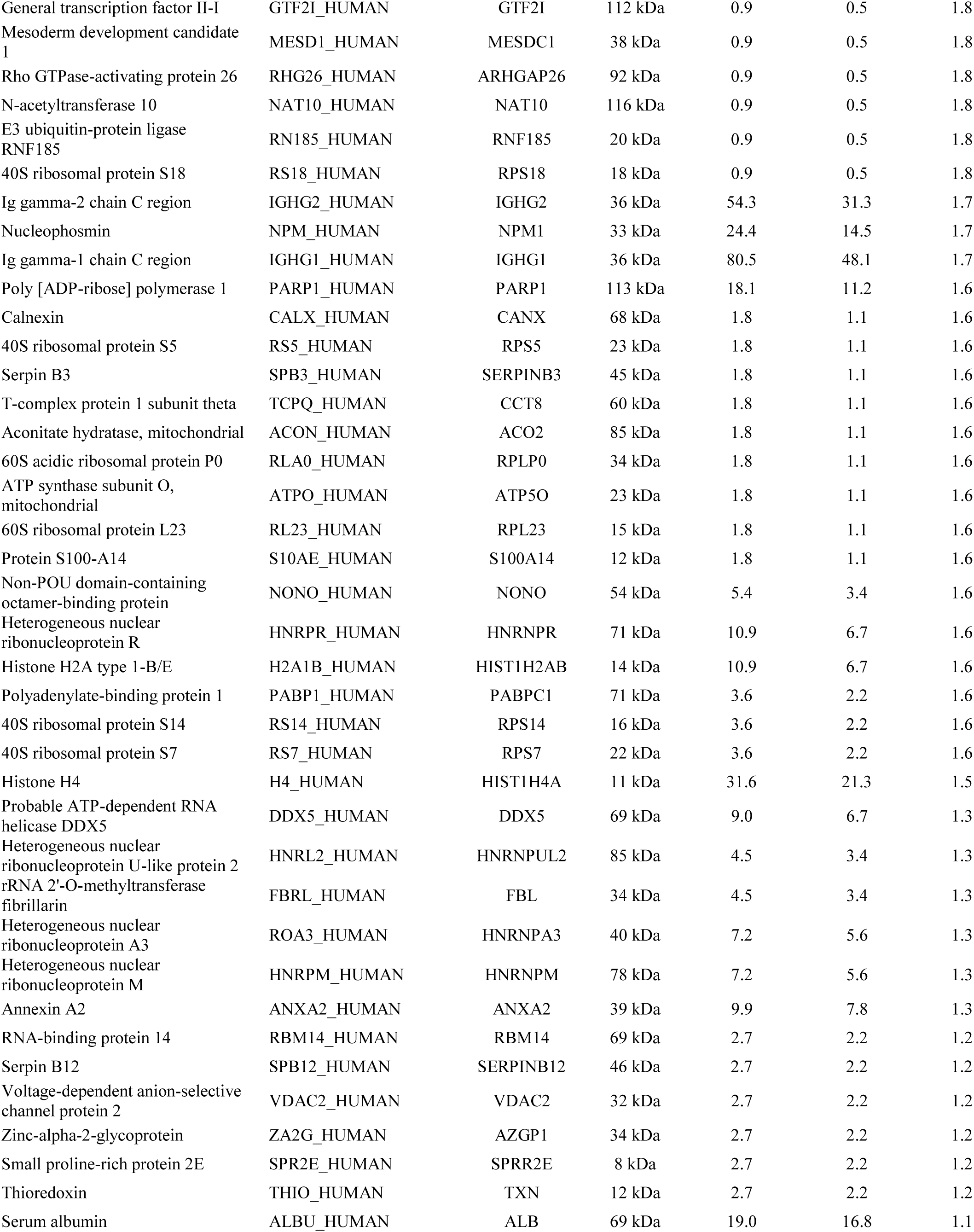
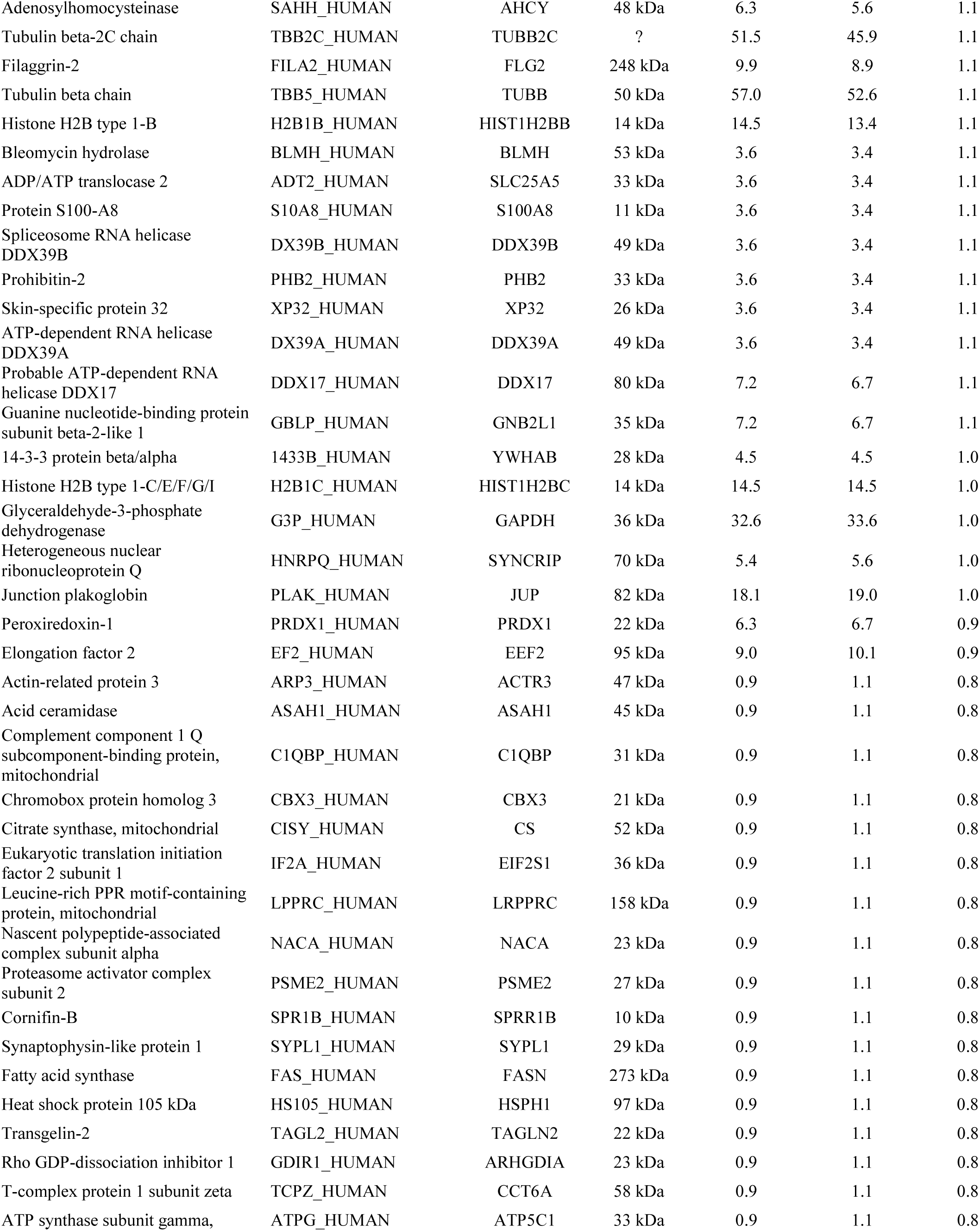
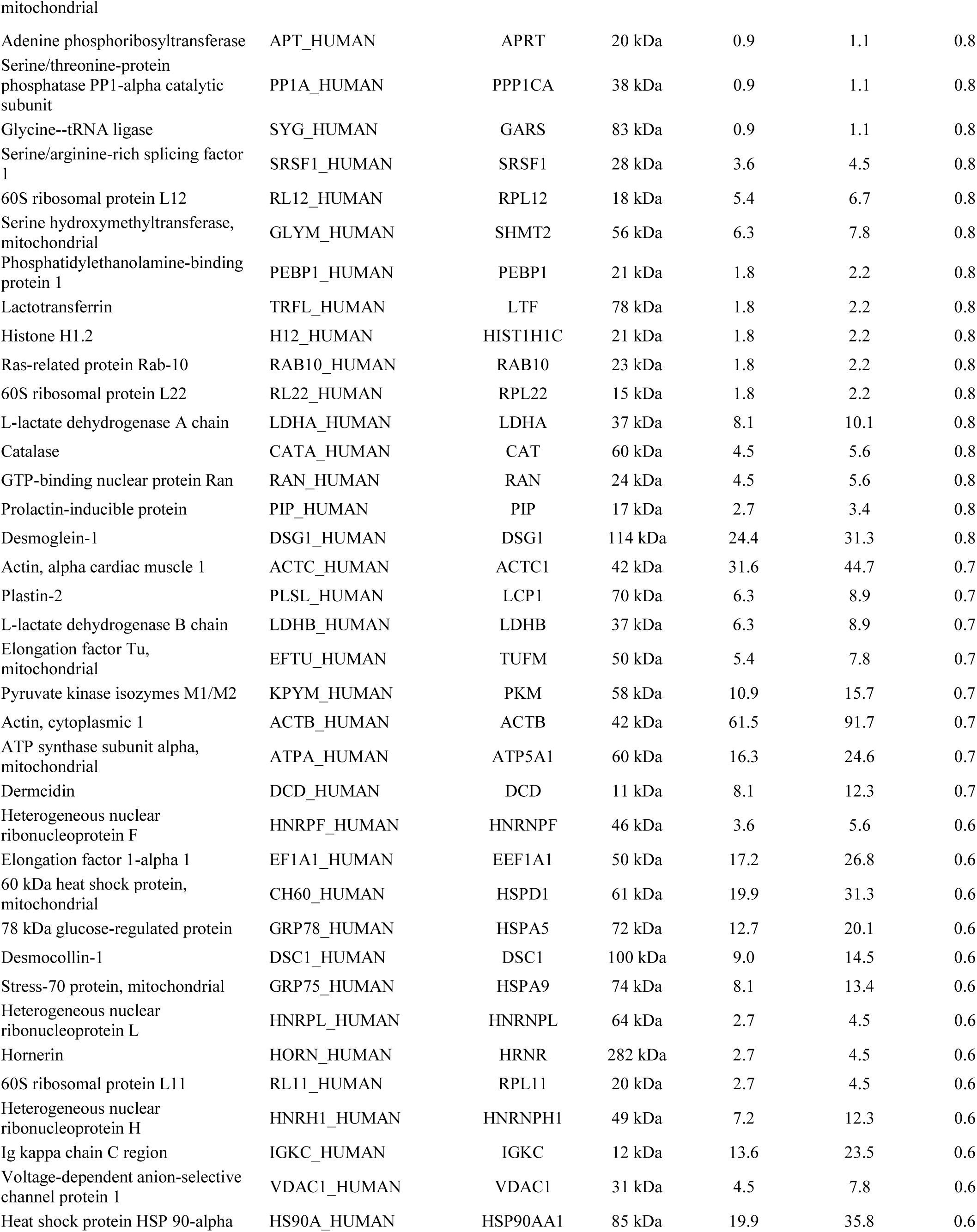
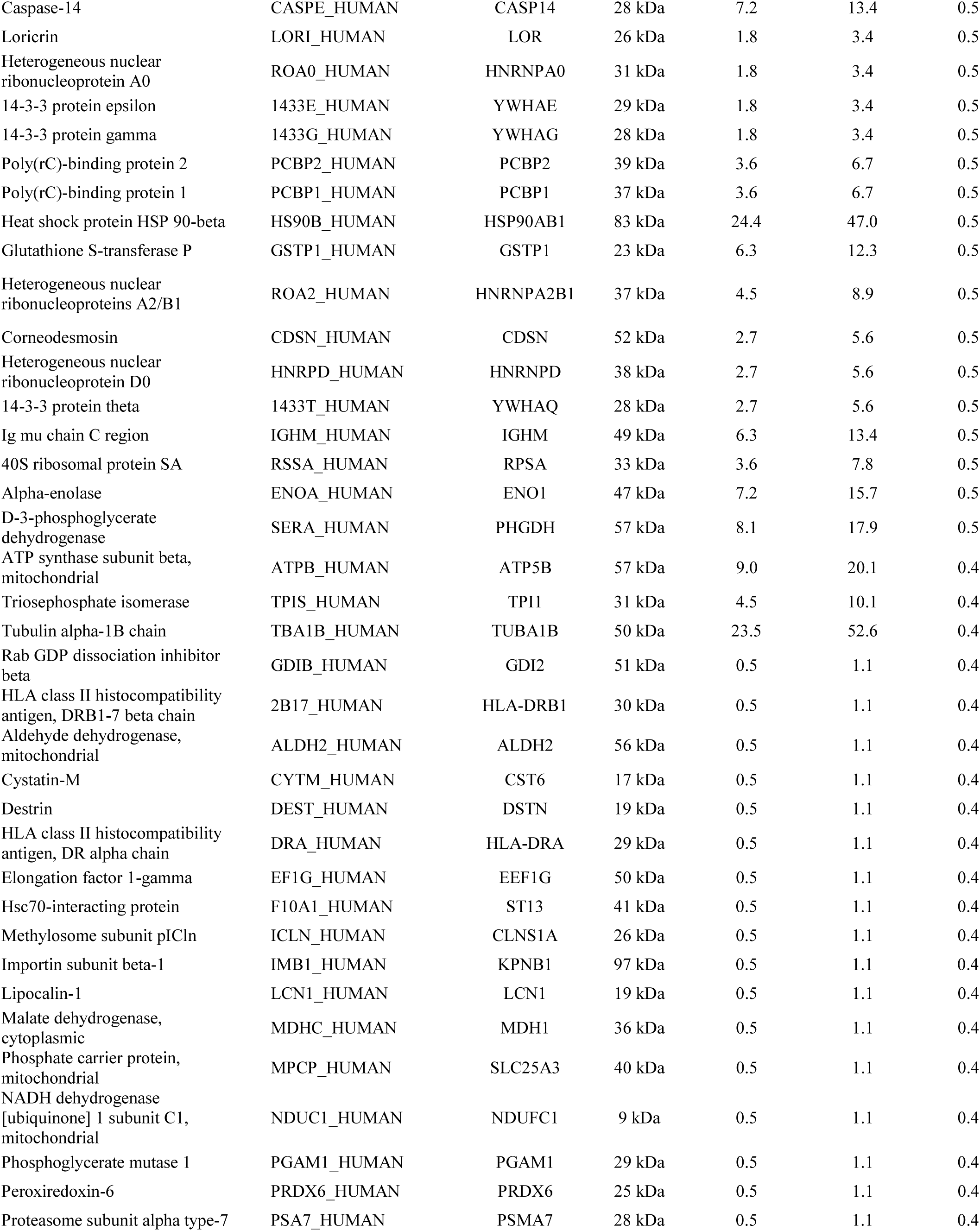
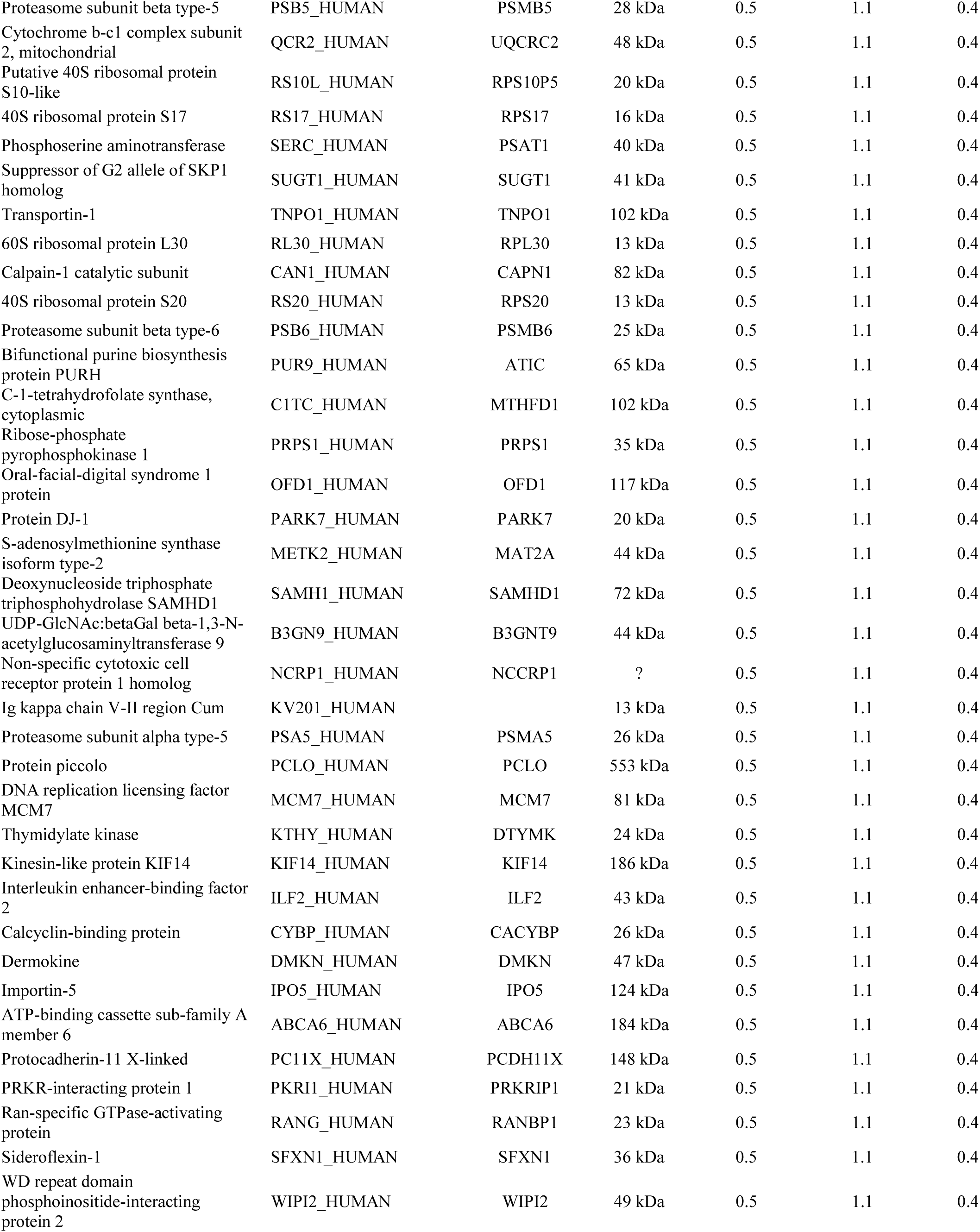
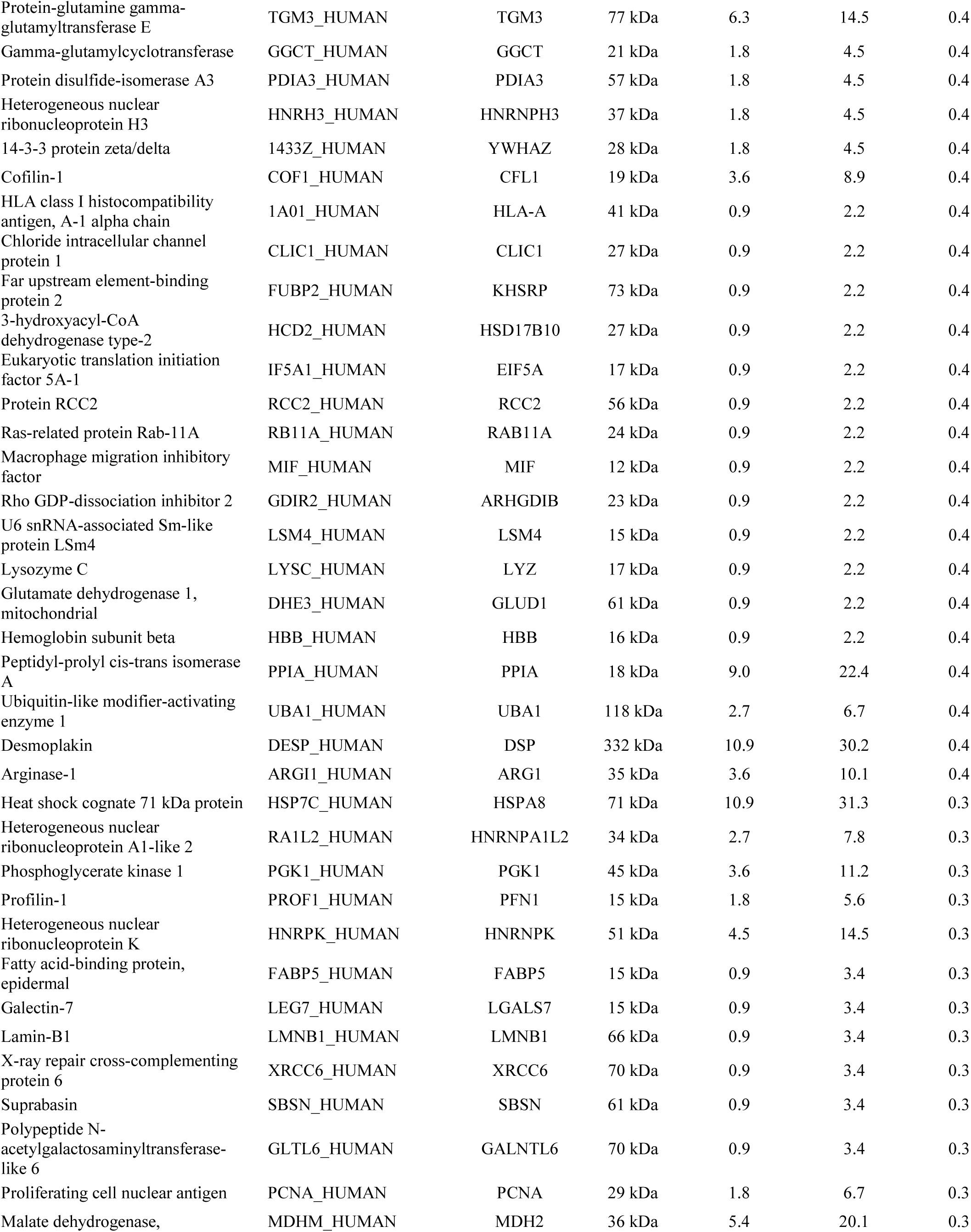
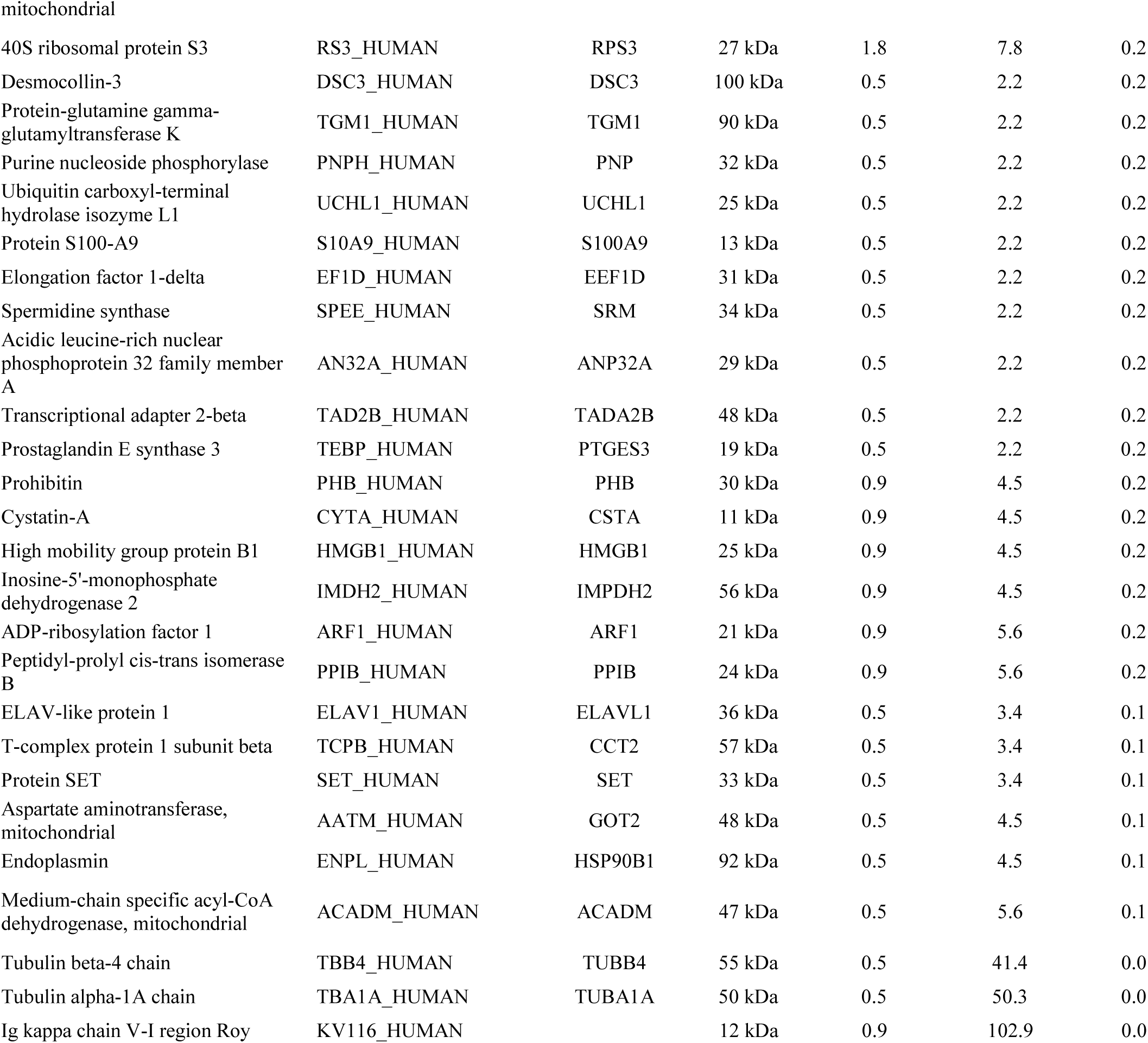

**Supplemental Table 4.**
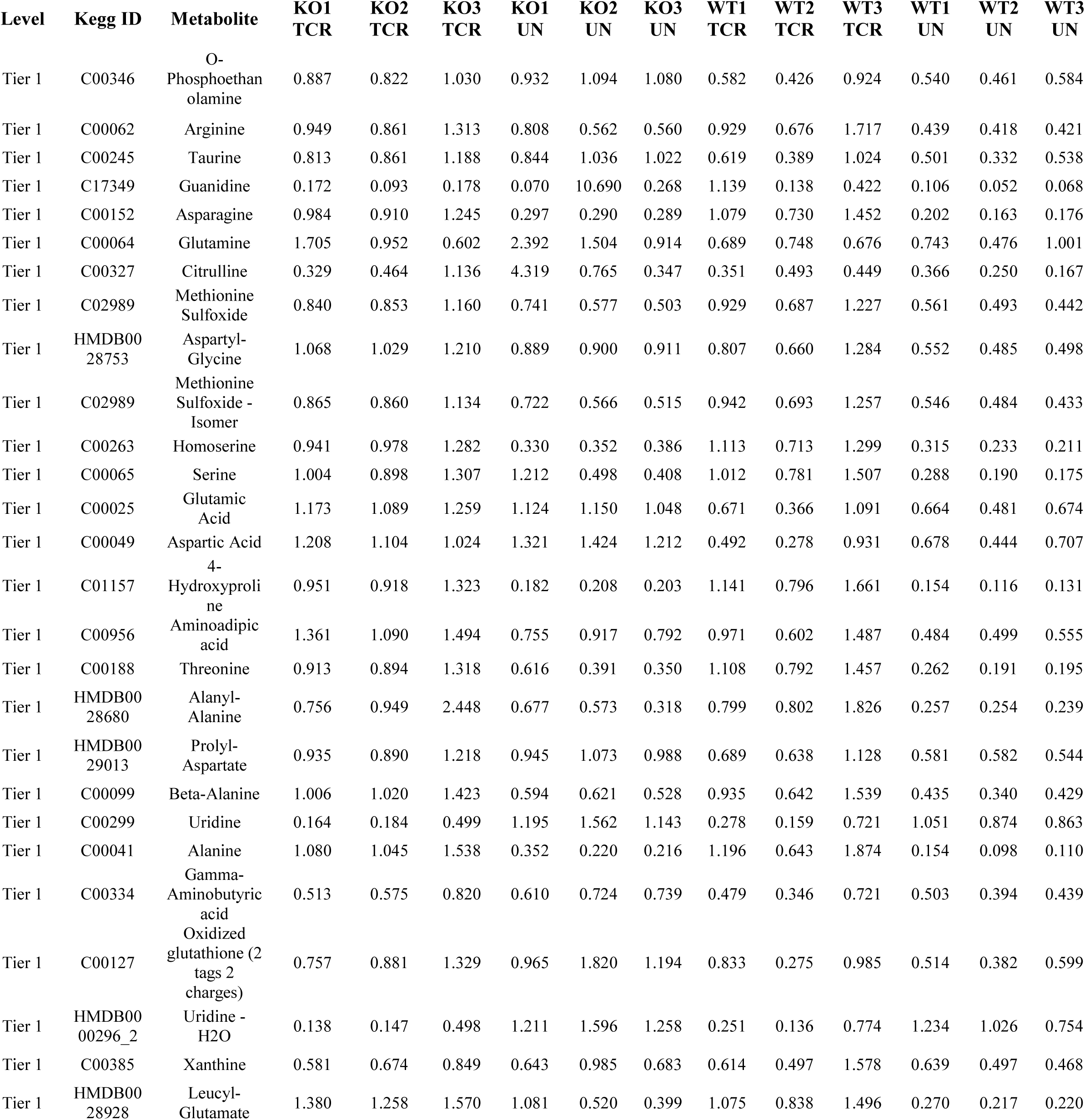
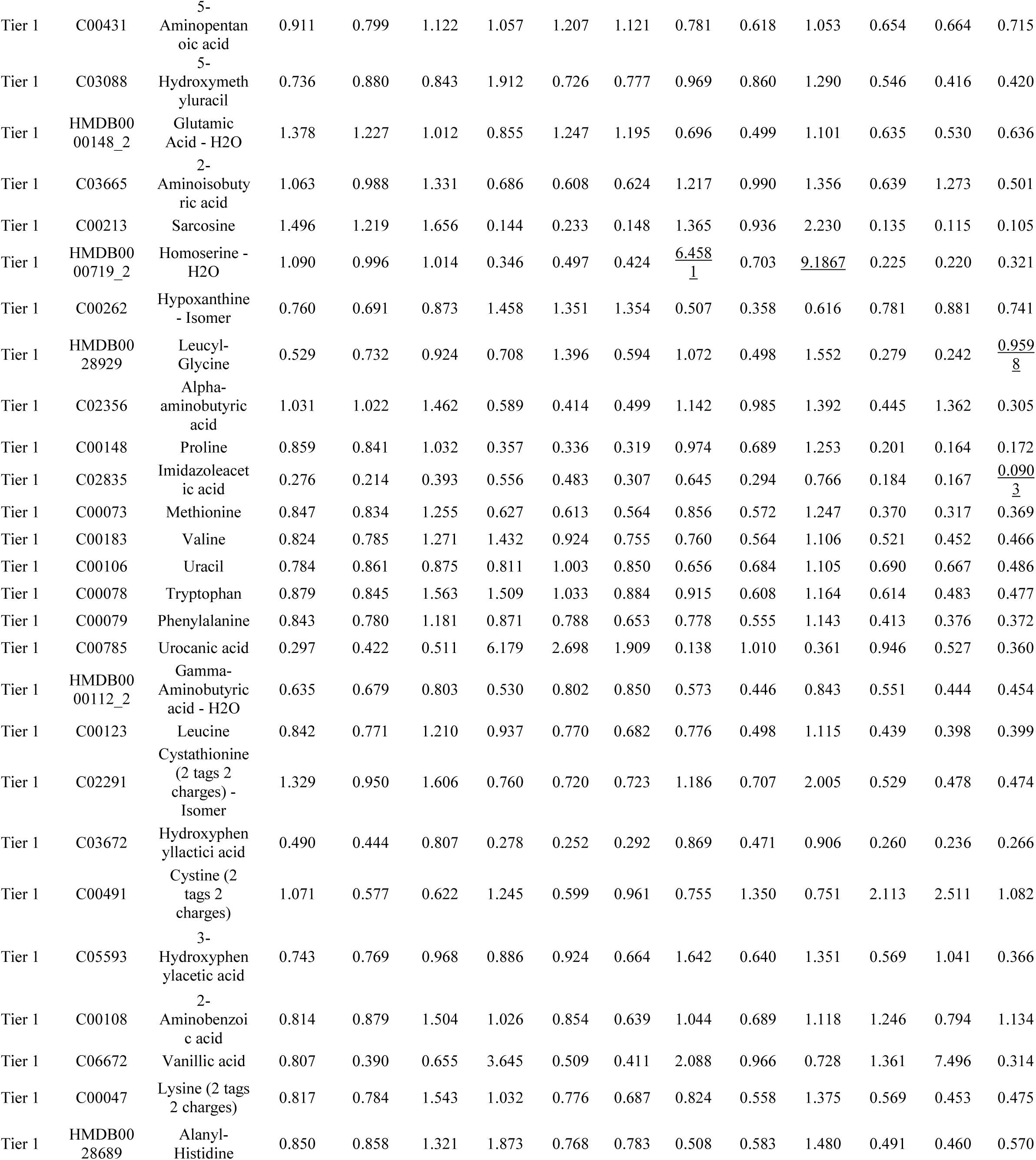
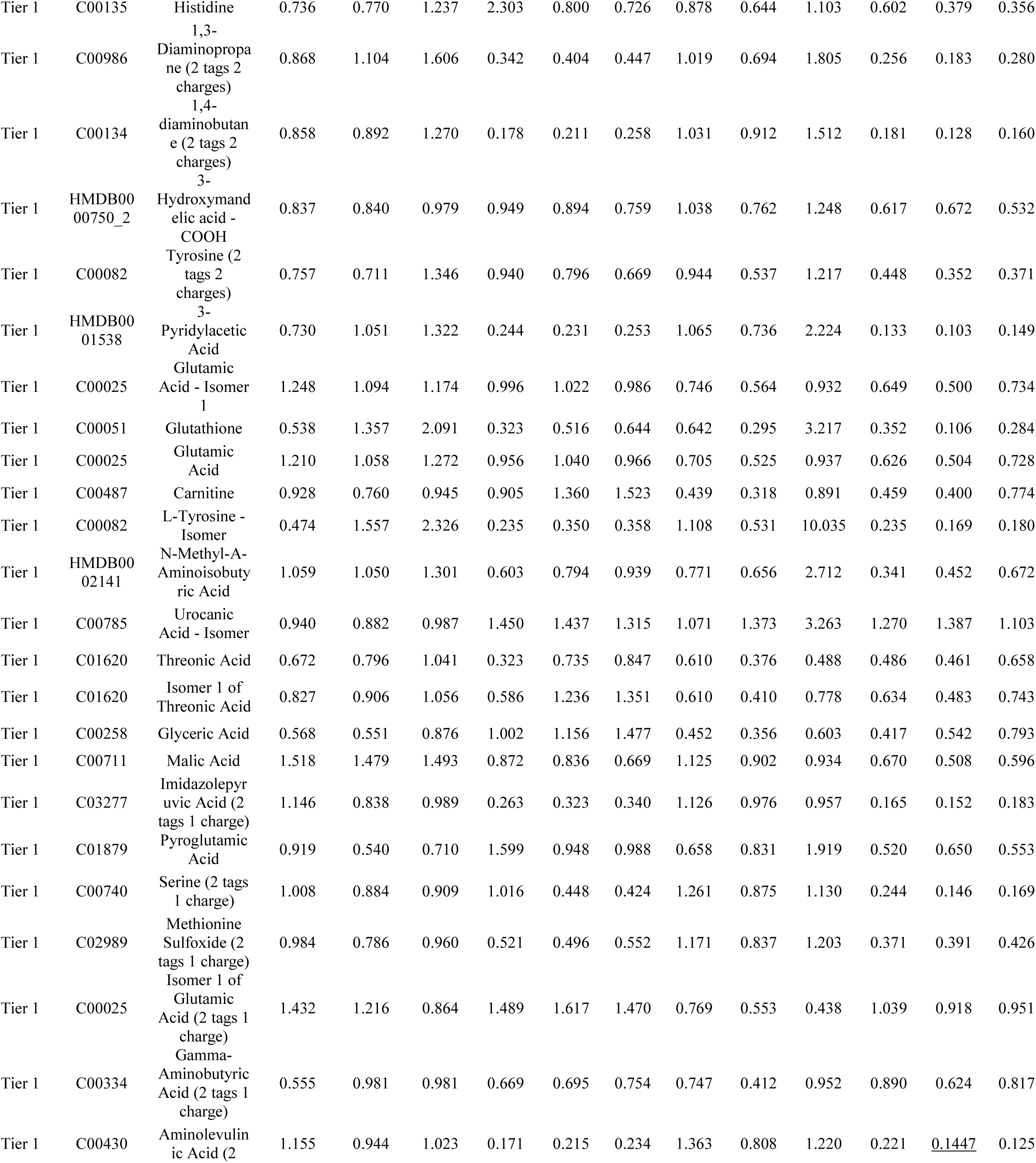
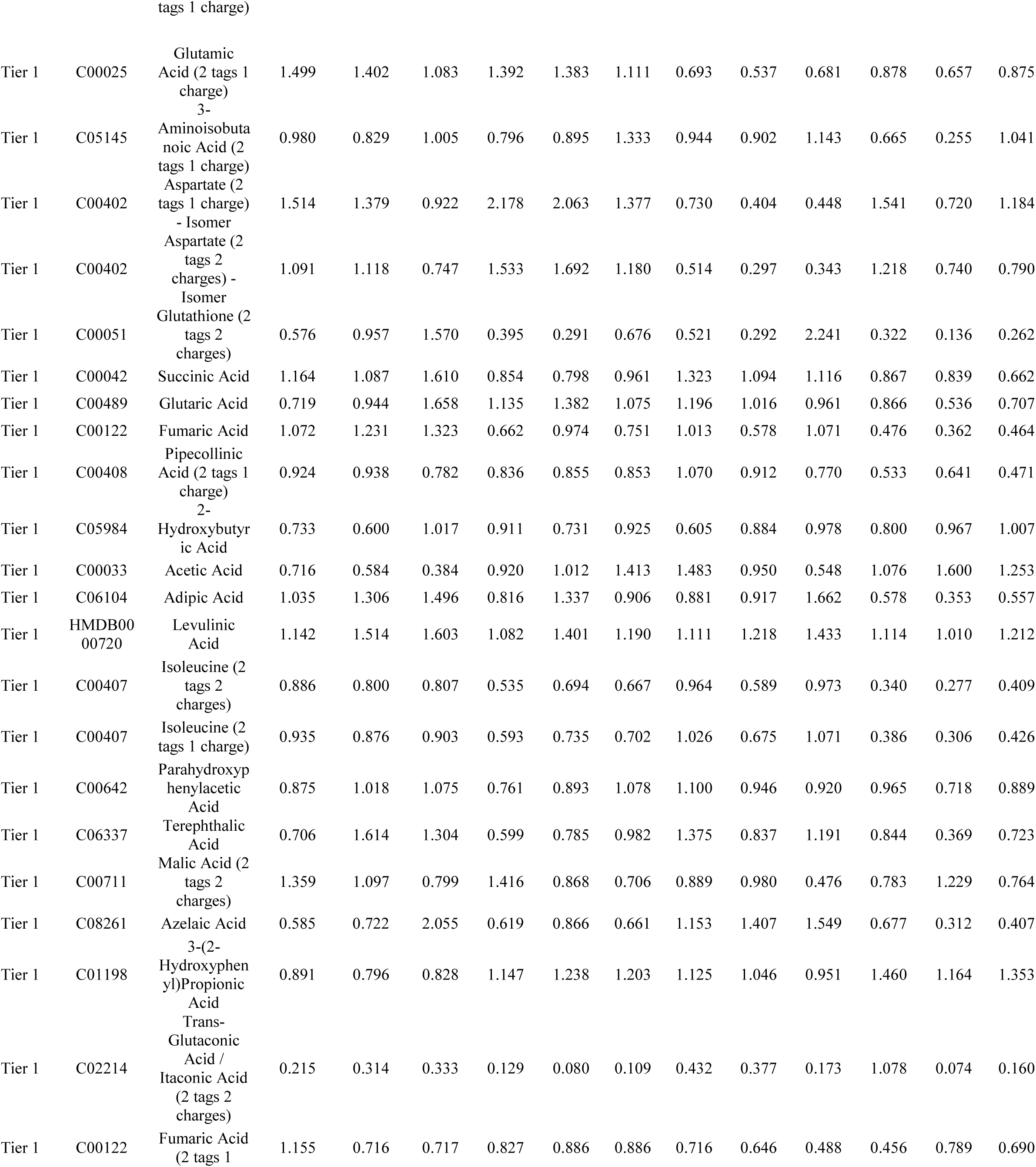
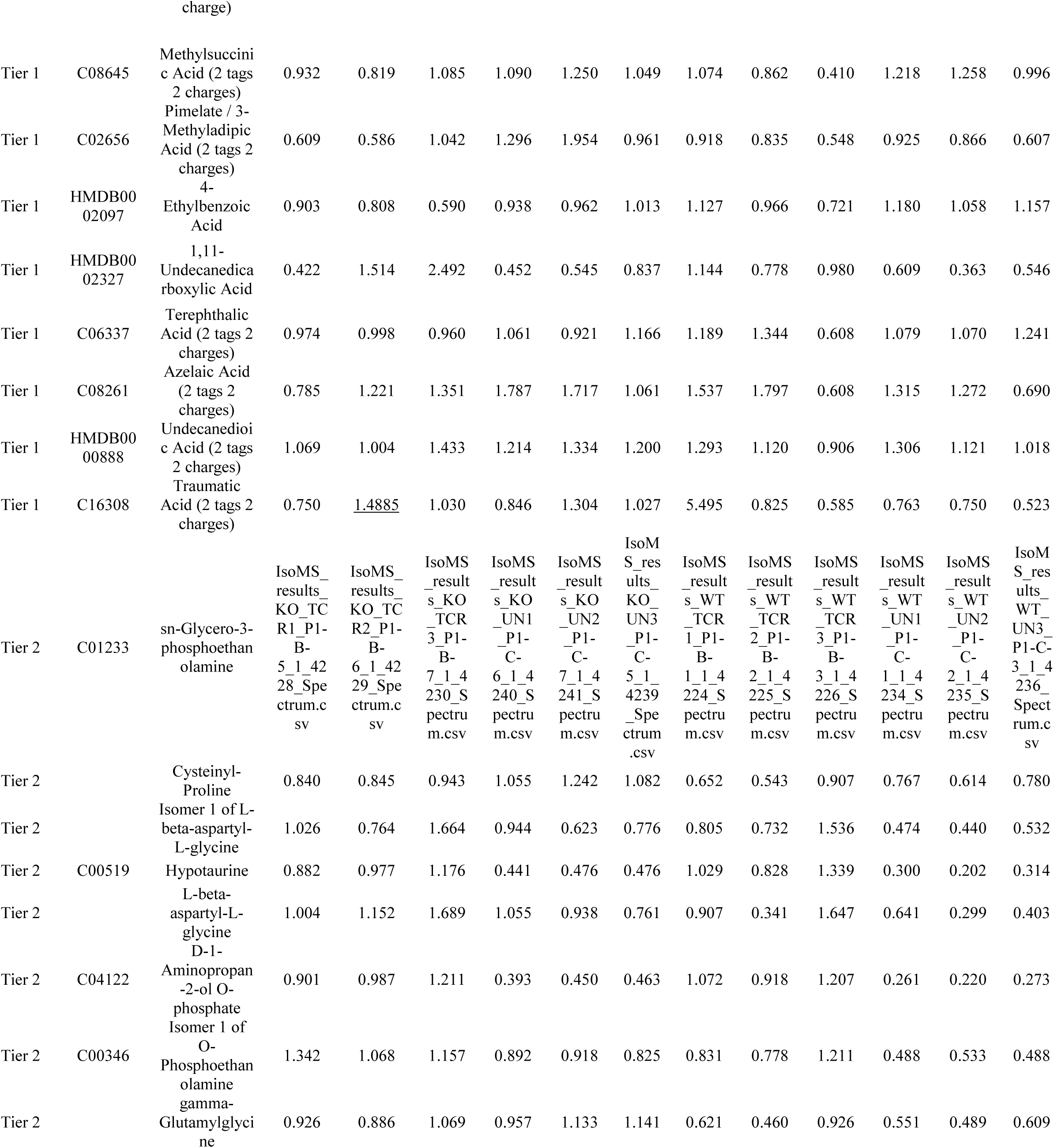
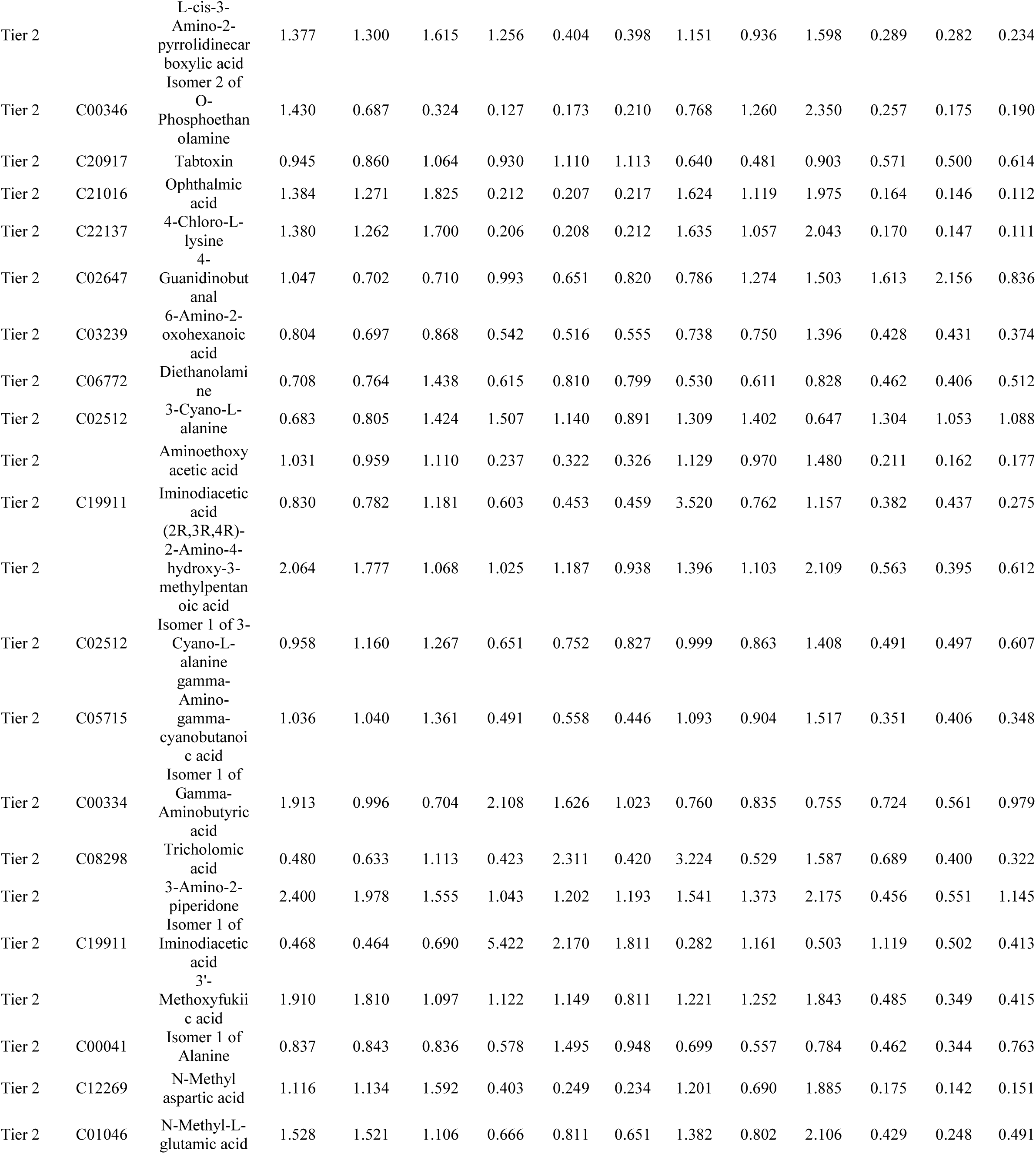
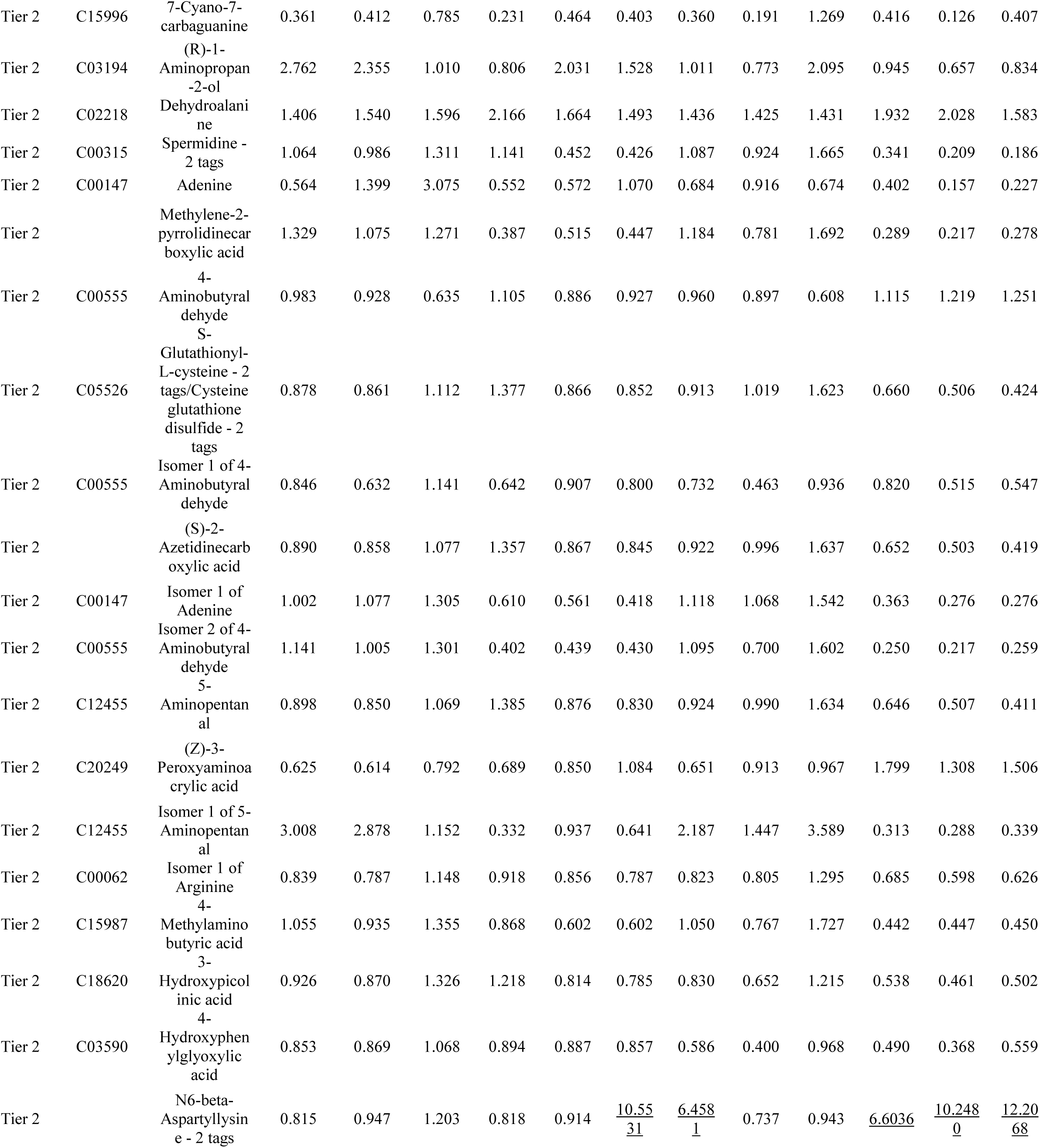
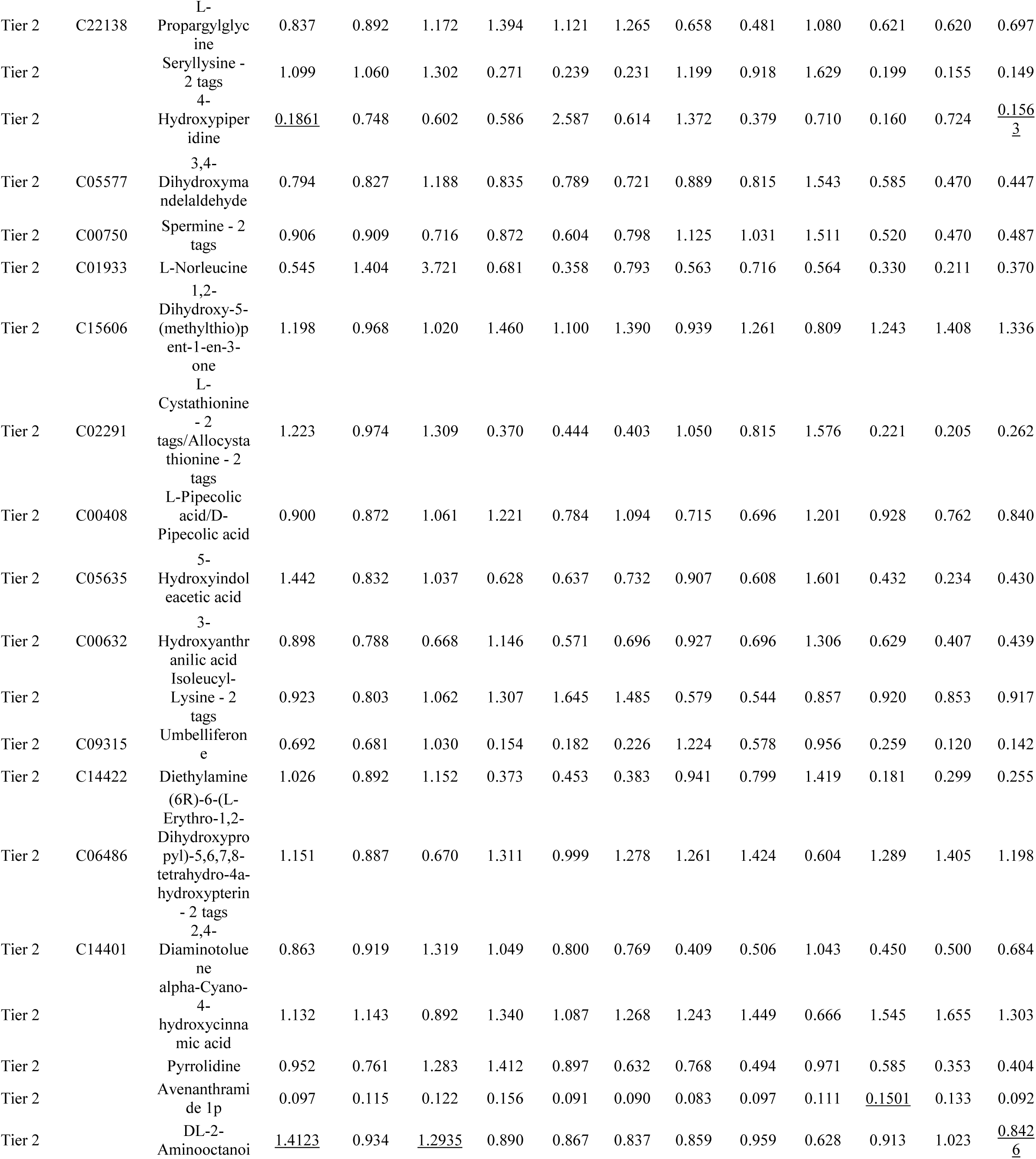
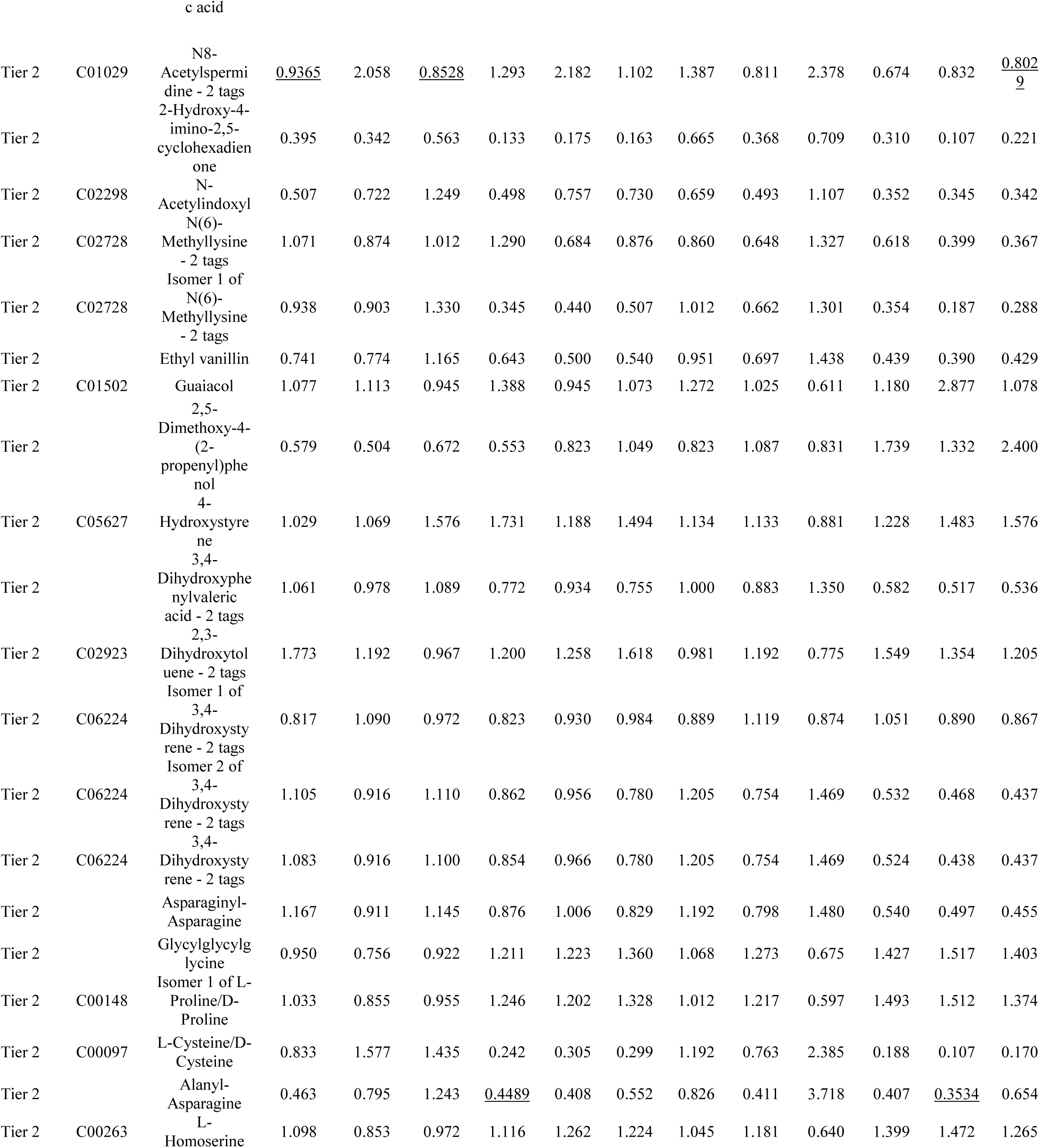
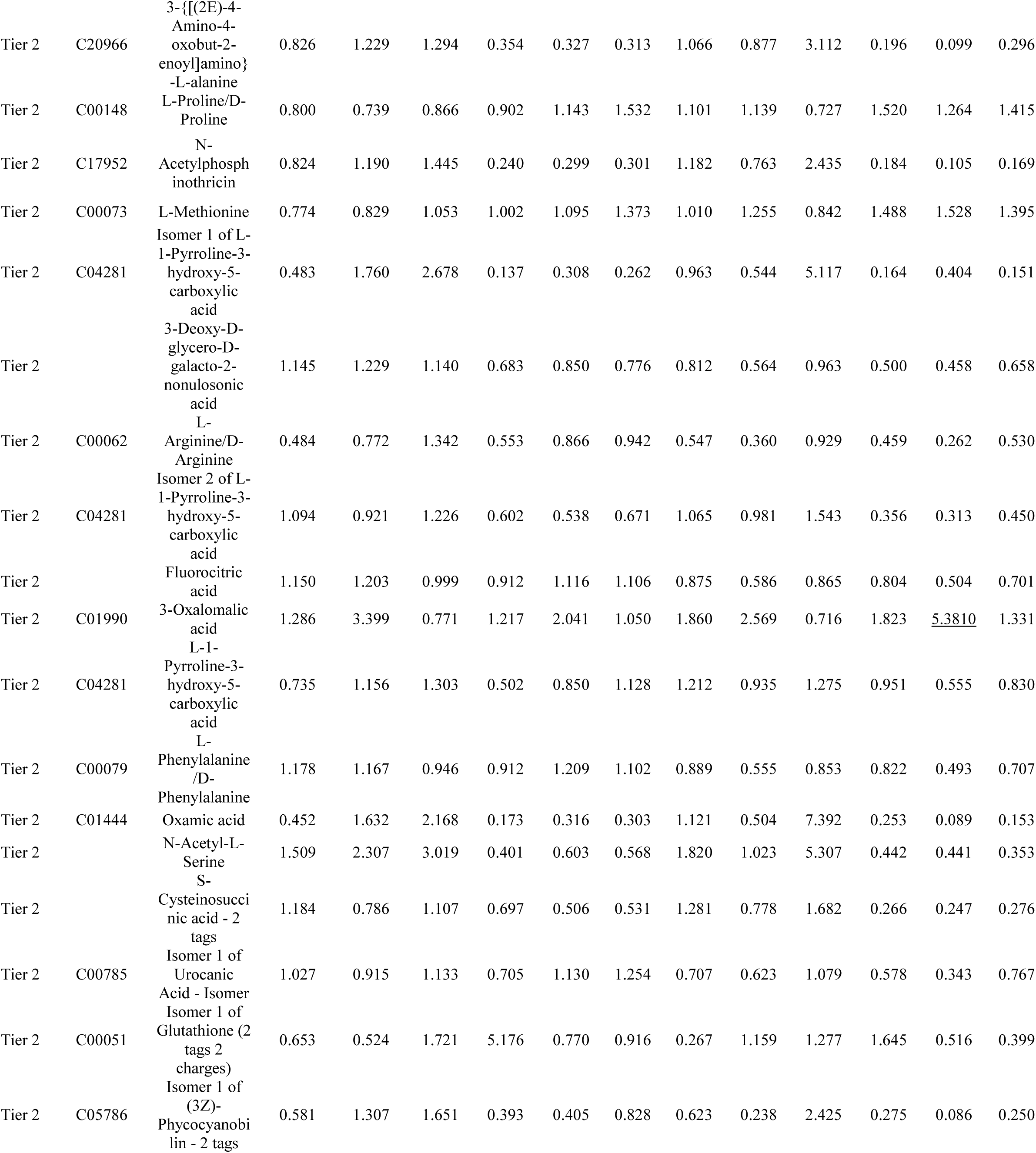
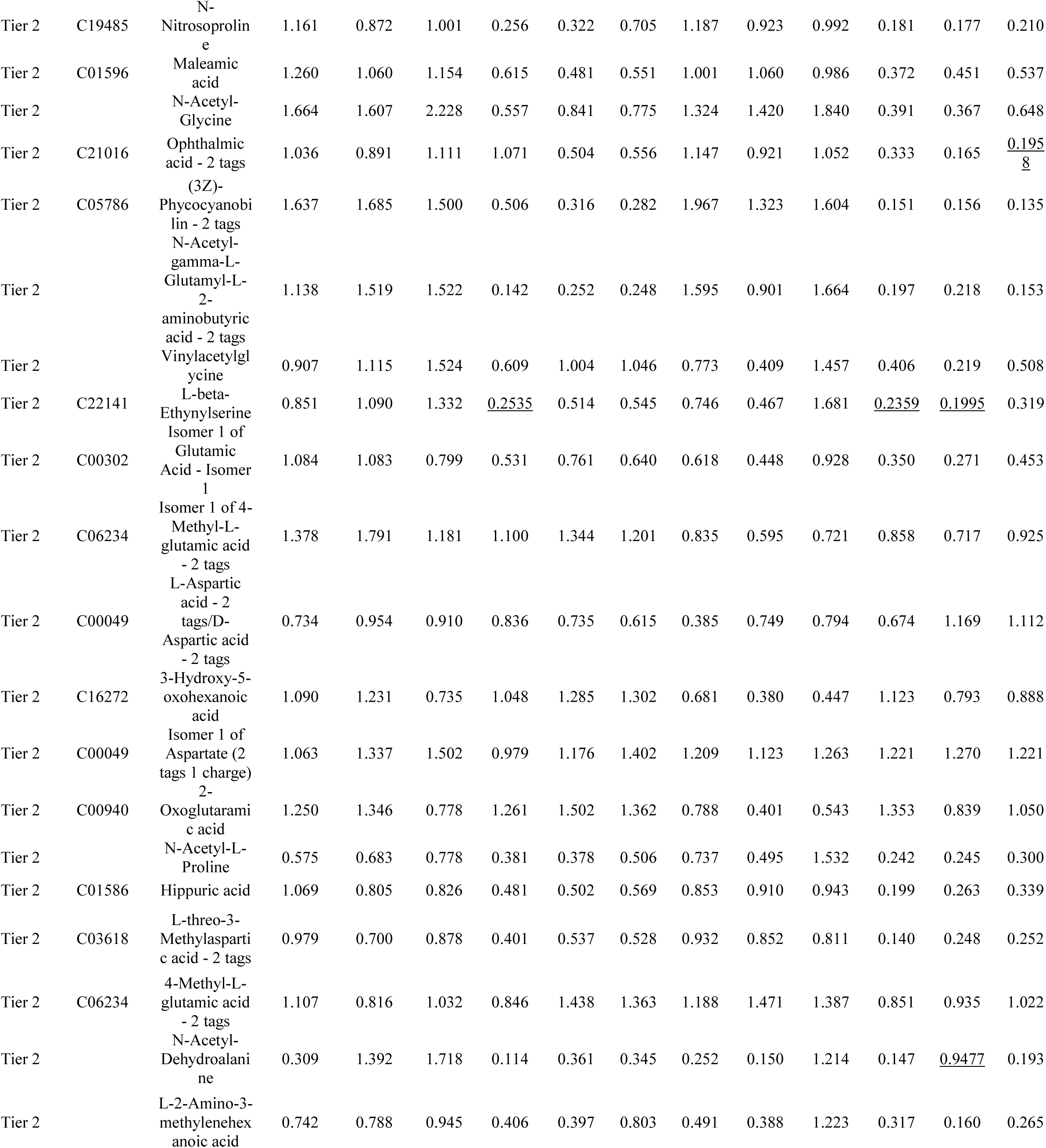
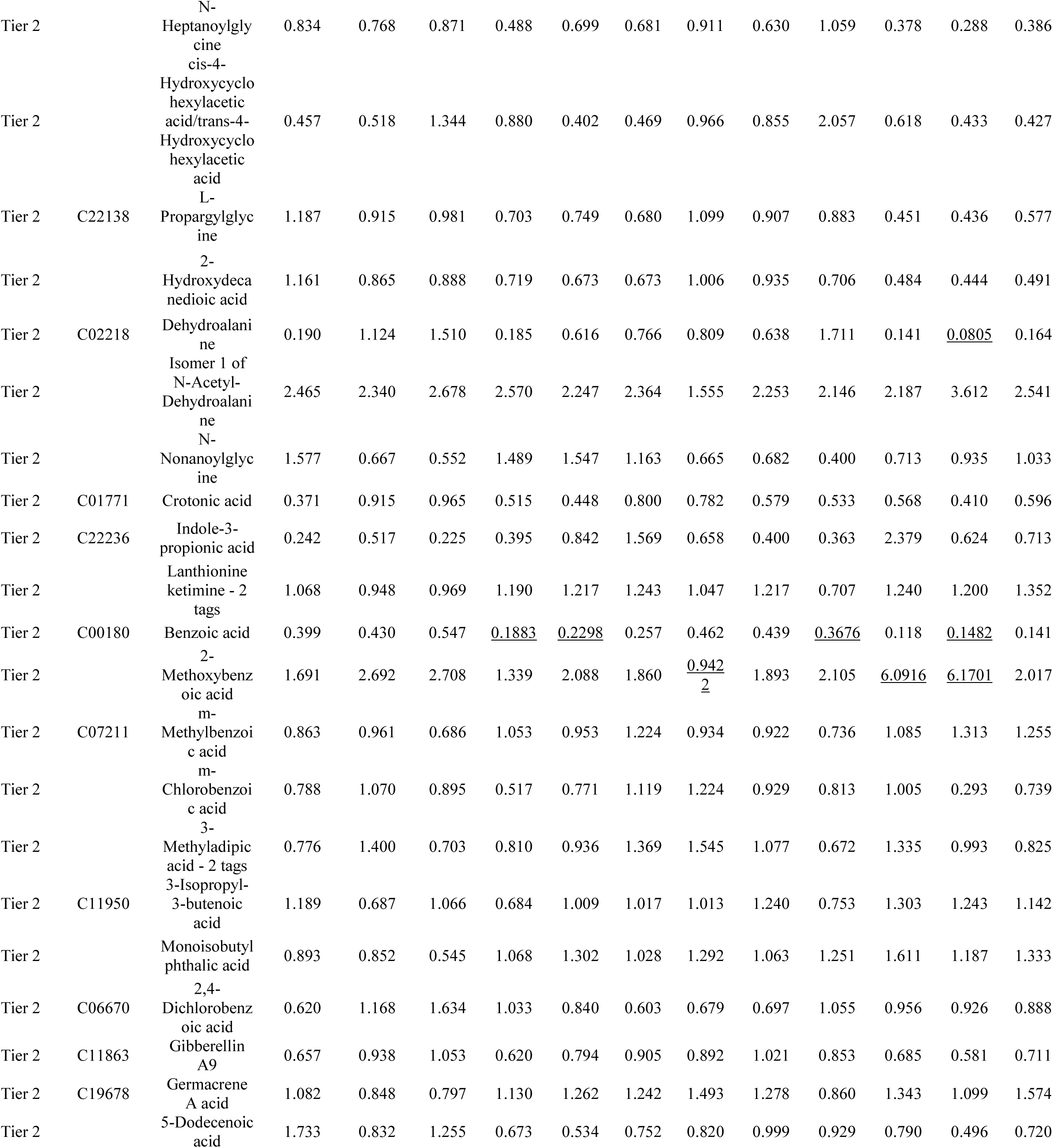
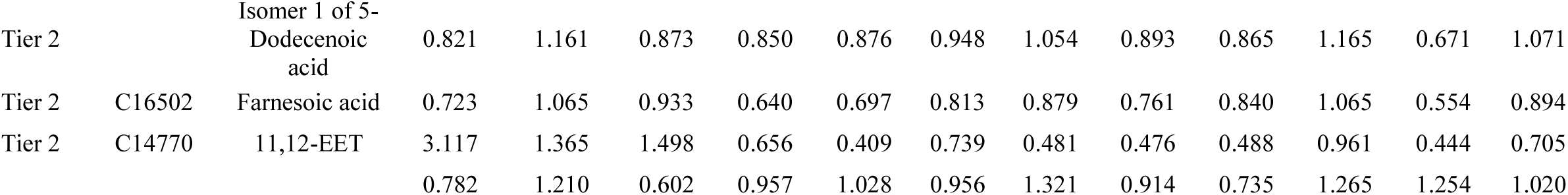

**Supplemental Table 5.**

| Genotype | Metabolite | FC<br>(TCR/Unstim) | log2(FC) | raw.pval | (-)<br>LOG10(p) |
| --- | --- | --- | --- | --- | --- |
| KO | Uridine | 0.16797 | -2.5737 | 0.00173 | 2.7619 |
| KO | Acetic acid | 0.42243 | -1.2432 | 0.035406 | 1.4509 |
| KO | Glyceric acid | 0.4382 | -1.1903 | 0.015953 | 1.7972 |
| KO | Hypoxanthine | 0.4552 | -1.1354 | 1.574e-05 | 4.803 |
| KO | Imidazoleacetic acid | 0.52931 | -0.9178 | 0.040071 | 1.3972 |
| KO | 3-(2-Hydroxyphenyl)propanoic acid | 0.57711 | -0.79308 | 0.002979 | 2.526 |
| KO | L-Propargylglycine | 0.62152 | -0.68613 | 0.002868 | 2.5424 |
| KO | Germacrene A acid | 0.62307 | -0.68252 | 0.027278 | 1.5642 |
| KO | Glycerylphosphorylethanolamine | 0.63593 | -0.65306 | 0.004522 | 2.3447 |
| KO | 1,2-Dihydroxy-3-keto-5-methylthiopentene | 0.66631 | -0.58574 | 0.044568 | 1.351 |
| KO | Malate | 1.5595 | 0.64106 | 0.016142 | 1.7921 |
| KO | Beta-Alanine | 1.5998 | 0.67793 | 0.000632 | 3.1992 |
| KO | 3,5-Dihydroxybenzoic acid | 1.6453 | 0.71833 | 0.004807 | 2.3182 |
| KO | Hydroxyphenyllactic acid | 1.6881 | 0.75539 | 0.022497 | 1.6479 |
| KO | Maleamate | 1.7388 | 0.79807 | 0.015687 | 1.8045 |
| KO | Hypotaurine | 1.7678 | 0.82193 | 0.000283 | 3.5488 |
| KO | gamma-Amino-gamma-cyanobutanoate | 1.8658 | 0.89979 | 0.000149 | 3.8277 |
| KO | L-Threonine | 1.8682 | 0.90165 | 0.009025 | 2.0445 |
| KO | L-Alpha-aminobutyric acid | 1.8916 | 0.91964 | 0.001466 | 2.8339 |
| KO | D-1-Aminopropan-2-ol O-phosphate | 1.9219 | 0.9425 | 0.00035 | 3.4565 |
| KO | L-beta-Ethynylserine | 1.9994 | 0.99958 | 0.019721 | 1.7051 |
| KO | N-Nitrosodiethylamine | 2.0688 | 1.0488 | 0.000928 | 3.0326 |
| KO | L-Proline | 2.2009 | 1.1381 | 1.2518e-05 | 4.9025 |
| KO | Glutaconic acid | 2.2052 | 1.1409 | 0.01044 | 1.9813 |
| KO | 1,3-Diaminopropane | 2.3917 | 1.258 | 0.006246 | 2.2044 |
| KO | L-Homoserine | 2.422 | 1.2762 | 5.1884e-05 | 4.285 |
| KO | Imidazol-5-yl-pyruvate | 2.6359 | 1.3983 | 0.009586 | 2.0184 |
| KO | 3-{{(2E)-4-Amino-4-oxobut-2-enoyl}amino}-L-alanine | 2.7394 | 1.4538 | 0.004614 | 2.3359 |
| KO | L-Asparagine | 2.9055 | 1.5388 | 1.8139e-05 | 4.7414 |
| KO | N-Acetylphosphinothricin | 3.3098 | 1.7267 | 0.003019 | 2.5202 |
| KO | Umbelliferone | 3.4229 | 1.7752 | 0.000453 | 3.344 |
| KO | 3-Pyridylacetic acid | 3.4262 | 1.7766 | 0.0029 | 2.5376 |
| KO | N-Methyl-D-aspartic acid | 3.5231 | 1.8169 | 0.000235 | 3.6282 |
| KO | L-Cysteine | 3.6911 | 1.8841 | 0.01618 | 1.791 |
| KO | Putrescine | 3.7442 | 1.9046 | 0.000186 | 3.7301 |
| KO | L-Alanine | 3.7695 | 1.9144 | 0.000238 | 3.6236 |
| KO | 5-Aminolevulinic acid | 4.1353 | 2.048 | 0.002524 | 2.5979 |
| KO | 4-Hydroxyproline | 4.3388 | 2.1173 | 3.7249e-05 | 4.4289 |
| KO | 4-Chloro-L-lysine | 5.6273 | 2.4924 | 9.8473e-06 | 5.0067 |
| KO | Ophthalmic acid | 5.6933 | 2.5093 | 2.2798e-05 | 4.6421 |
| KO | Sarcosine | 6.7796 | 2.7612 | 0.000256 | 3.592 |
| WT | Uridine | 0.20089 | -2.3156 | 0.000421 | 3.376 |
| WT | D-Aspartic acid | 0.2459 | -2.0238 | 0.010779 | 1.9674 |
| WT | L-Cystine | 0.27618 | -1.8563 | 0.050599 | 1.2959 |
| WT | 2-Keto-glutaramic acid | 0.28106 | -1.831 | 0.001871 | 2.728 |
| WT | 3-Hydroxy-5-oxohexanoate | 0.28413 | -1.8154 | 0.000541 | 3.2672 |
| WT | (Z)-3-Peroxyaminoacrylate | 0.29332 | -1.7695 | 0.00035 | 3.4554 |
| WT | Hypoxanthine | 0.31407 | -1.6709 | 0.002074 | 2.6833 |
| WT | Methylsuccinic acid | 0.36988 | -1.4349 | 0.011397 | 1.9432 |
| WT | 11,12-Epoxyeicosatrienoic acid | 0.37051 | -1.4324 | 0.020113 | 1.6965 |
| WT | 4-Aminobutiraldehyde | 0.37093 | -1.4308 | 0.005079 | 2.2942 |
| WT | m-Methylbenzoate | 0.37874 | -1.4007 | 0.007591 | 2.1197 |
| WT | 4-Hydroxystyrene | 0.39251 | -1.3492 | 0.01063 | 1.9735 |
| WT | 2,3-Dihydroxytoluene | 0.39303 | -1.3473 | 0.004751 | 2.3232 |
| WT | Acetic acid | 0.40527 | -1.303 | 0.044035 | 1.3562 |
| WT | Glyceric acid | 0.40848 | -1.2917 | 0.047896 | 1.3197 |
| WT | 2-Aminoacrylic acid | 0.40992 | -1.2866 | 0.005417 | 2.2663 |
| WT | 1,2-Dihydroxy-3-keto-5-methylthiopentene | 0.41074 | -1.2837 | 0.012075 | 1.9181 |
| WT | 3-(2-Hydroxyphenyl)propanoic acid | 0.42081 | -1.2488 | 0.000779 | 3.1084 |
| WT | 3-Isopropylbut-3-enoic acid | 0.4467 | -1.1626 | 0.010577 | 1.9756 |
| WT | 4-Ethylbenzoic acid | 0.44698 | -1.1617 | 0.003679 | 2.4343 |
| WT | 4-Methyl-L-glutamate | 0.4515 | -1.1472 | 0.00205 | 2.6882 |
| WT | 2,4-Dichlorobenzoate | 0.45301 | -1.1424 | 0.000743 | 3.1288 |
| WT | L-Aspartic acid | 0.4608 | -1.1178 | 0.0262 | 1.5817 |
| WT | 2-Hydroxybutyric acid | 0.46664 | -1.0996 | 0.017976 | 1.7453 |
| WT | 2-Aminobenzoic acid | 0.46679 | -1.0991 | 0.004439 | 2.3527 |
| WT | Threonic acid | 0.47753 | -1.0663 | 0.020052 | 1.6978 |
| WT | Germacrene A acid | 0.49065 | -1.0272 | 0.024678 | 1.6077 |
| WT | Hypotaurine | 2.0302 | 1.0216 | 0.001048 | 2.9796 |
| WT | L-threo-3-Methylaspartate | 2.1181 | 1.0828 | 0.030249 | 1.5193 |
| WT | L-Homoserine | 2.1209 | 1.0847 | 0.00251 | 2.6003 |
| WT | N-Nitrosodiethylamine | 2.1597 | 1.1108 | 0.008924 | 2.0495 |
| WT | D-1-Aminopropan-2-ol O-phosphate | 2.2157 | 1.1478 | 0.000227 | 3.6448 |
| WT | Alanylalanine | 2.2887 | 1.1945 | 0.046521 | 1.3323 |
| WT | 1,3-Diaminopropane | 2.4539 | 1.2951 | 0.028802 | 1.5406 |
| WT | Leucyl-Glutamate | 2.4764 | 1.3083 | 0.000526 | 3.2793 |
| WT | Spermidine | 2.5905 | 1.3732 | 0.00304 | 2.5171 |
| WT | L-Serine | 2.602 | 1.3796 | 0.00282 | 2.5498 |
| WT | L-Threonine | 2.6651 | 1.4142 | 0.000613 | 3.2127 |
| WT | L-Proline | 2.7843 | 1.4773 | 0.000365 | 3.4375 |
| WT | Umbelliferone | 2.7856 | 1.478 | 0.025007 | 1.6019 |
| WT | N-Nitrosoproline | 2.8747 | 1.5234 | 0.004092 | 2.3881 |
| WT | L-Asparagine | 3.0737 | 1.62 | 0.001218 | 2.9143 |
| WT | D-Serine | 3.0916 | 1.6284 | 0.001591 | 2.7984 |
| WT | Imidazol-5-yl-pyruvate | 3.2467 | 1.699 | 0.004773 | 2.3212 |
| WT | 5-Aminolevulinic acid | 3.6188 | 1.8555 | 0.002611 | 2.5831 |
| WT | Putrescine | 3.8125 | 1.9307 | 0.000273 | 3.5646 |
| WT | N-Methyl-D-aspartic acid | 4.0345 | 2.0124 | 0.011655 | 1.9335 |
| WT | 3-{{[(2E)-4-Amino-4-oxobut-2-enoyl]amino}-L-alanine | 4.2044 | 2.0719 | 0.080598 | 1.0937 |
| WT | 4-Hydroxyproline | 4.5693 | 2.192 | 0.001395 | 2.8556 |
| WT | L-Cysteine | 4.6594 | 2.2202 | 0.027113 | 1.5668 |
| WT | N-Acetylphosphinothricin | 4.766 | 2.2528 | 0.029868 | 1.5248 |
| WT | L-Alanine | 5.1529 | 2.3654 | 0.012532 | 1.902 |
| WT | 3-Pyridylacetic acid | 5.1781 | 2.3724 | 0.024599 | 1.6091 |
| WT | 4-Chloro-L-lysine | 5.6512 | 2.4986 | 0.000482 | 3.3168 |
| WT | Ophthalmic acid | 5.7367 | 2.5202 | 0.000111 | 3.953 |
| WT | Sarcosine | 6.4243 | 2.6835 | 0.004039 | 2.3938 |

**Supplemental Table 6.**
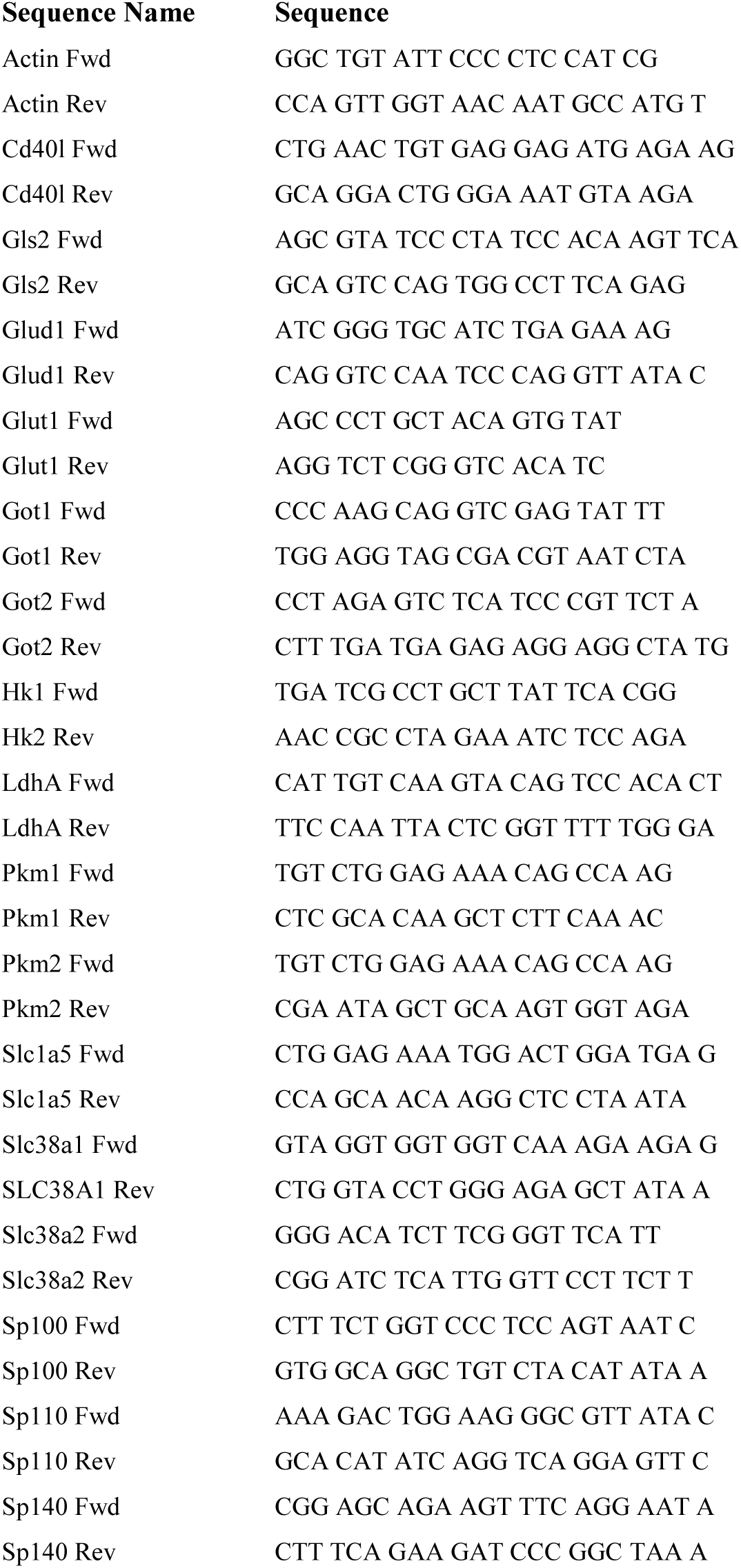

